# Single recipient cell tracking of tellurium-labeled extracellular vesicle proteomes (TeLEV) identifies EV-driven immunomodulation

**DOI:** 10.1101/2025.11.01.685872

**Authors:** Daniel Bachurski, Rahil Gholamipoorfard, Yong-Jia Bu, Patrick Hoelker, Lisa Wessendorf, Hendrik Jestrabek, Luca D. Schreurs, David Stahl, Amin Mokhlesi, Philipp Gödel, Felix Gaedke, Elias Ranjbari, Selen Seyhan, Ulrike Resch, Luisa Marie Schmidt, Tobias Tertel, France Rose, Cláudio Pinheiro, Maribel L. Corona, Anton von Lom, Alexander Frederik vom Stein, Phuong-Hien Nguyen, Katrin S. Reiners, An Hendrix, Guillaume van Niel, Marcus Krüger, Katarzyna Bozek, Lydia Meder, Per Malmberg, Paula Cramer, Barbara Eichhorst, Martin Peifer, Roland T. Ullrich, Astrid Schauss, Christian Pallasch, Paul J. Bröckelmann, Ron D. Jachimowicz, Christian Preußer, Bernd Giebel, Elke Pogge von Strandmann, Mark Nitz, Michael Hallek, Nima Abedpour

## Abstract

Extracellular vesicles (EVs) mediate tumor-immune cell communication by carrying protein cargo that can immediately modulate signaling and antigen presentation. Yet mapping the uptake of primary EV proteomes by human immune cells at single-cell resolution has been constrained by a lack of labeling strategies. We show here that TeLEV, a tellurium-based metabolic mass tagging approach that incorporates L-2-tellurienylalanine (TePhe) into EV proteomes, can produce a biologically rare monoisotopic signal, which is detectable by mass cytometry, imaging mass cytometry, and nanoscale SIMS, without perturbing EV morphology, yield, or proteome composition. We applied TeLEV to label primary malignant B-cell-derived EVs (MBC-EVs) from chronic lymphocytic leukemia (CLL) patients and could follow EV uptake by distinct cell populations of healthy donor peripheral blood mononuclear cells. MBC-EV uptake occurred predominantly in cells of myeloid lineages. In direct control experiments with matched secreted proteins, a machine learning approach identified CD123, CD127, and CD25 as key markers distinguishing primary MBC-EV recipients from matched secreted protein recipient cells. Nanoscale imaging enabled localization of EV-delivered proteins within heterochromatin, whereas Te-labeled secreted proteins accumulated in the cytoplasm of recipient cells. We then generated a pan-immune EV uptake atlas by tracing the uptake of primary and cell-line EVs from nine cell lines and six donors with chronic lymphocytic leukemia into 2,977,094 recipient cells across 43 cell types and subpopulations. We found that the uptake of MBC-EVs by myeloid recipients induced monocyte-derived dendritic-cell polarization characterized by the co-expression of the interleukin-receptor triad (IL-RT: CD123, CD127, CD25) identified above. Time-resolved EV uptake analysis showed a rapid, transient expression of CD123/CD127 followed by CD25, both tightly coupled to MBC-EV uptake by myeloid cells. The intensity of IL-RT expression correlated with that of PD-L1 and BCL-2. Using different STAT degraders to bidirectionally modify the EV-induced STAT5 signal, we observed that MBC-EV uptake and IL-RT, PD-L1, and BCL-2 expression increased with STAT3 degradation and decreased with STAT5 degradation. To investigate the functional consequences of the MBC-EV-induced changes, we showed that MBC-EVs in the presence of IL-2 induced a high-CD25 immune state with low cytotoxic and high B cell proliferation. Taken together, TeLEV represents a novel tool for single-cell tracking of EV proteomes, revealing STAT5-dependent immune remodeling of recipient cells.

## Introduction

Extracellular vesicles (EVs), particularly those derived from tumors, are recognized as potent mediators of intercellular communication, transferring complex protein cargoes that significantly modulate immune responses^1–5^. For instance, PD-L1 facilitates immune escape either through direct expression on melanoma-derived EVs or by induction on immunomodulatory monocytes via malignant B cell-derived EVs (MBC-EVs) in lymphoma^6,7^. Despite their established importance, deciphering the precise functional consequences of EV exchange at a single-cell level remains a major challenge due to methodological limitations^8^. The central hurdle is the inability to reliably track the fate of the primary EV proteome and identify recipient cells at single-cell resolution^9–12^. Current labeling strategies are severely limited, as they often alter EV biology or function, lack specificity, or fail to distinguish EV-encapsulated cargo from contaminating secreted proteins^10,11,13^. This ambiguity prevents researchers from causally linking the delivery of specific EV cargo with phenotypic changes in recipient cells^13–16^. Progress in the field urgently requires a minimally disruptive, proteome-sensitive labeling strategy that includes rigorous internal controls to isolate EV-specific effects and is compatible with high-resolution analysis^11,12,17^.

To address this critical methodological bottleneck, we developed TeLEV (Tellurium-based labeling of the extracellular vesicle proteome), a novel metabolic mass-tagging strategy. TeLEV utilizes isotopically enriched L-2-tellurienylalanine (TePhe)^18^, a phenylalanine mimic that incorporates directly into the nascent EV proteome. We established that this approach is minimally perturbing and non-toxic at working concentrations, preserving EV morphology and composition^18–20^. This generates a biologically rare, monoisotopic tellurium signal that enables sensitive, cross-platform detection via mass cytometry, imaging mass cytometry, and correlative TEM-nanoSIMS^21–23^. Crucially, the TeLEV workflow allows the generation of matched controls—Te-labeled EVs and Te-labeled secreted proteins isolated from the same secretome—providing the necessary controls to precisely attribute recipient cell phenotypes to EV-encapsulated cargo.

We applied TeLEV to investigate how MBC-EVs reshape the human immune landscape. By profiling a total of 8.9 million peripheral blood mononuclear cells exposed to EVs from primary donors and cell lines, we constructed, among other outputs, a comprehensive pan-immune tumor-derived EV uptake atlas, revealing preferential uptake by myeloid populations. By distinguishing EV uptake from matched soluble protein controls, we identified an EV-specific induction of a coordinated interleukin receptor triad (IL-RT; CD25, CD123, CD127) and polarization toward a monocyte-derived dendritic cell (moDC)-like state. Furthermore, nanoscale imaging revealed distinct subcellular fates: internalized EV proteomes localized predominantly to the heterochromatin of recipient cells, whereas secreted proteins accumulated in the cytoplasm.

To causally interrogate the mechanisms driving this EV-dependent remodeling, we integrated TeLEV analysis with selective proteolysis targeting chimera degraders for STAT3 (SD-36)^24,25^ and STAT5 (AK-2292)^26^ allowing us to bidirectionally modify STAT5 signaling. We discovered that MBC-EVs activate STAT5 signaling in myeloid cells, which is necessary for orchestrating the interleukin receptor programs and monocytic polarization. Conversely, STAT3 degradation enhanced both EV uptake and the resulting phenotypes, indicating STAT3 acts as a negative regulator of the STAT5-driven EV response. This study establishes TeLEV as a powerful, high-resolution platform for dissecting EV-mediated communication, resolving long-standing methodological ambiguities in the field, and uncovering a key STAT5-dependent mechanism by which tumor EVs orchestrate immune modulation.

## Main

### Cross-platform recipient cell tracking with matched controls and nanoscale localization

We established TeLEV, a tellurium-based metabolic mass tag in which primary cells or cell lines are cultured with L-2-tellurienylalanine (TePhe), the conditioned medium is fractionated by size-exclusion chromatography, and Te-labeled MBC-EVs (F3–F5) are analyzed alongside matched Te-labeled secreted proteins (F9–F11) from the same secretome (Fig. 1a)^27^. This matched-fractions design allows direct attribution of monoisotopic tellurium to EV-mediated cargo versus freely secreted proteins and enables readout across suspension mass cytometry, imaging mass cytometry, and correlative transmission electron microscopy (TEM) and nanoSIMS (Fig. 1b,c,d). At single-cell resolution, the approach produced robust Te signals in peripheral blood mononuclear cells (PBMCs) exposed to MBC-EVs for 16 h with negligible background in PBS-HAT controls, and it cleanly distinguished uptake of EV cargo from uptake of matched Te-labeled soluble proteins by high-dimensional analysis of recipient cell phenotypes (Fig. 1b and Extended Data Fig. 1a,b,c). Imaging mass cytometry on sister cultures reproduced heterogeneous recipient-cell intensities with the expected per-pixel sensitivity difference relative to suspension mass cytometry (Fig. 1c)^22^. Correlative TEM-nanoSIMS^28^ localized the Te tag derived from Te-labeled primary MBC-EVs predominantly to electron-dense heterochromatin in myeloid recipients. In contrast, the signal from Te-labeled secreted proteins was found to be concentrated in the cytoplasm (Fig. 1d). Together, Fig. 1 demonstrates that TeLEV allows for the sensitive, specific tracking of EV proteomes across diverse technologies, from population-level uptake to nanoscale localization, while distinguishing EV-mediated from non-EV-mediated protein transfer.

**Fig. 1.**
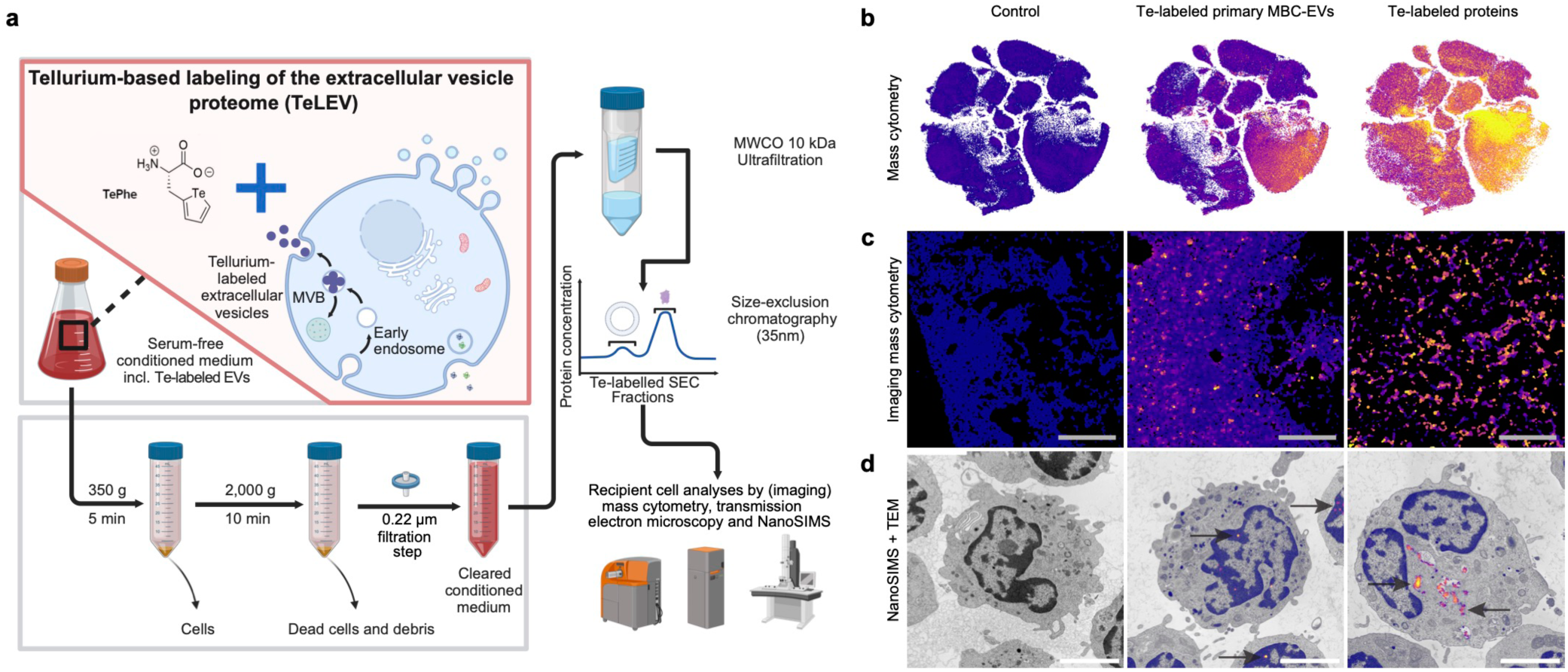
Single recipient cell tracking of tellurium-labeled EV proteomes. a, Schematic of the TeLEV (tellurium-based labeling of the extracellular vesicle proteome) mass-tag labeling workflow yielding Te-labeled MBC-EVs. Primary chronic lymphocytic leukemia cells or cell lines were cultured for 48 h in chemically defined, serum-free medium supplemented with 50 µM L-2-tellurienylalanine (TePhe). Conditioned medium was collected, 0.22 µm filtered, concentrated by 10 kDa ultrafiltration (Amicon 10 kDa RC), and fractionated to separate EVs from soluble secreted proteins by size-exclusion chromatography (35 nm resin; IZON qEV original 2). b, Mass cytometry of PBMC recipient cells showing a two-dimensional t-SNE embedding colored by the TeLEV signal originating from primary CLL donors. Columns show PBS-HAT vehicle control (left), CLL-derived MBC-EVs (middle), and Te-labeled soluble proteins (right). *HAT* denotes HEPES, human albumin, and trehalose. Data were acquired on a Helios mass cytometer. Primary MBC-EVs and secreted proteins shown in Fig. 1 were isolated from 25 mL of cell-conditioned medium (from 250M primary CLL cells over 48 h), resulting in 1.4 x 10^10^ MBC-EVs used per EV condition. c, Imaging mass cytometry of PBMC recipients exposed to PBS–HAT control, CLL-derived MBC-EVs, or Te-labeled secreted proteins. Representative fields are shown (acquired on a Hyperion imaging mass cytometer). Scale bar, 200 µm. d, Correlative ultrastructural and isotopic mapping of PBMC recipients. Transmission electron microscopy (TEM) on 70 nm sections was followed by nanoSIMS of the same regions of the same grids (NanoSIMS 50L; CAMECA, Gennevilliers, France) to detect tellurium enrichment. Arrows indicate TeLEV signal (heat-map overlay) localized predominantly in heterochromatin of myeloid recipients after Te-EV exposure (middle) and in the cytoplasm after exposure to Te-labeled soluble proteins (right). Scale bar, 2 µm.

### TeLEV preserves EV morphology, abundance, and proteome composition

Because labeling approaches involving modifications to EV-secreting cells could potentially distort the generation of primary EVs or their destiny, we investigated the effects of the TeLEV procedure on EV morphology, abundance, and protein composition^29^. Triplex immunogold TEM for CD9, CD63, and CD81 revealed indistinguishable labeling and preserved cup-shaped morphology in primary Te-labeled MBC-EVs and unlabeled controls. In the absence of uranyl acetate contrast, primary Te-labeled MBC-EVs remained visible, consistent with intrinsic tellurium-derived contrast that reveals fine ultrastructural features (Fig. 2a). Moreover, nanoparticle tracking analysis (NTA) of MBC-EVs generated from 10 primary CLL donors showed overlapping size distributions. Fractionation by size-exclusion chromatography in 3 donors produced concordant particle and protein profiles, with MBC-EV-enriched fractions peaking at F3–F5 and soluble proteins enriched in later fractions (Fig. 2b,c)^30^. Pooled MBC-EV fractions (F3–F5) exhibited no difference in particle counts or modal diameters between TeLEV and control preparations (Fig. 2d,e). Marker distribution and global proteomes were likewise unaffected. Immunoblots placed canonical EV markers (CD81, Flotillin-1, Syntenin) in F3–F5 and excluded GM130 from MBC-EV fractions in both conditions, with total protein enriched in late soluble fractions as expected (Fig. 2f). Shotgun proteomics recapitulated these patterns and revealed no systematic shift with TeLEV. For example, samples clustered by donor rather than labeling status in Pearson-correlation matrices and principal-component analysis, global log₂-intensity distributions, and the numbers of identified proteins were comparable between conditions, and differential testing detected no significantly changed proteins at FDR < 0.05 (s₀ > 0.1), including GO-annotated EV proteins (Fig. 2g–m). Together, these data suggested that TeLEV did not measurably perturb MBC-EV morphology, yield, size distribution, or proteome composition across donors, justifying downstream single-cell recipient analyses.

**Fig. 2.**
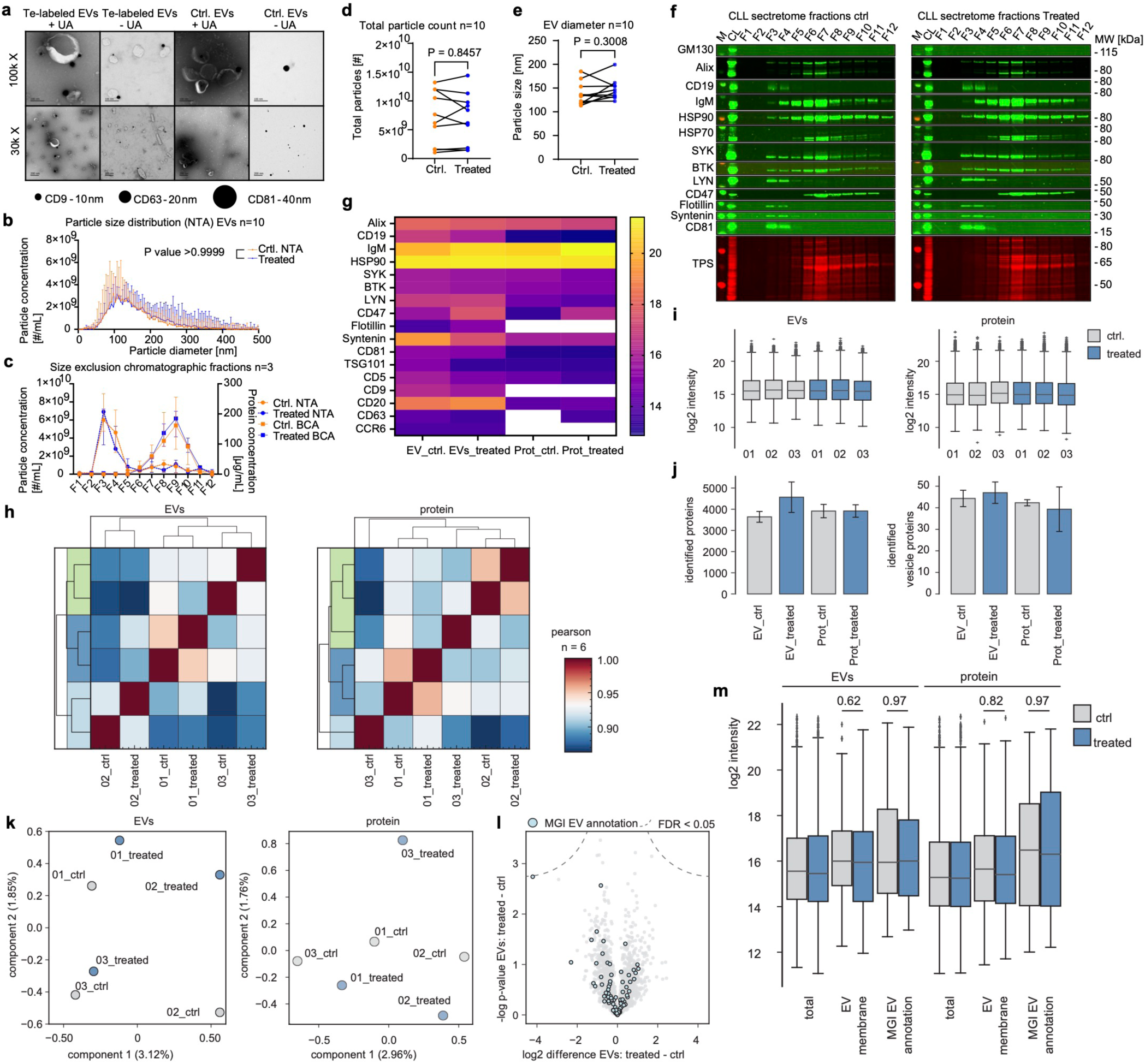
TeLEV does not perturb EV proteomes. a, Triplex immunogold labeling of tetraspanins on MBC-EVs. Primary CLL-derived MBC-EVs and unlabeled control MBC-EVs were stained with anti-tetraspanin antibodies coupled to distinct gold sizes: CD9 (10 nm), CD63 (20 nm), and CD81 (40 nm), with and without uranyl-acetate contrasting. Representative micrographs are shown. Scale bars, 100 nm (100,000x) and 200 nm (30,000x). MBC-EVs and secreted proteins shown in Fig. 2 were isolated from 50 mL of cell-conditioned medium (from 500M primary CLL cells over 48 h). b, Particle sizing of purified primary MBC-EVs. Nanoparticle tracking analysis (NTA) reported particle size distributions. Statistical significance was evaluated using the Kolmogorov-Smirnov test. Data are represented as mean + SEM. c, Size-exclusion chromatography (SEC) fractionation of conditioned medium. 1 mL fractions were collected and analyzed by NTA (particle counts) and BCA (protein). F1–F2, void volume; F3–F5, EV-enriched (MBC-EVs) fractions; F9–F11, secreted-protein fractions. SEC used 35 nm IZON resin following 10 kDa ultrafiltration. Data are represented as mean ± SEM. d, NTA particle counts for pooled primary MBC-EV fractions (F3–F5). No significant changes were identified based on Wilcoxon matched pairs signed rank test (p value = 0.8457). e, NTA particle diameters for pooled primary MBC-EV fractions (F3–F5). No significant changes were identified based on Wilcoxon matched pairs signed rank test (p value = 0.3008). f, Immunoblot analysis of SEC fractions from primary-leukemic donors. Fractions were concentrated and loaded at equal volumes; total-protein staining shows higher abundance in soluble-protein fractions. EV and non-EV markers are shown for TeLEV and control conditions. To avoid overloading of the gel, a 10 µg cut-off of protein fractions was chosen, which was not exceeded by the EV fractions. g, Proteomics comparison of isolated primary MBC-EVs and soluble secreted proteins with and without TeLEV treatment. Heatmap shows log₂ intensity values for selected marker proteins across conditions (n = 3 paired CLL donors). h, Pearson correlation of protein intensities from isolated primary MBC-EVs (left) and secreted proteins (right), showing no clustering based on TeLEV treatment. i, Bar plots of log_2_ intensities from isolated primary MBC-EVs (left) and secreted proteins (right) in control and TeLEV-treated samples. j, Mean number of total identified proteins (left) and EV proteins (right) in control and TeLEV-treated samples of isolated primary MBC-EVs and secreted proteins. Data are represented as mean ± SEM. k, Principal component analysis of isolated primary MBC-EVs (left) and secreted proteins (right), showing no grouping based on TeLEV treatment. l, Volcano plot showing differences of log_2_ intensities between TeLEV-treated and control samples in isolated primary MBC-EVs. No significantly abundant proteins were identified (Student’s t test: FDR < 0.05, s0 > 0.1). GO term-annotated “extracellular vesicle” (GO:1903561) proteins are marked in light blue. m, Boxplot of mean log_2_ intensities for control and TeLEV-treated samples of isolated primary MBC-EVs and secreted proteins based on the GO term “vesicle membrane” (GO:0012506) and MGI GO EV annotation (GO:1903561). Data are represented as mean ± SEM. No significantly changed proteins were identified based on unpaired t-test (*p < 0.05, **p < 0.01, ***p < 0.001).

### Preferential MBC-EV uptake by myeloid cells

We next mapped primary MBC-EV uptake by different immune cells at single-cell resolution after 16 h EV exposure (Supplementary Fig. 1). After data cleaning, 785,260 mass cytometry events were integrated to construct a reference PBMC landscape of primary MBC-EV uptake^23^. Unsupervised FlowSOM clustering followed by manual curation identified 43 immune populations and subpopulations, including rare plasmacytoid DCs (pDCs), myeloid DCs (mDCs), γδ T cells, regulatory T cells, and residual low-density neutrophils (Fig. 3a,b). Within this scope, TeLEV quantification revealed MBC-EV-protein uptake across all major cell lineages. However, TeLEV signal was clearly enhanced in myeloid cells, distributed across classical, intermediate, and non-classical monocytes, residual neutrophils, mDCs, and pDCs, but showed lower levels in all lymphoid subsets (Fig. 3c, left and middle columns)^31^. Cell density distributions showed condition-specific redistributions in myeloid and T-cell compartments relative to vehicle controls, as indicated by shifts in density maxima in the t-SNE space. (Fig. 3c, right column). In comparison, matched Te-labeled soluble proteins from the same secretomes were also preferentially taken up by myeloid cell populations and exhibited similar uptake patterns. Te-labeled proteins also resulted in two separate density peaks in classical monocytes and naïve CD4 T cells. Additionally, Te-labeled soluble protein uptake was overall 4.3 times higher (median Te signal of all cells exposed to primary MBC-EVs 0.5199 vs. matched Te-labeled secreted proteins 2.2486), reflecting the greater abundance of labeled proteins in matched samples (Fig. 2c,f and Fig. 3c, middle column). To characterize primary MBC-EV-specific phenotypes, we trained separate random-forest classifiers on cells in the top quintile of Te signal for MBC-EVs or soluble proteins (each relative to control)^32^. We evaluated the EV-to-protein log-ratio derived from permutation-based feature importance values (Fig. 3d). Interleukin receptor (IL-R) markers dominated this ratio: CD123 (IL3Rα), CD127 (IL7Rα), and CD25 (IL2Rα) (IL-R triad, IL-RT). CD25, in particular, highlighted TeLEV-positive monocytes and mDCs, including IL-RT-positive cells (Extended Data Fig. 2a,d and Fig. 3c). These findings suggested that primary MBC-EV uptake enhanced specifically IL-RT expression in different myeloid recipient cells.

**Fig. 3.**
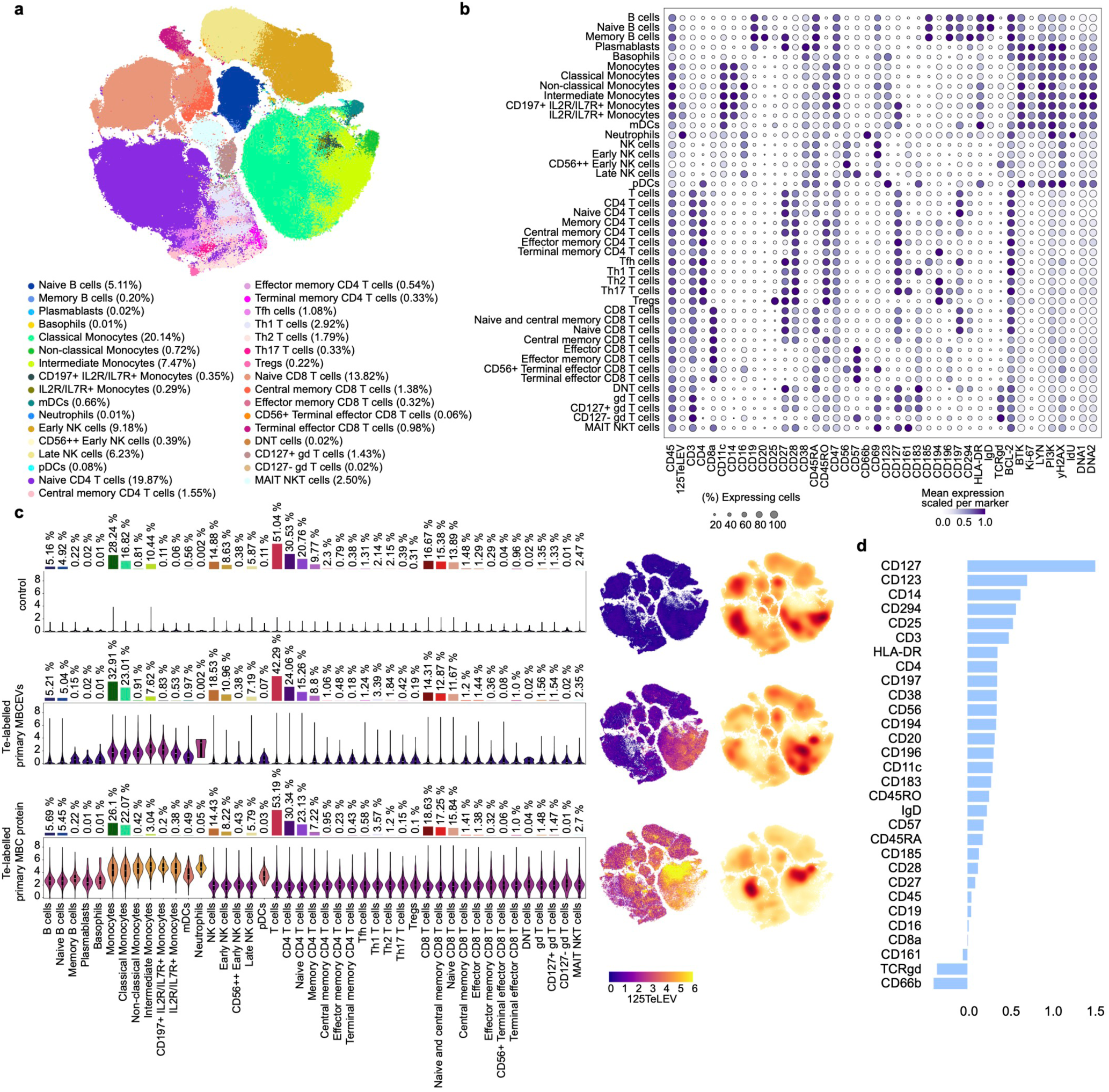
Single-cell mapping of PBMC uptake of Te-labeled EVs and soluble proteins secreted by CLL cells. a, t-SNE projection of PBMC recipient cells (785,260 cells in total) of PBS-HAT control (329,774 cells), primary MBC-EVs (226,157cells), and matched Te-labeled secreted proteins (229,329 cells), colored according to cell type. Numerical annotations indicate the relative proportions of each cell lineage. Primary MBC-EVs and secreted proteins shown in Fig. 3 were isolated from 25 mL of cell-conditioned medium (from 250M primary CLL cells over 48 h), resulting in 1.4 x 10^10^ MBC-EVs used per EV condition. b, Dot plot depicting the expression of selected markers across major cell types and their subtypes from three conditions, PBS-HAT control, primary Te-labeled MBC-EVs, and matched Te-labeled secreted proteins. Dot size corresponds to the proportion of cells expressing each marker within a given cell type, and color intensity indicates the mean expression level. c, Comparison of TeLEV signal distribution across major cell types and their subtypes in each indicated condition, with colors representing the average TeLEV signal level in each population. Bars above the plots show the relative proportions of each population in the corresponding condition (left). White lines denote the median, edges the IQR, and whiskers either 1.5 × IQR or minima/maxima (if no point exceeded 1.5 × IQR; minima/maxima are indicated by the violin plot range). t-SNE projection of cells for each condition showing TeLEV signal (middle). Two-dimensional density distributions of cells in t-SNE embedding space, stratified by conditions (right). d, Feature importance of EV-to-protein log ratios as determined by the random forest classifier, where higher positive values indicate stronger discriminative power in primary MBC-EVs compared to matched Te-labeled secreted proteins.

### MBC-EVs induce an IL-RT and a CD197⁺ moDC-like state in myeloid cells

To create a comprehensive immune recipient atlas of EVs from different sources, we investigated 2,977,094 PBMC recipient cells across sixteen conditions (vehicle plus fifteen EVs from nine cell lines and six primary CLL donors, Supplementary Fig. 2). EV input was normalized to the amount of conditioned medium rather than particle count (reported in Supplementary Table 1) due to inaccuracies, method limitations, and detection thresholds of all current EV quantification methods^11^. A unified t-SNE embedding served as a reference for all comparisons and revealed widespread EV uptake after 16 h (Fig. 4a,b). We modeled cell-type proportions using a Poisson generalized linear mixed model (GLMM) and identified condition-specific frequency shifts in PBMC recipients, mainly affecting the myeloid compartment (Fig. 4c). In PBMC recipients, we observed that certain immune cells, such as IL-R+ and IL-R+/CD197+ monocytes and mDCs, showed an inverse relationship with the frequency of other immune cells, such as non-classical monocytes and neutrophils. This pattern was observed in EV conditions in two of six primary CLL donors (Patients 1 and 4). Additionally, similar cell relationships were found in two cell lines, Ramos (a Burkitt lymphoma) and Su-DHL-4 (a DLBCL line). A high-resolution re-embedding of 276,867 myeloid cells confirmed a recipient cell signature only observed in MBC-EVs but not in malignant non-B cell EVs with enrichment of mDCs, and a CD197⁺ (CCR7+) monocyte subset, suggesting a moDC differentiation induced by MBC-EVs (Fig. 4d,e). These moDC precursors had lower HLA-DR expression than conventional myeloid DCs, reduced CD14 expression, and lacked CD183 (CXCR3). This signature was absent with malignant EVs from non-B cells, but was not elicited by all malignant B cell EVs. Unexpectedly, EVs from the CLL lines OSU-CLL and MEC-1 did not reproduce it, indicating dependence on EV source and patient context (Fig. 4f). Across B cell EV sources, the global responses to Ramos and to CLL patient 1 and 4 were most similar, whereas the two CLL cell lines diverged and lacked the MBC signature (Fig. 4f). Likewise, in the first two principal component space derived from the mean expression of lineage-defining markers across monocytes and mDCs as well as pDCs, the four EV sources exhibiting the MBC signature clustered together, with the highest-variability axis (PC1) separating them from the remaining EV sources (Extended Data Fig. 3a). CD127, CD25, and CD123, followed by CD197, were the strongest contributors to PC1 (Extended Data Fig. 3b,c). Particle number did not account for the signature, as CLL patient 1 and Ramos EV preparations contained fewer particles than all other cell lines tested (Supplementary Table 1). In the shared response pattern, myeloid recipient cells increased levels of CD25, CD123, and CD127 (the IL-RT), with the strongest co-expression observed in Patient 1, Patient 4, and Ramos cells (Fig. 4e–g and Extended Data Fig. 3a,b,c). Some IL-RT-positive monocytes also showed increased CD197 expression, indicating a transition toward a moDC-like state. Ternary plots showed that MBC-EVs increased CD25 expression of myeloid and lymphoid populations, while some non-B-cell EVs (such as those from HEK293T and HS-5) increased CD127 (Fig. 4g). Notably, CD25 induction extended to non-myeloid lineages, including innate-like T cells (MAIT, NKT, and T cells), NK cells, and B cells, revealing cell-type- and source-specific IL-RT signatures (Supplementary Table 2 and Extended Data Fig. 4c). While subsets such as CD127- γδ T cells displayed coordinated upregulation of the IL-RT after 16 h, MAIT/NKT cells and CD56++ early NK cells exhibited an inverse profile—rising CD25 with declining CD127—suggesting divergent temporal or pathway-dependent regulation (Extended Data Fig. 4c). Two controls confirmed that these effects were specific to EVs, as the feature-importance analysis identified CD25 as a distinguishing marker for EVs compared to soluble proteins (Fig. 3d and Extended Data Fig. 2b,c). Unlabeled EVs from Patient 1 produced the same IL-RT pattern (Extended Data Fig. 5).

**Fig. 4.**
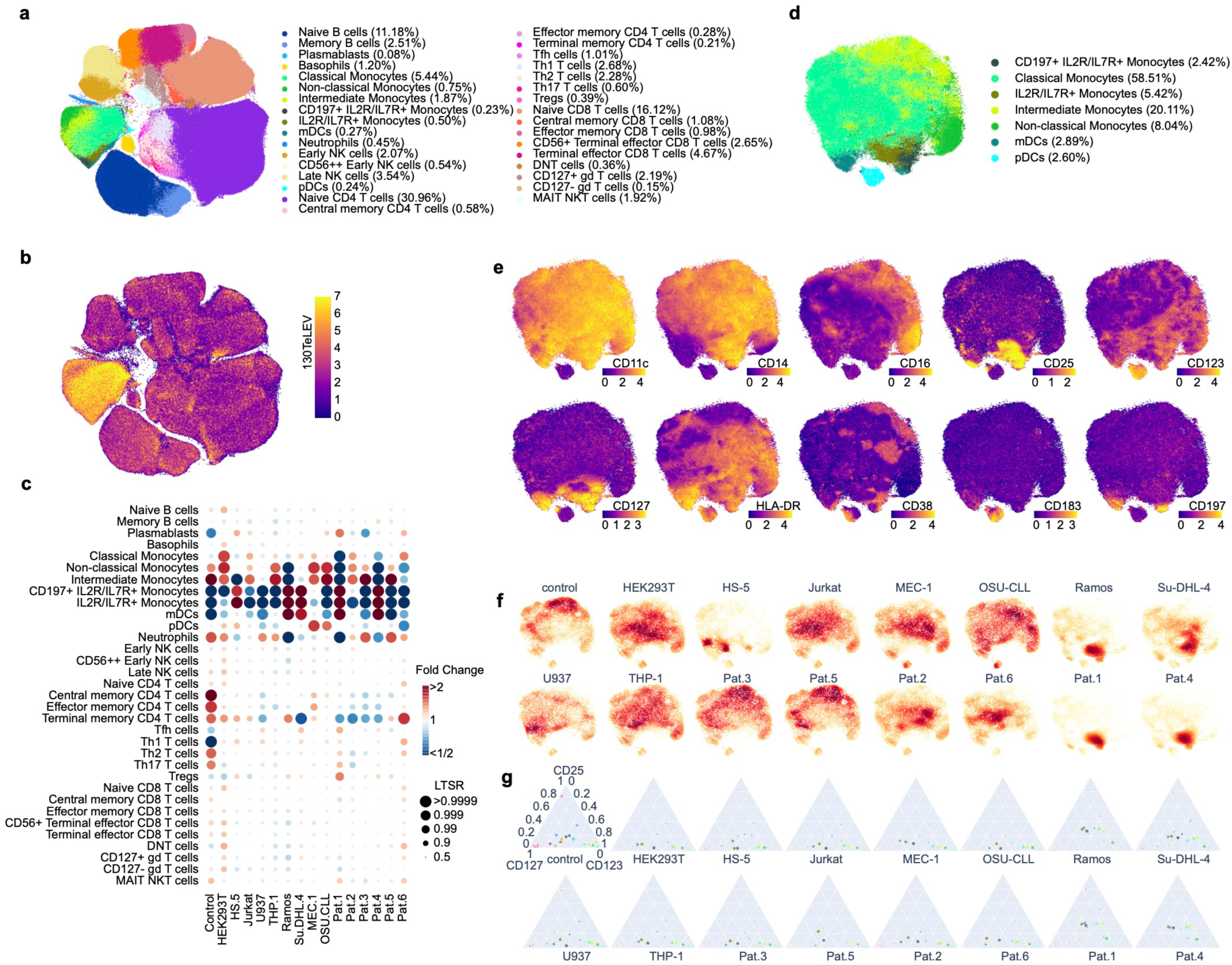
Pan-immune single-cell atlas identifies an IL-RT induced by B-cell malignant EVs. a, t-SNE projection of PBMC recipient cells across 16 conditions (PBS-HAT control plus 15 EV sources, in total 2,977,094 cells), colored according to cell type. Numerical annotations indicate the relative proportions of each cell lineage. EVs shown in Fig. 4 were isolated from 25 mL of cell-conditioned medium (from 250M primary CLL or 25M cells (all cell lines) over 48 h). The EV uptake conditions were normalized according to the input volume of cell-conditioned media (25 mL over 48 h), and EV counts and buffer volumes are reported in Supplementary Table 1. b, t-SNE projection of PBMC recipients across 16 conditions showing TeLEV signal. c, Fold changes in cell type proportions across conditions, estimated using a Poisson generalized linear mixed model that accounts for confounding factors (Methods). Dot size represents the probability of change, measured as the local true sign rate (LTSR). d, t-SNE projection of myeloid cells across 16 conditions (276, 867 cells), colored according to subtypes. e, t-SNE plots showing expression of monocytic and DC lineage markers and interleukin receptors on myeloid cells. f, Two-dimensional density distributions of myeloid cells in t-SNE embedding space, stratified by conditions. g, A cell type–color–coded ternary plot depicting expression levels of CD25, CD123, and CD127. Circle size reflects the average TeLEV signal level within each cell type. Colors correspond to those in the legend shown in (a)

### Biphasic IL-RT coupled to early myeloid uptake in EV uptake kinetics

We then characterized dynamics by exposing healthy-donor PBMCs to primary MBC-EVs across eleven time points (0–48 h) and jointly analyzing 1,512,100 single cells in a common embedding (Fig. 5a). The tellurium signal became detectable by fifteen min, with myeloid cells showing the earliest and strongest uptake (Fig. 5b). Comparing the uptake dynamics of secreted Te-labeled proteins at 2 h to primary Te-labeled MBC-EV uptake, we observed that MBC-EV uptake was more heterogeneous than the relatively uniform protein uptake in the monocyte compartment (Extended Data Fig. 6). Early cell composition profiles were stable for roughly the first 2 h and then progressively rebalanced after ∼4 h, including an increase in CD197⁺ monocytes consistent with the emergence of a moDC-like state (Fig. 5c). By 16 h, uptake patterns converged with those observed in the broader study: monocytes, mDCs, and pDCs dominated primary MBC-EV acquisition, and mDCs/CD197^+^ IL-R^+^ monocytes displayed the highest Te levels (Fig. 5d). Quantifying IL-RT trajectories within myeloid lineages revealed a biphasic program tightly coupled to uptake. In monocytes, CD123 and CD127 rose within 15–30 min, dipped toward baseline by ∼1 hour, and then entered a second phase in which CD127 peaked near 8 h while CD123 continued to increase through 48 h; CD25 exhibited a delayed induction that lagged CD127 by ∼8 h and remained elevated thereafter (Fig. 5e-f). pDCs and mDCs showed analogous but lineage-specific kinetics, with earlier and higher CD123 in pDCs and more sustained CD25 in mDCs (Extended Data Fig. 7a). Taken together, primary MBC-EV uptake induced a transient early CD123/CD127 rise followed by CD25 expression and changes of the immunophenotype of myeloid cells.

**Fig. 5.**
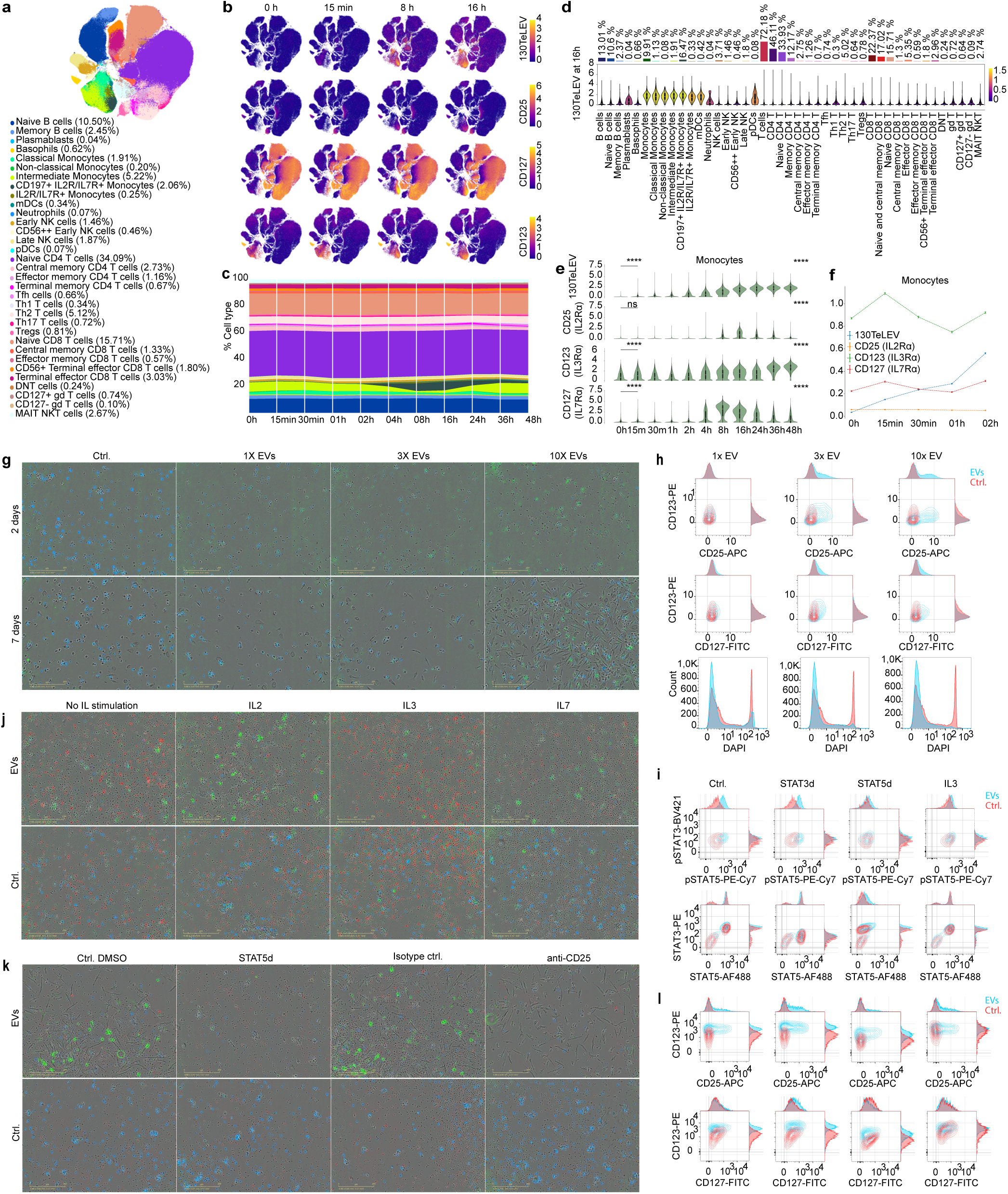
Temporal dynamics and functional consequence of the IL-RT. a, t-SNE projection of PBMC recipient cells across 11 time points (0-48 h, in total 1,512,100 cells) of EV uptake, colored according to cell type. Numerical annotations indicate the relative proportions of each cell lineage. Primary MBC-EVs and secreted proteins shown in Fig. 5a-f were isolated from 4.2 mL of cell-conditioned medium (from 42M primary CLL cells over 48 h), resulting in 2.7 x 10^9^ EVs used per time point condition. b, t-SNE plots showing TeLEV signal, CD25, CD127, and CD123 expression at selected time points (columns). c, Stacked bar plot depicting changes in cell type composition over 48 h. Colors correspond to those in the legend shown in (a). d, Comparison of TeLEV signal distribution across major cell types and their subtypes at 16 h, with colors representing the average TeLEV signal level in each population. Bars above the plots show the relative proportions of each population. White lines denote the median, edges the IQR, and whiskers either 1.5 × IQR or minima/maxima (if no point exceeded 1.5 × IQR; minima/maxima are indicated by the violin plot range). e, Comparison of TeLEV signal, CD25, CD123, and CD127 distributions within the Monocyte population (including Non-classical Monocytes, Intermediate Monocytes, Classical Monocytes, CD25/CD127⁺ Monocytes, and CD197⁺ CD25/CD127⁺ Monocytes) over 48 h. Statistical significance across all time points for each marker was evaluated using the Kruskal-Wallis test. Pairwise comparisons between 00 h and 00 h 15 min were performed using a one-sided Mann–Whitney U test. *p-values* < 0.05 were considered significant and are indicated in the figures as follows: \**p* < 0.05, \*\**p* < 0.01, \*\*\**p* < 0.001, and \*\*\*\**p* < 0.0001. f, Line plots depicting the mean expression levels of TeLEV signal, CD25, CD123, and CD127 within the Monocytes during the first 2 h of EV uptake. Error bars represent mean ± SEM. g, Live-cell immunocytochemistry of purified monocytes exposed to increasing doses of Ramos-derived EVs. Representative frames from day 2 and day 7 are shown. CD25 (Alexa Fluor 488; green) and Cytotox NIR (blue; loss of membrane integrity) are displayed after automated single-cell segmentation (Incucyte built-in). Cells were seeded at 25,000 per well and imaged every 2 h for 7 days using an Incucyte SX5 (20× objective). Scale bar, 200 µm. EVs and secreted proteins shown in Fig. 5g-l were isolated from 1.5 mL of cell-conditioned medium (from 1.5M Ramos cells over 48 h), resulting in 9 x 10^8^ EVs used per EV condition (3X condition, Fig. 5g,h and Extended Data Fig 7b). h, Flow cytometric contour plots of purified monocytes after 16 h of EV exposure across a dose titration (see Fig. legend 5g). CD25 (APC), CD123 (PE), and CD127 (FITC) were measured on a Miltenyi MACSQuant X with compensation using BioLegend compensation beads. DAPI was used as a viability dye for surface staining. i, Intracellular phospho-flow cytometry of purified monocytes incubated with EVs, showing total STAT5 (Alexa Fluor 488), total STAT3 (PE), pSTAT5 (PE-Cy7), and pSTAT3 (BV421). EV exposure was terminated by immediate formaldehyde fixation after 16 h prior to staining. Cleaved Caspase-3 was included as an intracellular apoptosis marker. Acquisition and compensation as in (h). j, Live-cell immunocytochemistry of EV-exposed healthy-donor PBMCs stimulated for 2 days with IL-2 (20 ng mL⁻¹), IL-3 (5 ng mL⁻¹), or IL-7 (25 ng mL⁻¹) in RPMI with serum. Representative images are shown (Incucyte SX5, 20×, 2 h intervals). Scale bar, 200 µm. k, Live-cell immunocytochemistry of EV-exposed healthy-donor PBMCs treated for 7 days with a STAT5 degrader (AK-2292) or anti-CD25 (Basiliximab), compared with DMSO and isotype controls. CD25 (Alexa Fluor 488; green), CD127 (PE; red), and Cytotox NIR (blue) are shown after automated segmentation and background subtraction. Cells were seeded at 50,000 per well and imaged on an Incucyte SX5 (20×, every 2 h). Scale bar, 200 µm. l, Flow-cytometric contour plots of healthy-donor PBMCs after 48 h of EV exposure across a dose titration (1× = 3×10^8^ EVs per 10^5^ cells), showing CD25 (APC), CD123 (PE), and CD127 (FITC). Acquisition and compensation as in (h).

### Interleukin priming links EV exposure to sustained CD25 induction and increased IL-2 responsiveness in myeloid cells

To further explore and validate the functional effects of MBC-EVs on the myeloid compartment, we conducted live-cell immunocytochemistry and (phospho)flow cytometry after exposure to MBC-EVs. To confirm our results on a larger scale and to account for heterogeneity and limited patient material, we chose Ramos-EVs as the MBC cell line model because it most closely resembles Patient 1 EVs in terms of IL-RT expression and moDC induction (Fig. 4). In purified human monocytes, live-cell immunocytochemistry across MBC-EV dose titration showed a dose-dependent increase in CD25; EV-treated monocytes displayed higher viability (low Cytotox NIR signal) and frequent polarized/elongated morphologies, with the highest-CD25 cells typically non-polarized but adjacent to polarized neighbors (Fig. 5g,h and Extended Data Fig. 7b). Early flow cytometry at 16 h confirmed MBC-EV-dependent increases in CD25 and CD127 across MBC-EV titration, with CD123 decreasing at the highest MBC-EV dose (Fig. 5h). The stimulation by various interleukin combinations revealed MBC-EV-dependent priming in CD14^+^ purified monocytes. IL-2 plus IL-7 in the presence of MBC-EVs produced pronounced polarization and maximal CD25 by day 7, whereas IL-2 plus IL-3 or IL-3 plus IL-7 did not produce comparable effects; adding IL-2 + IL-3 + IL-7 increased monocytic polarization and CD25, with stronger responses when combined with EVs (Extended Data Fig. 7c). Consistent with direct contributions from EV cargo to receptor–ligand landscapes, immunogold TEM detected IL-3, CD123, and CD127 on MBC-EVs (Extended Data Fig. 7d). Intracellular phospho-flow cytometry linked receptor remodeling to signaling: EV exposure increased pSTAT5 (MBC-EVs vs. Ctrl one-sided Mann–Whitney U test, p < 0.0001); a STAT3 degrader further enhanced EV-induced pSTAT5 (Stat3d + MBC-EVs vs. EVs one-sided Mann–Whitney U test, p < 0.0001); a STAT5 degrader reduced EV-induced pSTAT5 (Stat5d + MBC-EVs vs. MBC-EVs one-sided Mann–Whitney U test, p < 0.0001); and IL-3 alone partially reproduced the MBC-EV-induced pSTAT5 increase (IL3 vs. Ctrl one-sided Mann–Whitney U test, p < 0.0001 and MBC-EVs vs. IL3 one-sided Mann–Whitney U test, p < 0.0001) (Fig. 5i). In PBMC, MBC-EV exposure induced CD25+ clusters and enhanced IL-2 responsiveness, while IL-only controls showed the expected down-modulation of CD127 by IL-7 and the unexpected up-modulation by IL-3, underscoring that MBC-EVs reorganize IL-R programs and their downstream signaling properties rather than simply mirroring IL-only behavior (Fig. 5j)^33^. These findings align with the time-resolved biphasic IL-RT and help explain why MBC-EVs preferentially drive a high-CD25 state in myeloid recipients.

### STAT5 is required for CD25 induction and polarization and amplifies EV-driven programs

In healthy donor PBMCs imaged for seven days, a STAT5 degrader (AK-2292) abolished MBC-EV-driven monocyte polarization and prevented maximal CD25 induction, whereas anti-CD25 (Basiliximab) had little effect on MBC-EV-induced morphology; neither intervention affected the MBC-EV-associated viability advantage in our in vitro setting (Fig. 5k). In CD14+ purified monocytes, phospho-flow cytometric analysis revealed that STAT3 degradation augmented (STAT3d + MBC-EVs vs. MBC-EVs one-sided Mann–Whitney U test, p < 0.0001) and STAT5 degradation suppressed MBC-EV-induced CD25 (STAT5d + MBC-EVs vs. MBC-EVs one-sided Mann–Whitney U test, p < 0.0001); STAT5 degradation also lowered CD123 under both control (STAT5d vs. Ctrl one-sided Mann–Whitney U test, p<0.0001) and EV conditions (STAT5d + MBC-EVs vs. MBC-EVs one-sided Mann–Whitney U test, p<0.0001). IL-3 increased CD123 and CD127 (IL-3 vs. Ctrl, one-sided Mann–Whitney U test, both p<0.0001), but not CD25 (IL-3 vs. Ctrl, one-sided Mann–Whitney U test, p=1). CD123 and CD127 correlation evident in the controls was diminished in the presence of MBC-EVs (Fig. 5l).

These experiments suggest that STAT5 may act as a signaling node for CD25 induction and MBC-EV-dependent morphological polarization of myeloid recipients.

### STAT signaling modulates the uptake of MBC-EVs and tunes the IL-RT with checkpoint and survival co-regulation

To assess how STAT signaling influences EV responses, TeLEV tracking was combined with pharmacological degradation of STAT3 or STAT5 (± IL-2), and 3,590,520 PBMCs were profiled across sixteen conditions. Cell-type frequencies remained consistent, allowing direct comparison (Fig. 6a,b). STAT3 degradation globally increased Te-labeled MBC-EV uptake across most lineages, whereas STAT5 degradation decreased it (Fig. 6c,d). The uptake differences were not attributable to generalized activation, as CD69 levels were similar across IL-2-lacking conditions—EV alone, STAT3-degrader + MBC-EVs, and STAT5-degrader + MBC-EVs (Extended Data Fig. 8). In myeloid cells, MBC-EVs increased CD25, CD123, and CD127; STAT5 degradation reduced their expression, and STAT3 degradation further amplified EV-driven IL-R expression (Fig. 6e-i). EV-induced CD127 was less affected by STAT5 loss than CD123, and CD123 was reduced by STAT5 degradation even without MBC-EVs. STAT3 degradation alone did not induce the IL-RT program but enhanced the EV-induced response (Fig. 6g–k). MBC-EVs also increased PD-L1 and BCL-2 in myeloid and dendritic cell subsets, monocytes, moDCs, mDCs, and pDCs, with these changes heightened by STAT3 degradation and diminished by STAT5 degradation (Fig. 6j–m). At the PBMC level, IL-2 plus MBC-EVs led to high CD25 and lower Ki-67 in cytotoxic lineages, but higher Ki-67 in B cells, even with elevated CD69 (Fig. 6n–p). STAT3 and STAT5 modulated both EV cargo uptake and interleukin receptor expression, impacting checkpoint and survival markers.

**Fig. 6.**
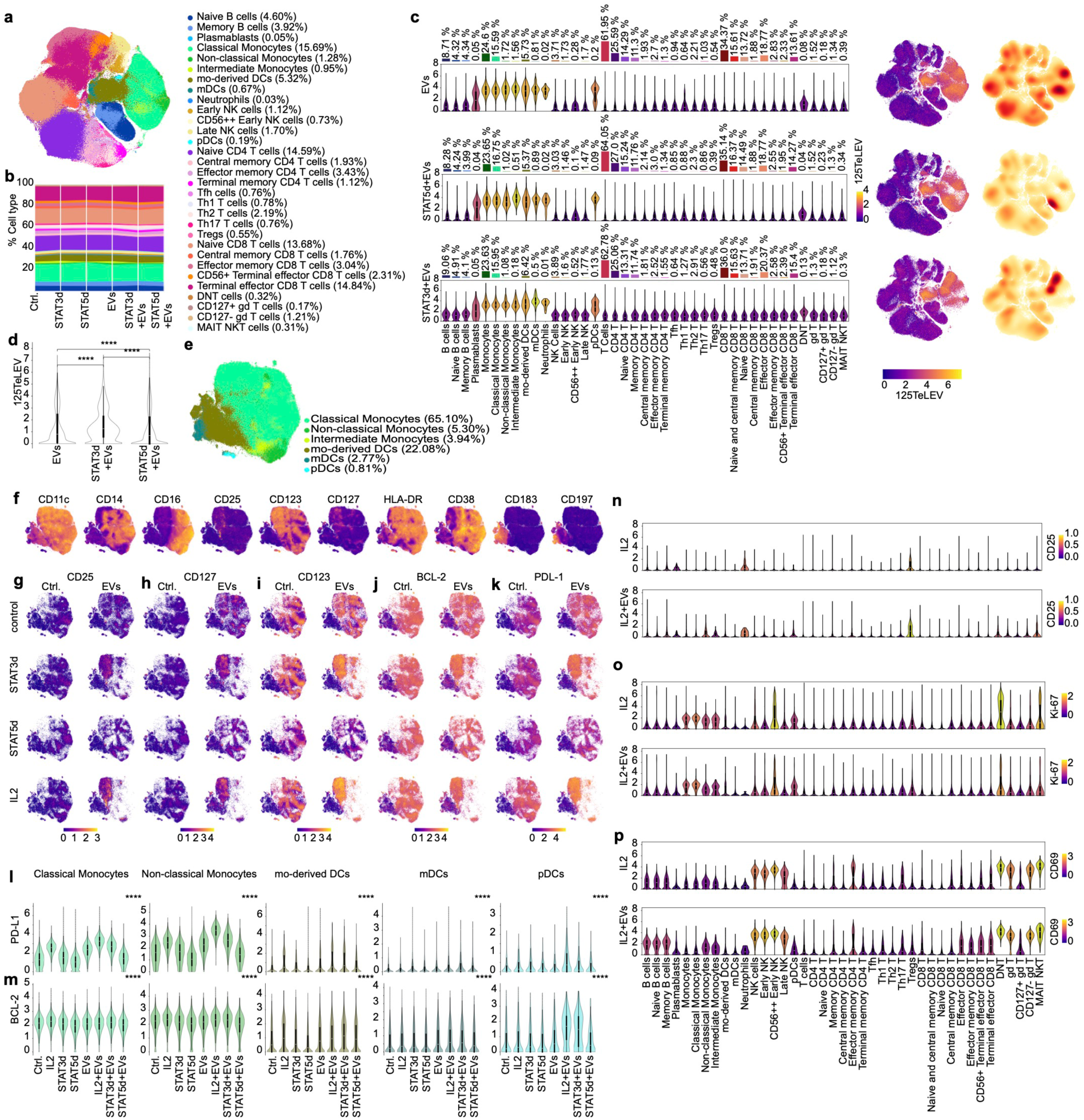
EV-driven immunomodulation by STAT5-regulated IL-RT. a, t-SNE projection of 16 conditions (in total 3,590,520 cells) colored according to cell type. Numerical annotations indicate the relative proportions of each cell lineage. MBC-EVs shown in Fig. 6 were isolated from 8 mL of cell-conditioned medium (from 8M Ramos cells over 48 h), resulting in 5 x 10^9^ EVs used per EV condition. b, Stacked bar plot depicting changes in cell type composition. Colors correspond to those in the legend shown in (a). c, Comparison of TeLEV signal distribution across major cell types and their subtypes. Shown are EVs (217,211 cells), STAT5d + EVs (226,884 cells), and STAT3d + EVs (206,018 cells) conditions, with colors representing the average TeLEV signal level in each population. Bars above the plots show the relative proportions of each population in the corresponding condition (left). White lines denote the median, edges the IQR and whiskers either 1.5 × IQR or minima/maxima (if no point exceeded 1.5 × IQR; minima/maxima are indicated by the violin plot range). t-SNE projection of cells for each condition showing TeLEV signal (middle). Two-dimensional density distributions of cells in t-SNE embedding space, stratified by conditions (right). d, Comparison of TeLEV signal distribution across three conditions shown in (c). Pairwise comparisons were performed using a two-sided Mann–Whitney U test. \*\*\*\**p* < 0.0001. e, t-SNE projection of myeloid population of 16 conditions (865,413 cells), colored according to subtypes. f, t-SNE plots showing expression of monocytic and DC lineage markers and interleukin receptors on myeloid cells. g-k, t-SNE plots showing CD25, CD127, CD123, BCL-2, and PD-L1 expressions across the indicated conditions. l, Comparison of PD-L1 expression distribution across the indicated conditions in Monocytes, moDCs, mDCs, and pDCs. For each cell type, Statistical significance across all conditions was evaluated using the Kruskal-Wallis test.\*\*\*\**p* < 0.0001. m, Same as in (l) for BCL-2. n-p, Comparison of CD25 (n), Ki-67 (o), and CD69 (p) expression distributions across major cell types and their subtypes, with colors representing the average respective marker expression level in each population. Shown are IL-2 (238,611 cells) and IL-2 + EVs (207,300 cells) conditions.

## Discussion

We established TeLEV, a tellurium-based metabolic label that preserved EV morphology and proteome composition. Across a total of 8.9M cells, it traced primary MBC-EV-proteomic transfer into primary single-recipient cells using mass cytometry, imaging mass cytometry, and nanoSIMS^34^. In PBMCs, uptake was myeloid-dominant. MBC-EVs induced a coordinated IL-RT (CD123, CD127, CD25) with kinetics that aligned with EV uptake. Two design elements guided interpretation. First, TeLEV labeled the MBC-EV proteome while generating matched soluble protein fractions as control conditions, allowing attribution of recipient cell phenotypes to EV-encapsulated proteins rather than co-secreted proteins^11,12,35^. Second, labeling did not change EV size distributions, yield, or proteome composition; unlabeled MBC-EVs produced the same recipient-cell patterns.

Across nine cell lines and six primary CLL donors, MBC-EVs reproducibly increased CD123, CD127, and CD25 in monocytes and DCs, with source-dependent differences. Importantly, differences in EV count did not explain these MBC-EV-induced interleukin receptor co-expression. MBC-EVs pushed populations toward higher CD25, whereas few non-B sources favored CD127. A subset of triad-positive monocytes acquired CD197 and features of a moDC-like state with lower HLA-DR and absent CD183. Unlabeled MBC-EVs reproduced the triad pattern seen with Te-labeled MBC-EVs, supporting EV specificity. Time-resolved analyses showed that myeloid cells rapidly take up MBC-EVs, triggering a rapid but transient increase in CD123 and CD127, followed by a later, longer-lasting increase in CD25^36^. This process was observed across various myeloid cell types and depended on EV dose and the surrounding cytokines^37^. The presence of IL-2 and IL-7, together with MBC-EVs, led to the highest CD25 expression and cell polarization in purified monocytes. In agreement with previous reports, MBC-EVs may deliver IL-3, which initiates STAT5 signaling^38–41^. In addition to IL-3, MBC-EVs also expressed CD123 and CD127. Exposure to MBC-EVs boosted pSTAT5, linking receptor changes to signaling activity. Functionally, PBMCs exposed to MBC-EVs and IL-2 became highly activated but did not proliferate as much, especially in cytotoxic cells, while B cells showed increased proliferation. Overall, these results outline a sequence where rapid MBC-EV uptake leads to early receptor priming and later CD25 upregulation, resulting in myeloid cells with enhanced survival and PD-L1 expression.

Our findings also demonstrate that STAT5 signaling is essential for mediating myeloid cell responses to MBC-EVs. Specifically, loss of STAT5 abolishes the capacity of myeloid cells to maximally upregulate CD25 expression and to acquire characteristic monocyte polarization, indicating that STAT5 is indispensable for the terminal stages of the EV-driven response^42^. In contrast, STAT3 degradation enhances EV uptake and amplifies the IL-RT response in myeloid cells, but this effect depends on the presence of MBC-EVs. This observation suggests that STAT3 functions as a negative regulator of the response, effectively acting as a brake on STAT5 activity^25^. Notably, STAT3 degradation further increases pSTAT5 levels following EV exposure, supporting the role of STAT3 in modulating STAT5 signaling during this process. A novel finding of this study is that the recipient cell’s STAT signaling state regulates EV uptake itself, suggesting a feed-forward regulatory loop. A high STAT5/low STAT3 basal state may prime myeloid cells to be more receptive to MBC-EVs, and the EV cargo then further amplifies this state. This is further supported by the reduced CD123 expression observed in the baseline condition exposed to a STAT5 degrader.

Pharmacological targeting of STAT3 (leading to increased pSTAT5) or STAT5 (resulting in decreased pSTAT5) enables fine-tuning of both EV- and IL-RT–driven responses^24–26^. Immune checkpoint and survival markers such as PD-L1 and BCL-2 exhibit parallel regulation: their expression is elevated in response to MBC-EVs, further enhanced by STAT3 loss, and attenuated by STAT5 loss, mirroring the regulatory patterns seen for CD25 and MBC-EV uptake^7,25,43^. In PBMCs, the combination of IL-2 and MBC-EVs increases CD25 expression, reduces proliferation of cytotoxic lineages, and enhances B cell proliferation compared with IL-2 alone. Collectively, these results indicate that the interplay between EVs and the STAT pathway orchestrates recipient cell adaptation, with STAT5 serving as a requisite driver of late CD25 induction and monocyte polarization, and STAT3 acting as a context-dependent suppressor by regulating STAT5 signaling. This regulatory framework is consistently observed for PD-L1 and BCL-2 across myeloid cell subsets.

Our findings have important implications. First, they help clarify why myeloid remodeling and immune evasion often accompany high EV levels in B-cell malignancies: MBC-EVs appear to shift monocytes toward a CD197⁺, PD-L1 high, BCL-2 high phenotype that supports cell migration, survival, and engagement of T-cell checkpoints, while also altering IL-2 signaling^7,44–46^. Second, the comprehensive EV uptake immune atlas developed here supplies quantitative metrics—such as EV uptake, IL-RT marker profiles, and EV-regulated immune profiles—that can be used to assess EV samples or guide experimental designs. Third, the discovery that STAT3 and STAT5 interact to fine-tune cell responses opens the door to experiments combining EV exposure with STAT pathway modulators or defined cytokine mixes to test specific mechanisms, and these strategies can be translated to animal models or patient samples to explore broader applications and therapeutic potential^47–49^. Finally, using a minimally disruptive, proteome-sensitive labeling strategy alongside matched soluble protein controls addresses ongoing uncertainty in the field, making it easier to determine whether observed effects are truly caused by MBC-EVs or by other secreted factors^50,51^.

Taken together, TeLEV represents a breakthrough technology for high-resolution analysis of EV proteomes at the single-cell level. Using this innovative platform revealed that MBC-EVs actively shape myeloid immune landscapes, orchestrating a sequence of receptor changes that promote survival and immune checkpoint readiness. The STAT3/STAT5 signaling axis emerges as a key regulator, with targeted STAT3 degradation modulating STAT5 activity to finely tune immune responses. By combining nanoscale imaging with large-scale cellular profiling and targeted pathway interventions, TeLEV offers a powerful framework for dissecting EV-driven immunomodulation in both healthy and disease contexts. This technology holds promise for guiding future therapeutic strategies and for controlling EV-mediated effects in B-cell malignancies.

## Methods

### Primary leukemic samples

Primary CLL cells and PBMCs isolated from the peripheral blood of CLL patients and healthy donor PBMCs were collected at the University Hospital of Cologne after written and informed consent according to the Declaration of Helsinki and with Institutional Review Board approvals at the University of Cologne, no. 13-091 (BioMaSOTA), no. 19-1559 (Buffy Coats), no 21-1317 (SFB1530) and 21-1472. PBMC isolation and CLL cell purification procedures were performed as previously described^52^. CLL cells were freshly isolated to produce cell-conditioned medium. Clinical information relative to patient samples is listed in Supplementary Table 3.

### Cell lines and cell culture

HS-5, THP-1, Su-DHL-4, OSU-CLL, MEC-1, Ramos, Jurkat, U-937, and HEK293T were cultured in RPMI 1640 (Gibco, #11875093) supplemented with GlutaMAX (Gibco, #61870036), 10% fetal bovine serum (Gibco, #10270106), and 1% penicillin–streptomycin (Gibco, #15140122). All cell lines were tested for Mycoplasma contamination by PCR. All cells were maintained at 37°C in a humidified incubator with 5% CO_2_. In all mass cytometry experiments, 1 x 10^6^ cells were analyzed per condition. For live-cell immunocytochemistry, cultures comprised 25,000 monocytes and 50,000 PBMCs. Flow cytometric analyses utilized a total of 100,000 cells per condition.

### Stimulation and treatment of recipient cells

Primary purified monocytes or PBMCs were stimulated with 20 ng/mL IL-2 (BioLegend, #589102), 5 ng/mL IL-3 (BioLegend, #578002), and 25 ng/mL IL-7 (BioLegend, #581902)^53–55^. STAT3 (SD-36, MedChemExpress, #HY-129602) and STAT5 degraders (AK-2292, MedChemExpress, #HY-148813) were used at 500 nM and compared to DMSO (PanReac AppliChem, #A3672) controls^25,26^. Anti-CD25 antibody (Basiliximab, MedChemExpress, #HY-10885) was used at 10 µg/mL and compared to an isotype control (Invitrogen, #MA5-55090). All stimulation, inhibition, and degradation conditions were simultaneously added with MBC-EVs.

### Healthy donor CD14⁺ monocyte and PBMC isolation

PBMCs were isolated from healthy donor buffy coats. Cells were washed twice in PBS containing 2% FCS and 1 mM EDTA, then counted by flow cytometry. Cell counts were determined by gating on singlets and DAPI-negative events. The necessary number of PBMCs was resuspended to a final concentration of 1 × 10⁸ cells/mL. CD14⁺ monocytes were purified by magnetic bead positive selection (EasySep Human CD14 Positive Selection Kit II, STEMCELL Technologies, #17818) with an EasySep Magnet (STEMCELL Technologies, #18001) using 100 µL of selection cocktail per 1 × 10⁸ cells. Labeled cells were incubated for 6 min at room temperature, magnetically separated, and the CD14- fraction was transferred to a new tube. The remaining CD14^+^ cells were washed twice with 10 mL of buffer, incubated for 3 min each time, and the supernatant was decanted. The previously collected CD14⁻ fraction was again incubated on the magnet for 6 min before collecting the final supernatant. Viability was confirmed by DAPI exclusion using flow cytometry. Purified cells were cryopreserved in FBS containing 10% DMSO, cooled at a controlled rate to –80°C, then transferred to -150°C storage for long-term preservation.

### EV separation

Primary MBC-EVs were isolated from EDTA-anticoagulated peripheral blood collected by venipuncture from patients with chronic lymphocytic leukemia who presented with leukocytosis ≥ 15,000 leukocytes per mL, and samples were processed immediately (Sarstedt, #003.1524). Leukemic B cells were enriched as untouched cells via immunodensity-based negative selection according to the manufacturer’s instructions using the RosetteSep Human B Cell Enrichment Cocktail (STEMCELL Technologies, #15064).

For EV production, the primary cells were adjusted to 1.0 × 10⁷ cells/mL, and the cell lines were adjusted to 1.0 × 10⁶ cells/mL, with viability greater than 95% as determined by DAPI exclusion. The cells were washed twice and cultured in serum-free medium for 48 h. Conditioned supernatants were harvested and clarified by centrifugation at 350 × g for 5 min at room temperature, followed by 2,000 × g for 10 min at 4 °C. The clarified medium was passed through a 0.22 µm PES membrane filter (Filtropur S, Sarstedt, #83.1826.001) after thawing and then concentrated in 10 kDa centrifugal ultrafiltration units (Amicon Ultra-15 10 kDa, Merck Millipore, #UFC901024) to a final volume of 0.5 mL before being applied to a qEVoriginal 35 nm size-exclusion column (qEVoriginal 35 nm Gen2, Izon, #ICO-35). The concentration step was performed at 4 °C.

Fractions were collected in 1 mL increments once the 0.5 mL load had fully entered the column. Fractions F3–F5 were retained as EV-enriched fractions, and fractions F9–F11 were collected as protein controls. Particle concentrations of the EV-enriched fractions were quantified by nanoparticle tracking analysis as described in the dedicated NTA section, and the samples were then reconcentrated. For functional assays, EV preparations were supplemented with in-house-prepared PBS-HAT as previously reported^56^. All EV samples were stored at −80 °C before use.

### TePhe synthesis

125TePhe and 130TePhe (L-2-tellurienylalanine) were synthesized by adapting previously reported methods for the synthesis of TePhe with natural-abundance Te^18,20^. Isotopically enriched Te was sourced from Trace Sciences (Lot 134-1, 92.8% isotopic enrichment for Te-125 and Lot 130Te/9, 99.40% isotopic enrichment for Te-130).

### Tellurium-based metabolic labeling of the extracellular vesicle proteome

Immediately after purification, primary CLL cells or cell lines are seeded for media conditioning (see EV separation). 50 µM of the isotopologues Te-125 or Te-130 L-2-tellurienylalanine (TePhe) are added as medium supplements for metabolic labeling of nascent secretomes.

### EV assay normalization

Current challenges in EV assay normalization stem from the lack of standardized methods for quantifying EVs across different systems^11,12^. Co-isolated secreted proteins often confound protein yields from EV preparations, and preparations deemed ’pure’ typically exhibit very low protein concentrations. Quantification techniques such as nanoparticle tracking analysis (NTA), nanoflow cytometry, and resistance pulse sensing are limited by device-specific detection thresholds, which may exclude smaller EVs (<60 nm) and introduce variability across cell systems^30^. Additionally, TeLEV normalization cannot rely on TePhe incorporation signals, as these are strongly modulated by cellular protein synthesis and secretion rates. To provide a pan-immune EV uptake atlas, we normalized EV input using 25 mL of conditioned medium collected over 48 h. This input corresponds to the secretome of 250 million primary cells (cultured at 1.0 × 10⁷ cells/mL) or 25 million cells for cell lines (cultured at 1.0 × 10⁶ cells/mL), consistent with the distinct seeding densities required for primary versus immortalized cultures and lower EV secretion rates of primary malignant B cells (Supplementary Table 1). Supplementary Table 1 details EV counts from NTA, volumes, and specifies the input material. We acknowledge that EV input normalization remains an unresolved challenge within current methodological constraints.

### Immunoblotting

Cells or MBC-EVs were lysed using RIPA buffer (Cell Signaling Technology, # 9806S). From the cell lysates, 10 µg of protein was loaded per lane. The secretome fractions were adjusted to a cut-off of 10 µg, whereas the EV fractions did not reach this level. The samples were separated on NuPAGE 4-12% Bis-Tris Gel (1.5mm, 15 wells, Invitrogen, # NP0336BOX) under reducing conditions and transferred to a nitrocellulose membrane (Amersham, Protran, # GE10600016). Protein expression was visualized using Revert 700 Total Protein Stain for Western Blot Normalization (LI-COR, # 926-11011). The membranes were blocked for 1 hour at room temperature with Intercept (TBS) Blocking Buffer (LI-COR, #927-60001) and then incubated overnight at 4 °C with unlabeled primary antibodies against GM130, Alix, CD19, IgM, HSP90, HSP70, Syntenin-1, and CD81. After washing, the blots were incubated with an IRDye 800CW Donkey anti-Rabbit IgG secondary antibody (1:15,000, LI-COR, #926-32213) for 1 hour at room temperature and imaged with a LI-COR Odyssey CLx Imager (Lincoln, Nebraska, USA). To allow re-probing with additional primary antibodies, membranes were stripped using NewBlot Nitro Stripping Buffer (5X, LI-COR, #928-40030) according to the manufacturer’s instructions and subsequently re-incubated with alternative primary antibodies.

### Nanoparticle tracking analysis

Measurements were performed on a ZetaView PMX-120-12F-R5 (Particle Metrix, Germany) equipped with a 488 nm, 40 mW laser and a CMOS camera. Samples were diluted in PBS to a final volume of 1 mL. Optimal concentrations were determined by pre-testing to achieve 100-300 particles per frame. Each measurement involved one cycle, during which 11 cell positions were scanned at a frame rate of 30 Hz. Instrument settings were: autofocus enabled, camera sensitivity 80.0, shutter 100, and cell temperature 25 °C. Videos were analyzed using ZetaView Software v8.06.01 SP1 with analysis thresholds of maximum particle size 1000 nm, minimum size 10 nm, and minimum brightness 20.

### Immunogold transmission electron microscopy

Formvar-coated copper grids (Science Services, Munich) were prepared with 5 µl of undiluted purified EV samples. The grids were incubated with the samples for 20 min, blocked for 30 min (Aurion, #905.001), and washed three times for 2 min in PBS containing 0.1% BSA-c (Aurion, #900.099), pH 7.4. Three gold-coupled antibodies or primary unlabeled antibodies (Supplementary Table 4) were diluted 1:25 in PBS with 0.1% BSA-c: CD9, 10 nm (ABCAM, #STN-273562), CD63, 20 nm (ABCAM, #STN-273564), and CD81, 40 nm (ABCAM, #AB273949) combined in one mix. Grids were incubated in drops of antibody solution for 60 min, washed 6 times for 5 min in drops of PBS with 0.1% BSA-c, followed by 3 2-minute washes in PBS. Unlabeled antibodies were detected with a Protein G Immunogold detection step at 1:20 dilution (Aurion, #810.133). After washing, grids were fixed for 5 min with 1% Glutaraldehyde (Sigma, #G5882-100ML) in PBS and washed three times for 2 min in ddH_2_O. Grids were stained for 4 min with 1.5% Uranyl acetate (Agar Scientific, #R1260A) and blotted dry. For each sample, an unstained duplicate was prepared. Images were captured with a Gatan OneView 4K camera on a JEM-2100Plus (Jeol) at 200 kV and analyzed with ImageJ (NIH).

### Cytospin and Imaging Mass Cytometry

Fully stained cells (see “Mass cytometric surface and intracellular protein detection” and Fig. 1b) were subjected to Cytospin on a Cytospin 4 centrifuge (Epredia, #A78300002) using Shandon EZ Single Cytofunnel assemblies (Epredia, #11972345) and Superfrost Plus glass slides (Epredia, #J1800AMNZ). Optimal cell density was determined by titration to 0.5 × 10^6^ cells in 0.5 mL PBS, yielding a single cell layer on the deposition area. After exposure to EVs, the PBMCs were washed and resuspended in PBS, then immediately loaded into the cytofunnel assemblies according to the manufacturer’s instructions. Assemblies were mounted with Superfrost Plus slides and centrifuged for 7 min at 800 rpm. Slides were removed and allowed to air dry at room temperature before further processing. IMC of Cytospin preparations was performed on a Hyperion imaging mass cytometer (Standard BioTools), ablating 750 µm x 750 µm representative regions of interest with a laser power of 3 (laser frequency 200 Hz).

The imaging mass cytometric images were converted from the mcd format to tiff files, and hot pixels were removed using a previously reported method^57^. We used the pretrained mesmer2 model from DeepCell to segment the cells^58^. Namely, images of the nuclear marker DNA and the cytoplasmic marker CD45 were combined into an RGB image (red = DNA, green = CD45, blue = none) and cut into 512 x 512-pixel patches with a 256-pixel overlap. The Mesmer2 model is called for each patch, yielding nuclear and whole-cell probability maps. The probability maps from all the patches are merged using averaging, and a watershed algorithm is then applied, as in the original DeepCell code. For each detected cell (whole-cell segmentation map), the averaged signal intensity for the marker 125TeLEV is computed and arcsinh-corrected with cofactor 5. For display, aggregates were removed using morphological operations. After otsu thresholding, the connected components are processed with a closing operation with a disk element of radius 1 and with an opening operation with a disk element of radius 2. If the connected component has a radius over 150 pixels, it is considered an aggregate and removed from the detections.

### Mass spectrometry sample preparation

Secreted proteins and EV samples collected in PBS were diluted with 1:1 (v:v) 8% SDS in PBS, homogenized by heating for 10 min at 95 °C, and sonicated with a Bioraptor sonicator; the sonication settings were 21 °C water with 10 cycles of 30 on and 30 off. 20 µg protein extract was reduced and alkylated for 10 min at 70 °C using 5 mM TCEP and 15 mM CAA. Protein digestion was performed following the SP3 protocol^59^. In brief, 20 µg of both hydrophobic and hydrophilic beads were added to the sample, and the beads were bound by adding 1:1 volume of acetonitrile (ACN). After 8 min of incubation time, magnetic beads were immobilized and washed 2x with 70% ethanol and acetonitrile. Proteins were digested with trypsin (substrate:enzyme ratio 100:1) and LysC (substrate:enzyme ratio 200:1) overnight at RT. Samples were acidified to 1% formic acid (FA), then cleaned up using house-made SDB-RPS tips. Desalted peptides were loaded onto Evotips, and separation was performed on an Evosep ONE LC system (Evosep, Denmark) equipped with 8 cm PepSep columns (EvoSep, Denmark) at 45 °C with a gradient length of 30SPD (44 min). Mobile phases were composed of 0.1% FA as solvent A and 0.1% FA in ACN as solvent B. The LC system was coupled to a timsTOF pro 2 using a CaptiveSpray source (both Bruker, Germany). Samples were measured in dia-PASEF mode, with ion mobility calibrated using three ions from the Agilent ESI-Low Tuning Mix, following vendor specifications. The dia-PASEF window ranged in dimension 1/k0 0.7 - 1.45, with 16x2 Th windows and in dimension m/z from 350 to 1250.

### LC-MS/MS sample analysis

The mass spectrometry proteomics data have been deposited to the ProteomeXchange Consortium via the PRIDE partner repository with the dataset identifier PXD069581^60^. Acquired spectra were analyzed with DIA-NN (V1.9.1) using library-free search against UniProt Homo Sapiens database (Apr. 2024)^61^. Mass ranges were set according to the settings of the mass spectrometer. Data were further processed using R (v. 4.2.2) with the libraries tidyverse, diann, and data.table, magrittr, FactoMineR, factoextra, and ggplot2, gprofiler, ggplot2. Data input was filtered for unique peptides, q-values < 0.01, and Lib.Q.Value <0.01, PG.Q.Value < 0.01, Global.Q.Value < 0.01, Quantity.Quality > 0.7, Fragment.count >= 4.

Further analysis was performed in Perseus (V 1.6.5.0) and InstantClue (V 0.10.10.20211105). Data were filtered to ensure 100% data completeness. Unpaired Student T-tests were performed on s0 = 0.1, FDR < 0.05, and 500 randomizations. Identified GO terms and MGI extracellular vesicle annotation were imported and compared for each group with an unpaired t-test.

### Correlative transmission electron microscopy and NanoSIMS analysis of EV recipients

PBMCs were centrifuged at 350 xg and fixed for 1 h in 2% Glutaraldehyde with 2.5 % Sucrose (Roth, #4621.1) and 3mM CaCl2 (Sigma, #C7902-500G) in 0.1M HEPES buffer (Gibco, #15630080). The supernatant was removed, and the pellets were mixed with 3% low-melting agarose in 0.2M HEPES buffer, incubated for 10 min at 37°C, and hardened at 4 °C for 30 min. Agarose blocks were cut into pieces of 1 mm³, washed four times for 15min with 0.1M HEPES buffer, and incubated with 1% Osmium tetroxide (Science Services, #E19190) and 1% Potassium hexacyanoferrate (Sigma, #P8131) for 1 h at 4 °C. After 3x5 min washes with 0.1 M Cacodylate buffer (Applichem, #A2140,0100), samples were dehydrated at 4 °C using an ascending ethanol series (50%, 70%, 90%, 3x100%) for 15 min each. The sample was incubated in a mixture of 50% ethanol/propylene oxide, then twice in pure propylene oxide for 15 min each. Infiltration was performed with a mixture of 50% Epon/propylenoxide and 75% Epon/propylenoxide for 2h each step and with pure Epon (Sigma, #45359-1EA-F) overnight at 4 °C. Samples were embedded into TAAB capsules (Agar Scientific, #G3744) and cured for 48 h at 60°C. Ultrathin sections of 70 nm were cut using an ultramicrotome (Leica Microsystems, UC6) and a diamond knife (Diatome, Biel, Switzerland) and picked up using 200 mesh finder Grids without film (Plano, #7FGC200) Sections were stained with 1.5 % uranyl acetate (Agar Scientific, #R1260A) for 15 min at 37°C and 3% Reynolds lead citrate solution made from Lead (II) nitrate (Roth, #HN32.1) and tri-Sodium citrate dihydrate (Roth #4088.3) for 4 min. Images were acquired using a JEM-2100 Plus Transmission Electron Microscope (JEOL) operating at 80kV equipped with a OneView 4K camera (Gatan). Mosaics of regions of interest were acquired using SerialEM ^62^. Location of the cell of interest was noted using the finder pattern. After TEM imaging, sections were transferred onto small 5x5 mm silicon wafer pieces (Plano, #G3390). For this, a small drop of water was added to the wafer, and the Grid was placed on the drop with the section facing the wafer. After the water had evaporated, the grid was removed. Wafers were shipped by placing them onto double-sided sticky tape (Tesa, #05338) inside a Petri dish. Images of the same cells were acquired using nanoSIMS. Registration of images obtained by electron microscopy and nanoSIMS was done using nuclei as fiducials with the plugin EC-CLEM from the software ICY^63^.

Analysis was performed at NanoSIMS Sweden, University of Gothenburg, using a NanoSIMS 50L (CAMECA, Gennevilliers, France). Samples were gold-coated, and a 16 keV Cs^+^ primary ion beam of 0.2 pA (D1-5) was used to raster the sample surface. The detectors were tuned to measure 12C2-, 12C14N-, and 130Te- secondary ions. Before imaging, the areas of interest were implanted to achieve a fluence of 5×10^5^ Cs^+^.cm^−2^ to enhance the ionization yield. Images were acquired on a field size of 10×10 μm^2^ with an image resolution of 512×512 pixels and a dwell time of 3 ms.pixel^−1^. 50 planes were accumulated for each sample. NanoSIMS images were visualized and analyzed with the WinImage software (CAMECA, Gennevilliers, France). Sequential image planes were collected, drift corrected, and accumulated to increase ion counts.

### Incucyte live cell immunocytochemistry

Live-cell immunocytochemistry on an Incucyte system was performed to quantify surface expression of CD25 and CD127 on primary human cells under standard incubator conditions, with a near-infrared (NIR) cytotoxicity readout. PBMCs were plated at 50,000 cells per well, and purified monocytes were plated at 25,000 cells per well in 100 µL of 0.22 µm-filtered, complete medium in Falcon 96-well black-wall, clear-bottom, tissue culture-treated imaging microplates (Corning Falcon, #353219). Plates were incubated for 1 h at 37 °C and 5% CO₂ before kinetic imaging was initiated. Imaging was conducted on an Incucyte instrument equipped with green, red, and NIR channels using a 20× objective. Phase-contrast and fluorescence images were acquired every 2 h for 48 – 7 days from four non-overlapping fields per well with hardware autofocus enabled. Alexa Fluor 488 was detected in the green channel, phycoerythrin (PE) in the red channel, and Cytotox NIR in the NIR channel. Exposure times were kept within the linear range across the time course using the vendor’s recommended presets and were held constant within experiments.

For live-cell immunocytochemistry, non-IL-2/IL-7-blocking anti-CD25-Alexa Fluor 488 and anti-CD127-PE were added at 1 µg mL⁻¹ each directly to cultures (Supplementary Table 4). Incucyte Cytotox NIR was added at a final concentration of 0.6 µM to report membrane compromise. Image analysis was performed in Incucyte software. Total cell masks were generated from phase-contrast images to estimate object counts, and fluorescence masks were generated in the green and red channels to calculate the fraction of CD25⁺ and CD127⁺ cells and mean integrated fluorescence per cell at each time point. Local background subtraction was applied for visualization.

### Flow cytometry (surface and intracellular phospho-staining)

For flow cytometric analysis, cells were cultured for 16 h or 2 days in V-bottom 96-well plates (100 µL/well). Cells were first incubated with 0.5 µL TrueStain FcX (BioLegend, #422302) per well, for 15 min at 4 °C. Subsequently, surface staining was performed using a pre-titrated antibody cocktail (see Supplementary Table 4) for 1 h at 4 °C in the dark. Following incubation, cells were washed twice with Cell Staining Buffer (CSB, BioLegend, #420201) and immediately processed for acquisition.

For intracellular flow cytometric analysis of phosphoproteins, cells were fixed by adding an equal volume of Fixation Buffer (BioLegend, #420801) directly to the culture medium and incubating for 15 min at 37°C. Fixed cells were washed three times in PBS by centrifugation at 900 × g for 2 min to remove residual fixative. Fixed cells were surface-stained as described above, then permeabilized for intracellular staining.

Permeabilization was performed using True-Phos Perm Buffer (BioLegend, #763344) by resuspending cells in 200 µL per well and incubating for at least 1 h at -20°C. Cells were then washed four times with CSB (900 × g, 2 min) and subjected to FcR blocking (as above), followed by overnight staining with an intracellular antibody cocktail at 4 °C.

After staining, cells were washed twice with CSB and resuspended in the same buffer for flow cytometric acquisition. Data were collected on a Miltenyi MACS Quant X and analyzed using FlowJo software (version 10.10, BD Biosciences).

### Antibodies

Antibodies for mass cytometry were either purchased pre-conjugated with heavy metals (Standard BioTools) or conjugated in-house using the Maxpar X8 chelating polymer kit (Standard BioTools) or the Maxpar MCP9 polymer kit (Standard BioTools) according to the manufacturer’s instructions. Details on antibody clones, corresponding heavy metal tags, and suppliers are provided in Supplementary Table 4.

### Mass cytometric surface and intracellular protein detection

Cells were exposed to EVs in 2 mL deep-well 96-well plates (Eppendorf, #0030502310). After EV incubation, the samples were washed twice before TrueStain FcX Fc-receptor block was added for 15 min at 4 °C. For the surface staining, the antibody master mix was added to each well, and the cells were incubated for 45 min at 4 °C with intermittent gentle vortexing. They were then washed twice in CSB at 350 × g for 5 min at 4 °C (BioLegend, #420201). For intracellular antigen detection, the cells were fixed and permeabilized for 45 min at 4 °C using the FoxP3 fixation/permeabilization buffer set (Invitrogen eBioscience, #00-5523-00), then washed twice in the corresponding permeabilization buffer at 900 × g for 5 min at 4 °C. Fc-receptor block was reapplied for 15 min at 4 °C. The intracellular or cytoplasmic antibody master mixes were then added, and the samples were incubated for 45 min at 4 °C with intermittent gentle vortexing, followed by two washes in permeabilization buffer at 900 × g for 5 min at 4 °C. The cells were fixed in 1.6% methanol-free paraformaldehyde for 10 min at room temperature (ThermoFisher, #28906), then washed and incubated overnight at 4 °C with MaxPar Intercalator-Ir 191/193 at a final concentration of 125 nM prepared in MaxPar fix/perm buffer (Standard BioTools, Intercalator #201192B, Fix and Perm #201067). On the following day, the samples were washed twice in CSB and prepared for acquisition.

### Mass cytometric data acquisition

Data were acquired on a Helios mass cytometer (Standard BioTools) with daily instrument quality control and tuning. Manual gating using FCSExpress7 (DeNovo Software) was applied to identify live single cells in the sample convolute based on 140Ce, event length, center, width, DNA (191Ir and 193Ir), and live/dead (103Rh) channels.

### Mass cytometry data preprocessing

Signal intensity measurements in mass cytometry (CyTOF) are often influenced by neighboring channels, leading to spillover effects due to inherent technological limitations. These interferences, known as spillover effects, can significantly reduce the accuracy of cell-type identification. To compensate for spillover effects, we use CytoSpill ^64^, a tool that quantifies and corrects spillover in CyTOF data without requiring single-stained controls. We used the control condition dataset to estimate the spillover matrix with the R package *CytoSpill* (v1.0.1), which was then applied to compensate for the other datasets. Following spillover correction, the single-cell data undergo an inverse hyperbolic sine transformation with a co-factor of 5 (arcsinh(x/5)).

### FlowSOM clustering of mass cytometry data and cell type annotation

Preprocessed mass cytometry data from multiple experiments were integrated for clustering. The lineage-defining markers 89Y-CD45, 141Pr-CD196, 143Nd-CD123, 144Nd-CD19, 145Nd-CD4, 146Nd-CD8a, 147Sm-CD11c, 148Nd-CD16, 149Sm-CD45RO, 150Nd-CD45RA, 151Eu-CD161, 152Sm-CD194, 153Eu-CD25, 154Sm-CD27, 155Gd-CD57, 156Gd-CD183, 158Gd-CD185, 160Gd-CD28, 161Dy-CD38, 163Dy-CD56, 164Dy-TCRγδ, 166Er-CD294, 167Er-CD197, 168Er-CD14, 170Er-CD3, 171Yb-CD20, 172Yb-CD66b, 173Yb-HLA-DR, 174Yb-IgD, 176Yb-CD127 were normalized using the 99.9th percentile prior to clustering. Cells were initially grouped into 125 clusters based on lineage marker expressions using the R package FlowSOM (v2.10.0)^65^. These clusters were subsequently manually merged and annotated according to biological relevance, yielding 43 distinct cell populations and subpopulations.

### Cell type composition analysis

A generalized linear mixed model (GLMM) with a Poisson outcome^66^ was used to model the impact of conditions on cell–type–specific counts, with conditions included as a covariate. The effect of this factor on cell-type composition was estimated through interaction terms with cell type. Models were fitted using the glmer function from the R package *lme4* (v1.1.37). The fold change is relative to the grand mean and adjusted. Statistical significance was evaluated using the local true sign rate (LTSR), which represents the probability that the estimated direction of effect is correct.

### Random Forest analysis

A random forest model was used to evaluate the impact of Te-labeled MBC-EVs and Te-labeled MBC-secreted proteins on recipient cells relative to control cells. For robust comparisons, cells with EV and protein signal levels in the top 20% (highest quintile) were selected. The primary MBC-EV (or Te-labeled protein) and control data were combined, and features were normalized using z-score transformation. This integrated dataset was used to train a random forest classifier with the RandomForestClassifier() function from the Python package scikit-learn (v1.4.0). To address class imbalance, class weights were applied, and hyperparameters were optimized via 5-fold cross-validation. Feature importance was calculated as the average across 10 nested cross-validation runs, with 5 permutations per feature, to estimate each feature’s contribution to group differentiation.

### Visualization

Visualizations were carried out in R (v.4.3.2) and Python (v.3.10) using scanpy (v.1.11.0)^67^, which is a scalable toolkit for analyzing single-cell data. For data visualization, high-dimensional single-cell data were reduced to two dimensions using the nonlinear dimensionality reduction algorithm t-SNE (t-distributed stochastic neighbor embedding)^68^. t-SNE plots were created and visualized using scanpy.

### EV-TRACK

We have submitted all relevant data of our experiments to the EV-TRACK knowledge database (EV-TRACK ID: EV250125) with an EV-METRIC of 86.57%^69^.

### Statistics

All statistical differences were calculated in GraphPad Prism, R, or Python. Statistical tests are indicated in the corresponding figure legends. The immunoblot analyses were repeated twice and showed consistent results. p values below 0.05 were considered statistically significant and are indicated in the figures as follows: *p < 0.05, **p < 0.01, ***p < 0.001, and ****p < 0.0001

## Acknowledgments

DB and MH are funded by the German Research Foundation (DFG) SFB1530-455784452 (subprojects B01, Z01, Z02) and the José Carreras Leukemia Foundation (Grant DJCLS 04R/2021). NA is supported by the Cancer Research Center Cologne Essen (CCCE) as well as by the German Research Foundation (DFG) SFB1530-455784452 (subproject Z03) and Project-ID 497777992. DB, DS, RTU, AVL, and HJ are supported by the Mildred Scheel Nachwuchszentrum Grant 70113307 from the German Cancer Aid. RDJ is funded by the German Research Foundation (DFG) FOR5504-496650118 (subproject TP05), JA2439/4-1, and SFB1530-455784452 (subproject C02), by the Ministry for Culture and Science of North Rhine-Westphalia (NW21-062A CANTAR), and by the Behrens-Weise-Foundation (for the Improvement of Human Health). DS was supported by an MD Research Stipend of the Else Kröner Forschungskolleg Clonal Evolution in Cancer, University Hospital Cologne, Cologne, Germany. DB is supported by the Exzellenz Initiieren (E.I.) – Stiftung Kölner Krebsforschung. DB and BE are supported by the German Federal Ministry of Education and Research (BMBF; project 01KU2108) within the ERA PerMed framework. We thank Dr. Dirk Strunk (Paracelsus Medical University) for his scientific insights and guidance. We also thank Dr. Jan van Deun (University of Erlangen-Nuremberg) and Dr. Anna Kashkanova (Max Planck Institute for the Science of Light, Erlangen) for their experimental support. Additionally, we appreciate Dr. Christian Jüngst and Beatrix Martiny (CECAD Imaging Facility, University of Cologne) for their experimental assistance. (Imaging) Mass cytometric data acquisition was carried out by Petra Hofmann (Department I of Internal Medicine) and Marion Müller (Institute of Pathology) at the CyTOF Facility of the Medical Faculty, University of Cologne, supported by the DFG-funded CRC1530 (Project Z02). Immunogold transmission electron microscopy was performed at the CECAD Imaging Facility using a Jeol JEM 2100 Plus, funded by DFG-INST 216/793-1 FUGG. We additionally thank the Regional Computing Center of the University of Cologne for providing computing time on the DFG-funded High Performance Computing system CHEOPS (Funding number: DFG-INST 216/512-1 FUGG) and for support. The NanoSIMS measurements were performed at the NanoSIMS Sweden Facility at the University of Gothenburg, SciLifeLab Swedish infrastructure. Graphical representations were created with BioRender (https://BioRender.com/hlo8aeq).

---

**Extended Data Fig. 1.**
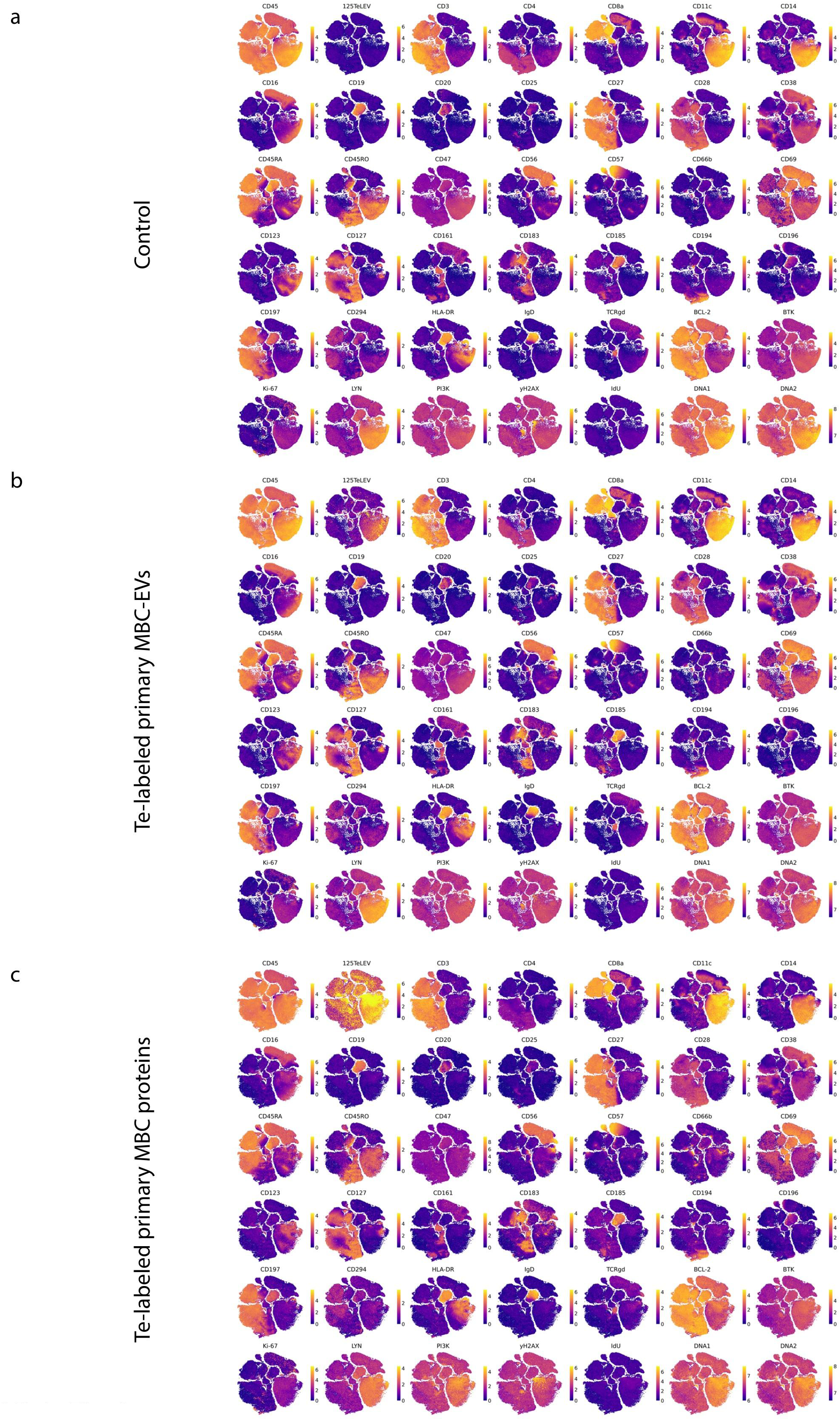
High-dimensional profiling of MBC-EV recipients. a-c, t-SNE plots showing selected marker expressions for the three conditions. PBS-HAT control, primary Te-labeled MBC-EVs, and matched Te-labeled secreted proteins.

**Extended Data Fig. 2.**
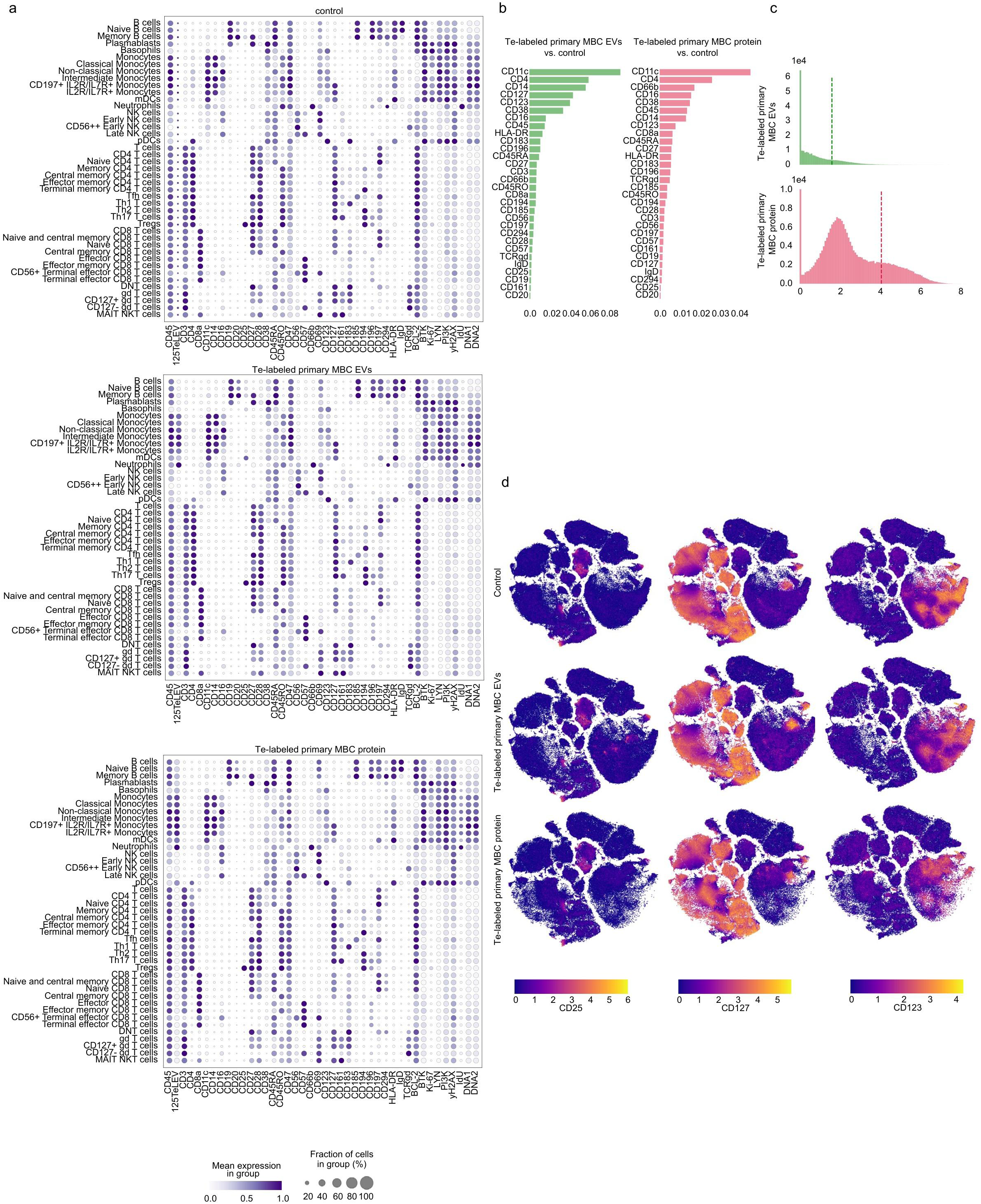
Characterization of Te-labeled CLL-derived EVs and secreted proteins across immune cell populations. a, Dot plot depicting the expression of selected markers across major cell types and their subtypes for three conditions: PBS-HAT control (top, 329,774 cells), primary Te-labeled MBC-EVs (middle, 226,157 cells), and Te-labeled secreted proteins (bottom, 229,329 cells). Dot size corresponds to the proportion of cells expressing each marker within a given cell type, and color intensity indicates the mean expression level. b, Feature importance of surface markers for distinguishing the primary MBC-EV from control data, as determined by a random forest classifier (left). The same analysis is shown for matched Te-labeled secreted proteins (right). c, TeLEV signal histograms for primary MBC-EVs (top) and matched secreted proteins (bottom). The dashed lines indicate the 80th-percentile cutoff. d, t-SNE plots showing CD25, CD127, and CD123 expressions for the three conditions.

**Extended Data Fig. 3.**
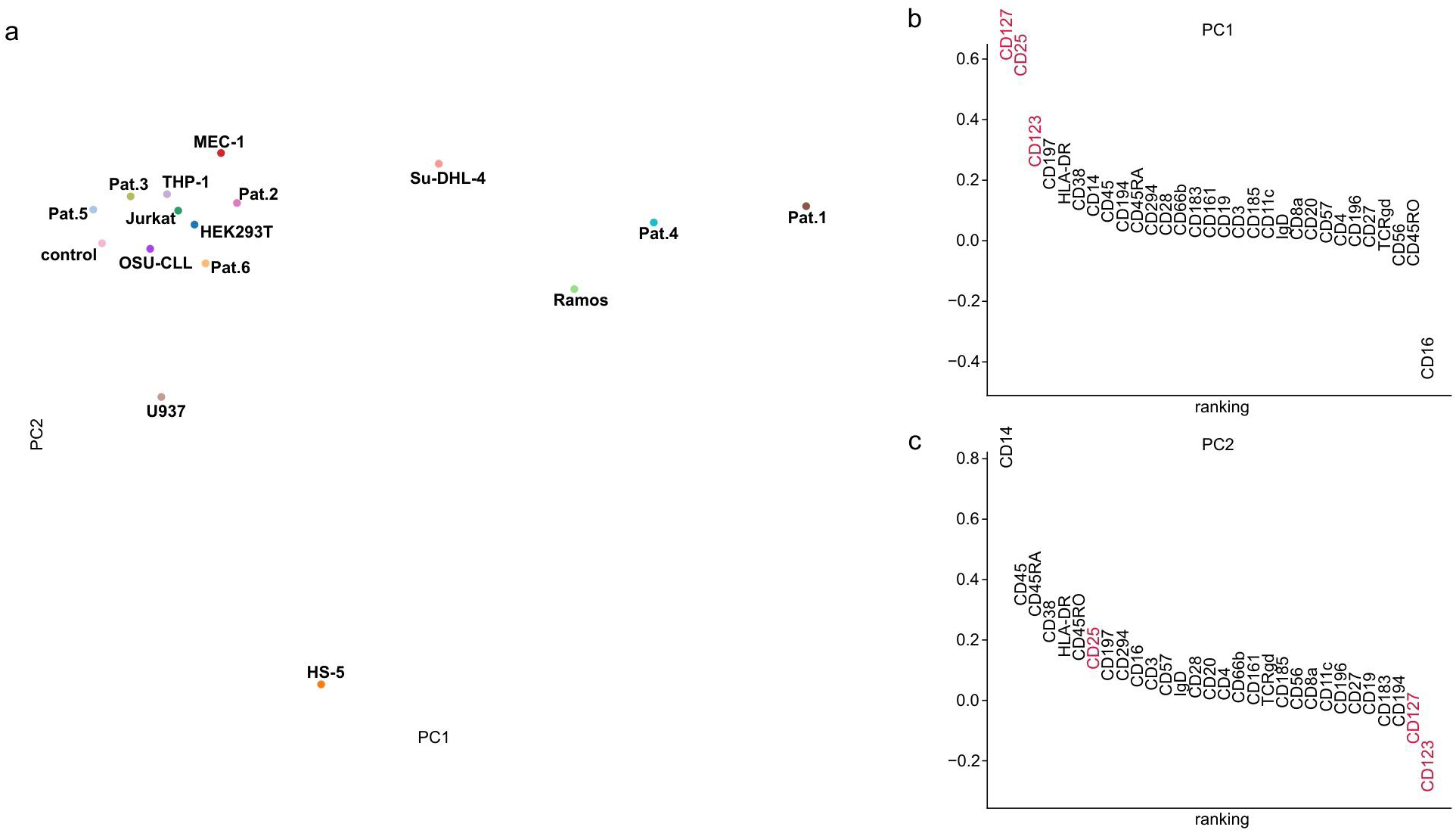
PCA analysis of lineage-defining markers. a, The mean expression of lineage-defining markers across monocytes, mDCs, and pDCs for different EV sources in principal components space (PC1and PC2). b, Lineage-defining markers ranked by the contribution to PC1. The highest-variability axis (PC1) separates four EV sources with the MBC signature from the remaining EV sources. CD127, CD25, CD123, and CD197 are the strongest contributors to PC1. c, Lineage-defining markers ranked by the contribution to PC2.

**Extended Data Fig. 4.**
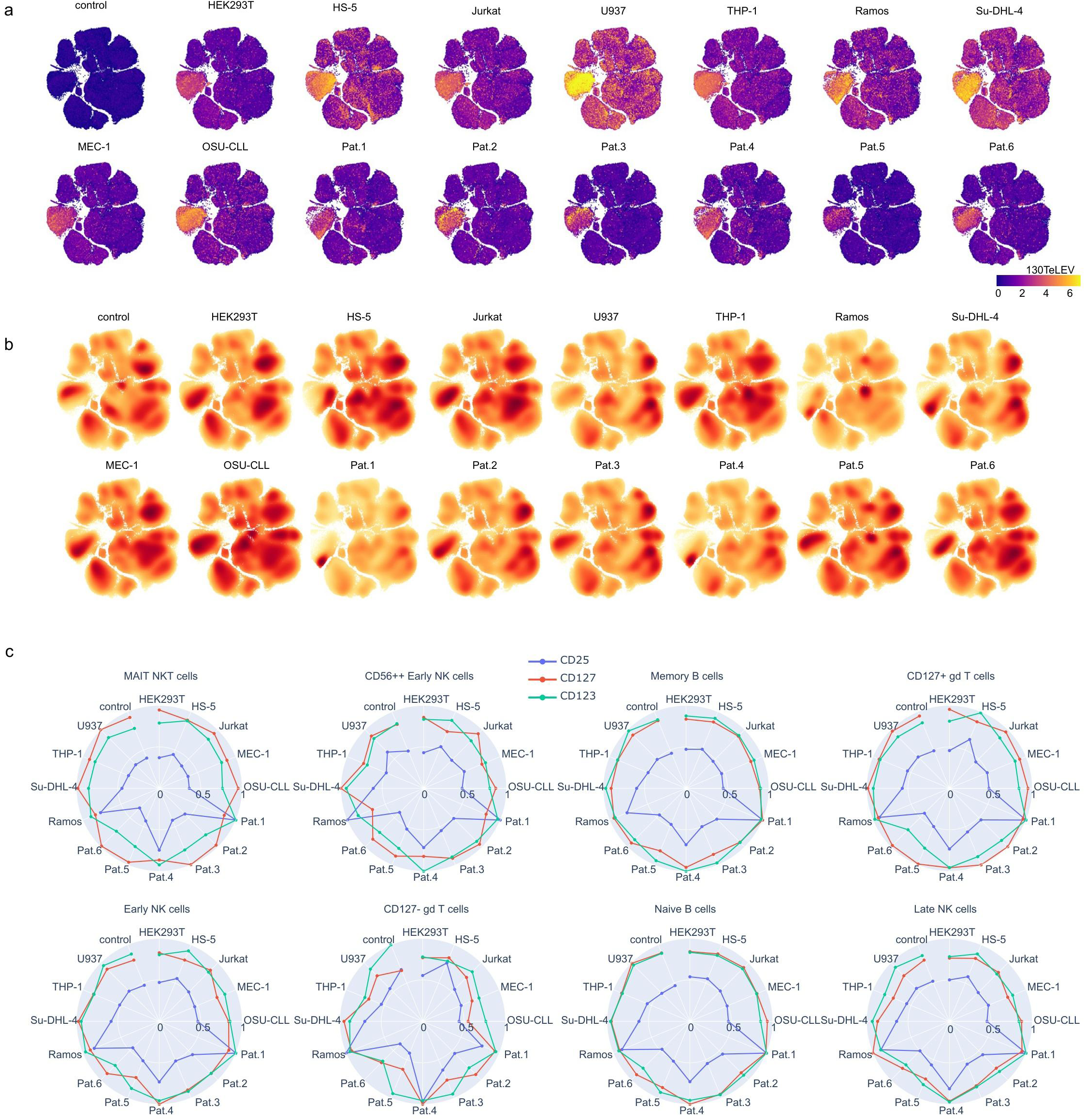
Comparison of EV uptake from primary and cell line-derived EV sources. a, t-SNE projection of PBMC recipient cells for 16 conditions (PBS-HAT control plus 15 EV sources) showing TeLEV signal. b, Two-dimensional density distributions of cells in t-SNE embedding space, stratified by conditions. c, IL-RT expression in the cell types that showed the highest differential expression of CD25 between four EV sources with the MBC signature (Patient 1, Patient 4, Ramos, and Su-DHL-4) and the remaining EV sources.

**Extended Data Fig. 5.**
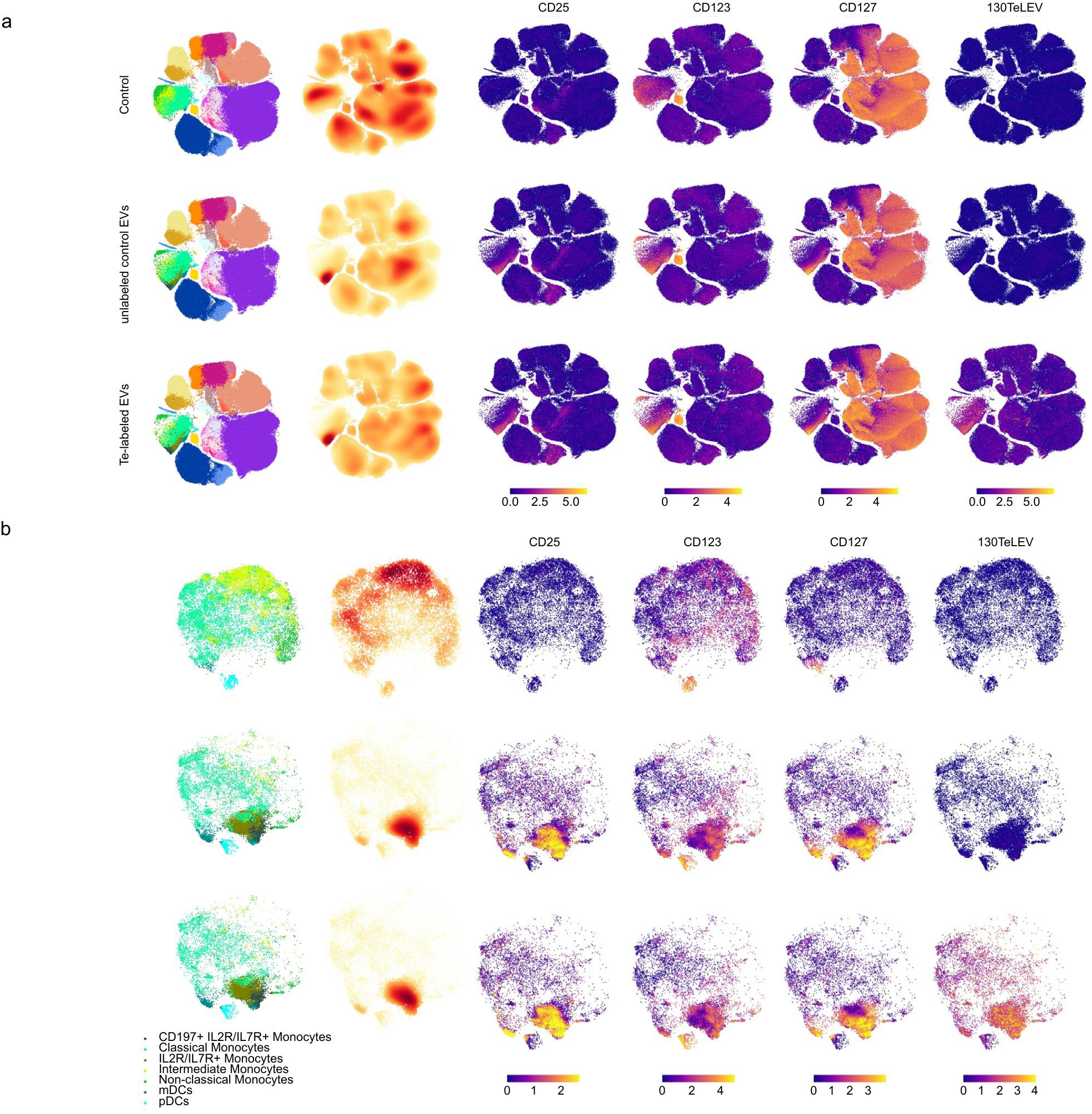
Recipient cell comparison of Te-labeled and unlabeled EV uptake. a, t-SNE projection of PBMC recipients of PBS-HAT control (200,468 cells), unlabeled control EVs (200,629 cells), and Te-labeled MBC-EVs (191,964 cells) from primary CLL cells colored according to cell type (column 1). Two-dimensional density distributions of cells in t-SNE embedding space, stratified by conditions (column 2). t-SNE plots showing CD25, CD123, and CD127 expression, and TeLEV signal (columns 3-6). The unlabeled control EV condition shows a minimal TeLEV background signal. b, t-SNE projection of the myeloid compartment. Shown are recipient cells of PBS-HAT control (17,445 cells), unlabeled control MBC-EVs (19,003 cells), and Te-labeled MBC-EVs derived from primary CLL cells (15,300 cells).

**Extended Data Fig. 6.**
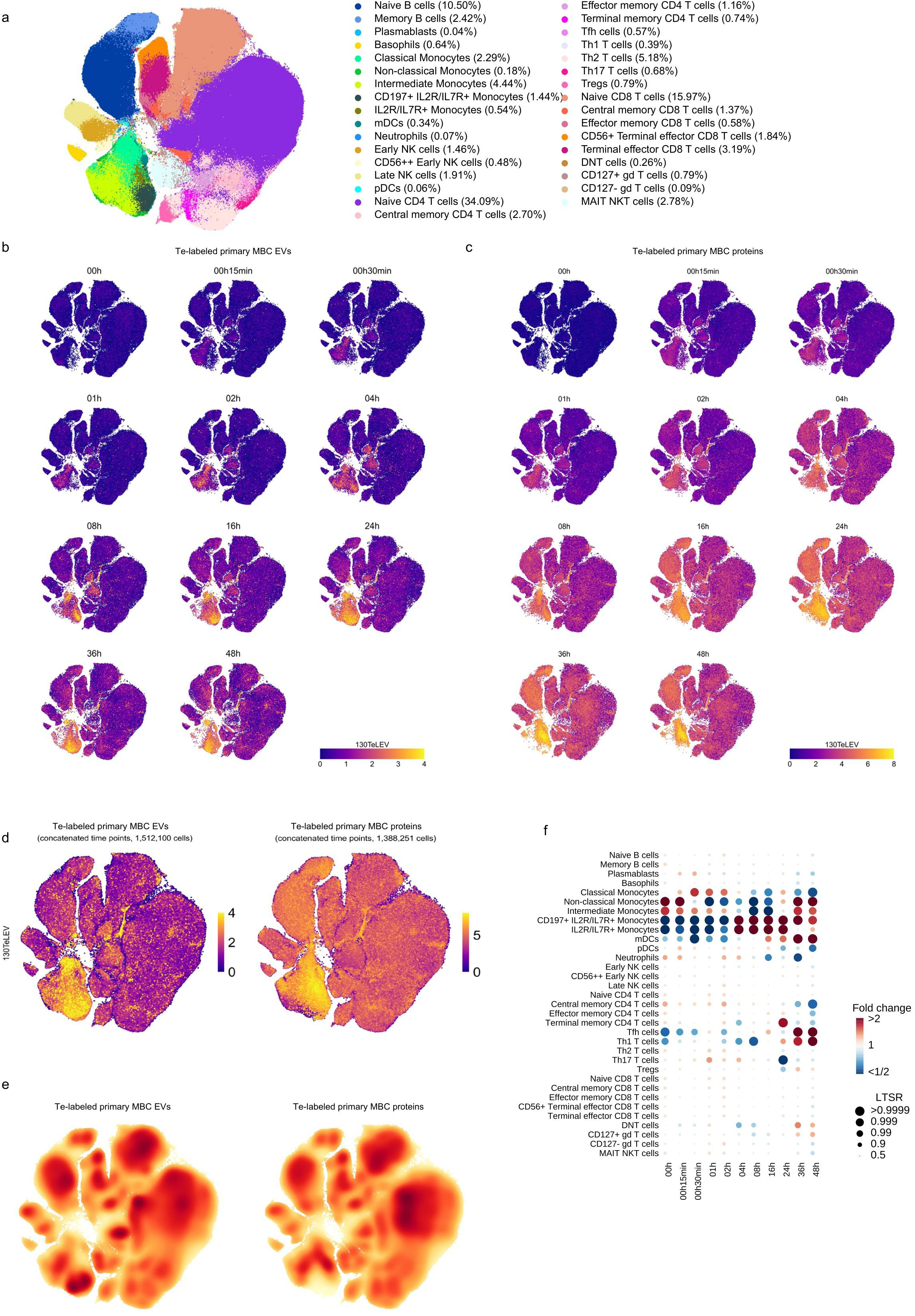
Time-resolved uptake of Te-labeled primary MBC-EVs and MBC-secreted proteins. a, t-SNE projection of PBMC recipient cells showing uptake dynamics across 11 time points for primary Te-labeled MBC-EVs and 11 time points for matched secreted proteins (in total 2,900,351 cells), colored according to cell type. Numerical annotations indicate the relative proportions of each cell lineage. b, t-SNE plots showing TeLEV expressions for 11 time points of MBC-EV uptake. c, t-SNE plots showing TeLEV expressions for 11 time points of Te-labeled secreted proteins uptake. d, t-SNE plots showing TeLEV expression for primary MBC-EV uptake (left, 1,512,100 cells) and matched Te-labeled secreted protein uptake (right, 1,388,251 cells) across concatenated time points. e, Two-dimensional density distributions of cells in t-SNE space for MBC-EV (left) and matched secreted protein conditions (right). f, Fold changes in cell type proportions across 11 time points of EV uptake, estimated using a Poisson generalized linear mixed model. Dot size represents the probability of change, measured as the local true sign rate (LTSR).

**Extended Data Fig. 7.**
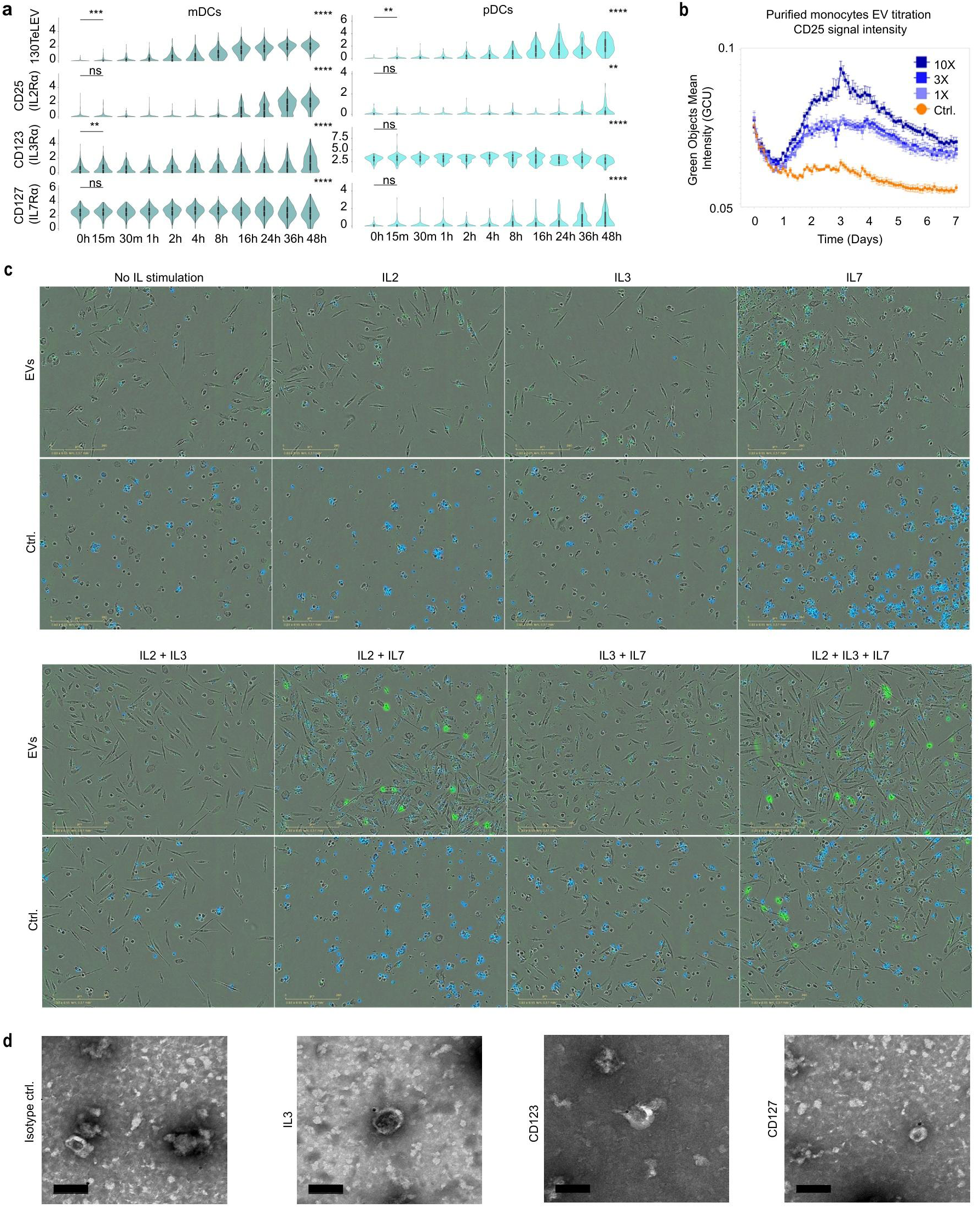
IL-RT-dependent myeloid polarization by MBC-EVs. a, Comparison of TeLEV signal, CD25, CD123, and CD127 distributions within mDCs (left) and pDCs (right) over 48 h. Statistical significance across all time points for each marker was evaluated using the Kruskal-Wallis test. Pairwise comparisons between 00 h and 00 h 15 min were performed using a one-sided Mann–Whitney U test. *p-values* < 0.05 were considered significant and are indicated in the figures as follows: \**p* < 0.05, \*\**p* < 0.01, \*\*\**p* < 0.001, and \*\*\*\**p* < 0.0001. b, Single-cell quantification of CD25 (Alexa Fluor 488) fluorescence over 7 days corresponding to (h). Values are unnormalized arbitrary units; data shown as mean ± SEM; n = 4 fields of view. c, Live-cell immunocytochemistry of EV-exposed monocytes stimulated with cytokines. Representative images from day 7 are shown for IL-2 (20 ng mL⁻¹), IL-3 (5 ng mL⁻¹), or, IL-7 (25 ng mL⁻¹), or IL-2 + IL-3, IL-2 + IL-7, IL-3 + IL-7, or IL-2 + IL-3 + IL-7 in RPMI with serum. CD25 (Alexa Fluor 488; green) and Cytotox NIR (blue) are shown after automated segmentation and background subtraction. Imaging on Incucyte SX5 (20×, 2 h intervals). Scale bar, 200 µm. d, Immunogold transmission electron microscopy of Ramos-EVs labeled for IL-3, CD123, and CD127 alongside an isotype control. Scale bar, 100 nm.

**Extended Data Fig. 8.**
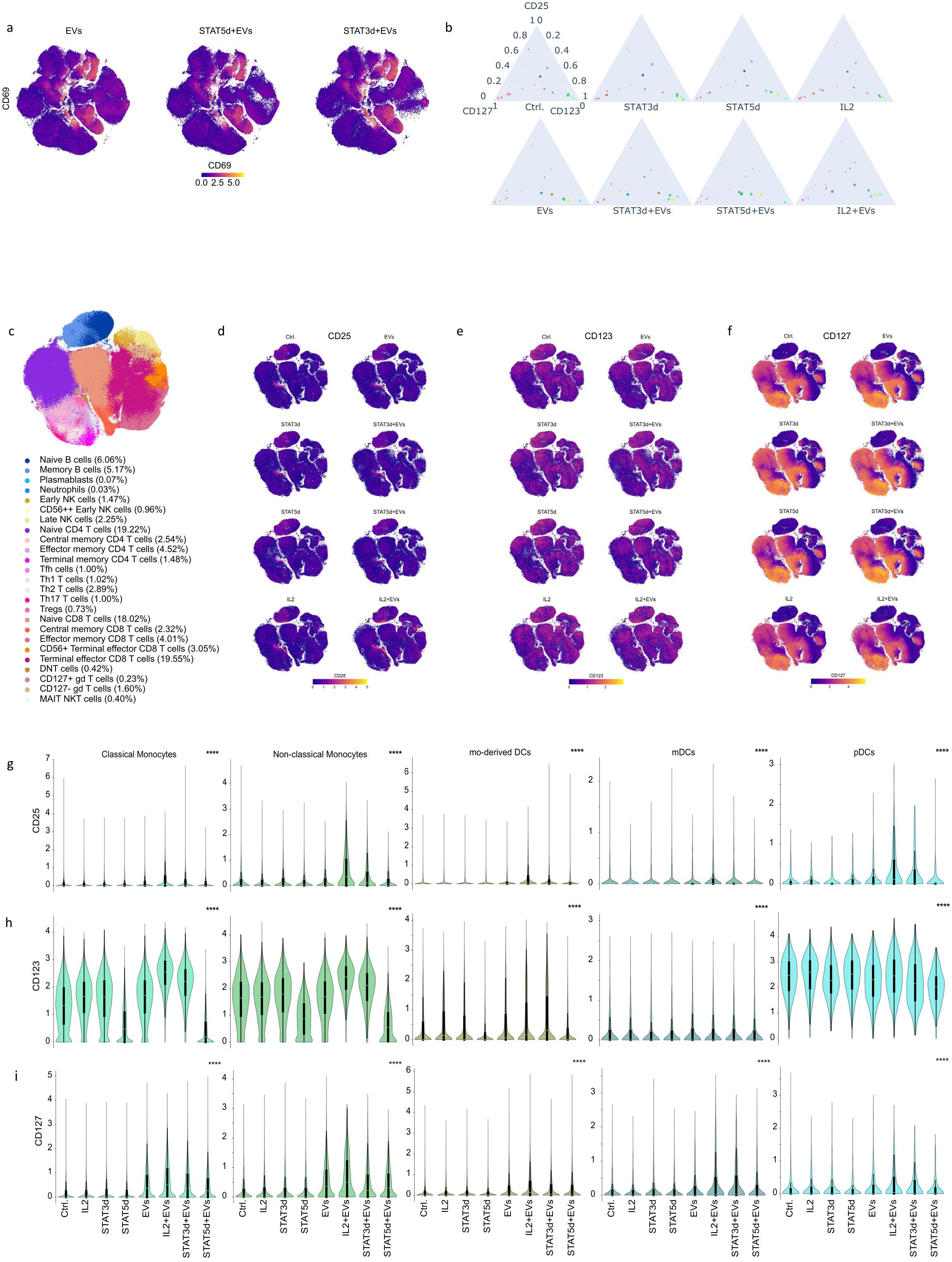
Global immune cell expression of the IL-RT. a, t-SNE projection of cells showing CD69 expression. Shown are MBC-EVs (217,211 cells), STAT5d + MBC-EVs (226,884 cells), and STAT3d + MBC-EVs (206,018 cells) conditions. b, A cell type–color–coded ternary plot depicting expression levels of CD25, CD123, and CD127. Circle size reflects the average TeLEV level within each cell type. c, t-SNE projection of the non-myeloid population of 16 conditions (2,725,107 cells), colored according to cell types. Numerical annotations indicate the relative proportions of each cell lineage. d-f, t-SNE plots showing CD25 (d), CD123 (e), and CD127 (f) expressions on the non-myeloid population across the indicated conditions. g-i, Comparison of CD25 (g), CD123 (h), and CD127 (i) expression distributions across the indicated conditions in Monocytes, moDCs, mDCs, and pDCs. For each cell type, Statistical significance across all conditions was evaluated using the Kruskal-Wallis test. \*\*\*\**p* < 0.0001. White lines denote the median, edges the IQR, and whiskers either 1.5 × IQR or minima/maxima (if no point exceeded 1.5 × IQR; minima/maxima are indicated by the violin plot range).

## Supplementary Figure legends

Supplementary Fig. 1 Immune recipient cell mapping of primary MBC-EV and MBC-secreted protein uptake. t-SNE projection of major cell types per condition. Shown are cell type overlays, two-dimensional density distributions in t-SNE space, and overlays of selected marker expressions of primary Te-labeled MBC-EV and matched secreted protein uptake.

Supplementary Fig. 2 Pan-immune cell atlas of EV uptake t-SNE projection of major cell types per condition. Shown are cell type overlays, two-dimensional density distributions in t-SNE space, and overlays of selected marker expressions.

Supplementary Table 1 EV quantification of the pan-immune EV uptake atlas Supplementary Table 2 MBC-EV-induced expression ranking of CD25 Supplementary Table 3 CLL patient characteristics

Supplementary Table 4 Antibodies and reagents

