## Supplementary Figure 1 for "Single recipient cell tracking of tellurium-labeled extracellular vesicle proteomes (TeLEV) identifies EV-driven immunomodulation"

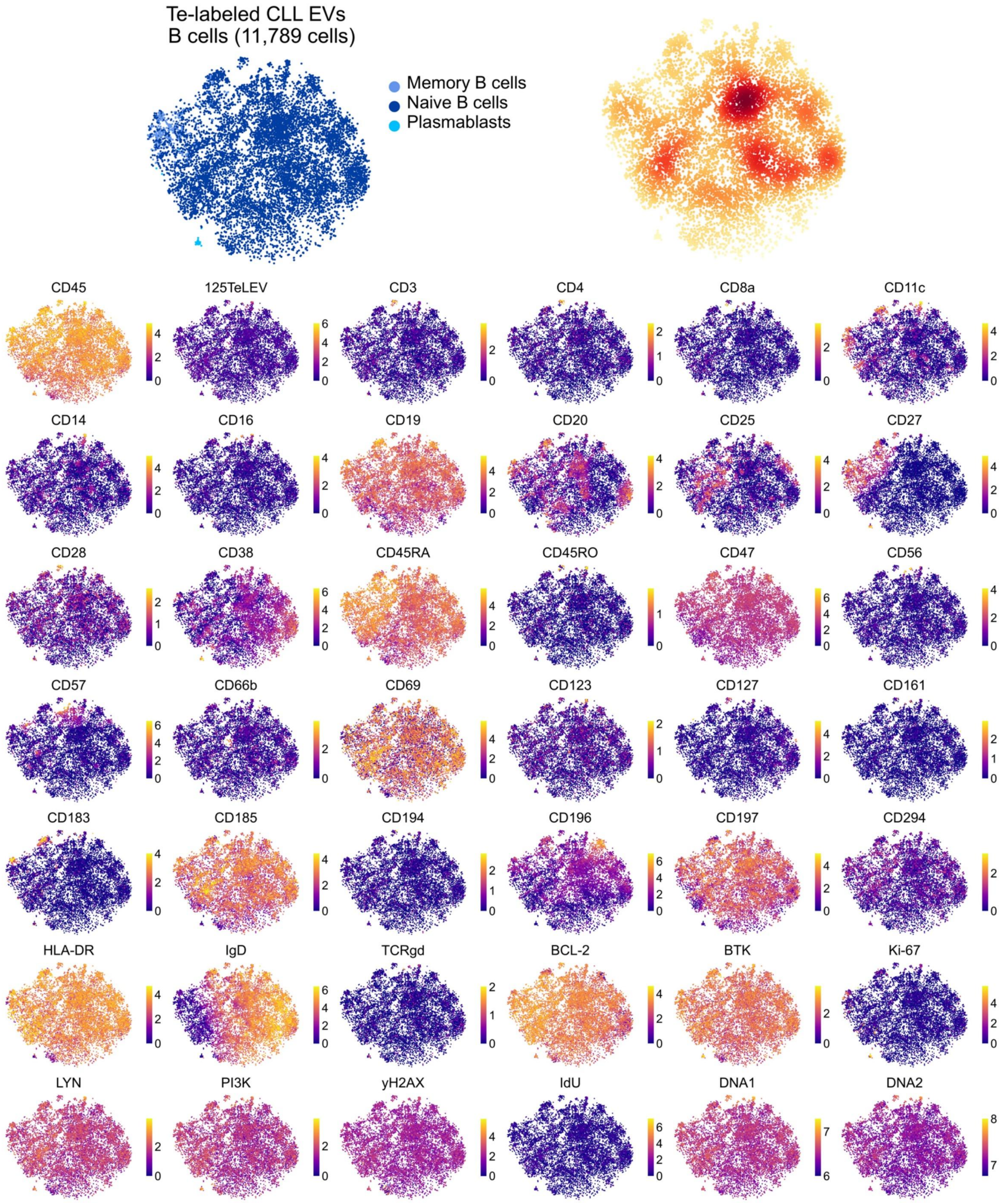
Te-labeled CLL EVs
B cells (11,789 cells)
Memory B cells
Naive B cells
Plasmablasts
CD45
125TeLEV
CD3
CD4
CD8a
CD11c
CD14
CD16
CD19
CD20
CD25
CD27
CD28
CD38
CD45RA
CD45RO
CD47
CD56
CD57
CD66b
CD69
CD123
CD127
CD161
CD183
CD185
CD194
CD196
CD197
CD294
HLA-DR
IgD
TCRgd
BCL-2
BTK
Ki-67
LYN
PI3K
yH2AX
IdU
DNA1
DNA2

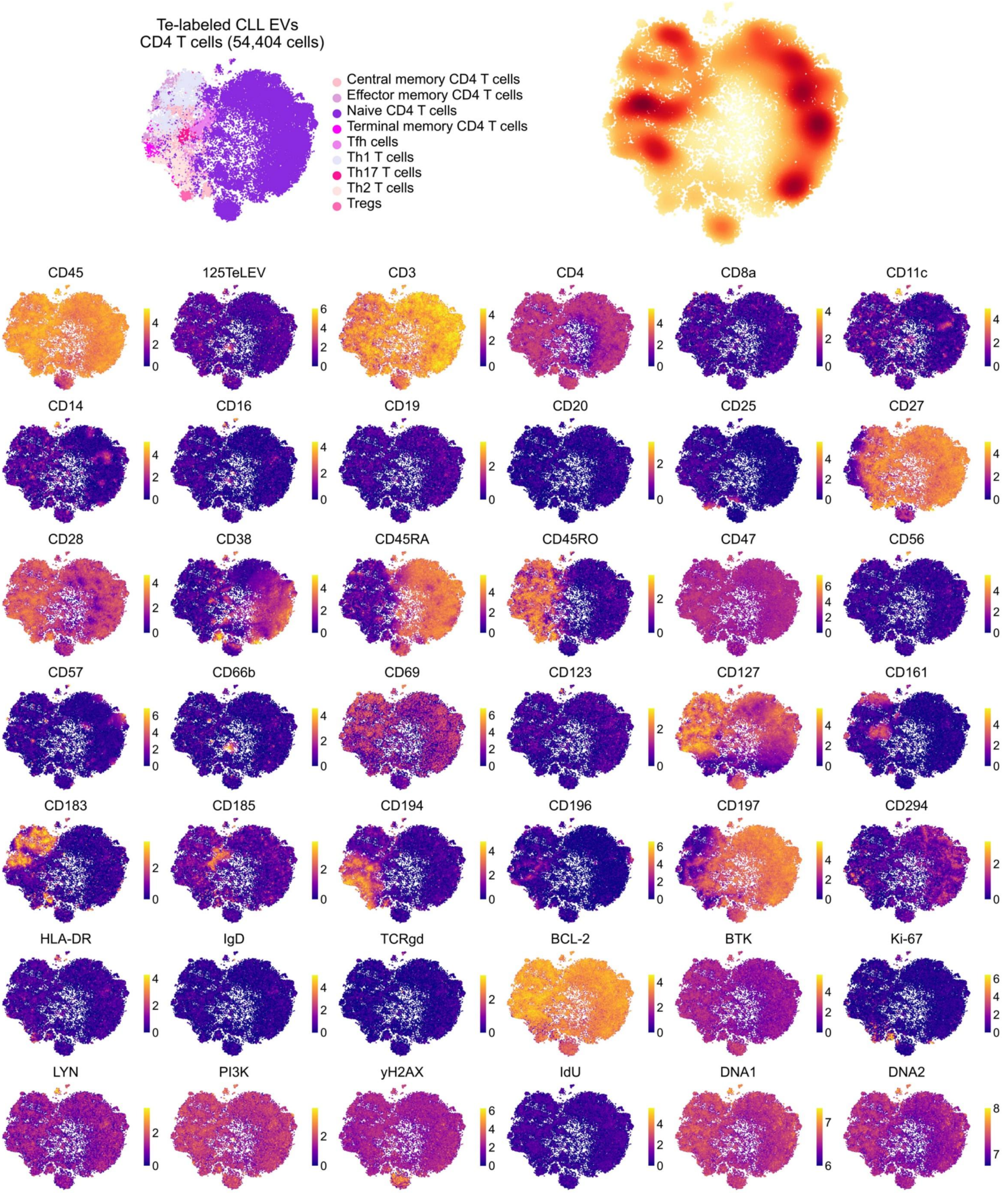

Te-labeled CLL EVs
CD4 T cells (54,404 cells)
Central memory CD4 T cells
Effector memory CD4 T cells
Naive CD4 T cells
Terminal memory CD4 T cells
Tfh cells
Th1 T cells
Th17 T cells
Th2 T cells
Tregs
CD45
125TeLEV
CD3
CD4
CD8a
CD11c
CD14
CD16
CD19
CD20
CD25
CD27
CD28
CD38
CD45RA
CD45RO
CD47
CD56
CD57
CD66b
CD69
CD123
CD127
CD161
CD183
CD185
CD194
CD196
CD197
CD294
HLA-DR
IgD
TCRgd
BCL-2
BTK
Ki-67
LYN
PI3K
yH2AX
IdU
DNA1
DNA2

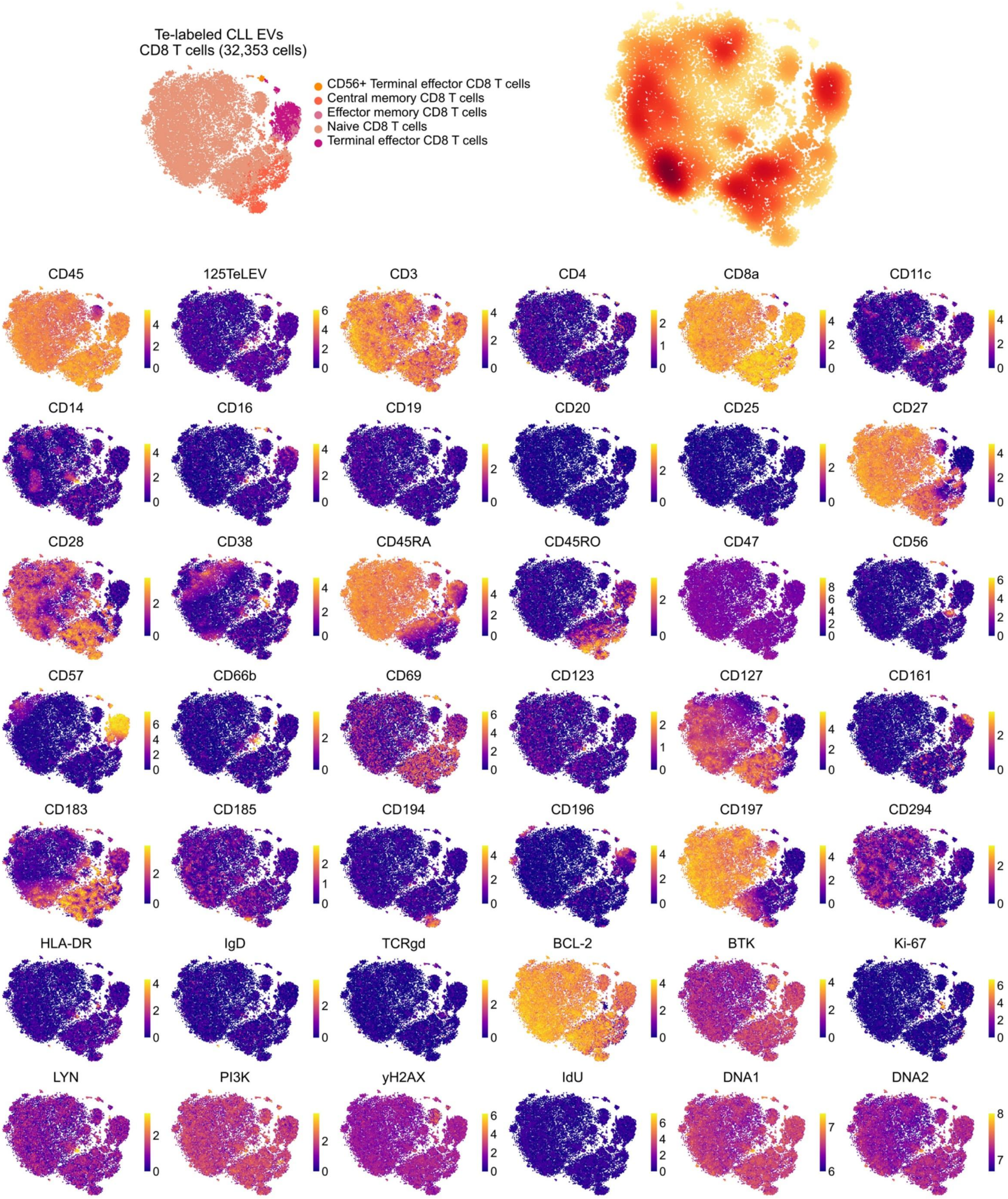
Te-labeled CLL EVs
CD8 T cells (32,353 cells)
CD56+ Terminal effector CD8 T cells
Central memory CD8 T cells
Effector memory CD8 T cells
Naive CD8 T cells
Terminal effector CD8 T cells
CD45
125TeLEV
CD3
CD4
CD8a
CD11c
CD14
CD16
CD19
CD20
CD25
CD27
CD28
CD38
CD45RA
CD45RO
CD47
CD56
CD57
CD66b
CD69
CD123
CD127
CD161
CD183
CD185
CD194
CD196
CD197
CD294
HLA-DR
IgD
TCRgd
BCL-2
BTK
Ki-67
LYN
PI3K
yH2AX
IdU
DNA1
DNA2

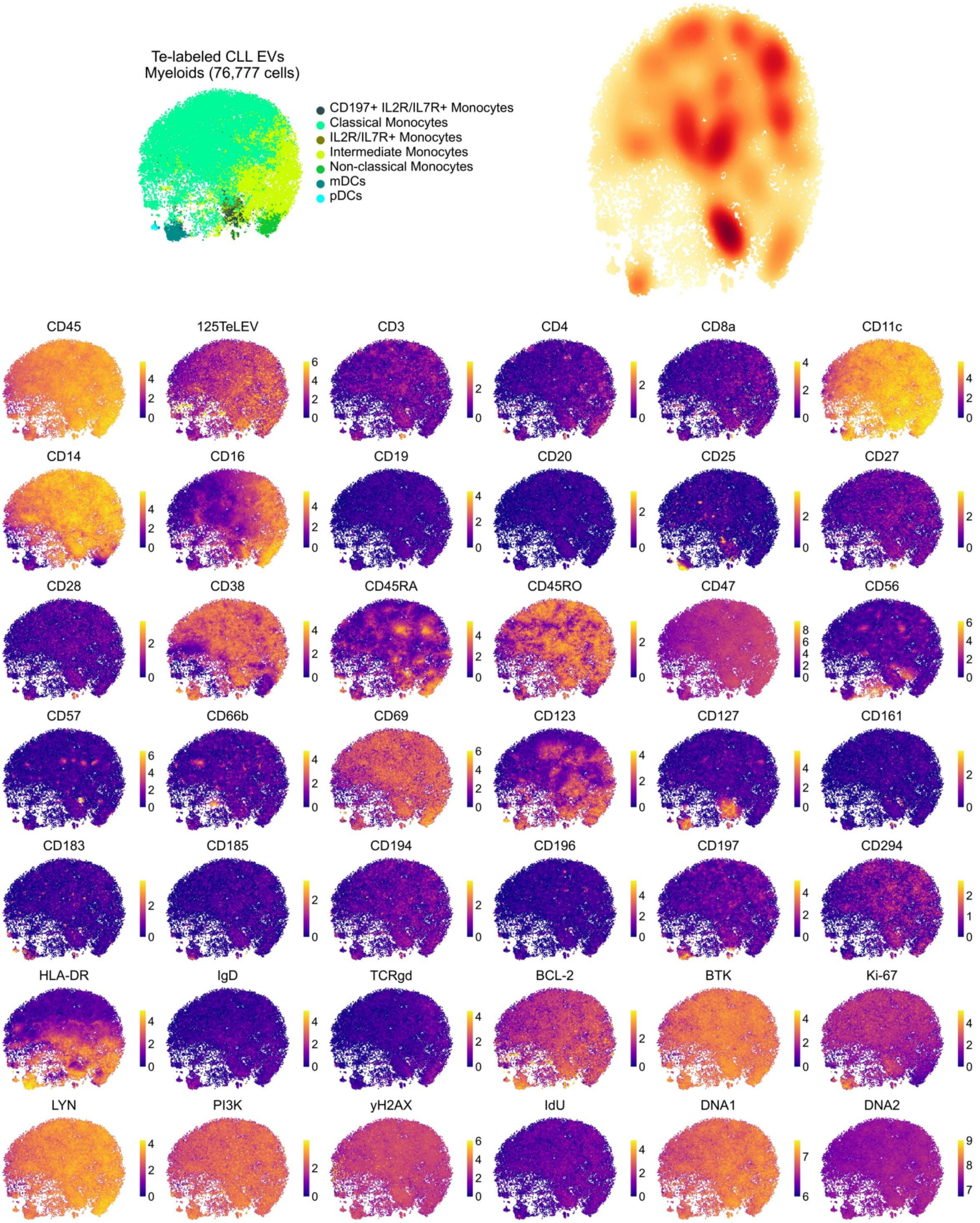
Te-labeled CLL EVs
Myeloids (76,777 cells)
CD197+ IL2R/IL7R+ Monocytes
Classical Monocytes
IL2R/IL7R+ Monocytes
Intermediate Monocytes
Non-classical Monocytes
mDCs
pDCs
CD45
125TeLEV
CD3
CD4
CD8a
CD11c
CD14
CD16
CD19
CD20
CD25
CD27
CD28
CD38
CD45RA
CD45RO
CD47
CD56
CD57
CD66b
CD69
CD123
CD127
CD161
CD183
CD185
CD194
CD196
CD197
CD294
HLA-DR
IgD
TCRgd
BCL-2
BTK
Ki-67
LYN
PI3K
yH2AX
IdU
DNA1
DNA2

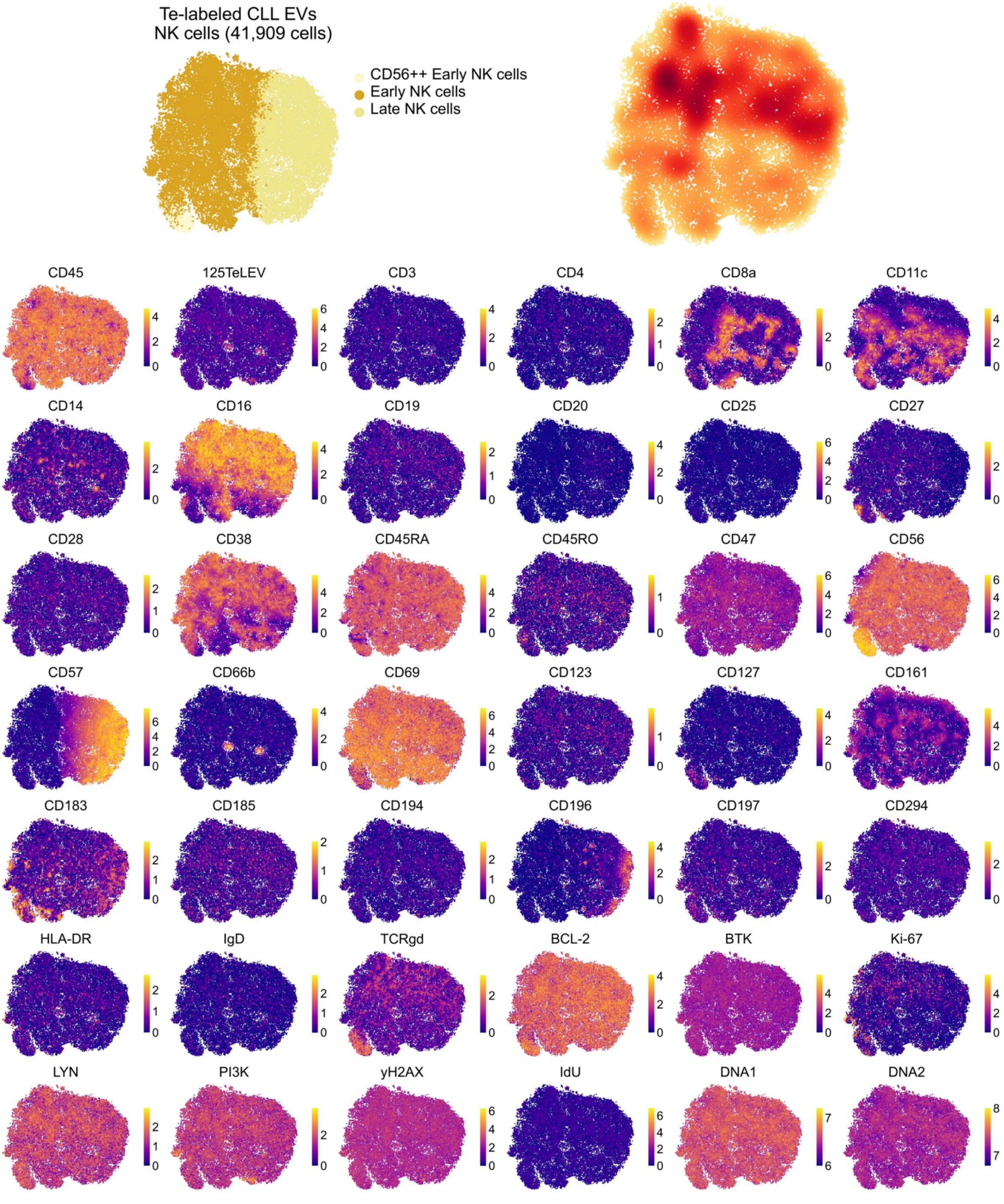

Te-labeled CLL EVs
NK cells (41,909 cells)
CD56++ Early NK cells
Early NK cells
Late NK cells
CD45
125TeLEV
CD3
CD4
CD8a
CD11c
CD14
CD16
CD19
CD20
CD25
CD27
CD28
CD38
CD45RA
CD45RO
CD47
CD56
CD57
CD66b
CD69
CD123
CD127
CD161
CD183
CD185
CD194
CD196
CD197
CD294
HLA-DR
IgD
TCRgd
BCL-2
BTK
Ki-67
LYN
PI3K
yH2AX
IdU
DNA1
DNA2

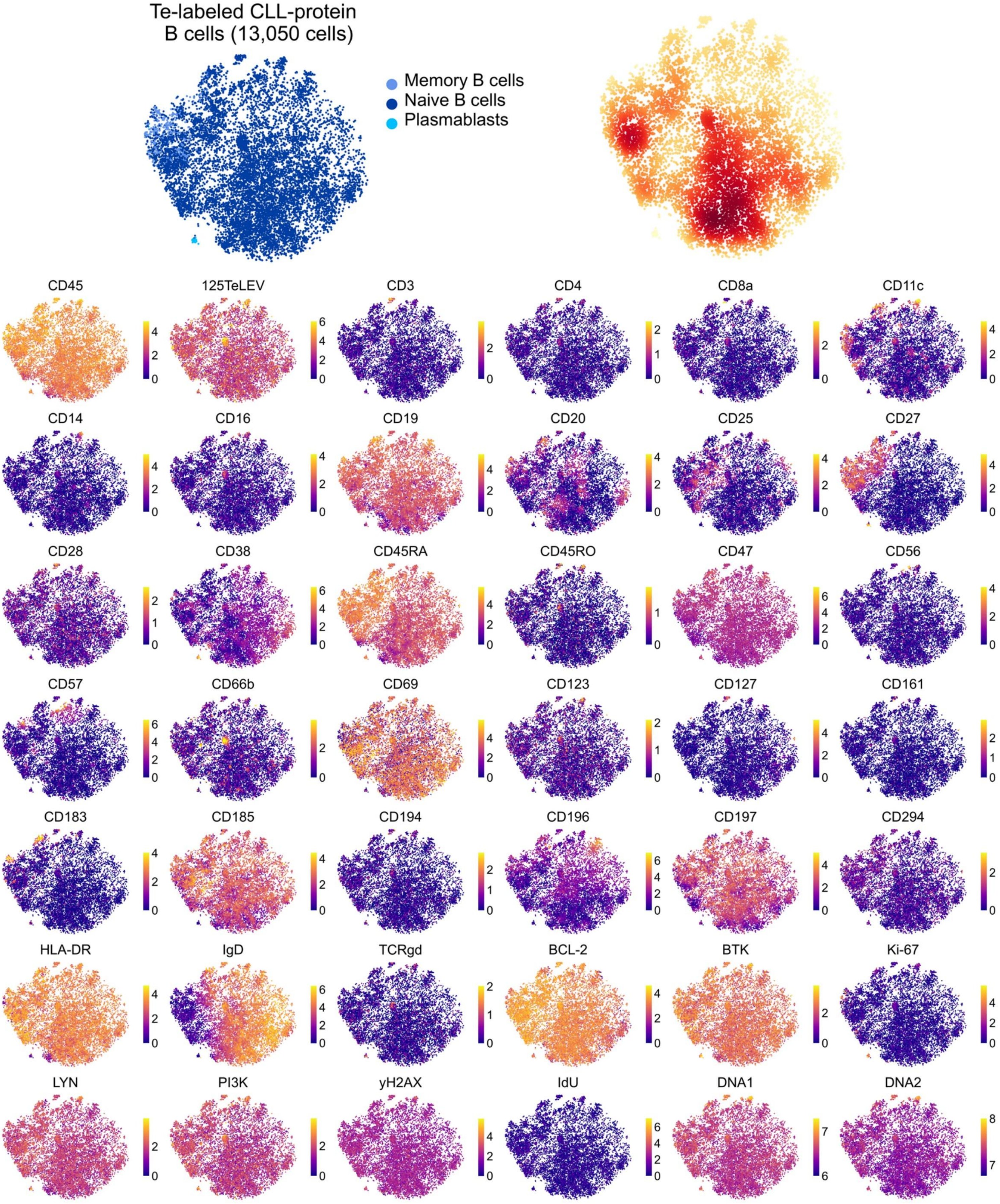
Te-labeled CLL-protein
B cells (13,050 cells)
Memory B cells
Naive B cells
Plasmablasts
CD45
125TeLEV
CD3
CD4
CD8a
CD11c
CD14
CD16
CD19
CD20
CD25
CD27
CD28
CD38
CD45RA
CD45RO
CD47
CD56
CD57
CD66b
CD69
CD123
CD127
CD161
CD183
CD185
CD194
CD196
CD197
CD294
HLA-DR
IgD
TCRgd
BCL-2
BTK
Ki-67
LYN
PI3K
yH2AX
IdU
DNA1
DNA2

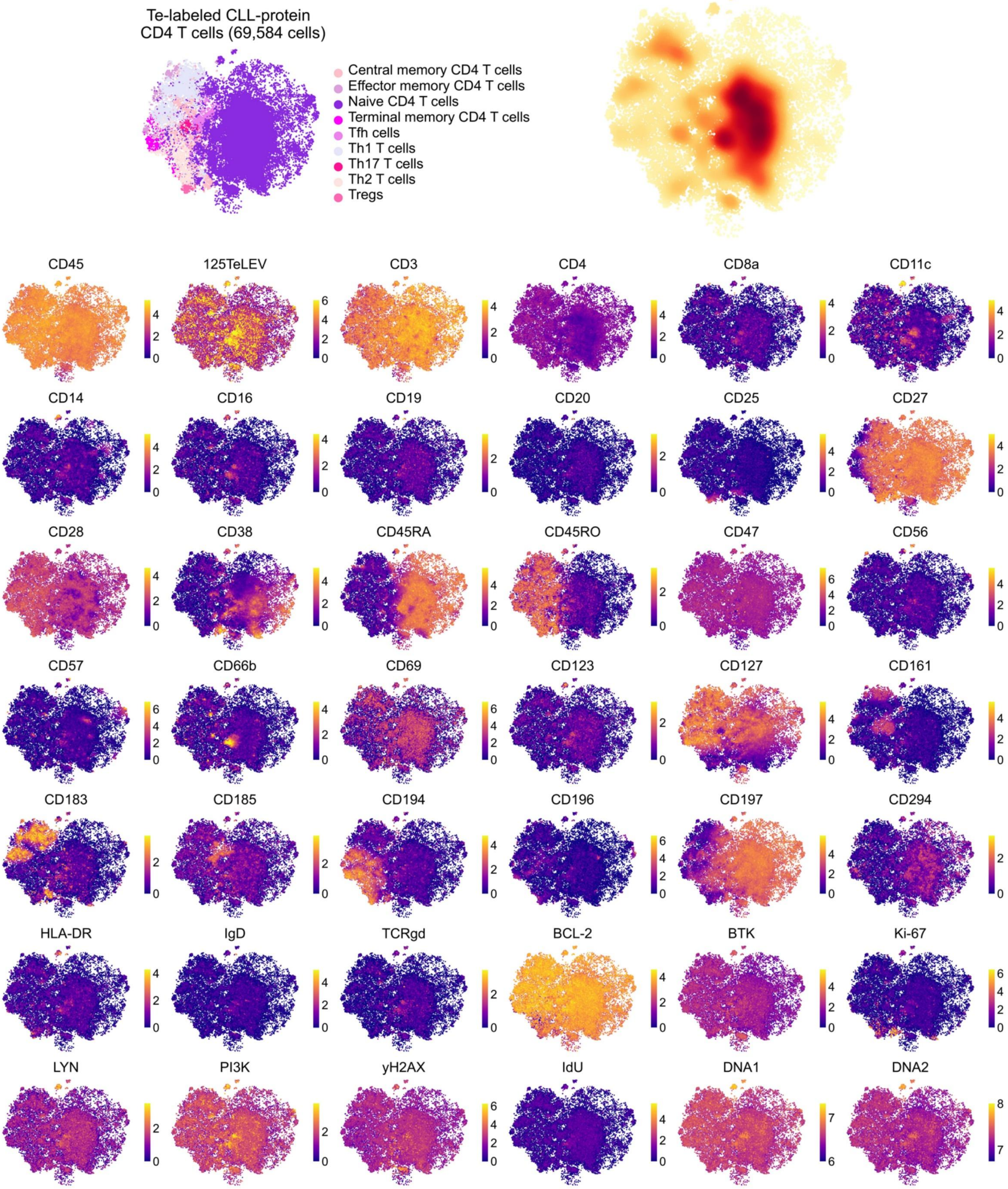
Te-labeled CLL-protein
CD4 T cells (69,584 cells)
Central memory CD4 T cells
Effector memory CD4 T cells
Naive CD4 T cells
Terminal memory CD4 T cells
Tfh cells
Th1 T cells
Th17 T cells
Th2 T cells
Tregs
CD45
125TeLEV
CD3
CD4
CD8a
CD11c
CD14
CD16
CD19
CD20
CD25
CD27
CD28
CD38
CD45RA
CD45RO
CD47
CD56
CD57
CD66b
CD69
CD123
CD127
CD161
CD183
CD185
CD194
CD196
CD197
CD294
HLA-DR
IgD
TCRgd
BCL-2
BTK
Ki-67
LYN
PI3K
yH2AX
IdU
DNA1
DNA2

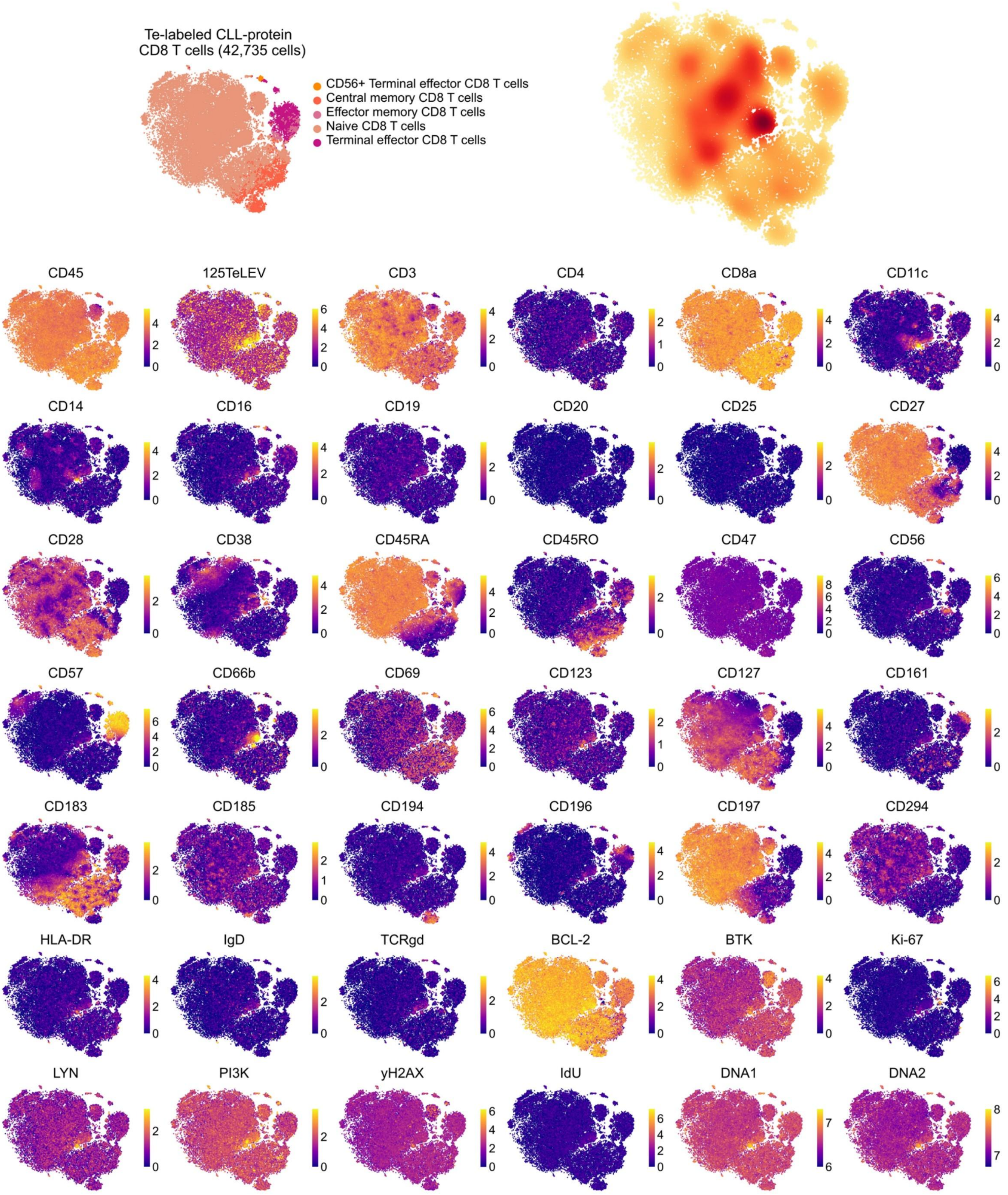
Te-labeled CLL-protein
CD8 T cells (42,735 cells)
CD56+ Terminal effector CD8 T cells
Central memory CD8 T cells
Effector memory CD8 T cells
Naive CD8 T cells
Terminal effector CD8 T cells
CD45
125TeLEV
CD3
CD4
CD8a
CD11c
CD14
CD16
CD19
CD20
CD25
CD27
CD28
CD38
CD45RA
CD45RO
CD47
CD56
CD57
CD66b
CD69
CD123
CD127
CD161
CD183
CD185
CD194
CD196
CD197
CD294
HLA-DR
IgD
TCRgd
BCL-2
BTK
Ki-67
LYN
PI3K
yH2AX
IdU
DNA1
DNA2

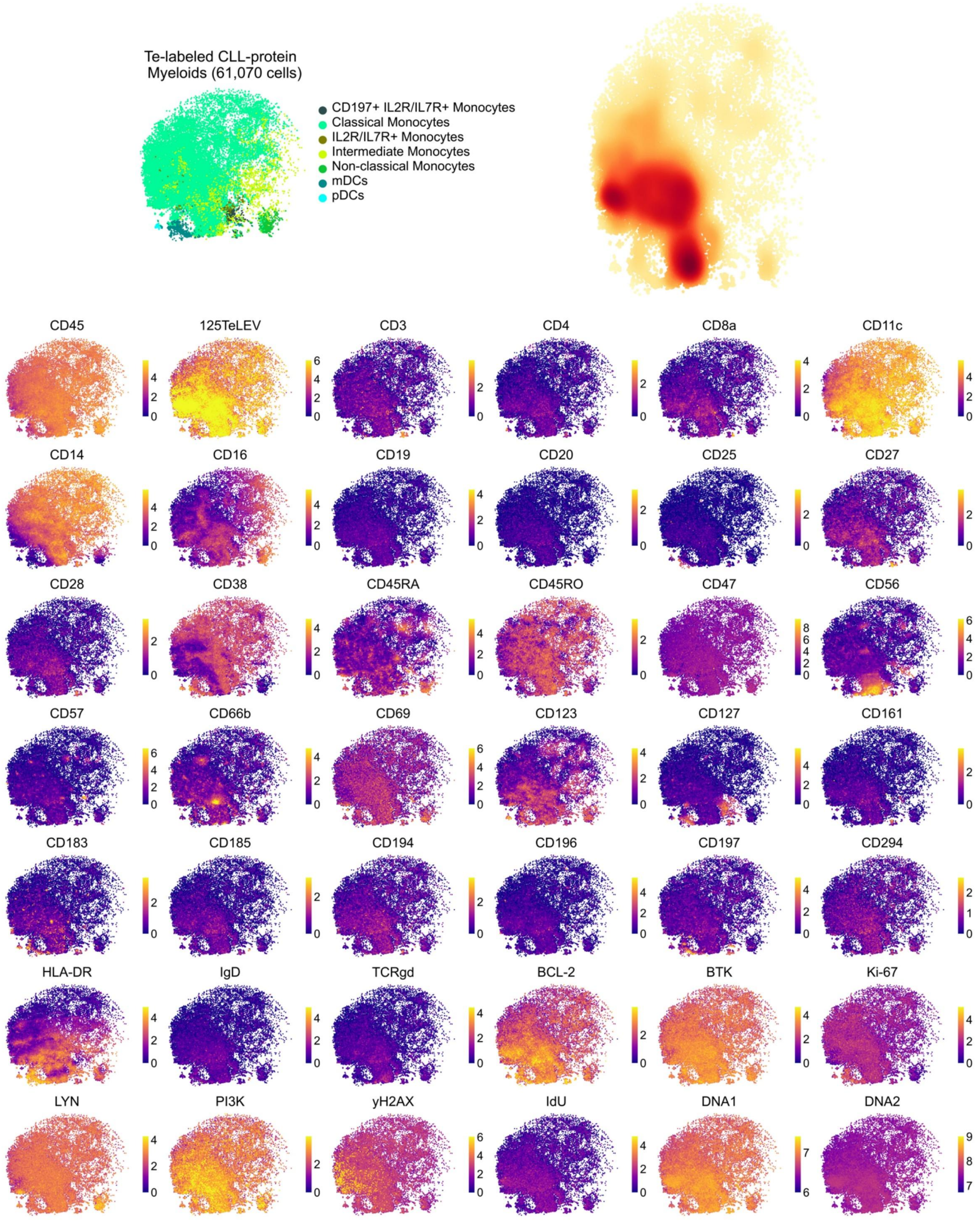
Te-labeled CLL-protein
Myeloids (61,070 cells)
CD197+ IL2R/IL7R+ Monocytes
Classical Monocytes
IL2R/IL7R+ Monocytes
Intermediate Monocytes
Non-classical Monocytes
mDCs
pDCs
CD45
125TeLEV
CD3
CD4
CD8a
CD11c
CD14
CD16
CD19
CD20
CD25
CD27
CD28
CD38
CD45RA
CD45RO
CD47
CD56
CD57
CD66b
CD69
CD123
CD127
CD161
CD183
CD185
CD194
CD196
CD197
CD294
HLA-DR
IgD
TCRgd
BCL-2
BTK
Ki-67
LYN
PI3K
yH2AX
IdU
DNA1
DNA2

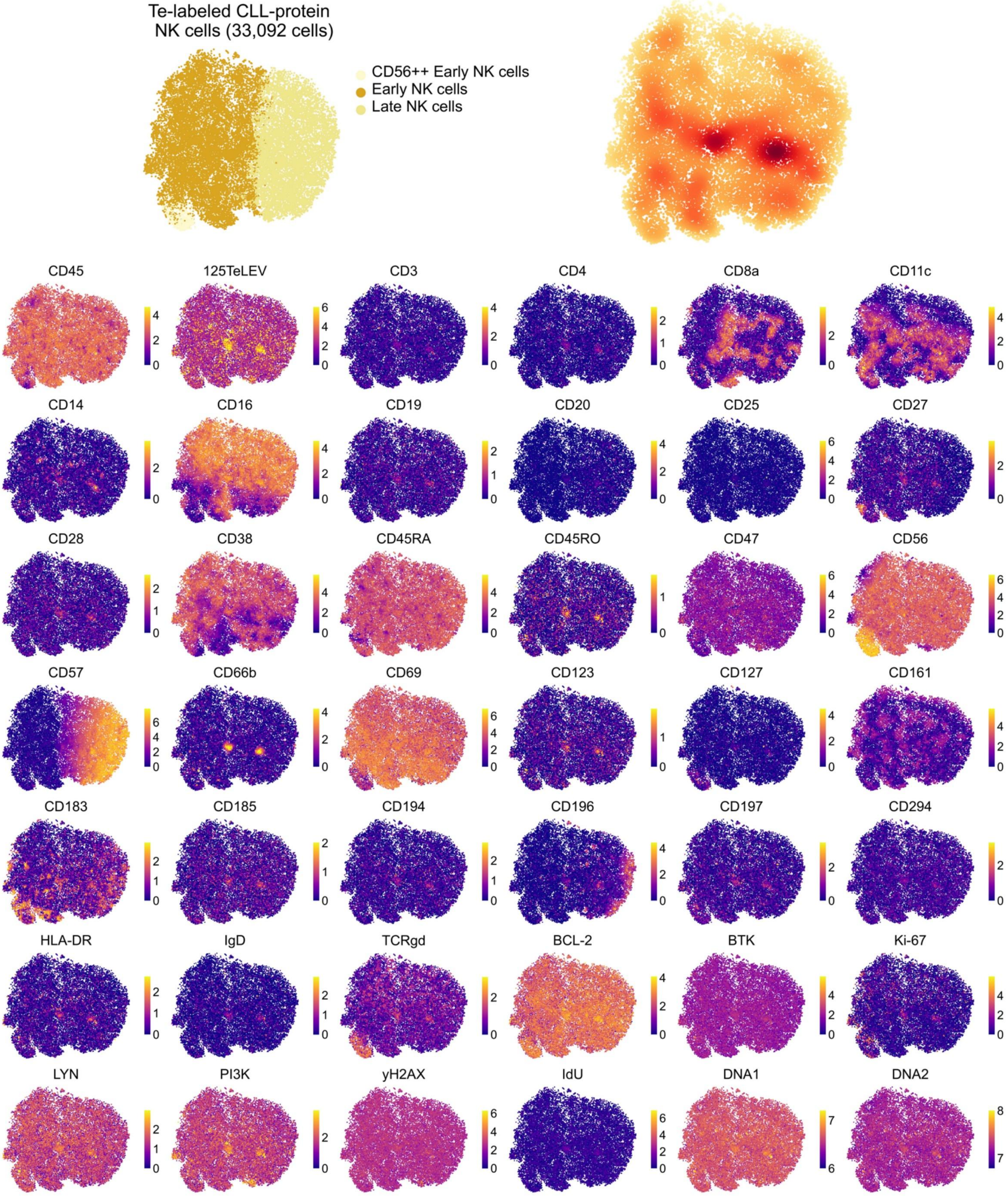

Te-labeled CLL-protein
NK cells (33,092 cells)
CD56++ Early NK cells
Early NK cells
Late NK cells
CD45
125TeLEV
CD3
CD4
CD8a
CD11c
CD14
CD16
CD19
CD20
CD25
CD27
CD28
CD38
CD45RA
CD45RO
CD47
CD56
CD57
CD66b
CD69
CD123
CD127
CD161
CD183
CD185
CD194
CD196
CD197
CD294
HLA-DR
IgD
TCRgd
BCL-2
BTK
Ki-67
LYN
PI3K
yH2AX
IdU
DNA1
DNA2

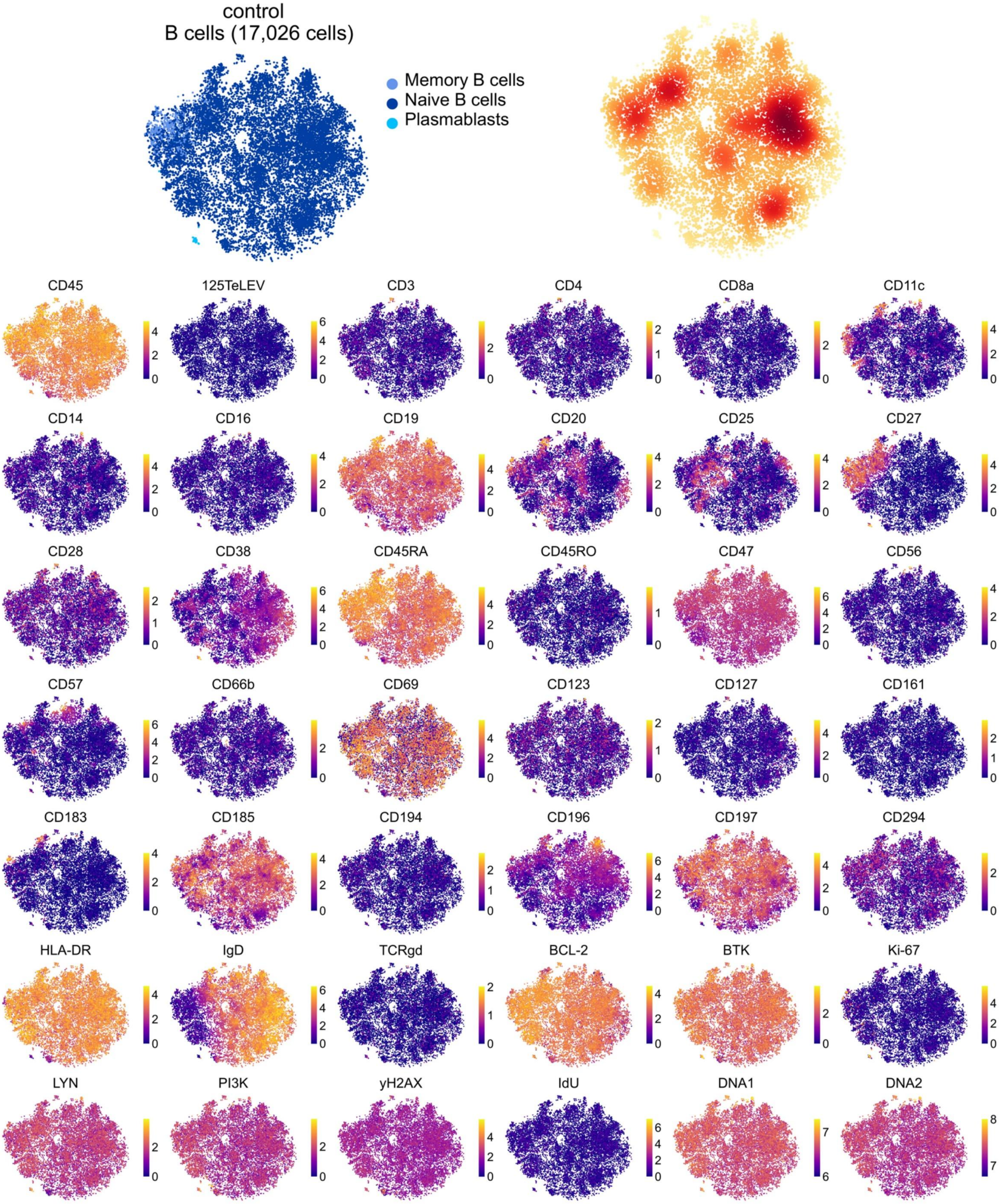
control
B cells (17,026 cells)
Memory B cells
Naive B cells
Plasmablasts
CD45
125TeLEV
CD3
CD4
CD8a
CD11c
CD14
CD16
CD19
CD20
CD25
CD27
CD28
CD38
CD45RA
CD45RO
CD47
CD56
CD57
CD66b
CD69
CD123
CD127
CD161
CD183
CD185
CD194
CD196
CD197
CD294
HLA-DR
IgD
TCRgd
BCL-2
BTK
Ki-67
LYN
PI3K
yH2AX
IdU
DNA1
DNA2

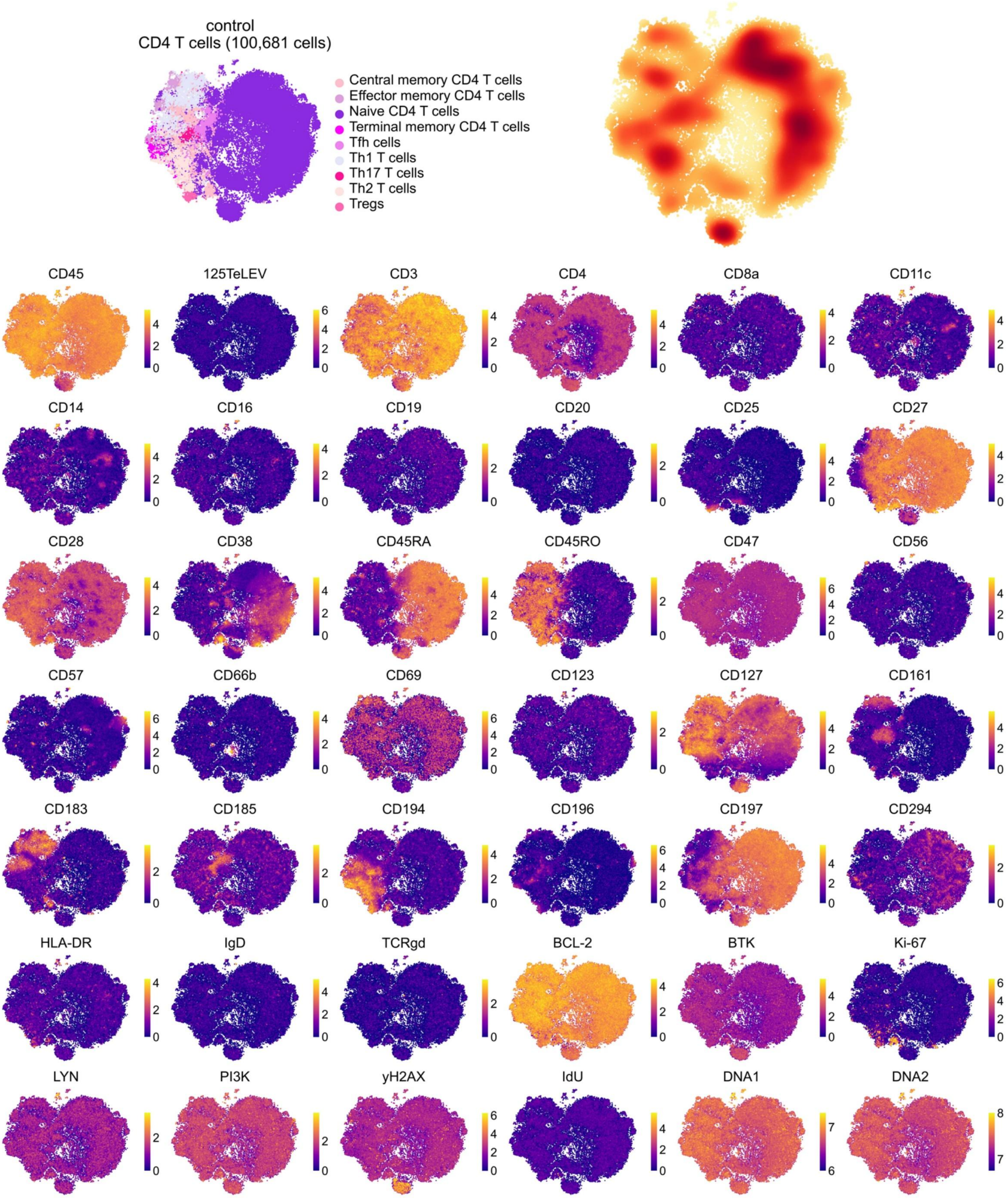
control
CD4 T cells (100,681 cells)
Central memory CD4 T cells
Effector memory CD4 T cells
Naive CD4 T cells
Terminal memory CD4 T cells
Tfh cells
Th1 T cells
Th17 T cells
Th2 T cells
Tregs
CD45
125TeLEV
CD3
CD4
CD8a
CD11c
CD14
CD16
CD19
CD20
CD25
CD27
CD28
CD38
CD45RA
CD45RO
CD47
CD56
CD57
CD66b
CD69
CD123
CD127
CD161
CD183
CD185
CD194
CD196
CD197
CD294
HLA-DR
IgD
TCRgd
BCL-2
BTK
Ki-67
LYN
PI3K
yH2AX
IdU
DNA1
DNA2

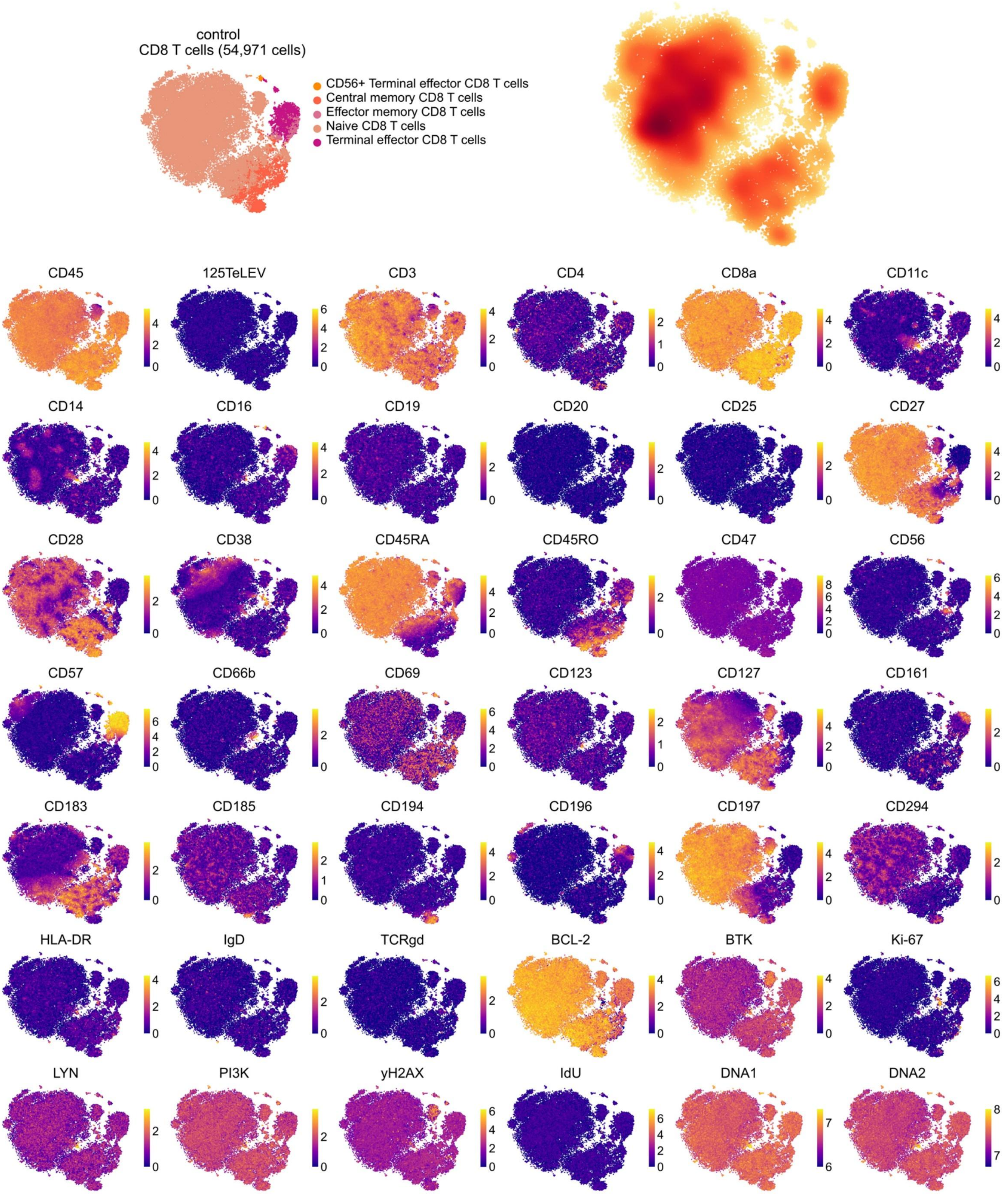
control
CD8 T cells (54,971 cells)
CD56+ Terminal effector CD8 T cells
Central memory CD8 T cells
Effector memory CD8 T cells
Naive CD8 T cells
Terminal effector CD8 T cells
CD45
125TeLEV
CD3
CD4
CD8a
CD11c
CD14
CD16
CD19
CD20
CD25
CD27
CD28
CD38
CD45RA
CD45RO
CD47
CD56
CD57
CD66b
CD69
CD123
CD127
CD161
CD183
CD185
CD194
CD196
CD197
CD294
HLA-DR
IgD
TCRgd
BCL-2
BTK
Ki-67
LYN
PI3K
yH2AX
IdU
DNA1
DNA2

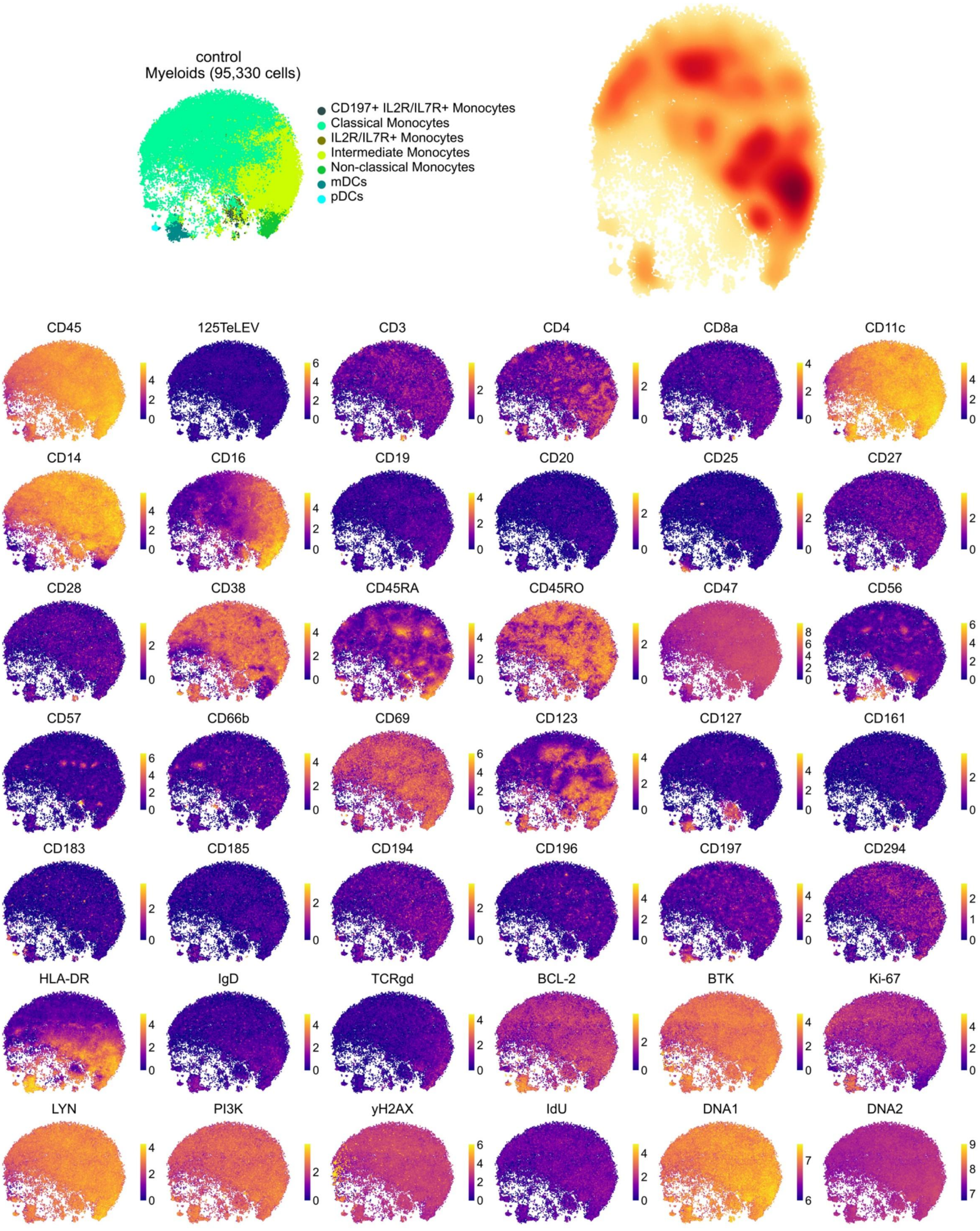
control
Myeloids (95,330 cells)
CD197+ IL2R/IL7R+ Monocytes
Classical Monocytes
IL2R/IL7R+ Monocytes
Intermediate Monocytes
Non-classical Monocytes
mDCs
pDCs
CD45
125TeLEV
CD3
CD4
CD8a
CD11c
CD14
CD16
CD19
CD20
CD25
CD27
CD28
CD38
CD45RA
CD45RO
CD47
CD56
CD57
CD66b
CD69
CD123
CD127
CD161
CD183
CD185
CD194
CD196
CD197
CD294
HLA-DR
IgD
TCRgd
BCL-2
BTK
Ki-67
LYN
PI3K
yH2AX
IdU
DNA1
DNA2

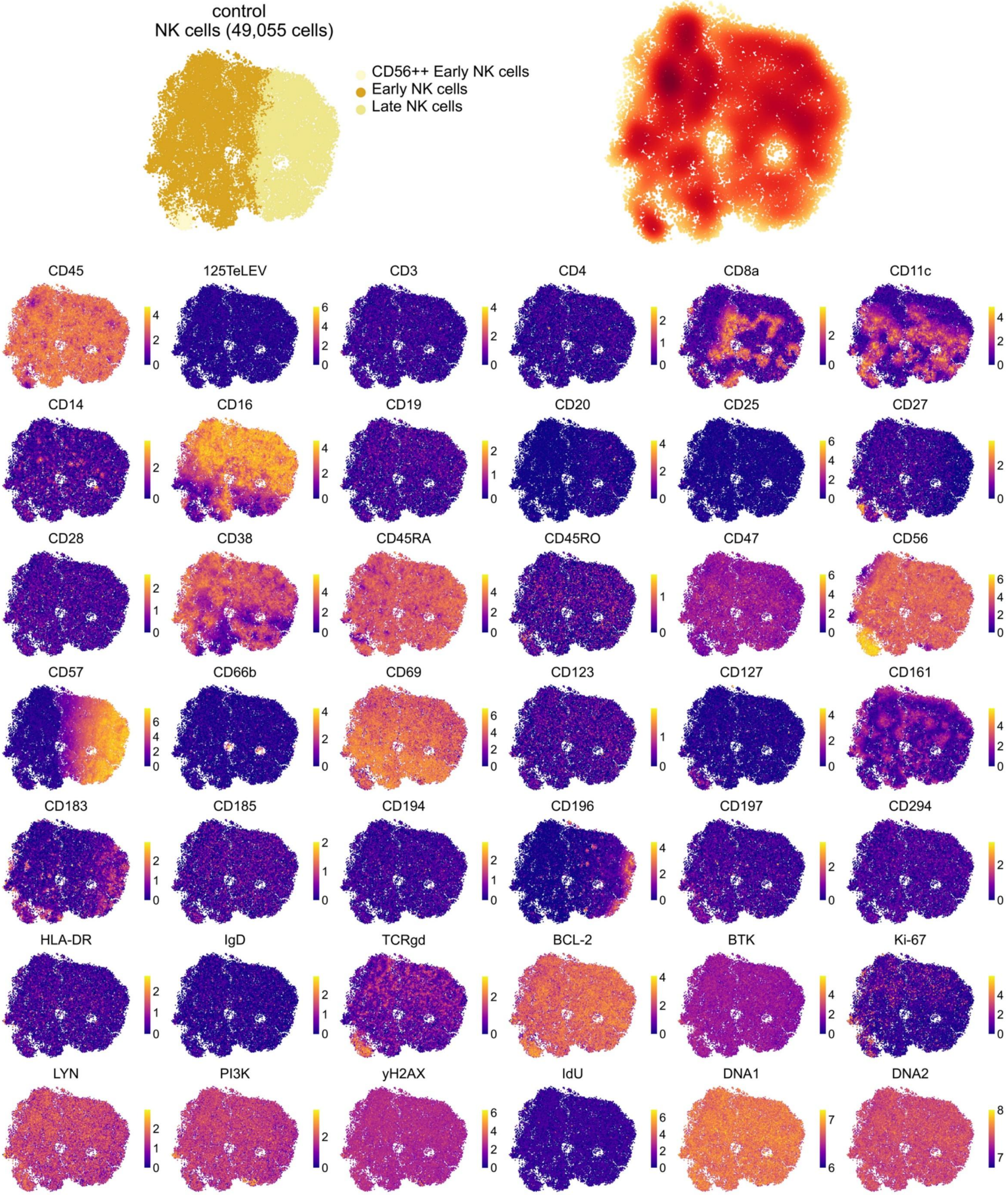
control
NK cells (49,055 cells)
CD56++ Early NK cells
Early NK cells
Late NK cells
CD45
125TeLEV
CD3
CD4
CD8a
CD11c
CD14
CD16
CD19
CD20
CD25
CD27
CD28
CD38
CD45RA
CD45RO
CD47
CD56
CD57
CD66b
CD69
CD123
CD127
CD161
CD183
CD185
CD194
CD196
CD197
CD294
HLA-DR
IgD
TCRgd
BCL-2
BTK
Ki-67
LYN
PI3K
yH2AX
IdU
DNA1
DNA2
