## Supplementary Figure 2 for "Single recipient cell tracking of tellurium-labeled extracellular vesicle proteomes (TeLEV) identifies EV-driven immunomodulation"

**Supplementary Fig. 2**

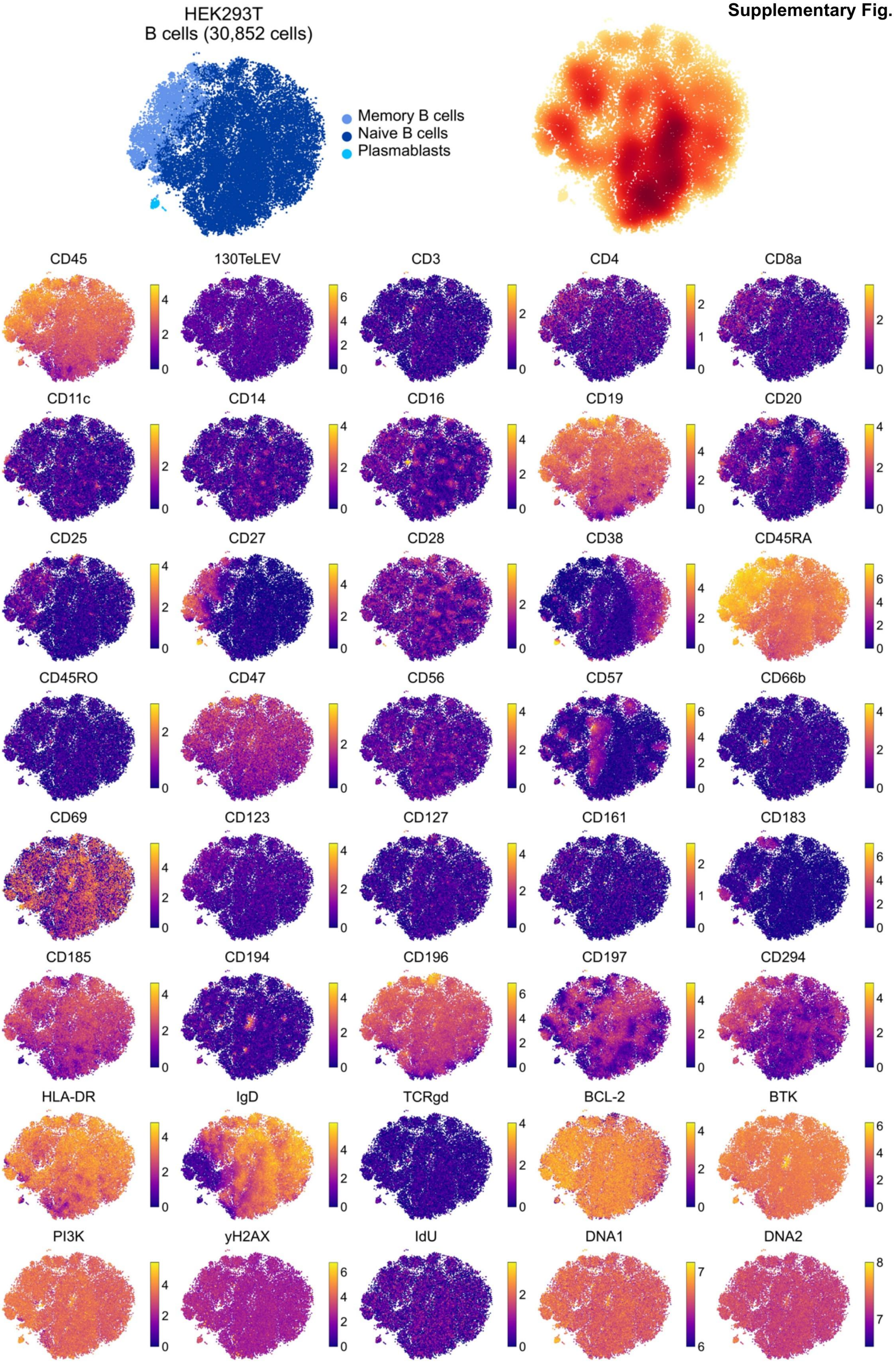

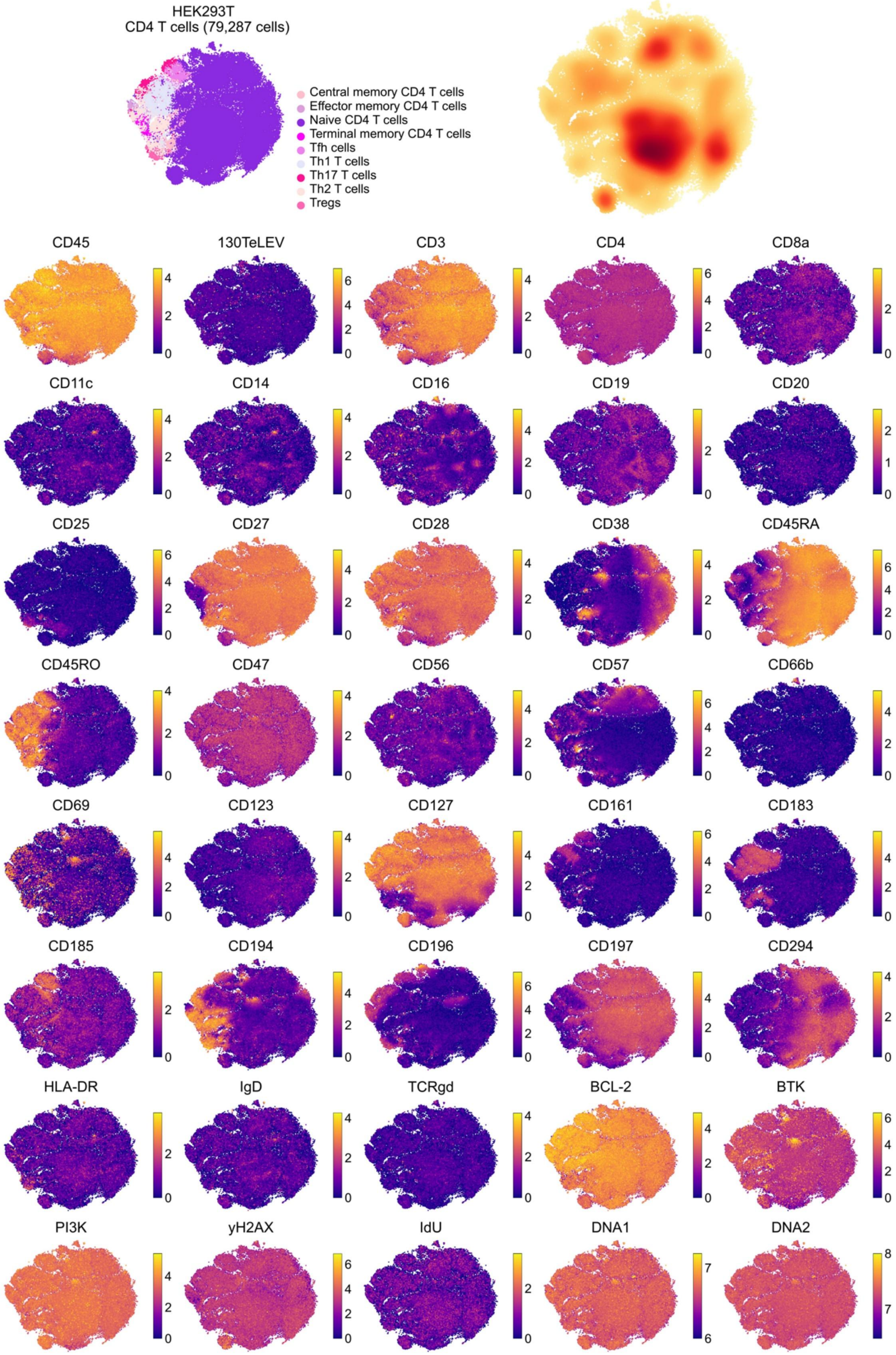
HEK293T
CD4 T cells (79,287 cells)
Central memory CD4 T cells
Effector memory CD4 T cells
Naive CD4 T cells
Terminal memory CD4 T cells
Tfh cells
Th1 T cells
Th17 T cells
Th2 T cells
Tregs
CD45
130TeLEV
CD3
CD4
CD8a
CD11c
CD14
CD16
CD19
CD20
CD25
CD27
CD28
CD38
CD45RA
CD45RO
CD47
CD56
CD57
CD66b
CD69
CD123
CD127
CD161
CD183
CD185
CD194
CD196
CD197
CD294
HLA-DR
IgD
TCRgd
BCL-2
BTK
PI3K
yH2AX
IdU
DNA1
DNA2

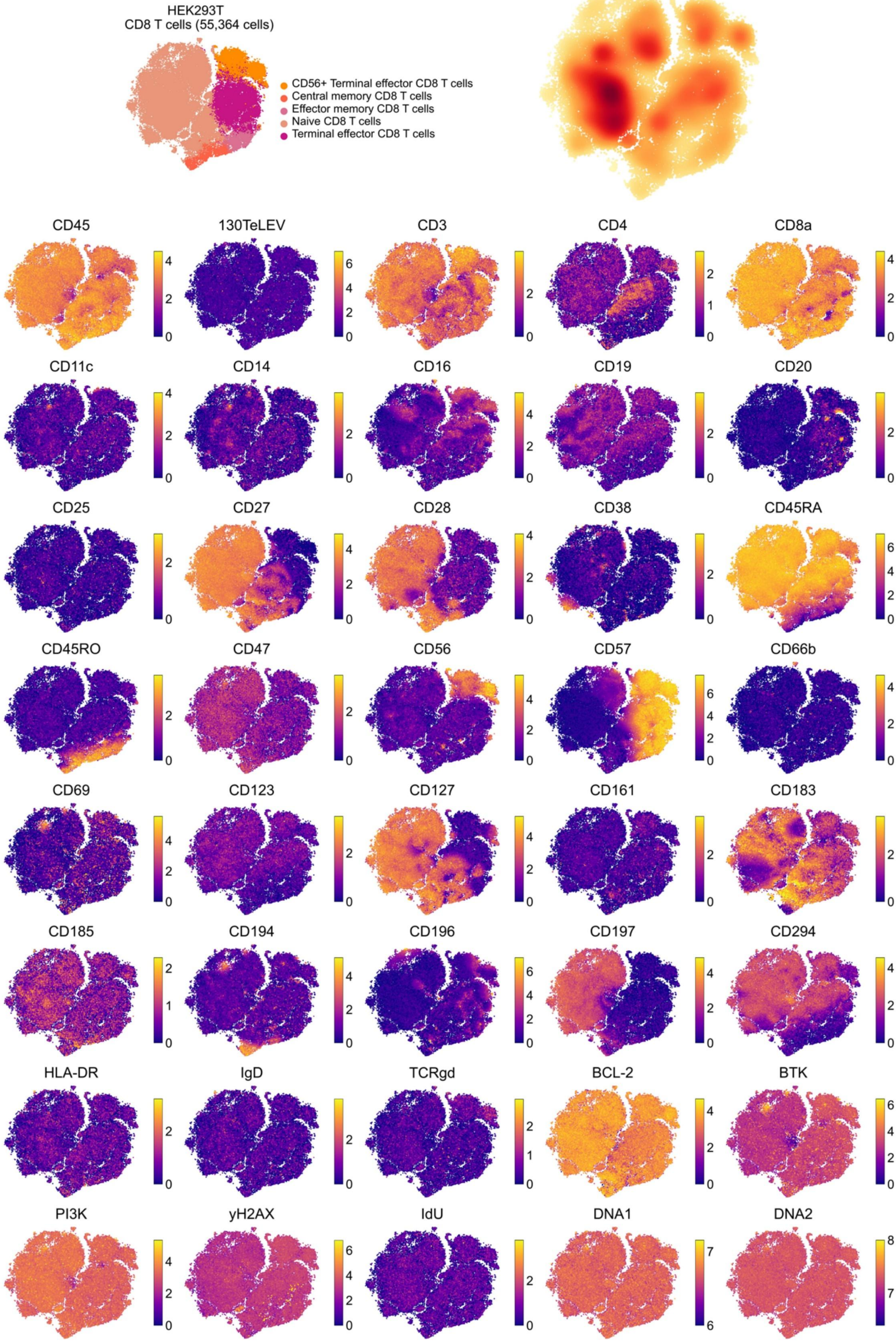
HEK293T
CD8 T cells (55,364 cells)
CD56+ Terminal effector CD8 T cells
Central memory CD8 T cells
Effector memory CD8 T cells
Naive CD8 T cells
Terminal effector CD8 T cells
CD45
130TeLEV
CD3
CD4
CD8a
CD11c
CD14
CD16
CD19
CD20
CD25
CD27
CD28
CD38
CD45RA
CD45RO
CD47
CD56
CD57
CD66b
CD69
CD123
CD127
CD161
CD183
CD185
CD194
CD196
CD197
CD294
HLA-DR
IgD
TCRgd
BCL-2
BTK
PI3K
yH2AX
IdU
DNA1
DNA2

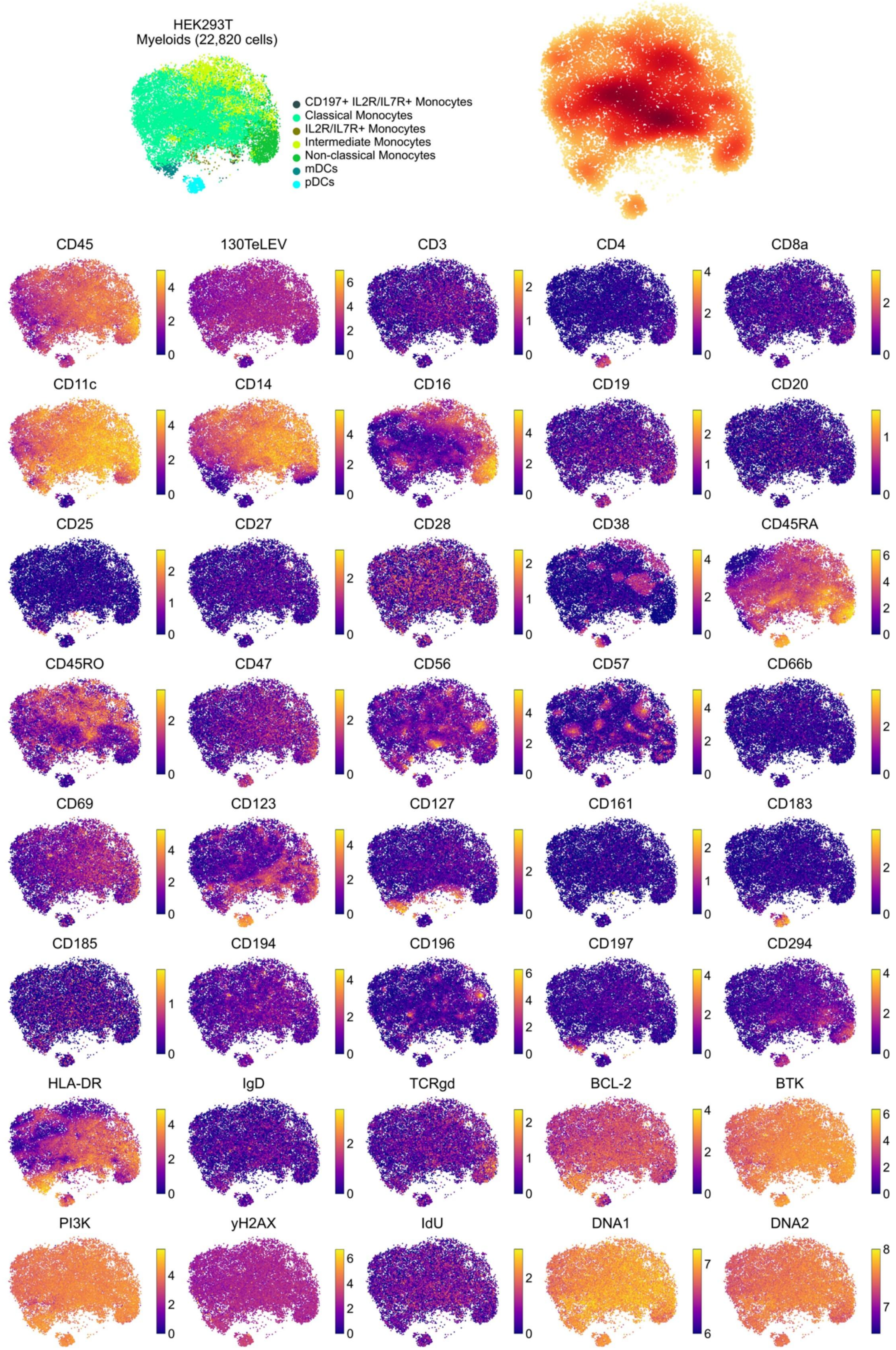
HEK293T
Myeloids (22,820 cells)
CD197+ IL2R/IL7R+ Monocytes
Classical Monocytes
IL2R/IL7R+ Monocytes
Intermediate Monocytes
Non-classical Monocytes
mDCs
pDCs
CD45
130TeLEV
CD3
CD4
CD8a
CD11c
CD14
CD16
CD19
CD20
CD25
CD27
CD28
CD38
CD45RA
CD45RO
CD47
CD56
CD57
CD66b
CD69
CD123
CD127
CD161
CD183
CD185
CD194
CD196
CD197
CD294
HLA-DR
IgD
TCRgd
BCL-2
BTK
PI3K
yH2AX
IdU
DNA1
DNA2

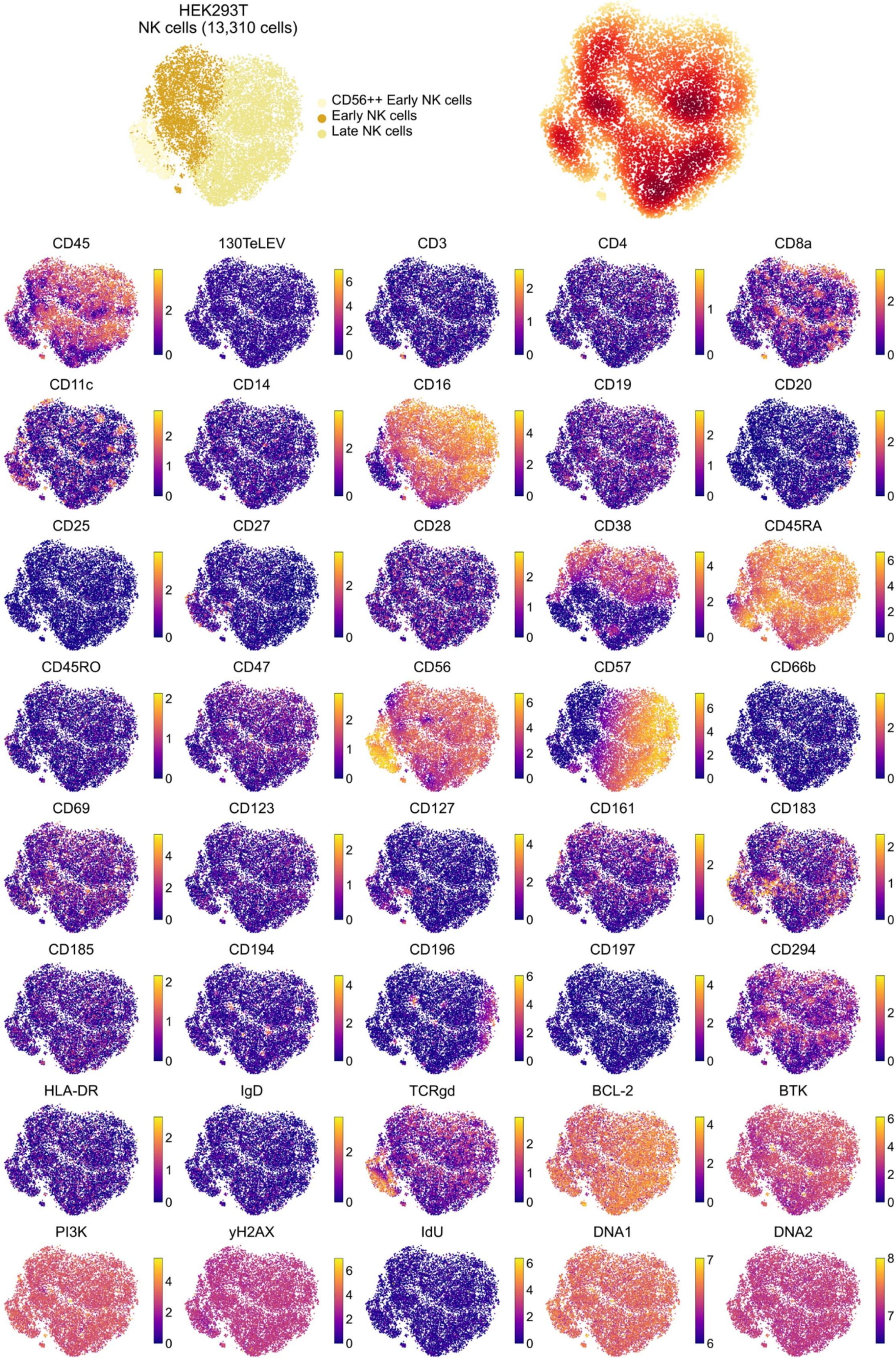

HEK293T
NK cells (13,310 cells)
CD56++ Early NK cells
Early NK cells
Late NK cells
CD45
130TeLEV
CD3
CD4
CD8a
CD11c
CD14
CD16
CD19
CD20
CD25
CD27
CD28
CD38
CD45RA
CD45RO
CD47
CD56
CD57
CD66b
CD69
CD123
CD127
CD161
CD183
CD185
CD194
CD196
CD197
CD294
HLA-DR
IgD
TCRgd
BCL-2
BTK
PI3K
yH2AX
IdU
DNA1
DNA2

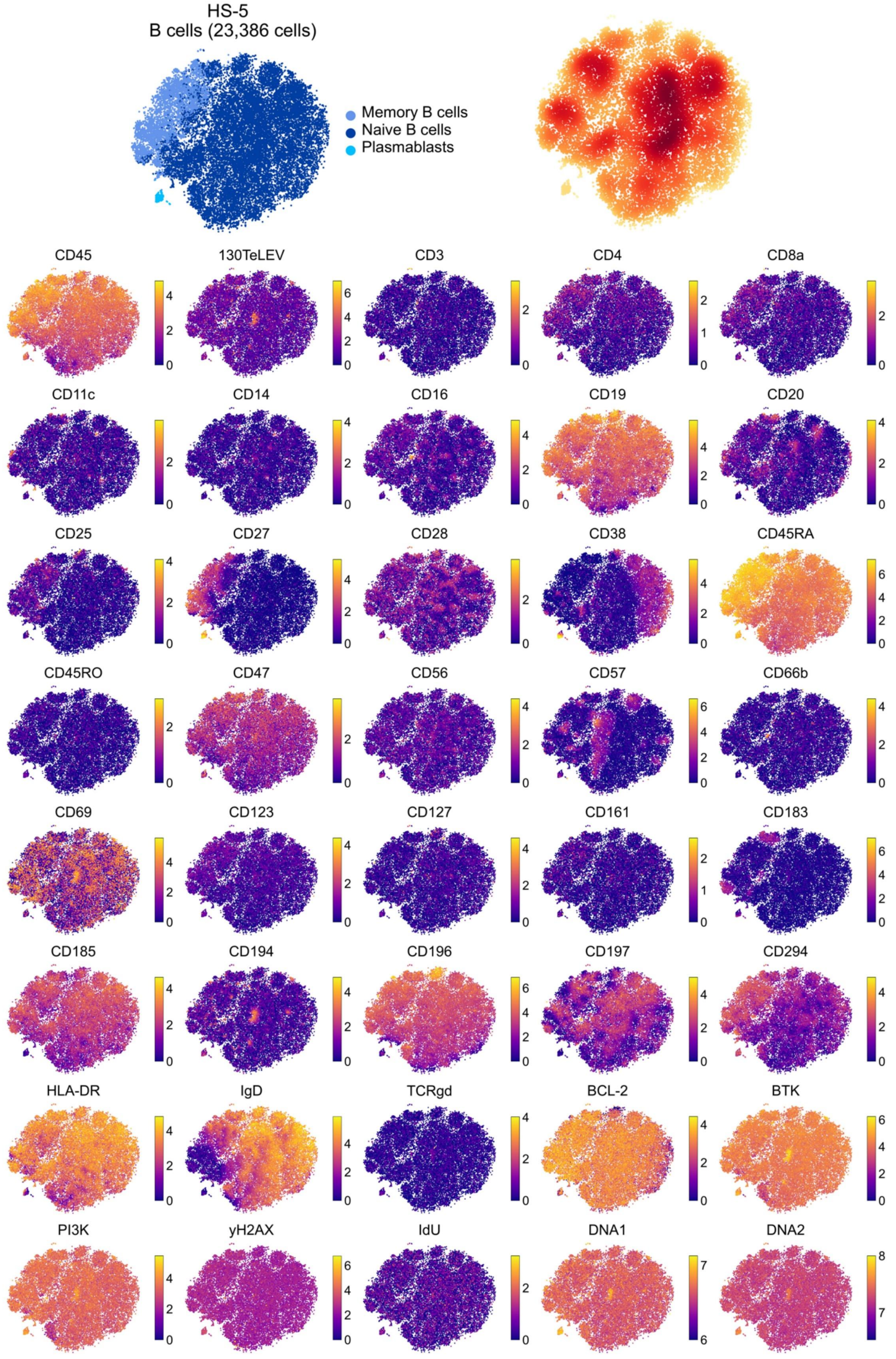
HS-5
B cells (23,386 cells)
Memory B cells
Naive B cells
Plasmablasts
CD45
130TeLEV
CD3
CD4
CD8a
CD11c
CD14
CD16
CD19
CD20
CD25
CD27
CD28
CD38
CD45RA
CD45RO
CD47
CD56
CD57
CD66b
CD69
CD123
CD127
CD161
CD183
CD185
CD194
CD196
CD197
CD294
HLA-DR
IgD
TCRgd
BCL-2
BTK
PI3K
yH2AX
IdU
DNA1
DNA2

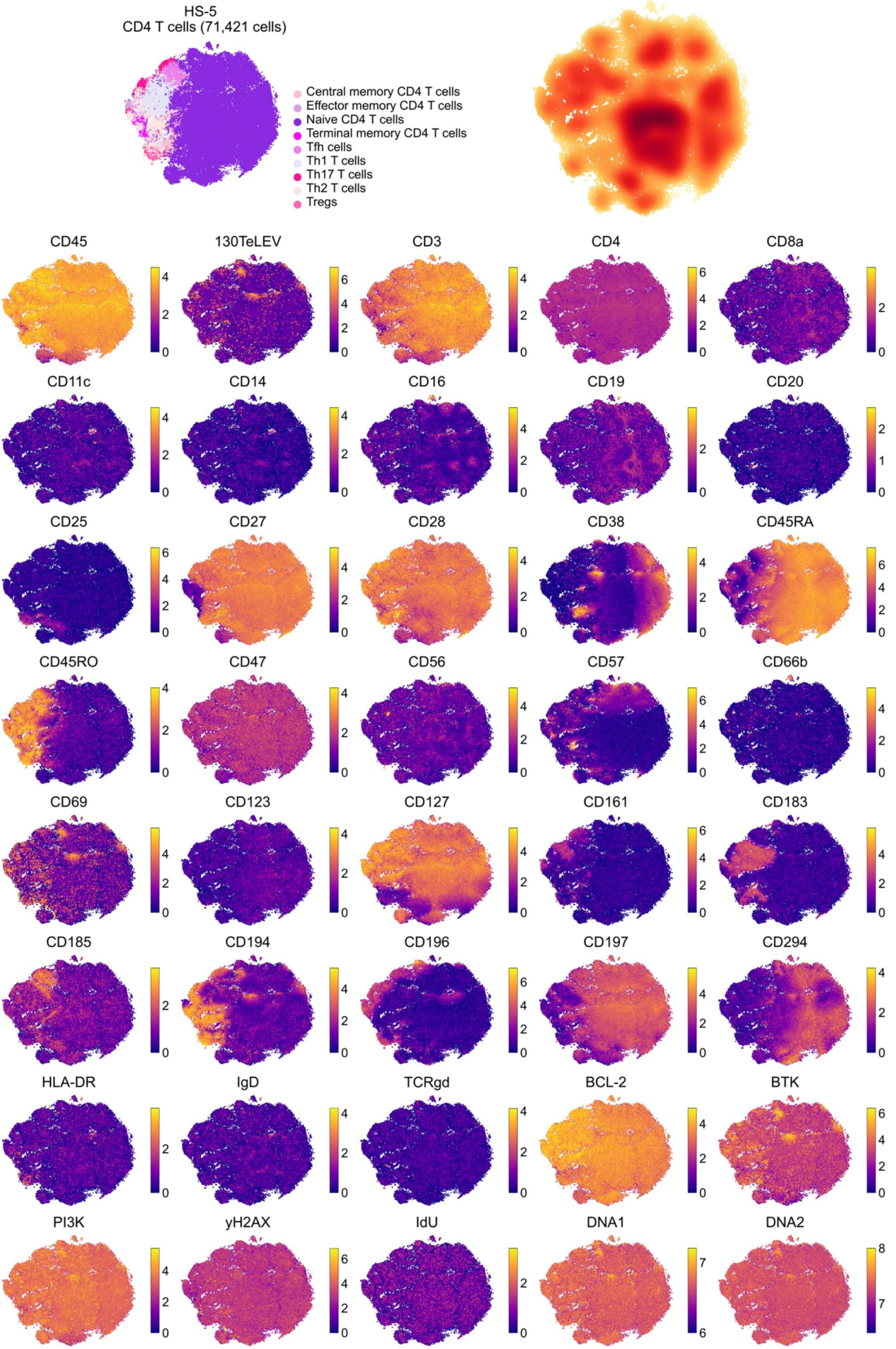

HS-5
CD4 T cells (71,421 cells)
Central memory CD4 T cells
Effector memory CD4 T cells
Naive CD4 T cells
Terminal memory CD4 T cells
Tfh cells
Th1 T cells
Th17 T cells
Th2 T cells
Tregs
CD45
130TeLEV
CD3
CD4
CD8a
CD11c
CD14
CD16
CD19
CD20
CD25
CD27
CD28
CD38
CD45RA
CD45RO
CD47
CD56
CD57
CD66b
CD69
CD123
CD127
CD161
CD183
CD185
CD194
CD196
CD197
CD294
HLA-DR
IgD
TCRgd
BCL-2
BTK
PI3K
yH2AX
IdU
DNA1
DNA2

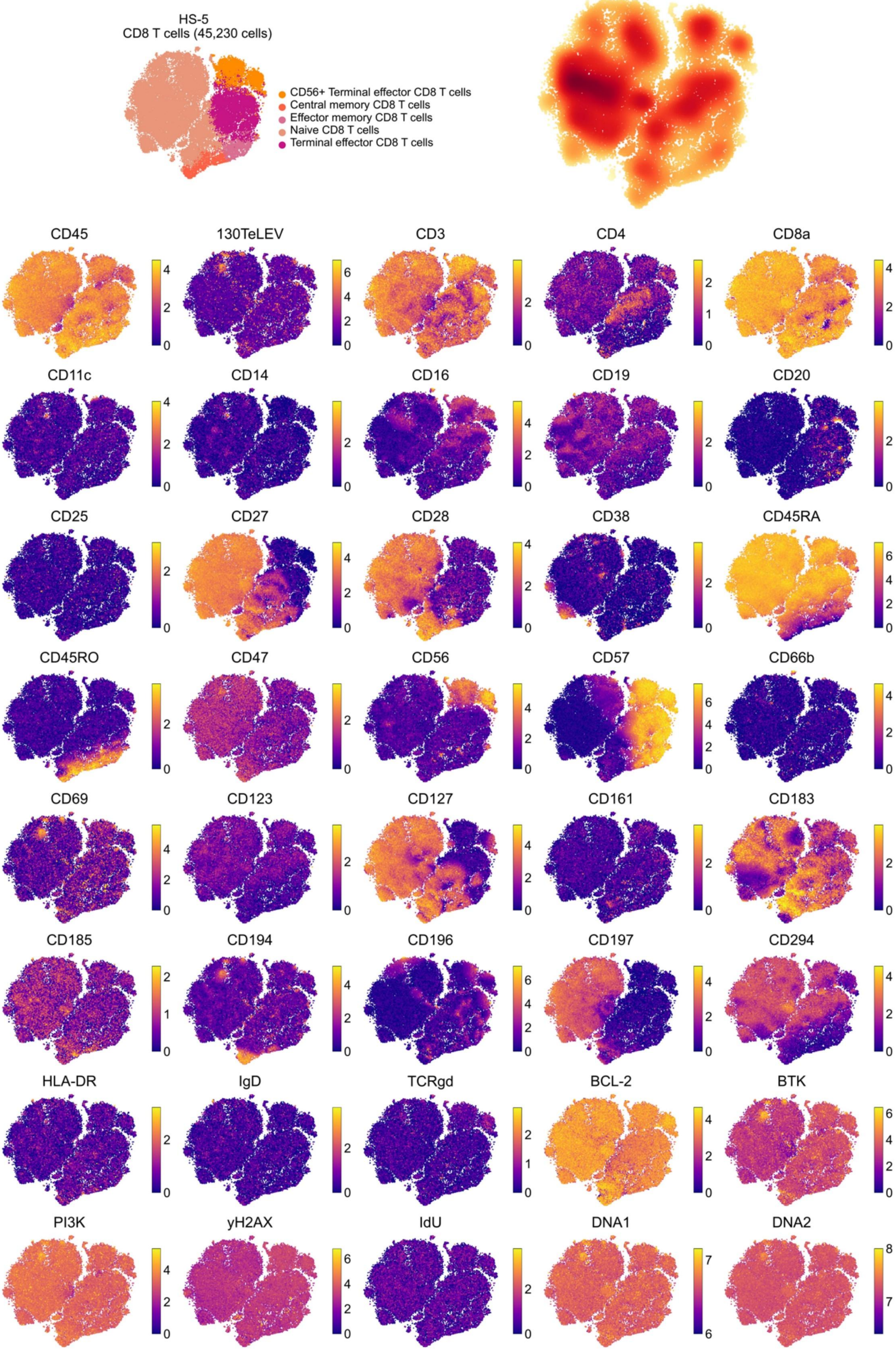
HS-5
CD8 T cells (45,230 cells)
CD56+ Terminal effector CD8 T cells
Central memory CD8 T cells
Effector memory CD8 T cells
Naive CD8 T cells
Terminal effector CD8 T cells
CD45
130TeLEV
CD3
CD4
CD8a
CD11c
CD14
CD16
CD19
CD20
CD25
CD27
CD28
CD38
CD45RA
CD45RO
CD47
CD56
CD57
CD66b
CD69
CD123
CD127
CD161
CD183
CD185
CD194
CD196
CD197
CD294
HLA-DR
IgD
TCRgd
BCL-2
BTK
PI3K
yH2AX
IdU
DNA1
DNA2

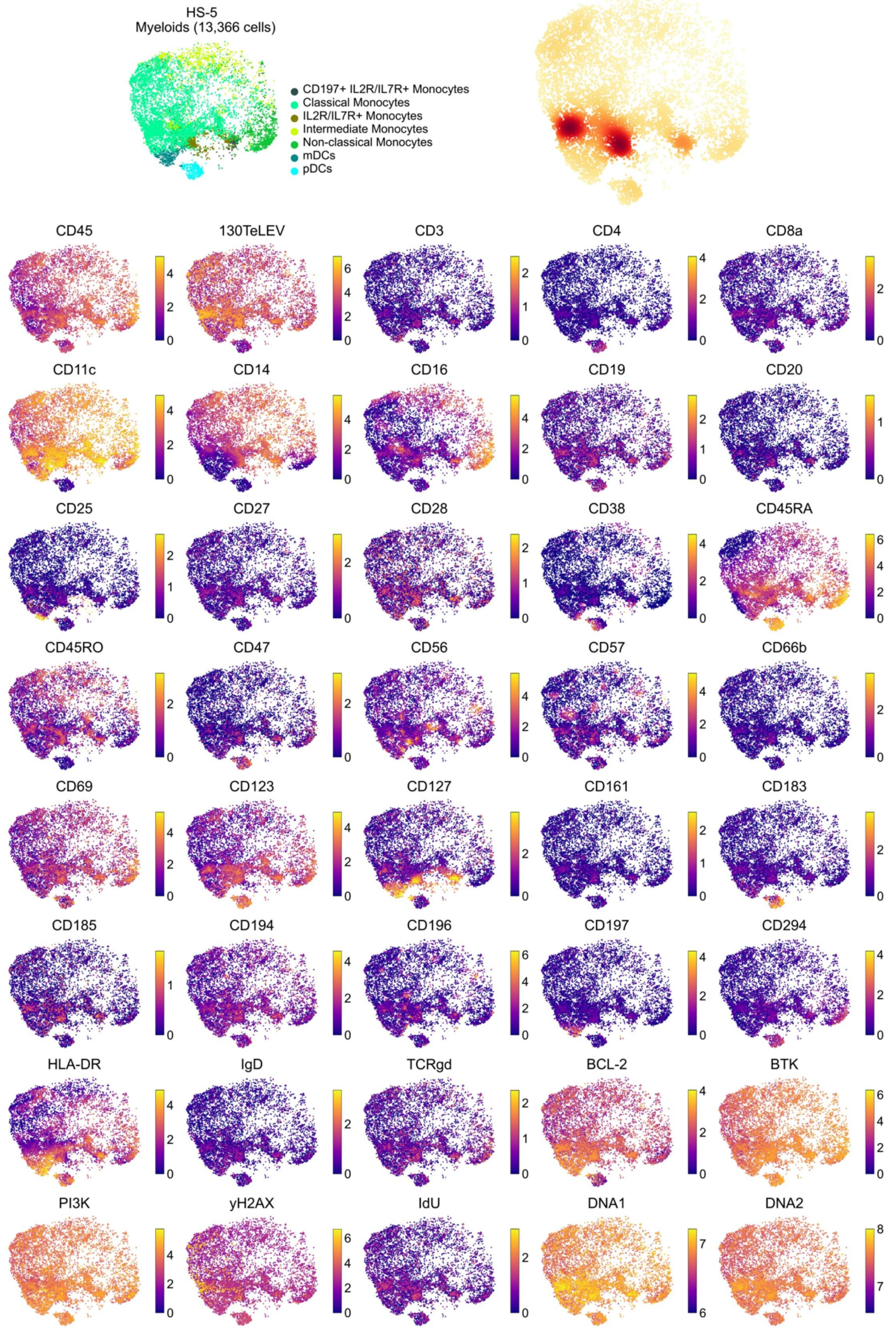

HS-5
Myeloids (13,366 cells)
CD197+ IL2R/IL7R+ Monocytes
Classical Monocytes
IL2R/IL7R+ Monocytes
Intermediate Monocytes
Non-classical Monocytes
mDCs
pDCs
CD45
130TeLEV
CD3
CD4
CD8a
CD11c
CD14
CD16
CD19
CD20
CD25
CD27
CD28
CD38
CD45RA
CD45RO
CD47
CD56
CD57
CD66b
CD69
CD123
CD127
CD161
CD183
CD185
CD194
CD196
CD197
CD294
HLA-DR
IgD
TCRgd
BCL-2
BTK
PI3K
yH2AX
IdU
DNA1
DNA2

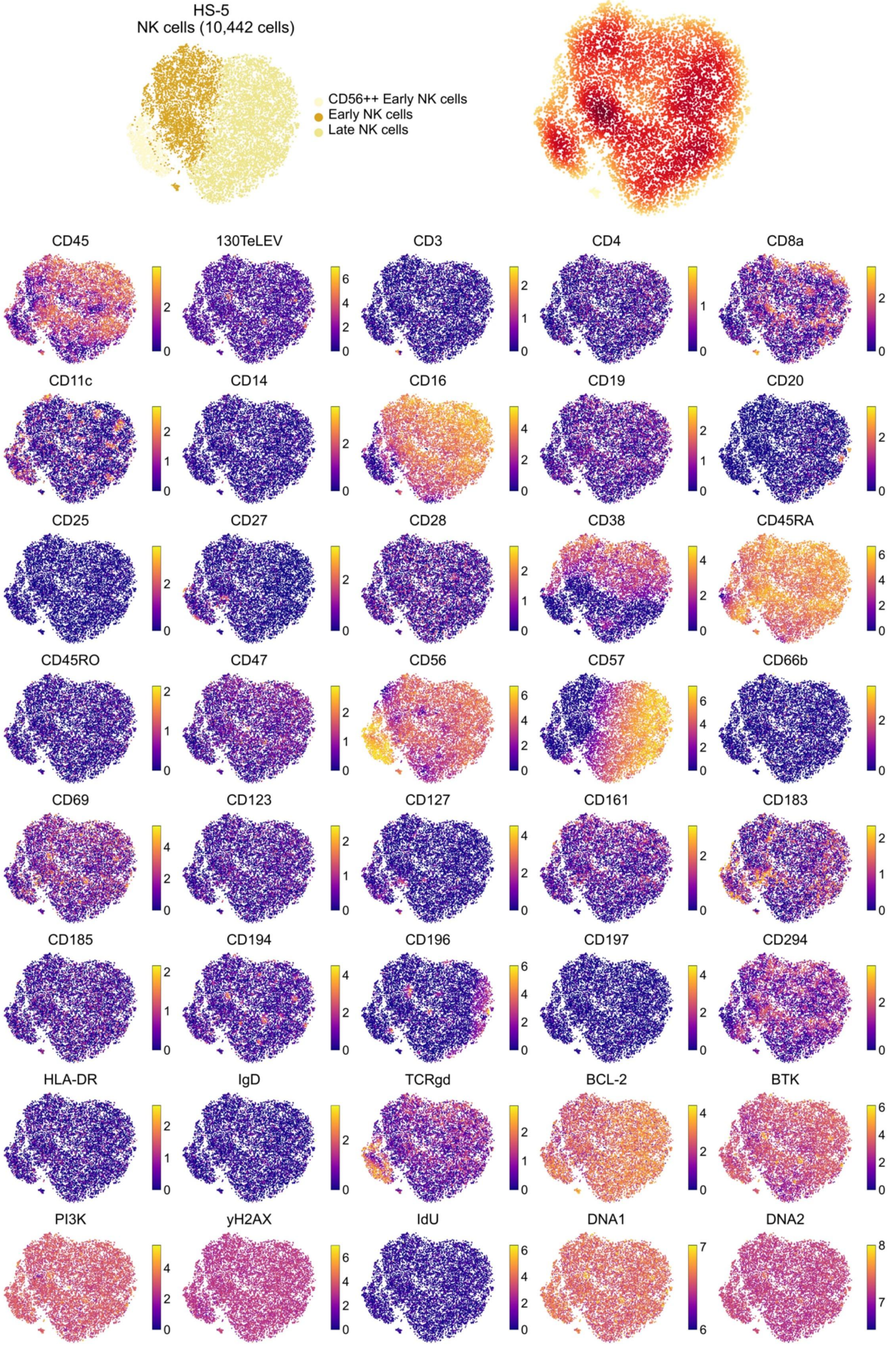
HS-5
NK cells (10,442 cells)
CD56++ Early NK cells
Early NK cells
Late NK cells
CD45
130TeLEV
CD3
CD4
CD8a
CD11c
CD14
CD16
CD19
CD20
CD25
CD27
CD28
CD38
CD45RA
CD45RO
CD47
CD56
CD57
CD66b
CD69
CD123
CD127
CD161
CD183
CD185
CD194
CD196
CD197
CD294
HLA-DR
IgD
TCRgd
BCL-2
BTK
PI3K
yH2AX
IdU
DNA1
DNA2

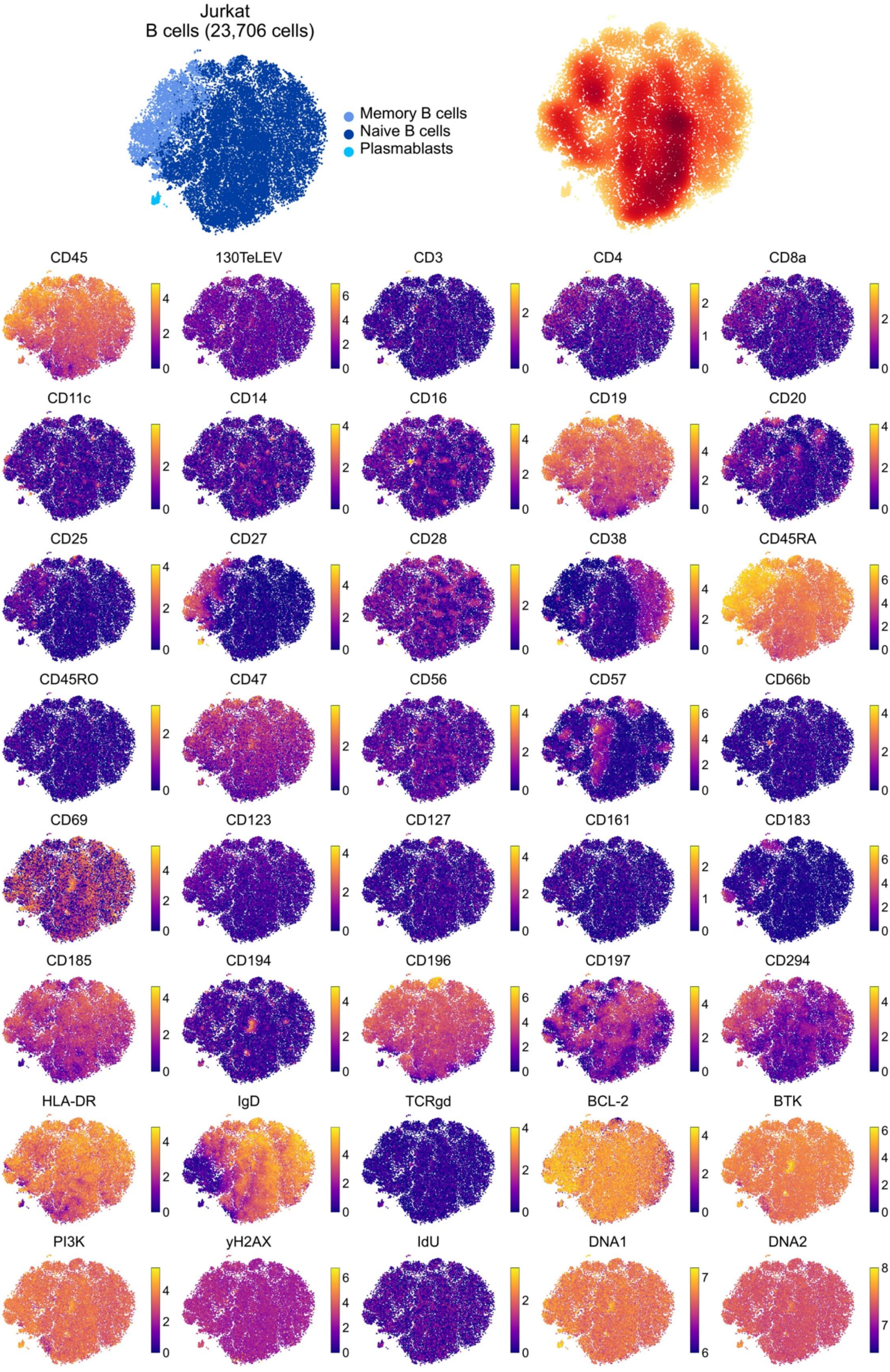

Jurkat
B cells (23,706 cells)
Memory B cells
Naive B cells
Plasmablasts
CD45
130TeLEV
CD3
CD4
CD8a
CD11c
CD14
CD16
CD19
CD20
CD25
CD27
CD28
CD38
CD45RA
CD45RO
CD47
CD56
CD57
CD66b
CD69
CD123
CD127
CD161
CD183
CD185
CD194
CD196
CD197
CD294
HLA-DR
IgD
TCRgd
BCL-2
BTK
PI3K
yH2AX
IdU
DNA1
DNA2

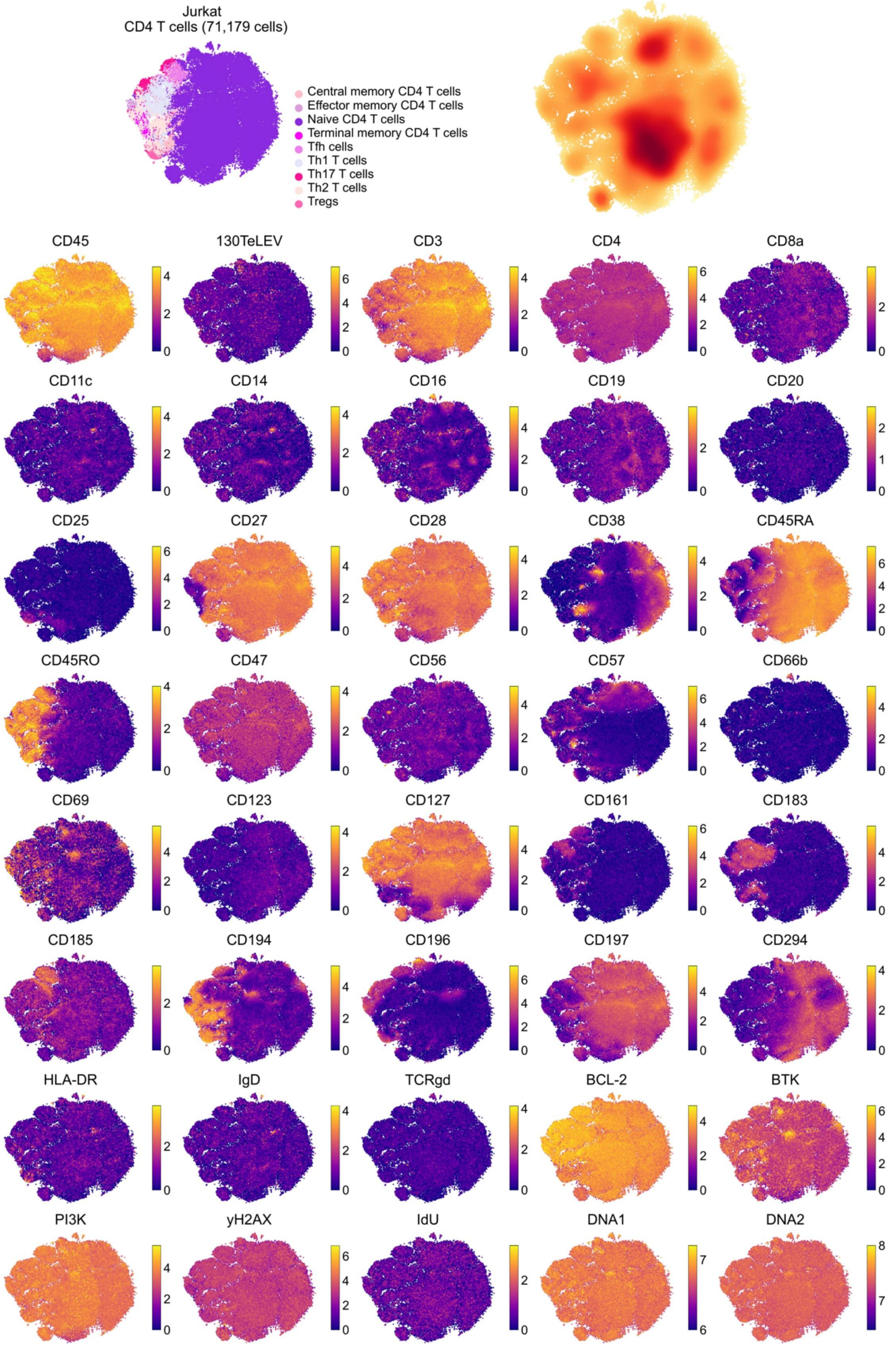
Jurkat
CD4 T cells (71,179 cells)
Central memory CD4 T cells
Effector memory CD4 T cells
Naive CD4 T cells
Terminal memory CD4 T cells
Tfh cells
Th1 T cells
Th17 T cells
Th2 T cells
Tregs
CD45
130TeLEV
CD3
CD4
CD8a
CD11c
CD14
CD16
CD19
CD20
CD25
CD27
CD28
CD38
CD45RA
CD45RO
CD47
CD56
CD57
CD66b
CD69
CD123
CD127
CD161
CD183
CD185
CD194
CD196
CD197
CD294
HLA-DR
IgD
TCRgd
BCL-2
BTK
PI3K
yH2AX
IdU
DNA1
DNA2

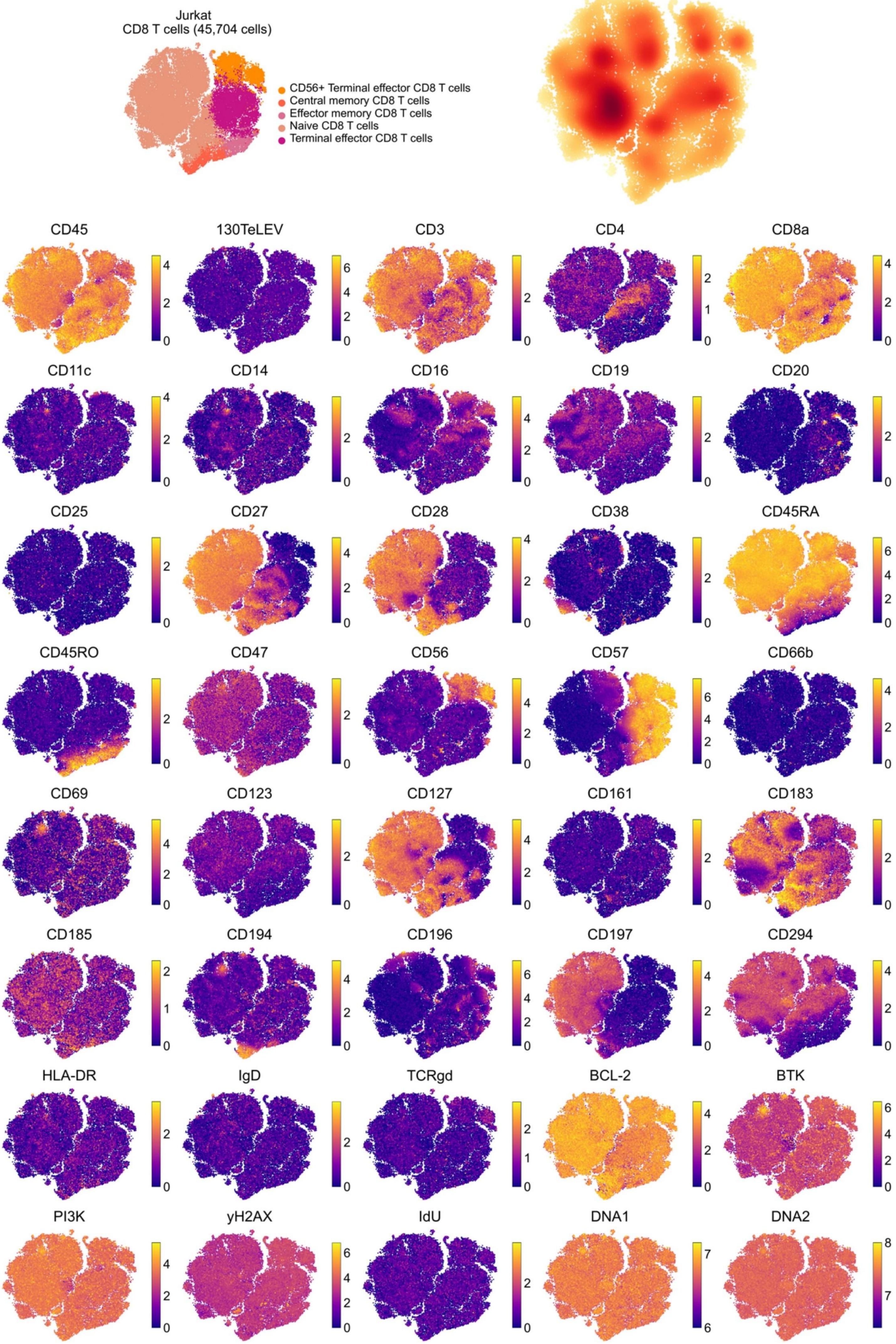
Jurkat
CD8 T cells (45,704 cells)
CD56+ Terminal effector CD8 T cells
Central memory CD8 T cells
Effector memory CD8 T cells
Naive CD8 T cells
Terminal effector CD8 T cells
CD45
130TeLEV
CD3
CD4
CD8a
CD11c
CD14
CD16
CD19
CD20
CD25
CD27
CD28
CD38
CD45RA
CD45RO
CD47
CD56
CD57
CD66b
CD69
CD123
CD127
CD161
CD183
CD185
CD194
CD196
CD197
CD294
HLA-DR
IgD
TCRgd
BCL-2
BTK
PI3K
yH2AX
IdU
DNA1
DNA2

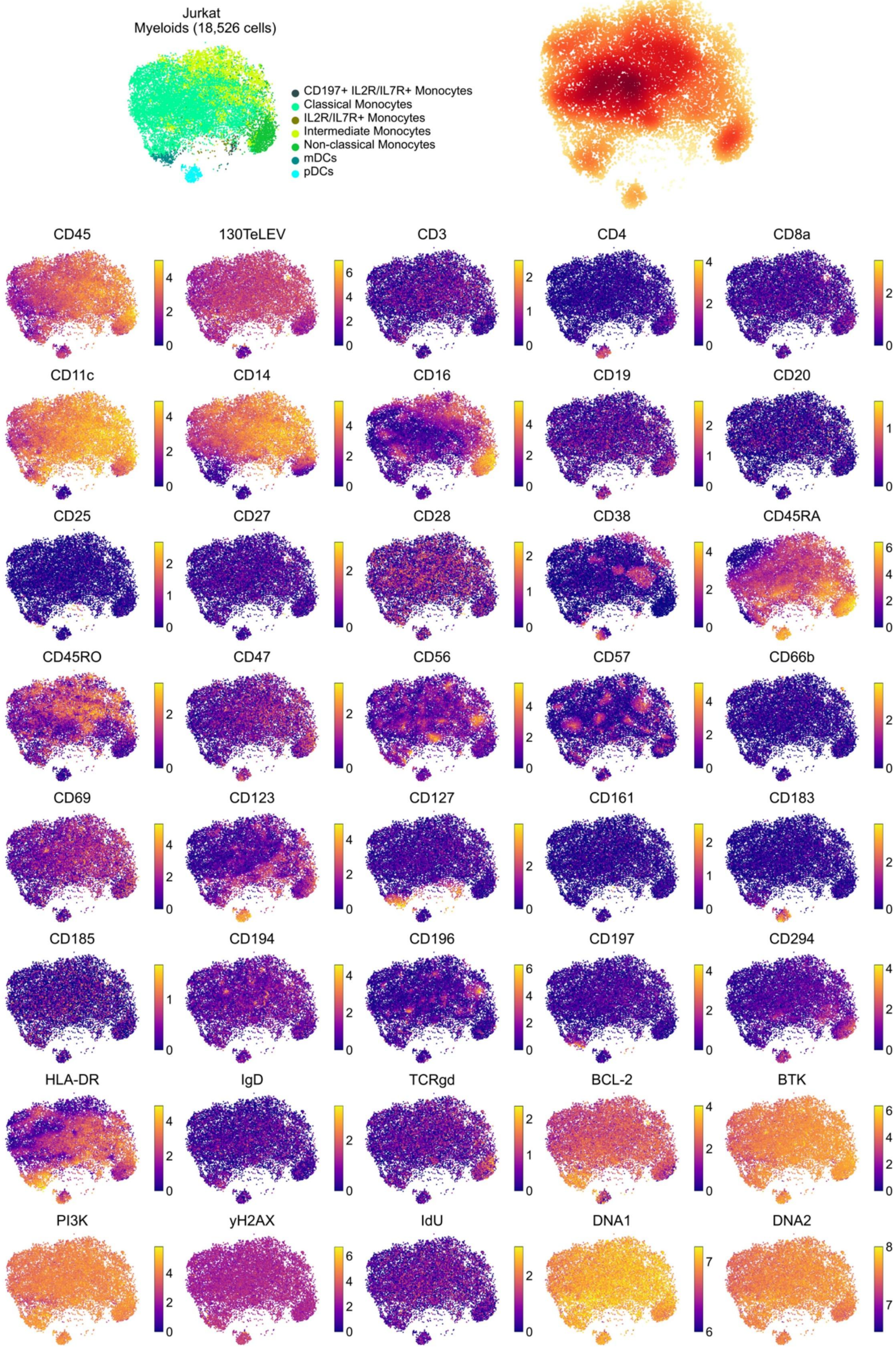

Jurkat
Myeloids (18,526 cells)
CD197+ IL2R/IL7R+ Monocytes
Classical Monocytes
IL2R/IL7R+ Monocytes
Intermediate Monocytes
Non-classical Monocytes
mDCs
pDCs
CD45
130TeLEV
CD3
CD4
CD8a
CD11c
CD14
CD16
CD19
CD20
CD25
CD27
CD28
CD38
CD45RA
CD45RO
CD47
CD56
CD57
CD66b
CD69
CD123
CD127
CD161
CD183
CD185
CD194
CD196
CD197
CD294
HLA-DR
IgD
TCRgd
BCL-2
BTK
PI3K
yH2AX
IdU
DNA1
DNA2

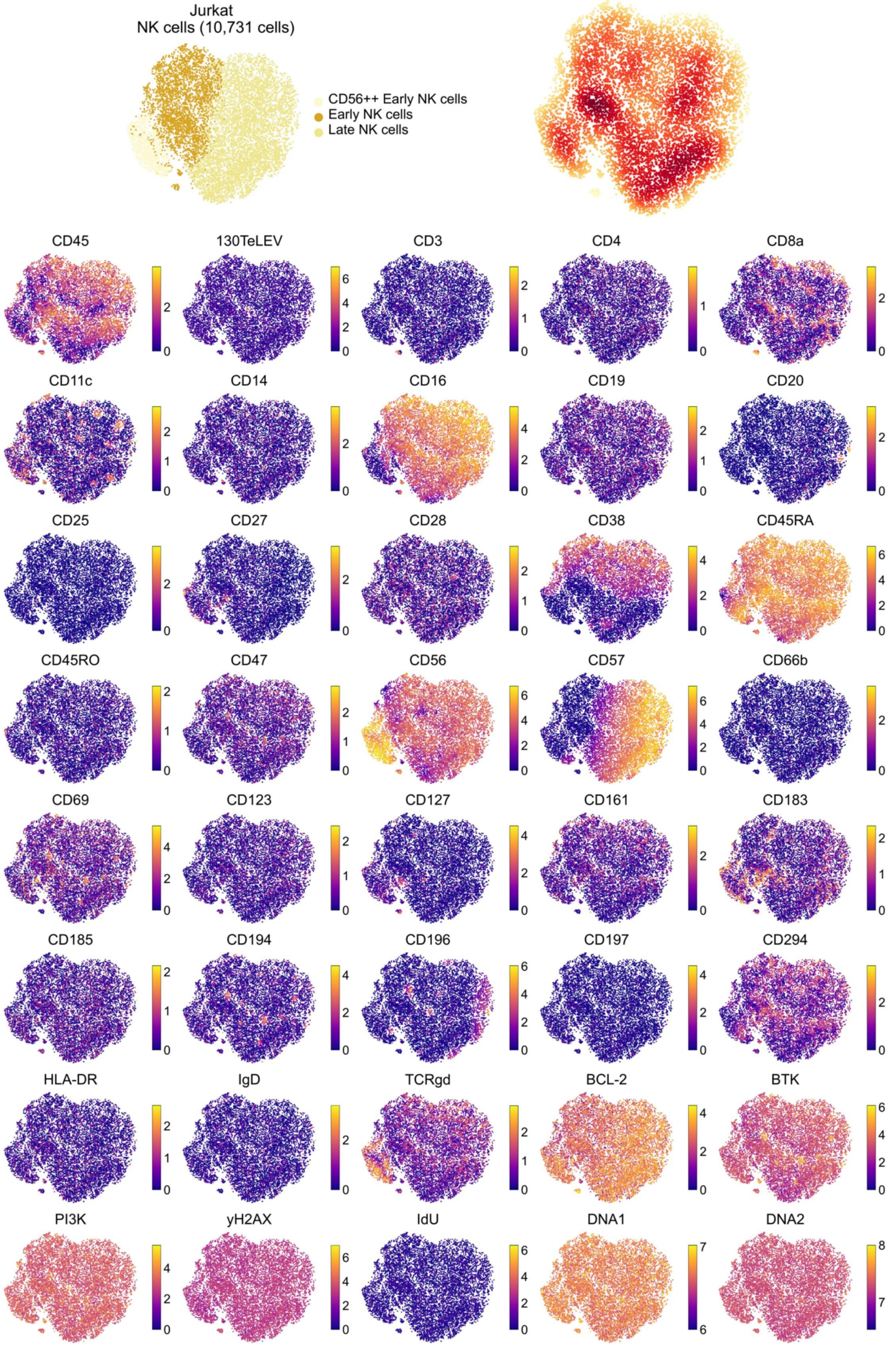

Jurkat
NK cells (10,731 cells)
CD56++ Early NK cells
Early NK cells
Late NK cells
CD45
130TeLEV
CD3
CD4
CD8a
CD11c
CD14
CD16
CD19
CD20
CD25
CD27
CD28
CD38
CD45RA
CD45RO
CD47
CD56
CD57
CD66b
CD69
CD123
CD127
CD161
CD183
CD185
CD194
CD196
CD197
CD294
HLA-DR
IgD
TCRgd
BCL-2
BTK
PI3K
yH2AX
IdU
DNA1
DNA2

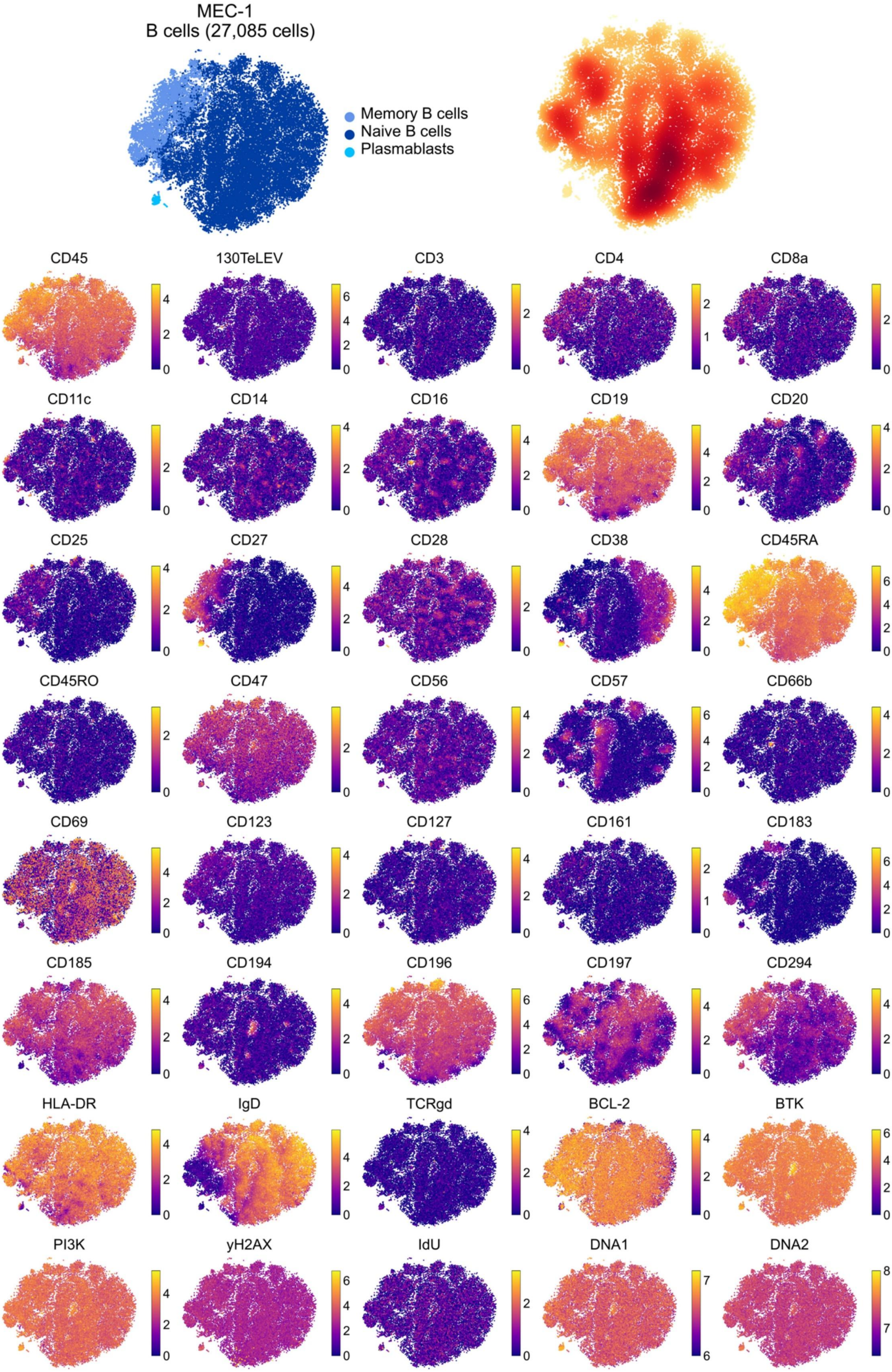
MEC-1
B cells (27,085 cells)
Memory B cells
Naive B cells
Plasmablasts
CD45
130TeLEV
CD3
CD4
CD8a
CD11c
CD14
CD16
CD19
CD20
CD25
CD27
CD28
CD38
CD45RA
CD45RO
CD47
CD56
CD57
CD66b
CD69
CD123
CD127
CD161
CD183
CD185
CD194
CD196
CD197
CD294
HLA-DR
IgD
TCRgd
BCL-2
BTK
PI3K
yH2AX
IdU
DNA1
DNA2

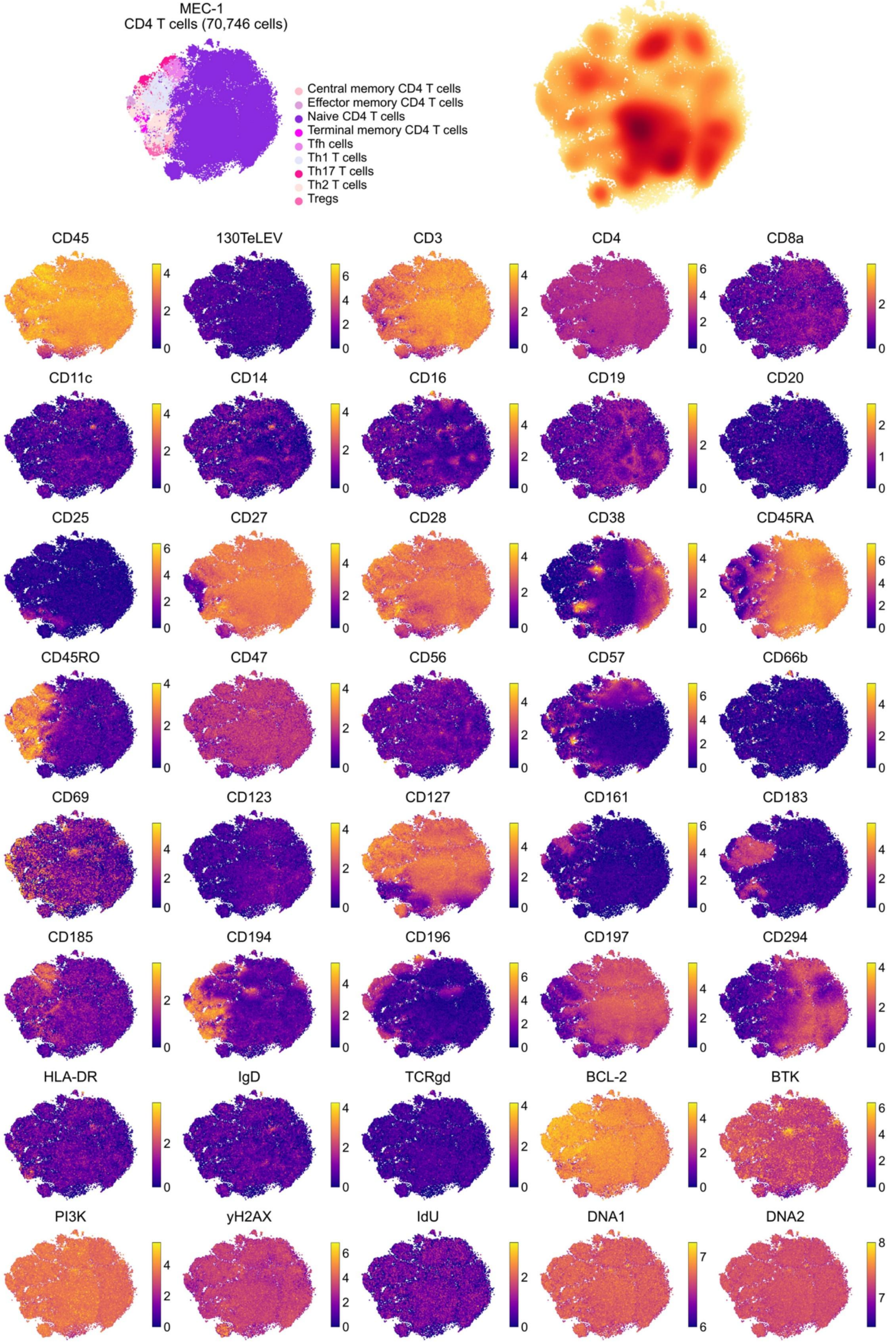
MEC-1
CD4 T cells (70,746 cells)
Central memory CD4 T cells
Effector memory CD4 T cells
Naive CD4 T cells
Terminal memory CD4 T cells
Tfh cells
Th1 T cells
Th17 T cells
Th2 T cells
Tregs
CD45
130TeLEV
CD3
CD4
CD8a
CD11c
CD14
CD16
CD19
CD20
CD25
CD27
CD28
CD38
CD45RA
CD45RO
CD47
CD56
CD57
CD66b
CD69
CD123
CD127
CD161
CD183
CD185
CD194
CD196
CD197
CD294
HLA-DR
IgD
TCRgd
BCL-2
BTK
PI3K
yH2AX
IdU
DNA1
DNA2

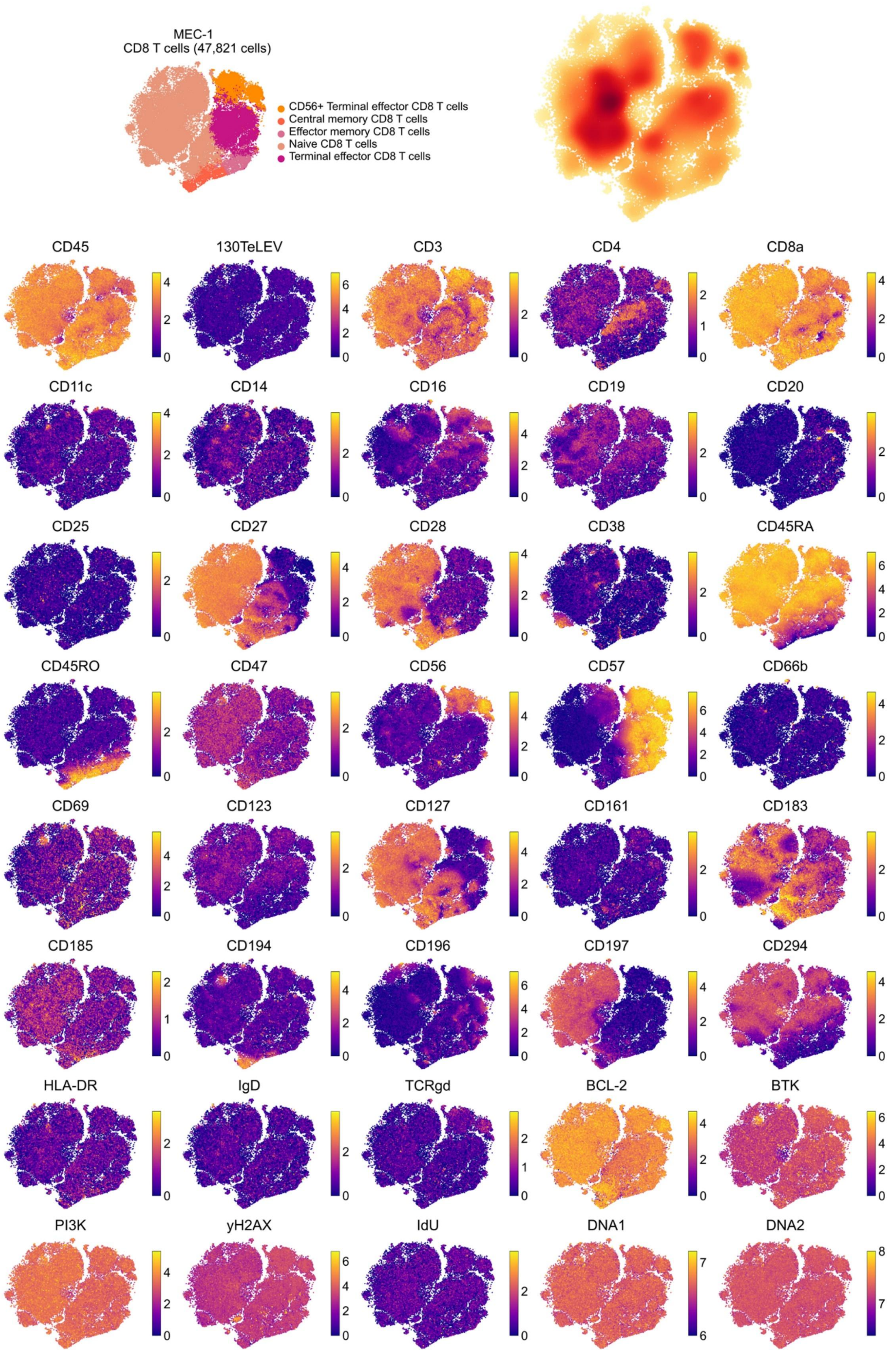

MEC-1
CD8 T cells (47,821 cells)
CD56+ Terminal effector CD8 T cells
Central memory CD8 T cells
Effector memory CD8 T cells
Naive CD8 T cells
Terminal effector CD8 T cells
CD45
130TeLEV
CD3
CD4
CD8a
CD11c
CD14
CD16
CD19
CD20
CD25
CD27
CD28
CD38
CD45RA
CD45RO
CD47
CD56
CD57
CD66b
CD69
CD123
CD127
CD161
CD183
CD185
CD194
CD196
CD197
CD294
HLA-DR
IgD
TCRgd
BCL-2
BTK
PI3K
yH2AX
IdU
DNA1
DNA2

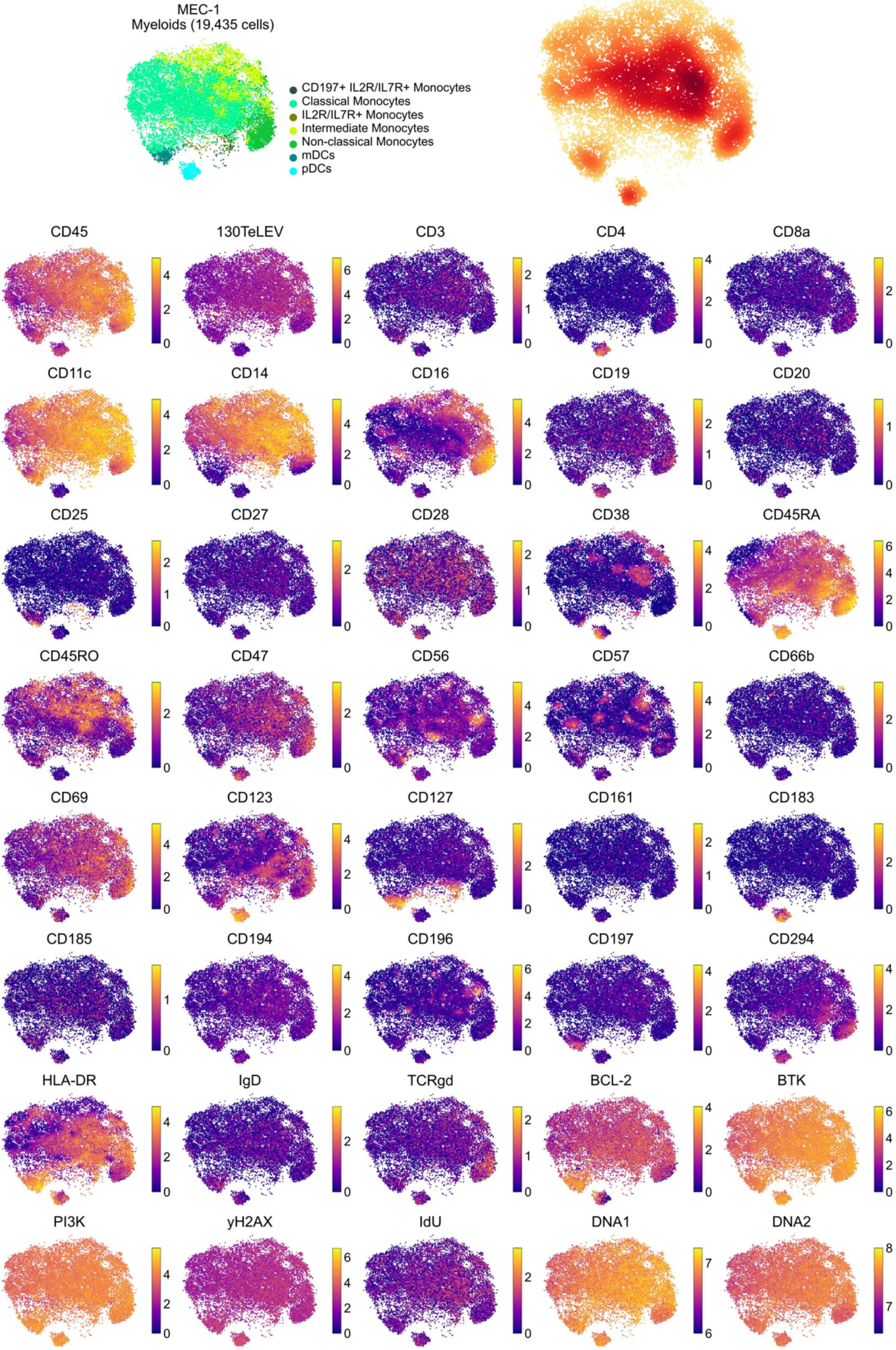

MEC-1
Myeloids (19,435 cells)
CD197+ IL2R/IL7R+ Monocytes
Classical Monocytes
IL2R/IL7R+ Monocytes
Intermediate Monocytes
Non-classical Monocytes
mDCs
pDCs
CD45
130TeLEV
CD3
CD4
CD8a
CD11c
CD14
CD16
CD19
CD20
CD25
CD27
CD28
CD38
CD45RA
CD45RO
CD47
CD56
CD57
CD66b
CD69
CD123
CD127
CD161
CD183
CD185
CD194
CD196
CD197
CD294
HLA-DR
IgD
TCRgd
BCL-2
BTK
PI3K
yH2AX
IdU
DNA1
DNA2

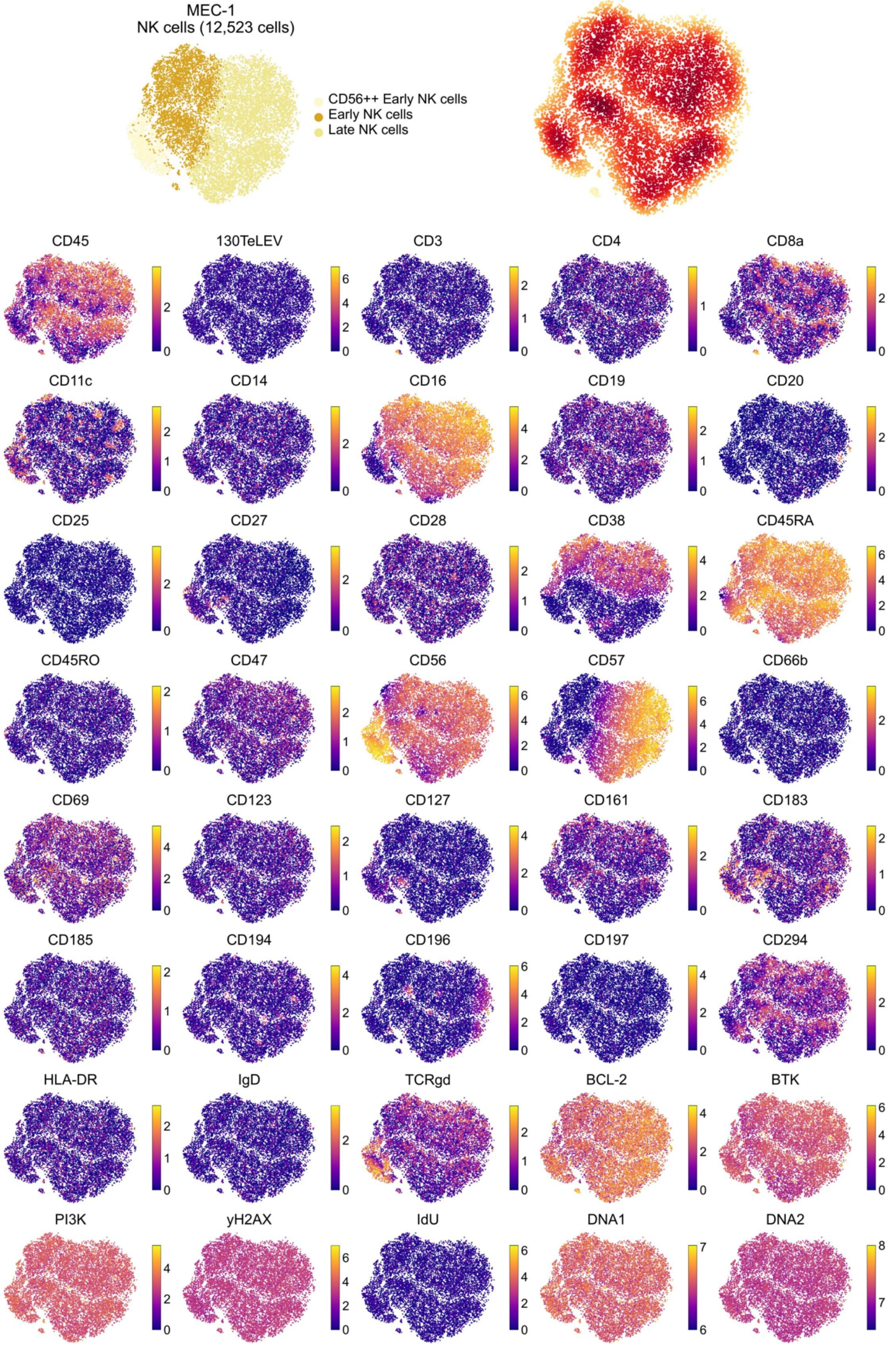

MEC-1
NK cells (12,523 cells)
CD56++ Early NK cells
Early NK cells
Late NK cells
CD45
130TeLEV
CD3
CD4
CD8a
CD11c
CD14
CD16
CD19
CD20
CD25
CD27
CD28
CD38
CD45RA
CD45RO
CD47
CD56
CD57
CD66b
CD69
CD123
CD127
CD161
CD183
CD185
CD194
CD196
CD197
CD294
HLA-DR
IgD
TCRgd
BCL-2
BTK
PI3K
yH2AX
IdU
DNA1
DNA2

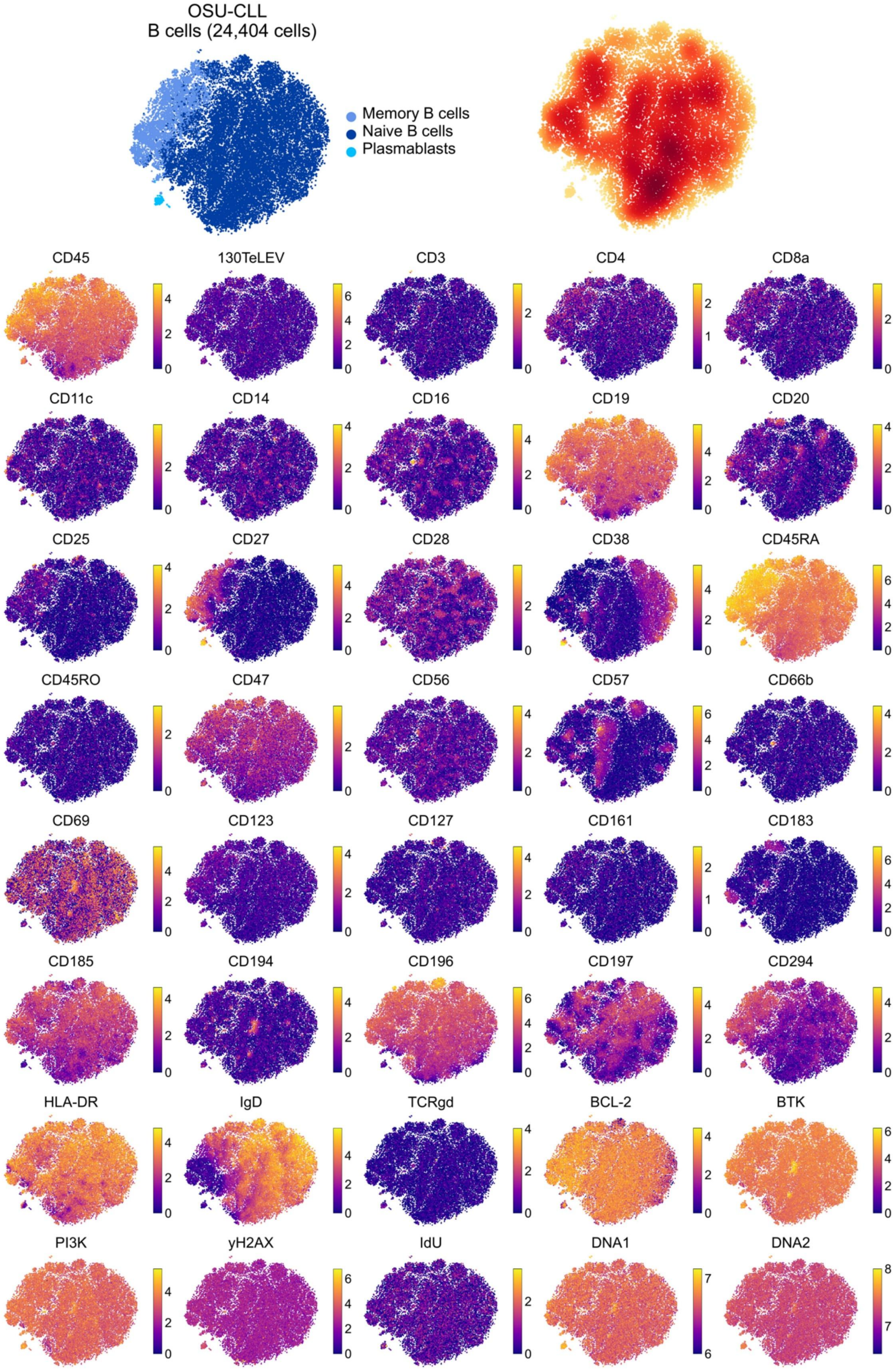

OSU-CLL
B cells (24,404 cells)
Memory B cells
Naive B cells
Plasmablasts
CD45
130TeLEV
CD3
CD4
CD8a
CD11c
CD14
CD16
CD19
CD20
CD25
CD27
CD28
CD38
CD45RA
CD45RO
CD47
CD56
CD57
CD66b
CD69
CD123
CD127
CD161
CD183
CD185
CD194
CD196
CD197
CD294
HLA-DR
IgD
TCRgd
BCL-2
BTK
PI3K
yH2AX
IdU
DNA1
DNA2

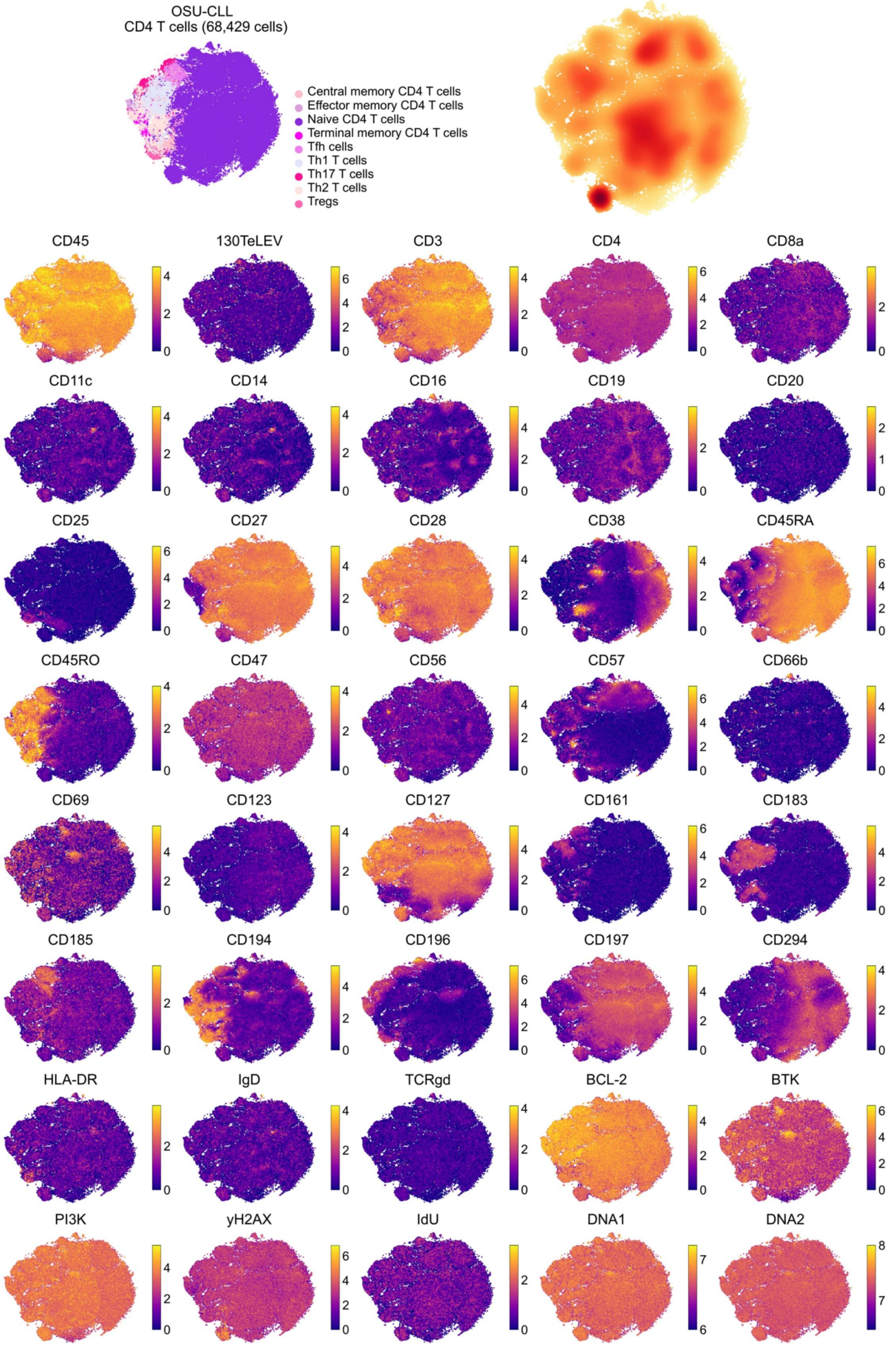
OSU-CLL
CD4 T cells (68,429 cells)
Central memory CD4 T cells
Effector memory CD4 T cells
Naive CD4 T cells
Terminal memory CD4 T cells
Tfh cells
Th1 T cells
Th17 T cells
Th2 T cells
Tregs
CD45
130TeLEV
CD3
CD4
CD8a
CD11c
CD14
CD16
CD19
CD20
CD25
CD27
CD28
CD38
CD45RA
CD45RO
CD47
CD56
CD57
CD66b
CD69
CD123
CD127
CD161
CD183
CD185
CD194
CD196
CD197
CD294
HLA-DR
IgD
TCRgd
BCL-2
BTK
PI3K
yH2AX
IdU
DNA1
DNA2

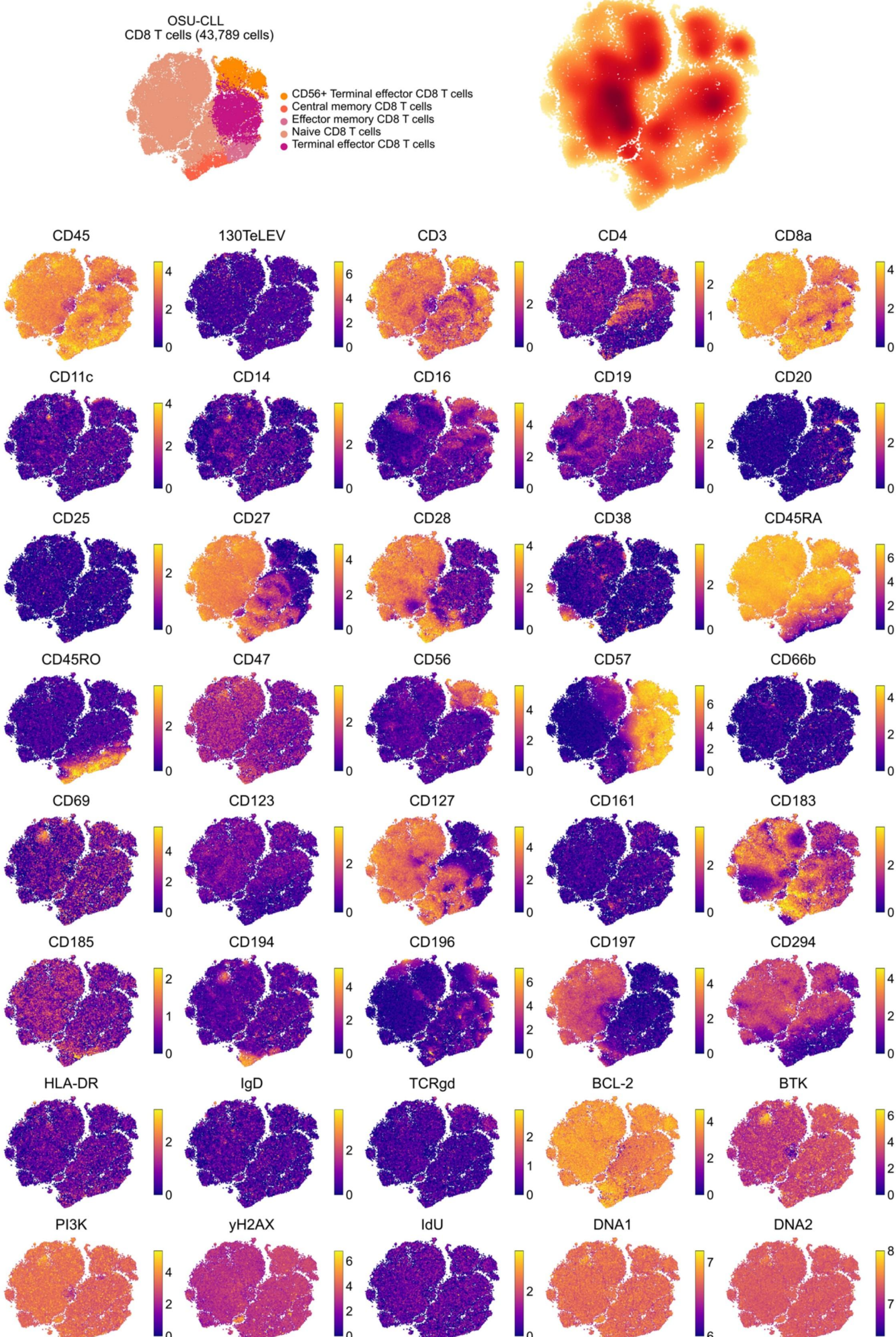
OSU-CLL
CD8 T cells (43,789 cells)
CD56+ Terminal effector CD8 T cells
Central memory CD8 T cells
Effector memory CD8 T cells
Naive CD8 T cells
Terminal effector CD8 T cells
CD45
130TeLEV
CD3
CD4
CD8a
CD11c
CD14
CD16
CD19
CD20
CD25
CD27
CD28
CD38
CD45RA
CD45RO
CD47
CD56
CD57
CD66b
CD69
CD123
CD127
CD161
CD183
CD185
CD194
CD196
CD197
CD294
HLA-DR
IgD
TCRgd
BCL-2
BTK
PI3K
yH2AX
IdU
DNA1
DNA2

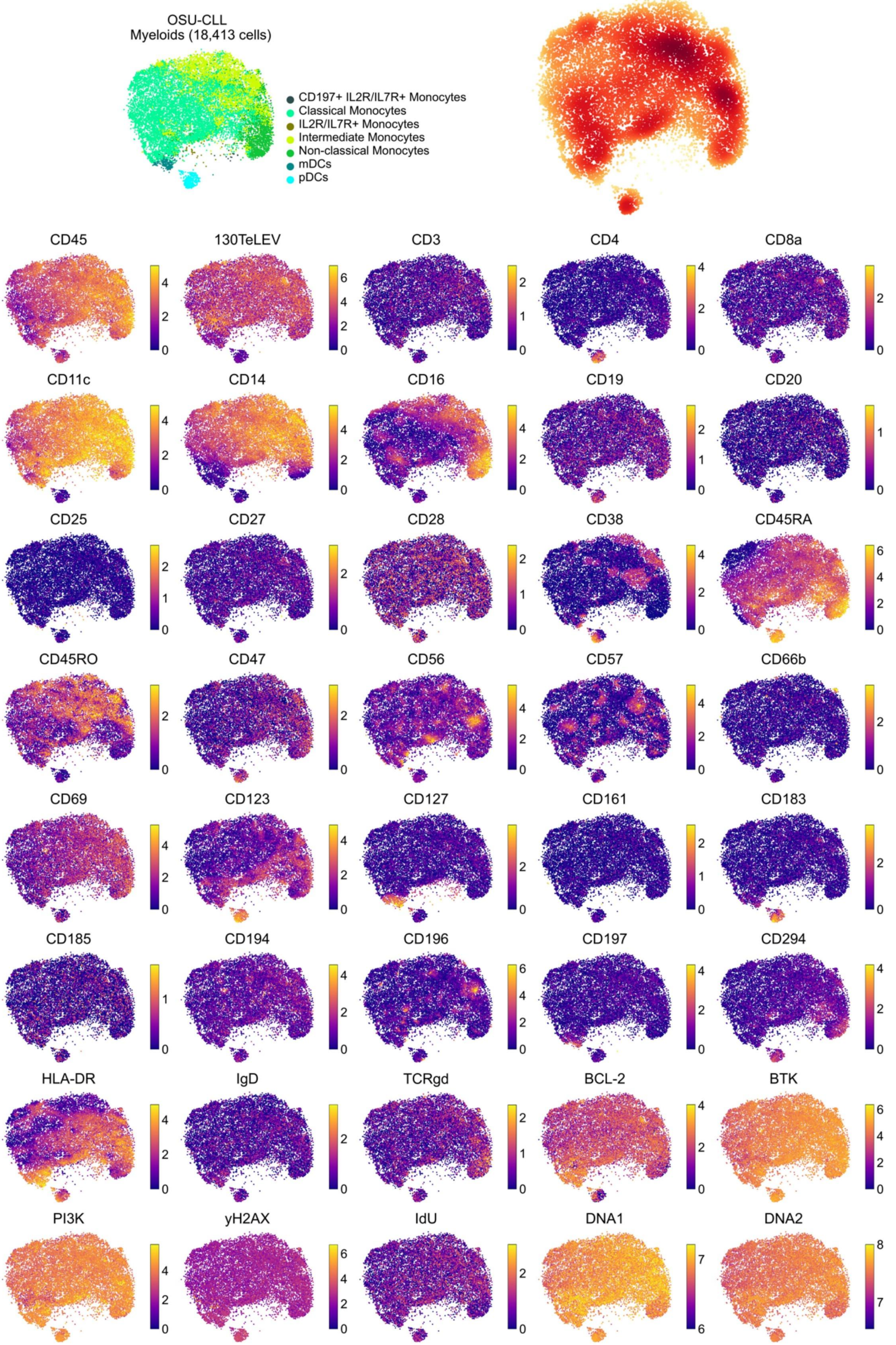

OSU-CLL
Myeloids (18,413 cells)
CD197+ IL2R/IL7R+ Monocytes
Classical Monocytes
IL2R/IL7R+ Monocytes
Intermediate Monocytes
Non-classical Monocytes
mDCs
pDCs
CD45
130TeLEV
CD3
CD4
CD8a
CD11c
CD14
CD16
CD19
CD20
CD25
CD27
CD28
CD38
CD45RA
CD45RO
CD47
CD56
CD57
CD66b
CD69
CD123
CD127
CD161
CD183
CD185
CD194
CD196
CD197
CD294
HLA-DR
IgD
TCRgd
BCL-2
BTK
PI3K
yH2AX
IdU
DNA1
DNA2

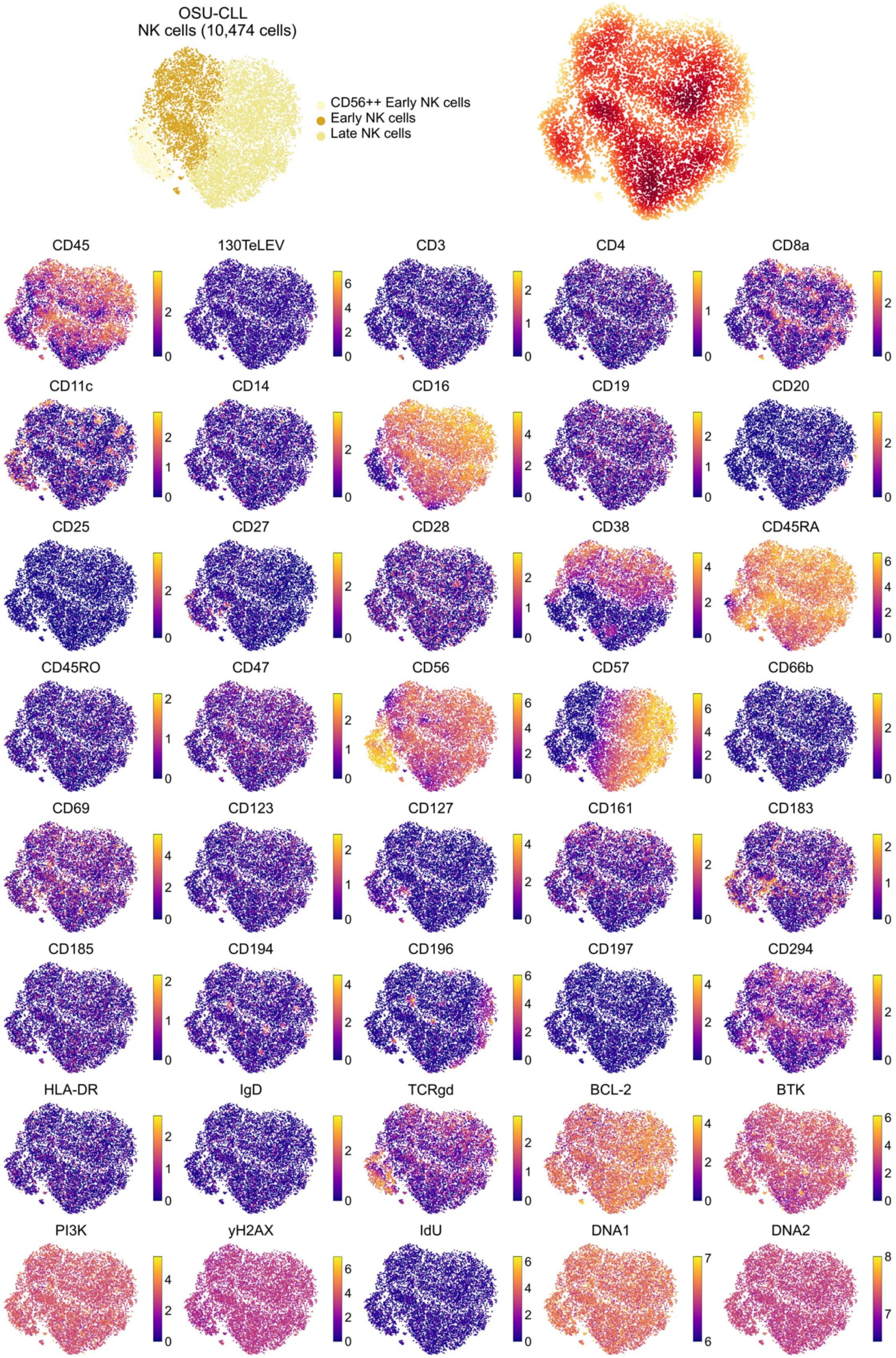

OSU-CLL
NK cells (10,474 cells)
CD56++ Early NK cells
Early NK cells
Late NK cells
CD45
130TeLEV
CD3
CD4
CD8a
CD11c
CD14
CD16
CD19
CD20
CD25
CD27
CD28
CD38
CD45RA
CD45RO
CD47
CD56
CD57
CD66b
CD69
CD123
CD127
CD161
CD183
CD185
CD194
CD196
CD197
CD294
HLA-DR
IgD
TCRgd
BCL-2
BTK
PI3K
yH2AX
IdU
DNA1
DNA2

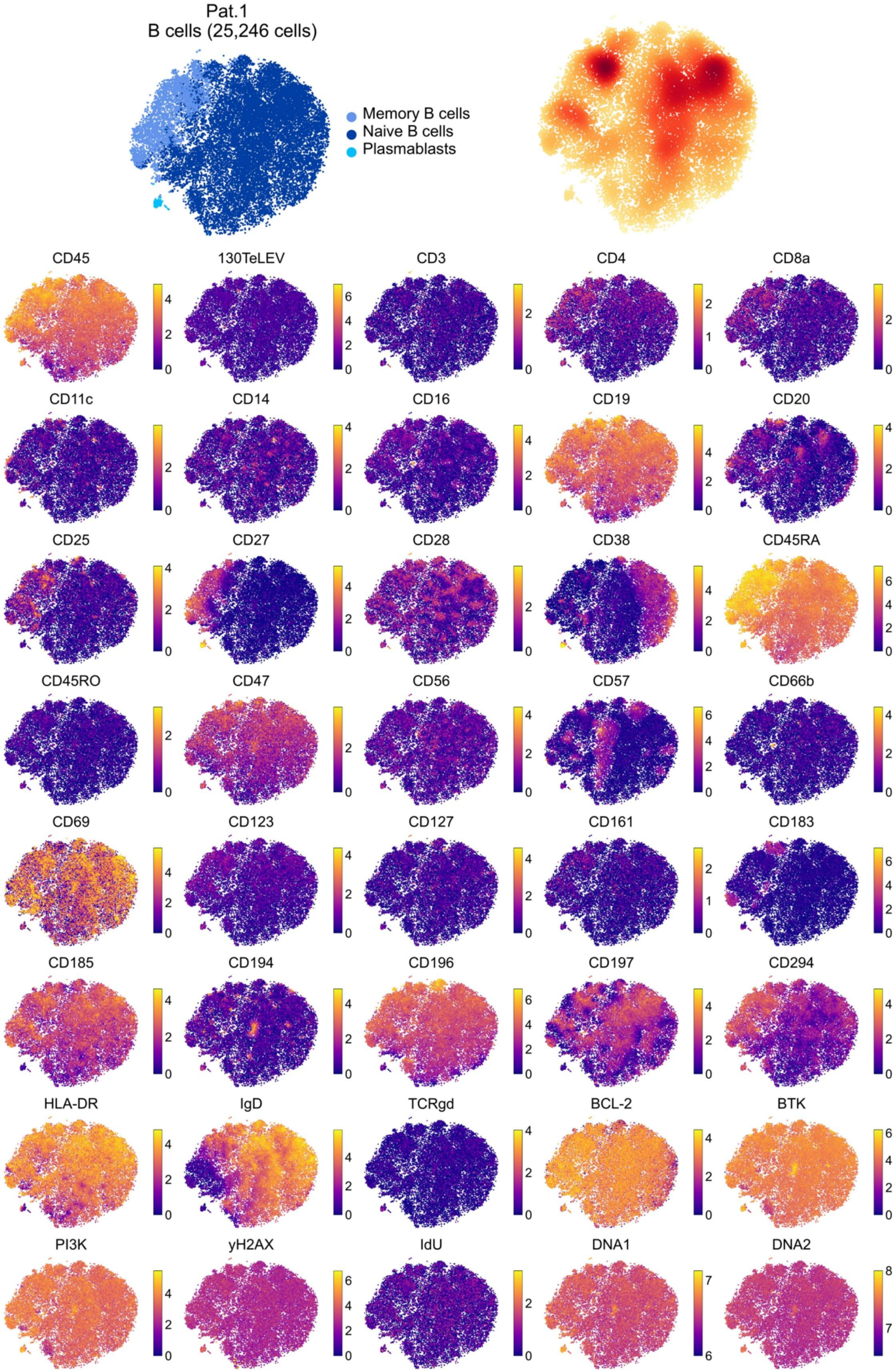

Pat.1
B cells (25,246 cells)
Memory B cells
Naive B cells
Plasmablasts
CD45
130TeLEV
CD3
CD4
CD8a
CD11c
CD14
CD16
CD19
CD20
CD25
CD27
CD28
CD38
CD45RA
CD45RO
CD47
CD56
CD57
CD66b
CD69
CD123
CD127
CD161
CD183
CD185
CD194
CD196
CD197
CD294
HLA-DR
IgD
TCRgd
BCL-2
BTK
PI3K
yH2AX
IdU
DNA1
DNA2

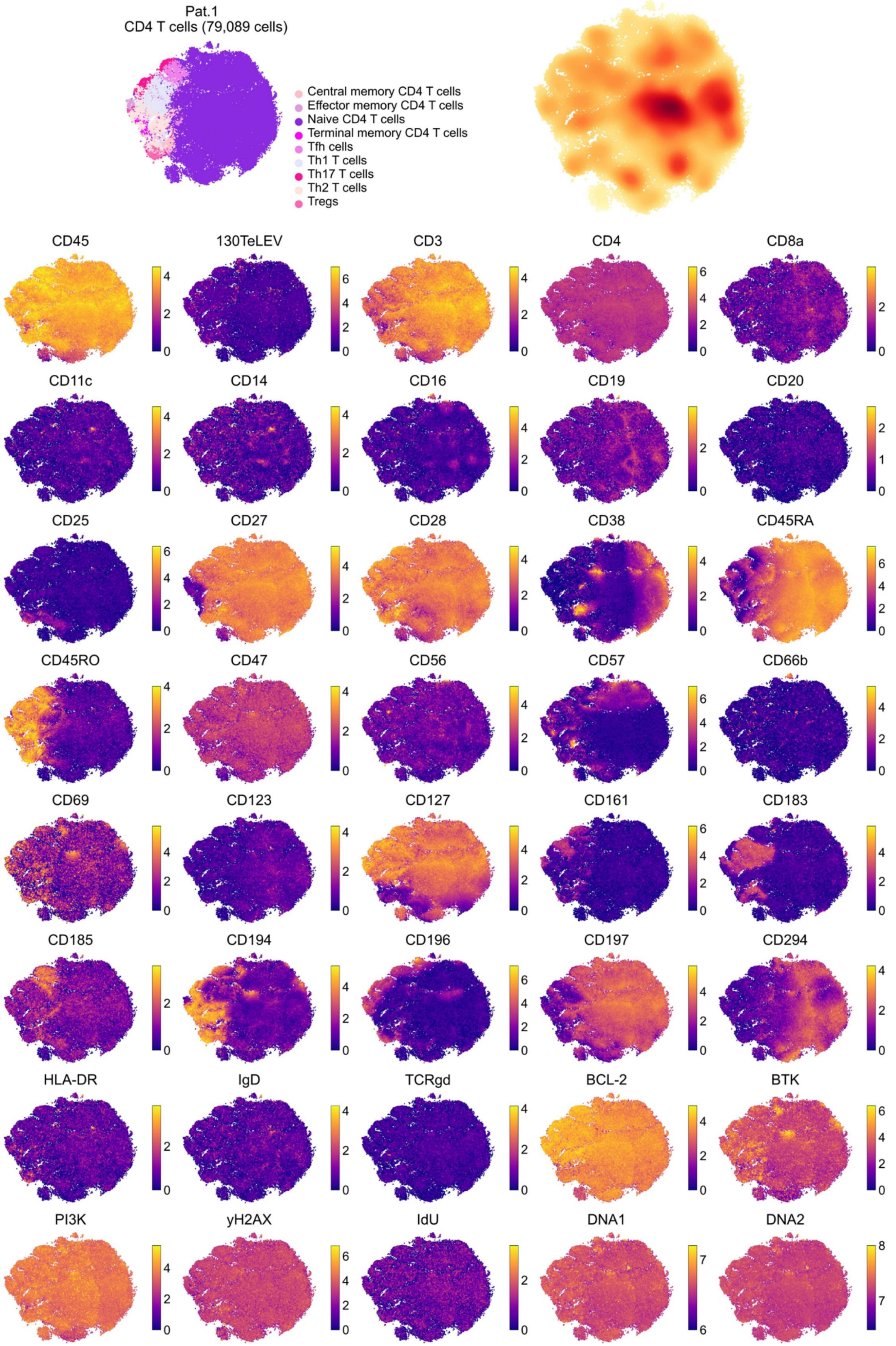
Pat.1
CD4 T cells (79,089 cells)
Central memory CD4 T cells
Effector memory CD4 T cells
Naive CD4 T cells
Terminal memory CD4 T cells
Tfh cells
Th1 T cells
Th17 T cells
Th2 T cells
Tregs
CD45
130TeLEV
CD3
CD4
CD8a
CD11c
CD14
CD16
CD19
CD20
CD25
CD27
CD28
CD38
CD45RA
CD45RO
CD47
CD56
CD57
CD66b
CD69
CD123
CD127
CD161
CD183
CD185
CD194
CD196
CD197
CD294
HLA-DR
IgD
TCRgd
BCL-2
BTK
PI3K
yH2AX
IdU
DNA1
DNA2

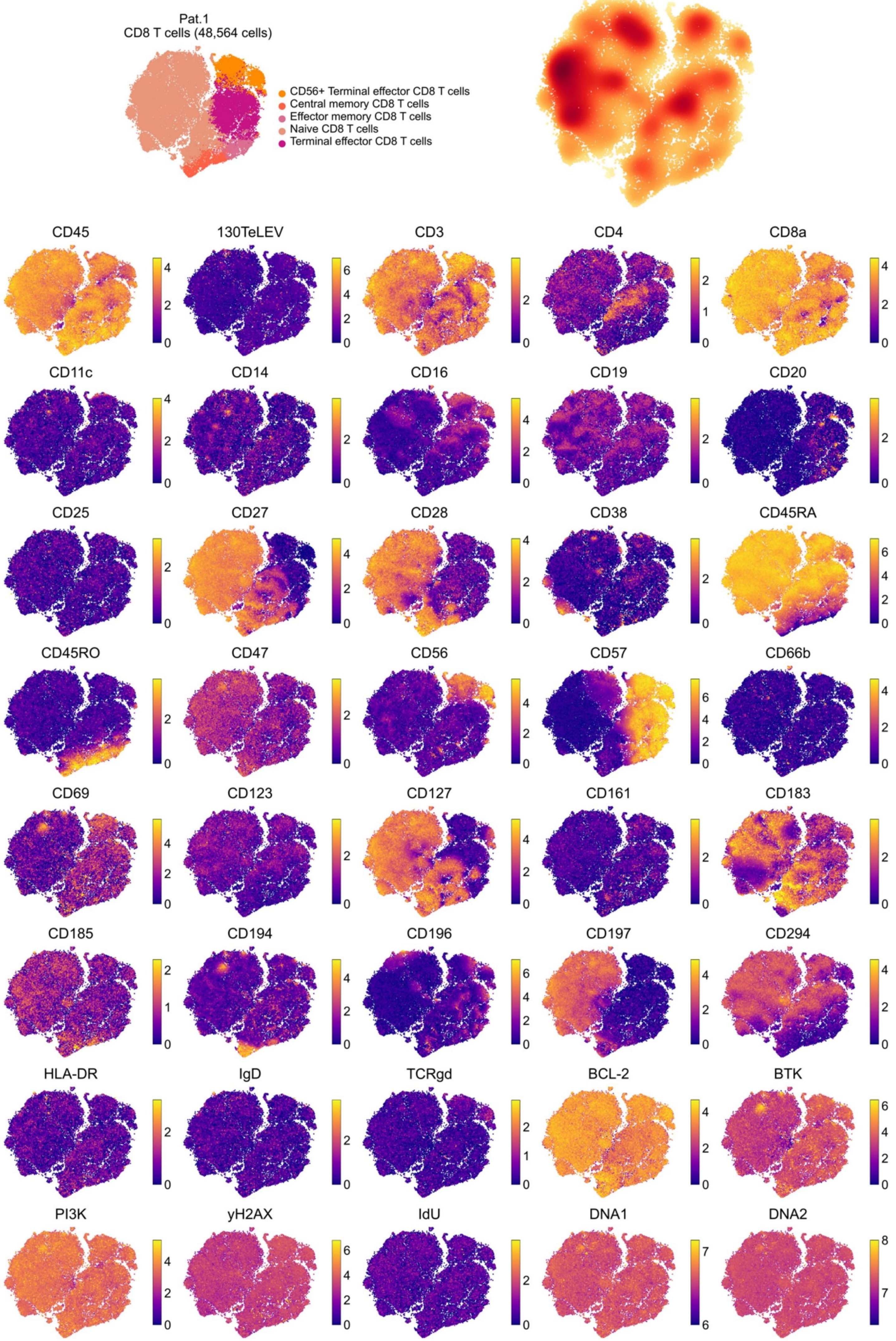
Pat.1
CD8 T cells (48,564 cells)
CD56+ Terminal effector CD8 T cells
Central memory CD8 T cells
Effector memory CD8 T cells
Naive CD8 T cells
Terminal effector CD8 T cells
CD45
130TeLEV
CD3
CD4
CD8a
CD11c
CD14
CD16
CD19
CD20
CD25
CD27
CD28
CD38
CD45RA
CD45RO
CD47
CD56
CD57
CD66b
CD69
CD123
CD127
CD161
CD183
CD185
CD194
CD196
CD197
CD294
HLA-DR
IgD
TCRgd
BCL-2
BTK
PI3K
yH2AX
IdU
DNA1
DNA2

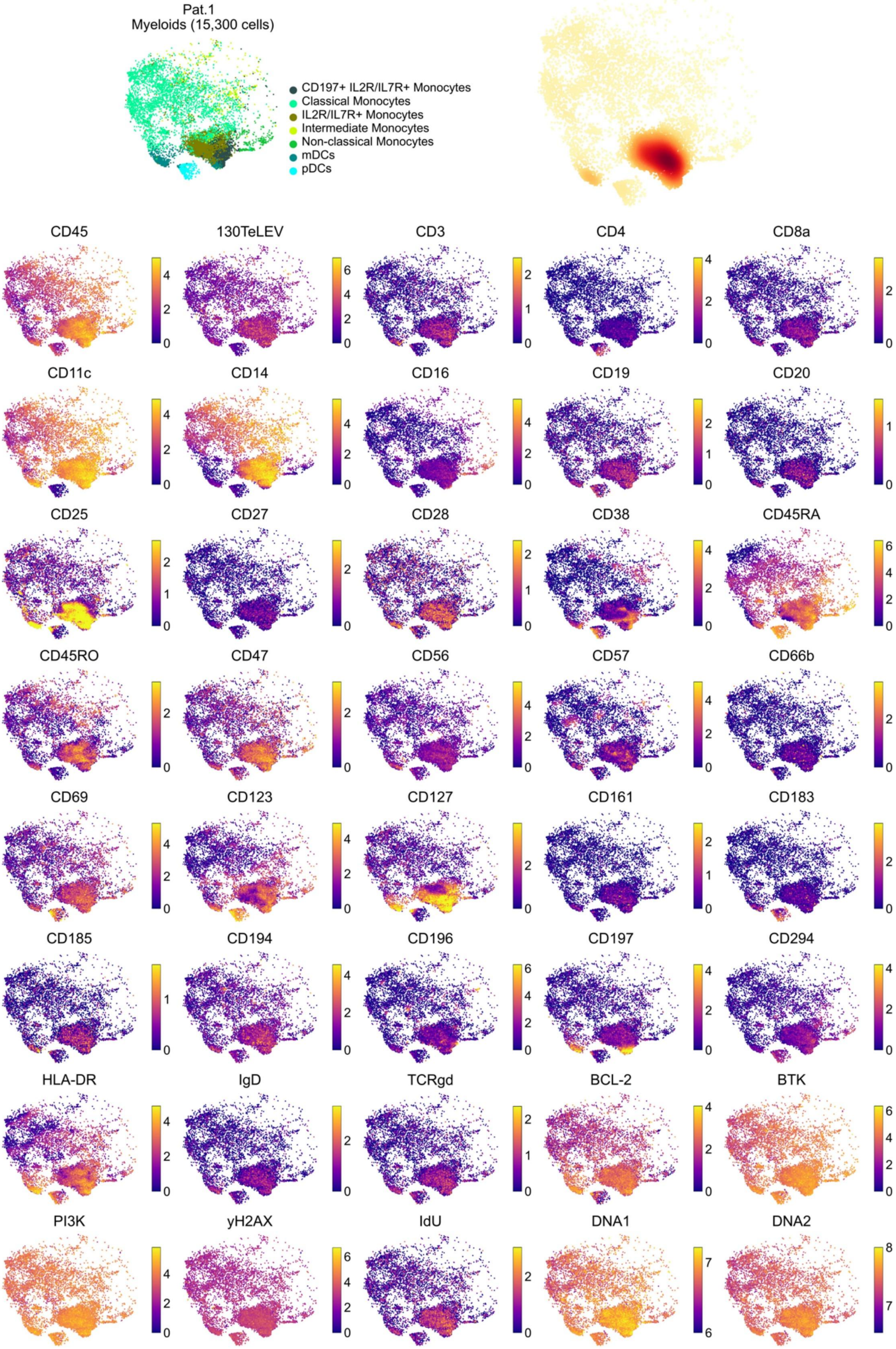
Pat.1
Myeloids (15,300 cells)
CD197+ IL2R/IL7R+ Monocytes
Classical Monocytes
IL2R/IL7R+ Monocytes
Intermediate Monocytes
Non-classical Monocytes
mDCs
pDCs
CD45
130TeLEV
CD3
CD4
CD8a
CD11c
CD14
CD16
CD19
CD20
CD25
CD27
CD28
CD38
CD45RA
CD45RO
CD47
CD56
CD57
CD66b
CD69
CD123
CD127
CD161
CD183
CD185
CD194
CD196
CD197
CD294
HLA-DR
IgD
TCRgd
BCL-2
BTK
PI3K
yH2AX
IdU
DNA1
DNA2

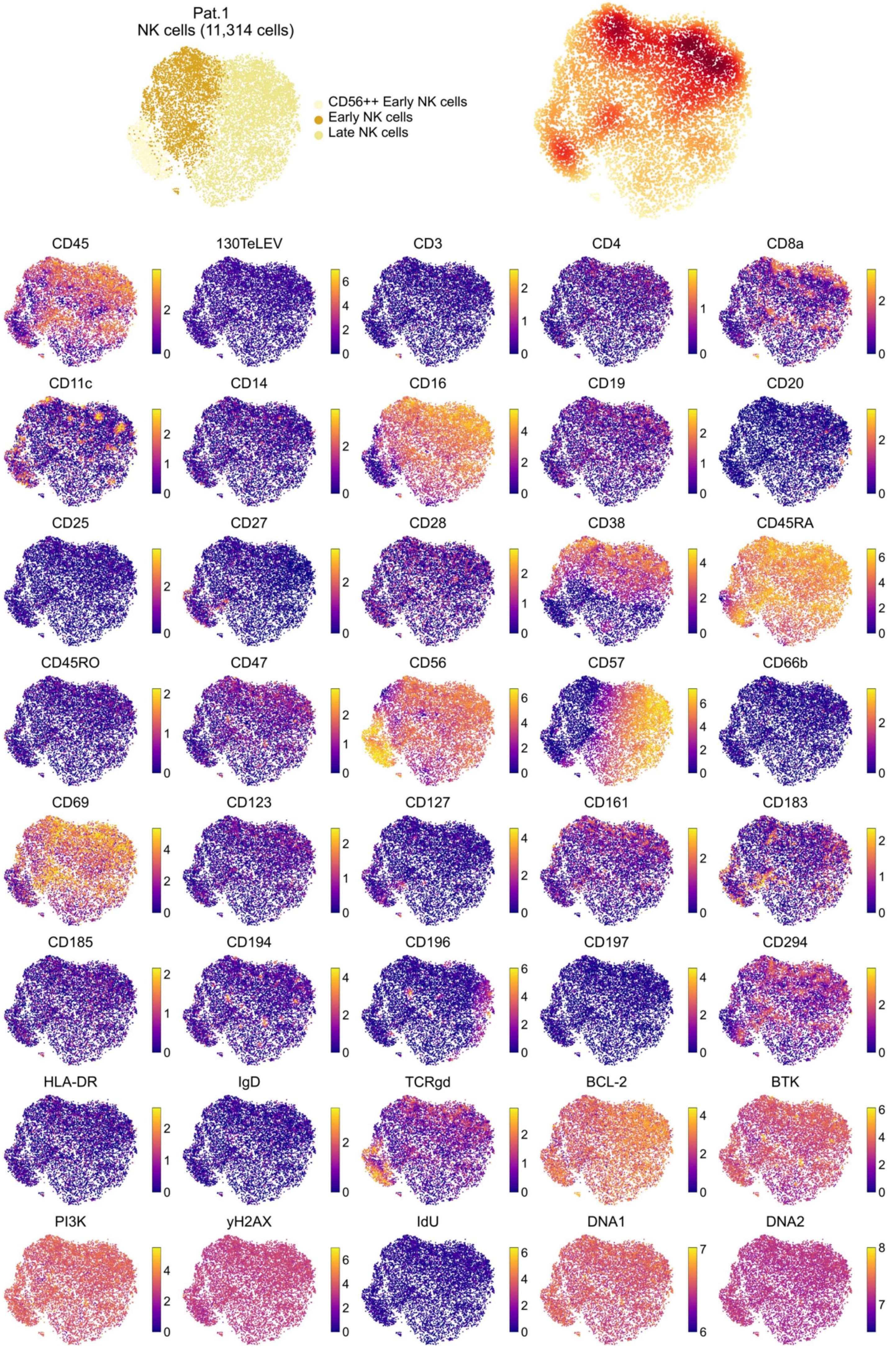
Pat.1
NK cells (11,314 cells)
CD56++ Early NK cells
Early NK cells
Late NK cells
CD45
130TeLEV
CD3
CD4
CD8a
CD11c
CD14
CD16
CD19
CD20
CD25
CD27
CD28
CD38
CD45RA
CD45RO
CD47
CD56
CD57
CD66b
CD69
CD123
CD127
CD161
CD183
CD185
CD194
CD196
CD197
CD294
HLA-DR
IgD
TCRgd
BCL-2
BTK
PI3K
yH2AX
IdU
DNA1
DNA2

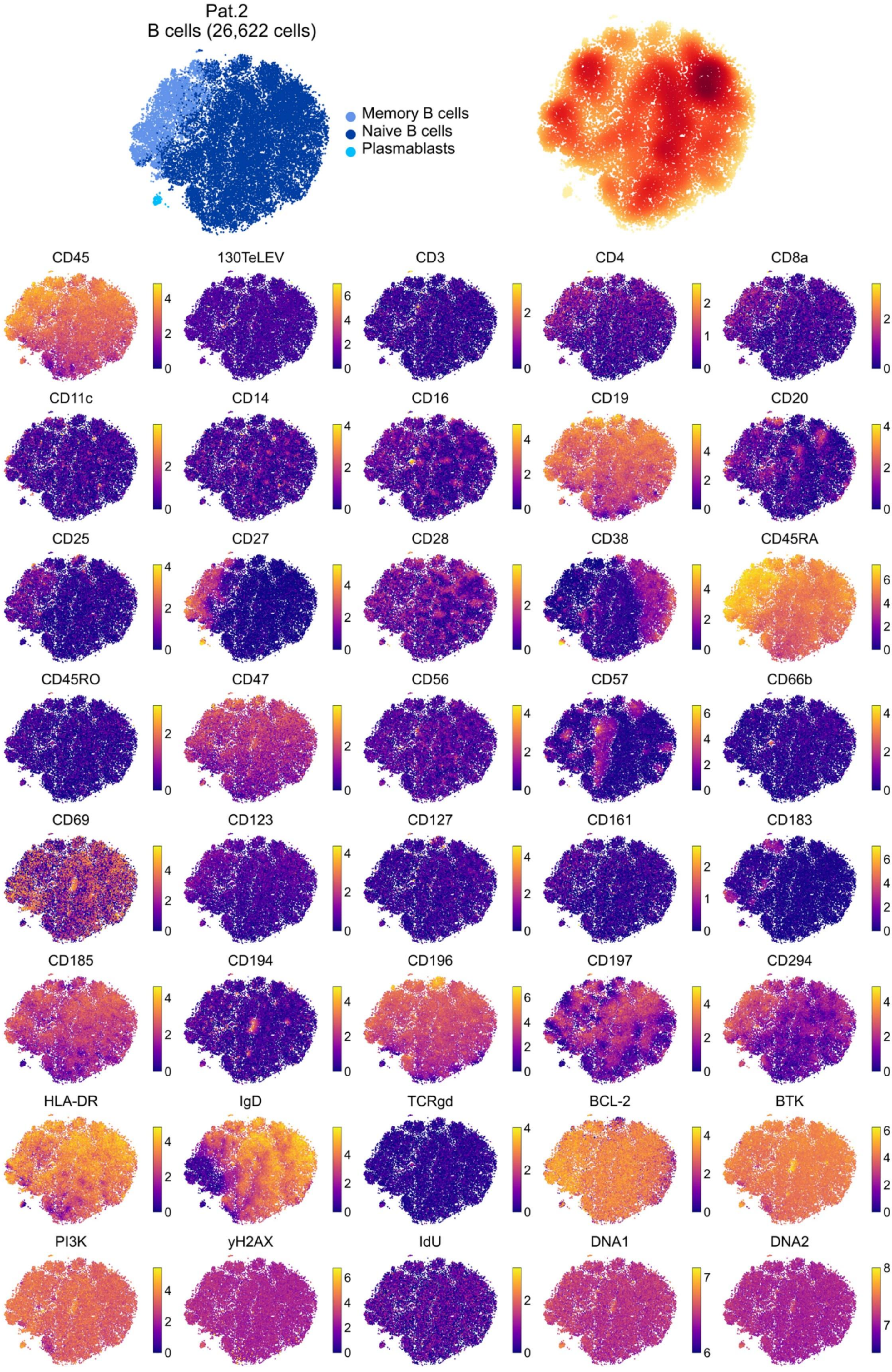

Pat.2
B cells (26,622 cells)
Memory B cells
Naive B cells
Plasmablasts
CD45
130TeLEV
CD3
CD4
CD8a
CD11c
CD14
CD16
CD19
CD20
CD25
CD27
CD28
CD38
CD45RA
CD45RO
CD47
CD56
CD57
CD66b
CD69
CD123
CD127
CD161
CD183
CD185
CD194
CD196
CD197
CD294
HLA-DR
IgD
TCRgd
BCL-2
BTK
PI3K
yH2AX
IdU
DNA1
DNA2

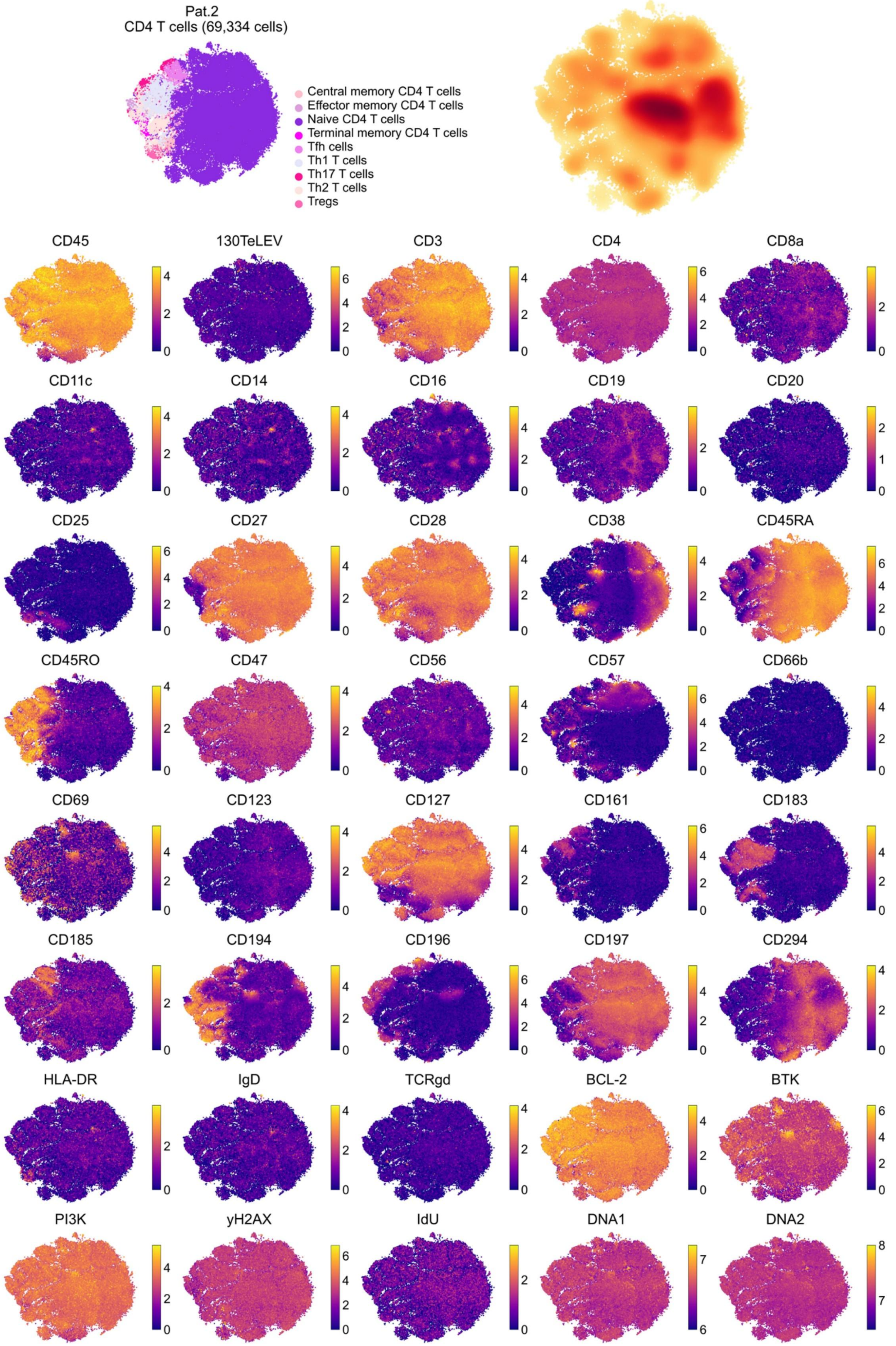

Pat.2
CD4 T cells (69,334 cells)
Central memory CD4 T cells
Effector memory CD4 T cells
Naive CD4 T cells
Terminal memory CD4 T cells
Tfh cells
Th1 T cells
Th17 T cells
Th2 T cells
Tregs
CD45
130TeLEV
CD3
CD4
CD8a
CD11c
CD14
CD16
CD19
CD20
CD25
CD27
CD28
CD38
CD45RA
CD45RO
CD47
CD56
CD57
CD66b
CD69
CD123
CD127
CD161
CD183
CD185
CD194
CD196
CD197
CD294
HLA-DR
IgD
TCRgd
BCL-2
BTK
PI3K
yH2AX
IdU
DNA1
DNA2

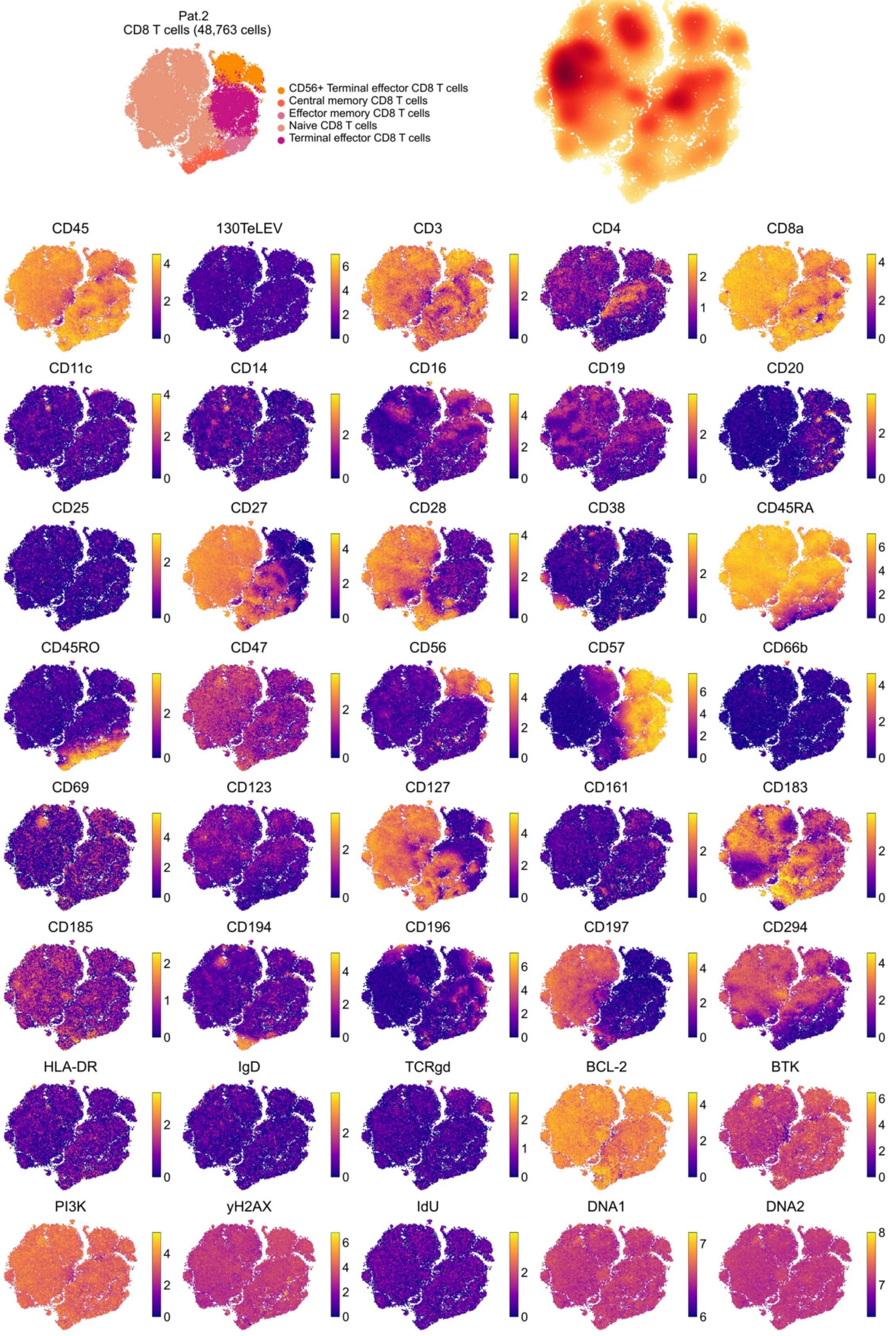
Pat.2
CD8 T cells (48,763 cells)
CD56+ Terminal effector CD8 T cells
Central memory CD8 T cells
Effector memory CD8 T cells
Naive CD8 T cells
Terminal effector CD8 T cells
CD45
130TeLEV
CD3
CD4
CD8a
CD11c
CD14
CD16
CD19
CD20
CD25
CD27
CD28
CD38
CD45RA
CD45RO
CD47
CD56
CD57
CD66b
CD69
CD123
CD127
CD161
CD183
CD185
CD194
CD196
CD197
CD294
HLA-DR
IgD
TCRgd
BCL-2
BTK
PI3K
yH2AX
IdU
DNA1
DNA2

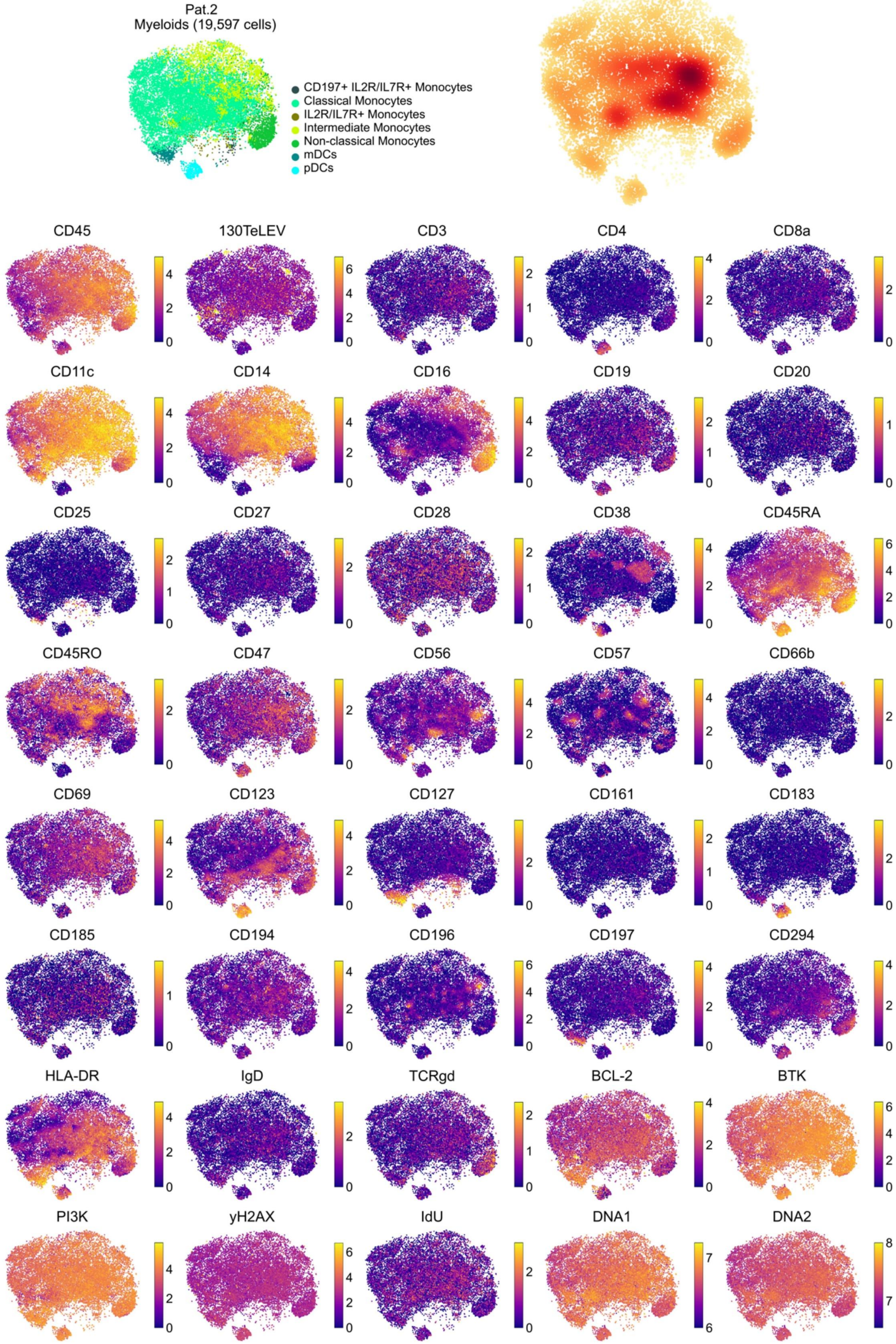
Pat.2
Myeloids (19,597 cells)
CD197+ IL2R/IL7R+ Monocytes
Classical Monocytes
IL2R/IL7R+ Monocytes
Intermediate Monocytes
Non-classical Monocytes
mDCs
pDCs
CD45
130TeLEV
CD3
CD4
CD8a
CD11c
CD14
CD16
CD19
CD20
CD25
CD27
CD28
CD38
CD45RA
CD45RO
CD47
CD56
CD57
CD66b
CD69
CD123
CD127
CD161
CD183
CD185
CD194
CD196
CD197
CD294
HLA-DR
IgD
TCRgd
BCL-2
BTK
PI3K
yH2AX
IdU
DNA1
DNA2

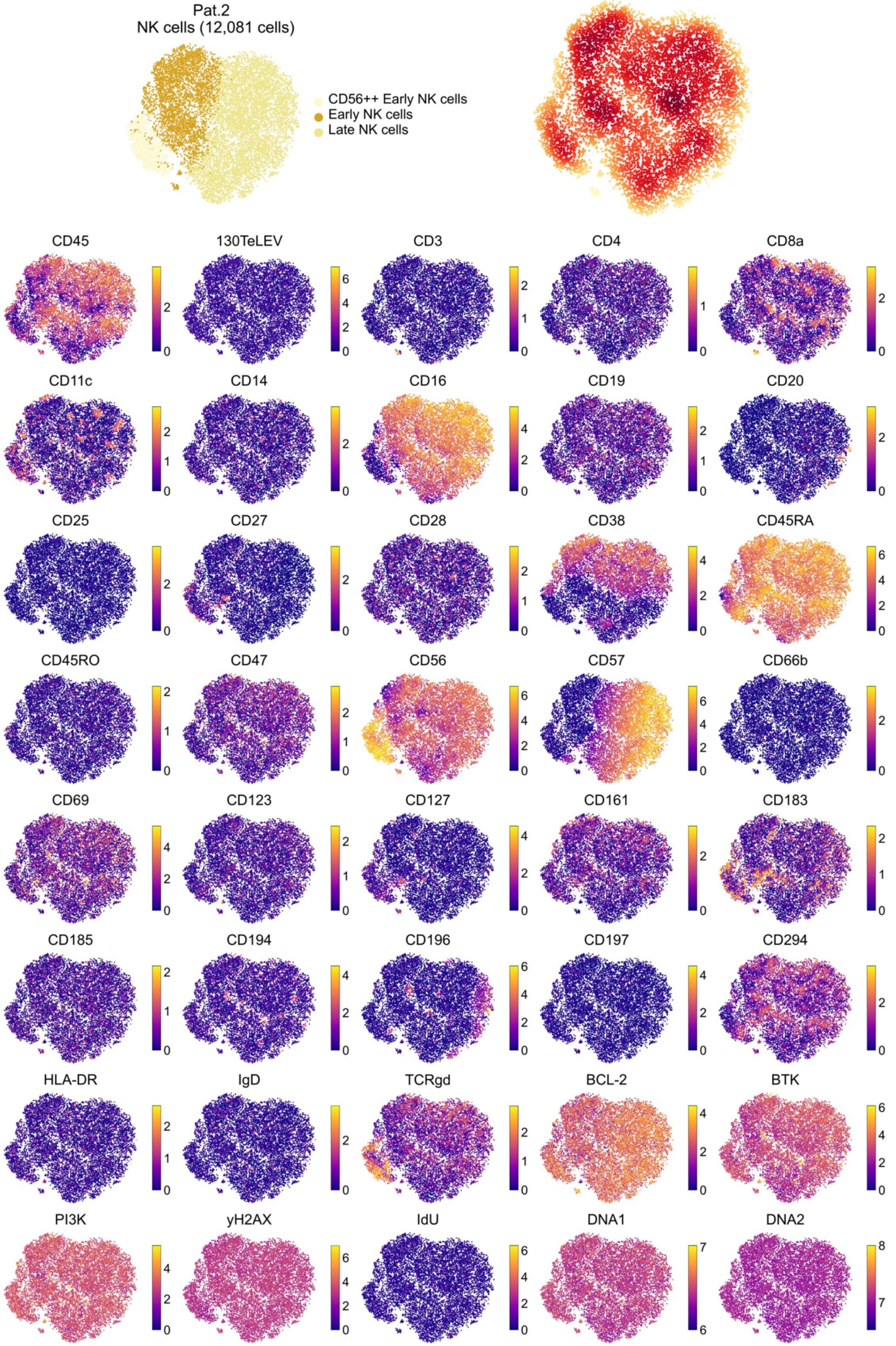
Pat.2
NK cells (12,081 cells)
CD56++ Early NK cells
Early NK cells
Late NK cells
CD45
130TeLEV
CD3
CD4
CD8a
CD11c
CD14
CD16
CD19
CD20
CD25
CD27
CD28
CD38
CD45RA
CD45RO
CD47
CD56
CD57
CD66b
CD69
CD123
CD127
CD161
CD183
CD185
CD194
CD196
CD197
CD294
HLA-DR
IgD
TCRgd
BCL-2
BTK
PI3K
yH2AX
IdU
DNA1
DNA2

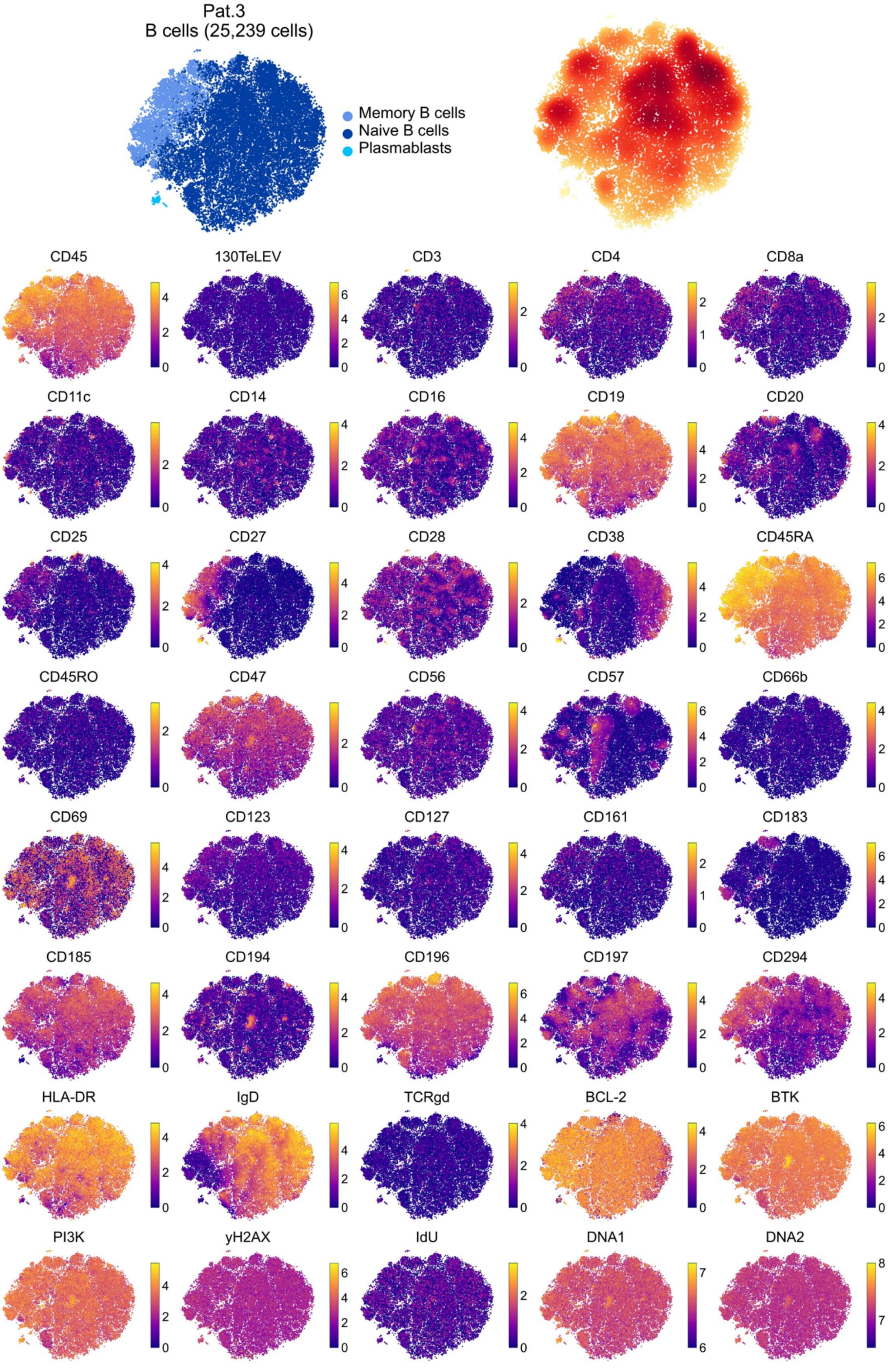
Pat.3
B cells (25,239 cells)
Memory B cells
Naive B cells
Plasmablasts
CD45
130TeLEV
CD3
CD4
CD8a
CD11c
CD14
CD16
CD19
CD20
CD25
CD27
CD28
CD38
CD45RA
CD45RO
CD47
CD56
CD57
CD66b
CD69
CD123
CD127
CD161
CD183
CD185
CD194
CD196
CD197
CD294
HLA-DR
IgD
TCRgd
BCL-2
BTK
PI3K
yH2AX
IdU
DNA1
DNA2

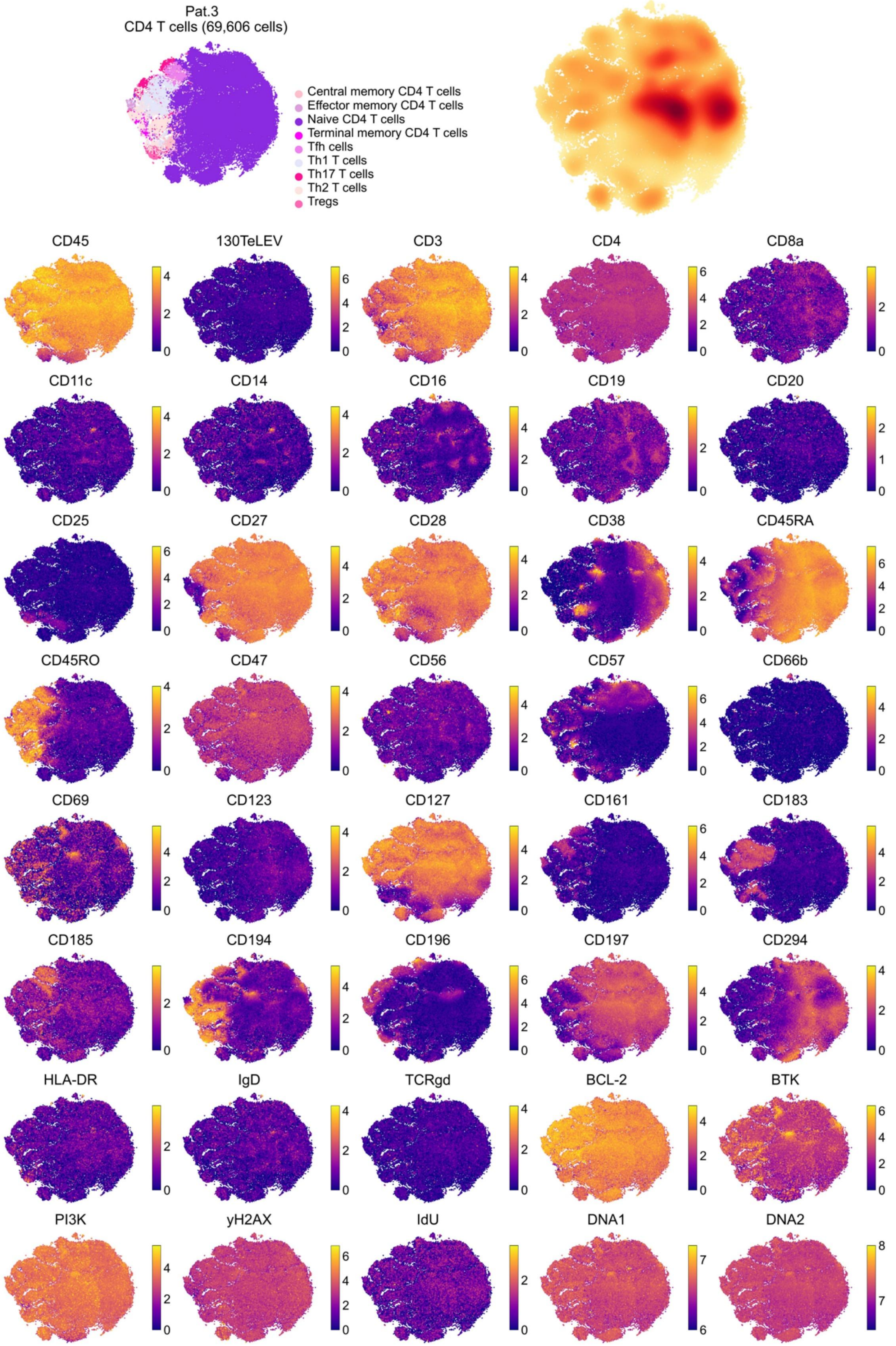
Pat.3
CD4 T cells (69,606 cells)
Central memory CD4 T cells
Effector memory CD4 T cells
Naive CD4 T cells
Terminal memory CD4 T cells
Tfh cells
Th1 T cells
Th17 T cells
Th2 T cells
Tregs
CD45
130TeLEV
CD3
CD4
CD8a
CD11c
CD14
CD16
CD19
CD20
CD25
CD27
CD28
CD38
CD45RA
CD45RO
CD47
CD56
CD57
CD66b
CD69
CD123
CD127
CD161
CD183
CD185
CD194
CD196
CD197
CD294
HLA-DR
IgD
TCRgd
BCL-2
BTK
PI3K
yH2AX
IdU
DNA1
DNA2

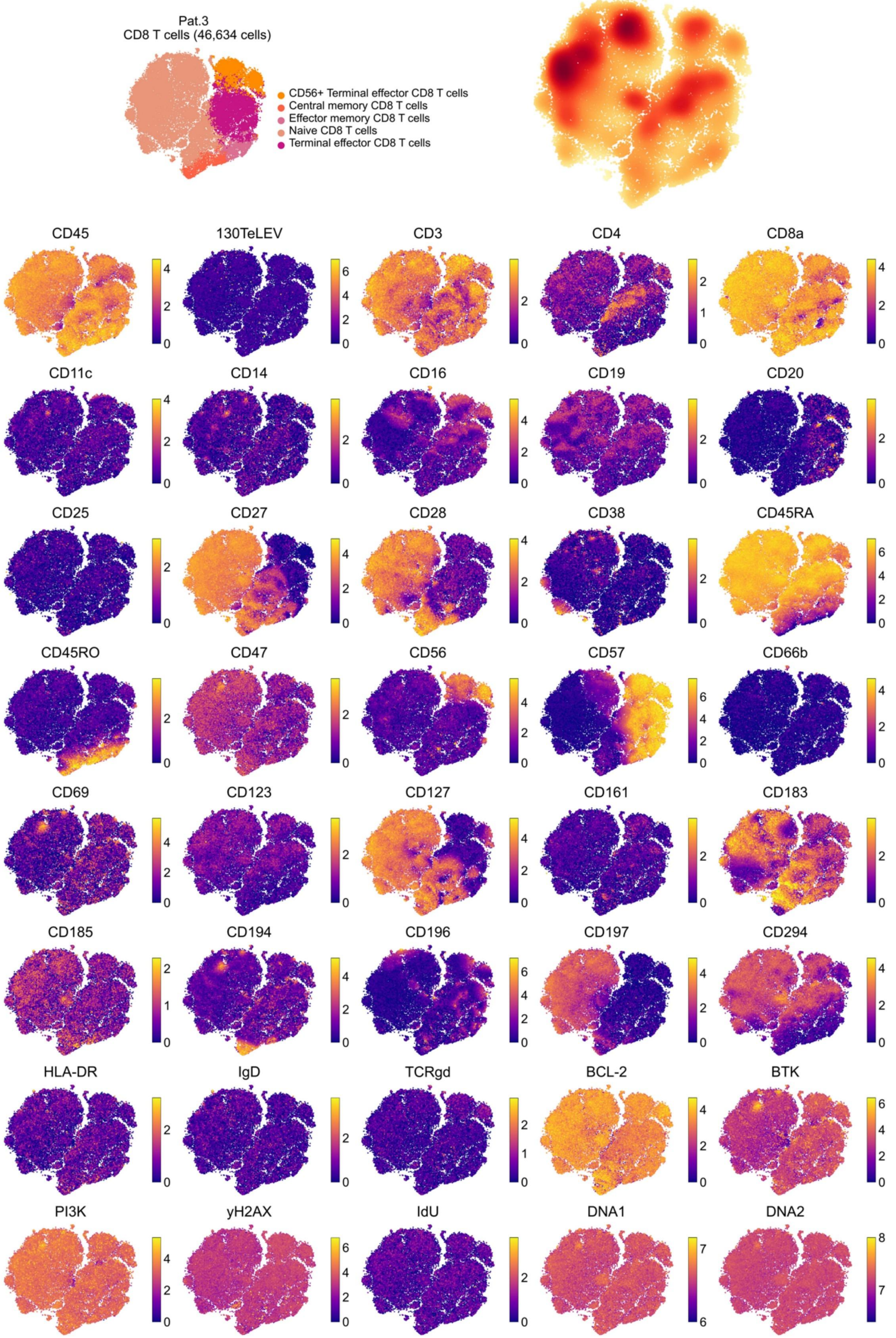
Pat.3
CD8 T cells (46,634 cells)
CD56+ Terminal effector CD8 T cells
Central memory CD8 T cells
Effector memory CD8 T cells
Naive CD8 T cells
Terminal effector CD8 T cells
CD45
130TeLEV
CD3
CD4
CD8a
CD11c
CD14
CD16
CD19
CD20
CD25
CD27
CD28
CD38
CD45RA
CD45RO
CD47
CD56
CD57
CD66b
CD69
CD123
CD127
CD161
CD183
CD185
CD194
CD196
CD197
CD294
HLA-DR
IgD
TCRgd
BCL-2
BTK
PI3K
yH2AX
IdU
DNA1
DNA2

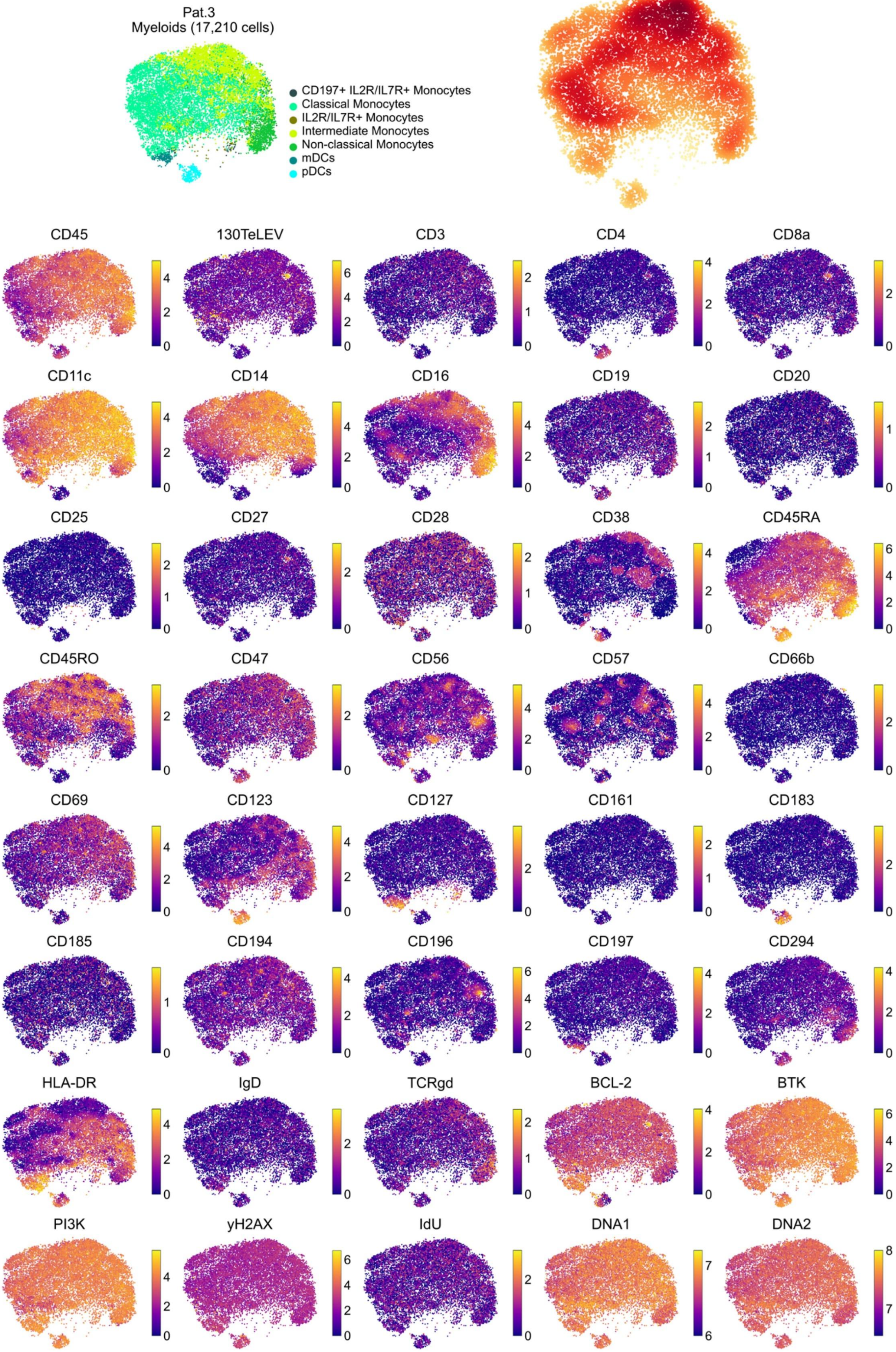

Pat.3
Myeloids (17,210 cells)
CD197+ IL2R/IL7R+ Monocytes
Classical Monocytes
IL2R/IL7R+ Monocytes
Intermediate Monocytes
Non-classical Monocytes
mDCs
pDCs
CD45
130TeLEV
CD3
CD4
CD8a
CD11c
CD14
CD16
CD19
CD20
CD25
CD27
CD28
CD38
CD45RA
CD45RO
CD47
CD56
CD57
CD66b
CD69
CD123
CD127
CD161
CD183
CD185
CD194
CD196
CD197
CD294
HLA-DR
IgD
TCRgd
BCL-2
BTK
PI3K
yH2AX
IdU
DNA1
DNA2

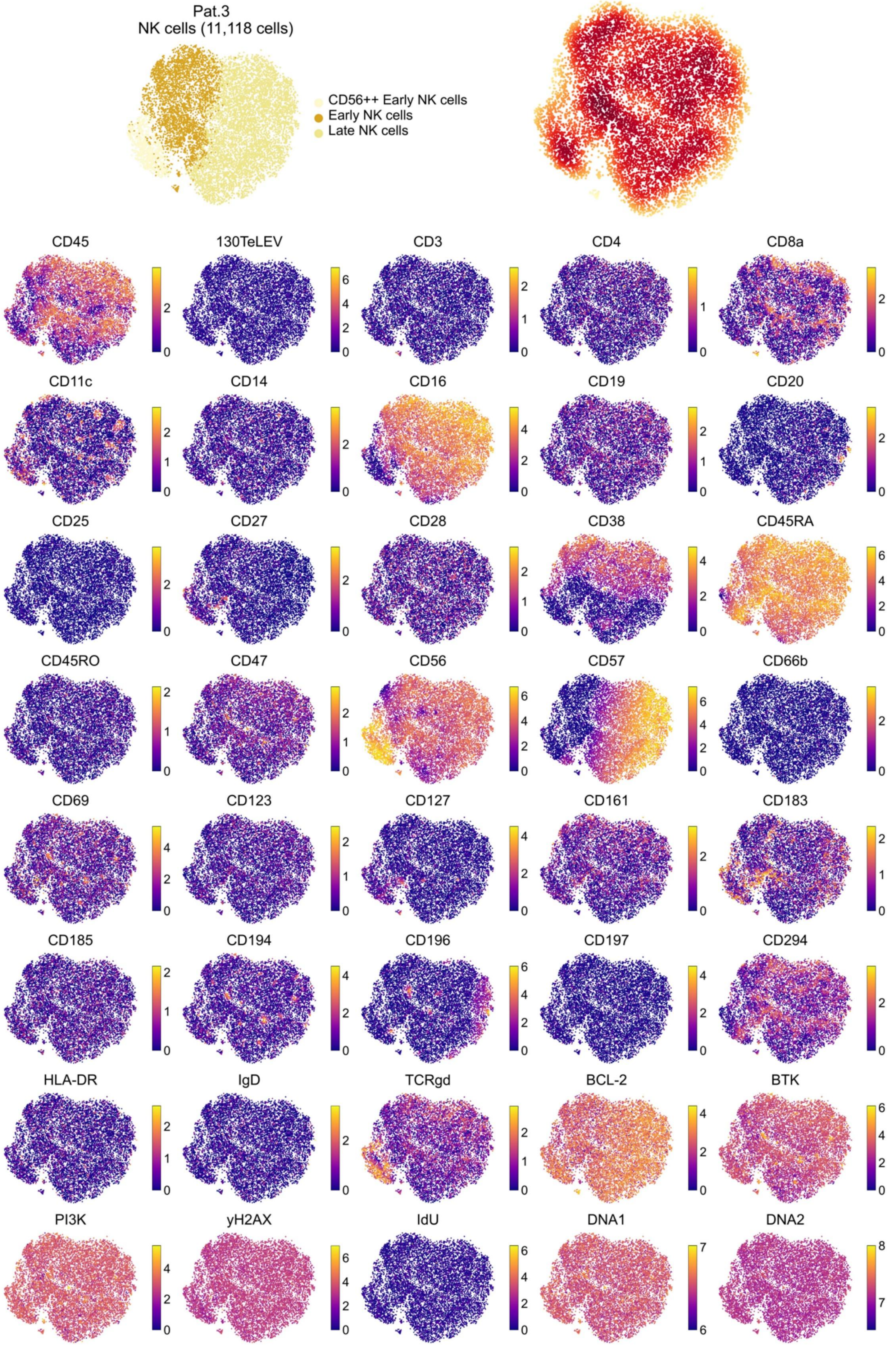
Pat.3
NK cells (11,118 cells)
CD56++ Early NK cells
Early NK cells
Late NK cells
CD45
130TeLEV
CD3
CD4
CD8a
CD11c
CD14
CD16
CD19
CD20
CD25
CD27
CD28
CD38
CD45RA
CD45RO
CD47
CD56
CD57
CD66b
CD69
CD123
CD127
CD161
CD183
CD185
CD194
CD196
CD197
CD294
HLA-DR
IgD
TCRgd
BCL-2
BTK
PI3K
yH2AX
IdU
DNA1
DNA2

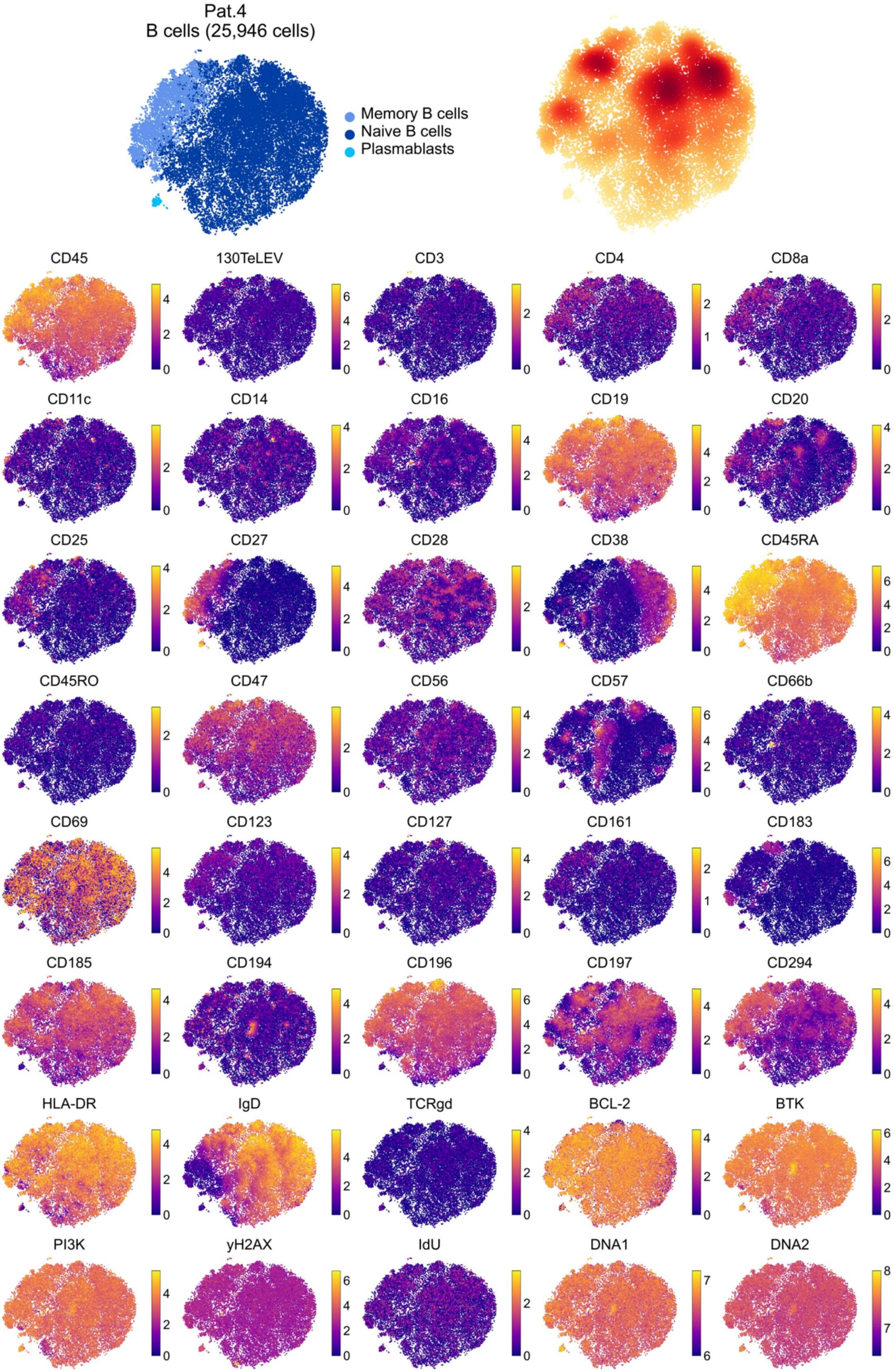
Pat.4
B cells (25,946 cells)
Memory B cells
Naive B cells
Plasmablasts
CD45
130TeLEV
CD3
CD4
CD8a
CD11c
CD14
CD16
CD19
CD20
CD25
CD27
CD28
CD38
CD45RA
CD45RO
CD47
CD56
CD57
CD66b
CD69
CD123
CD127
CD161
CD183
CD185
CD194
CD196
CD197
CD294
HLA-DR
IgD
TCRgd
BCL-2
BTK
PI3K
yH2AX
IdU
DNA1
DNA2

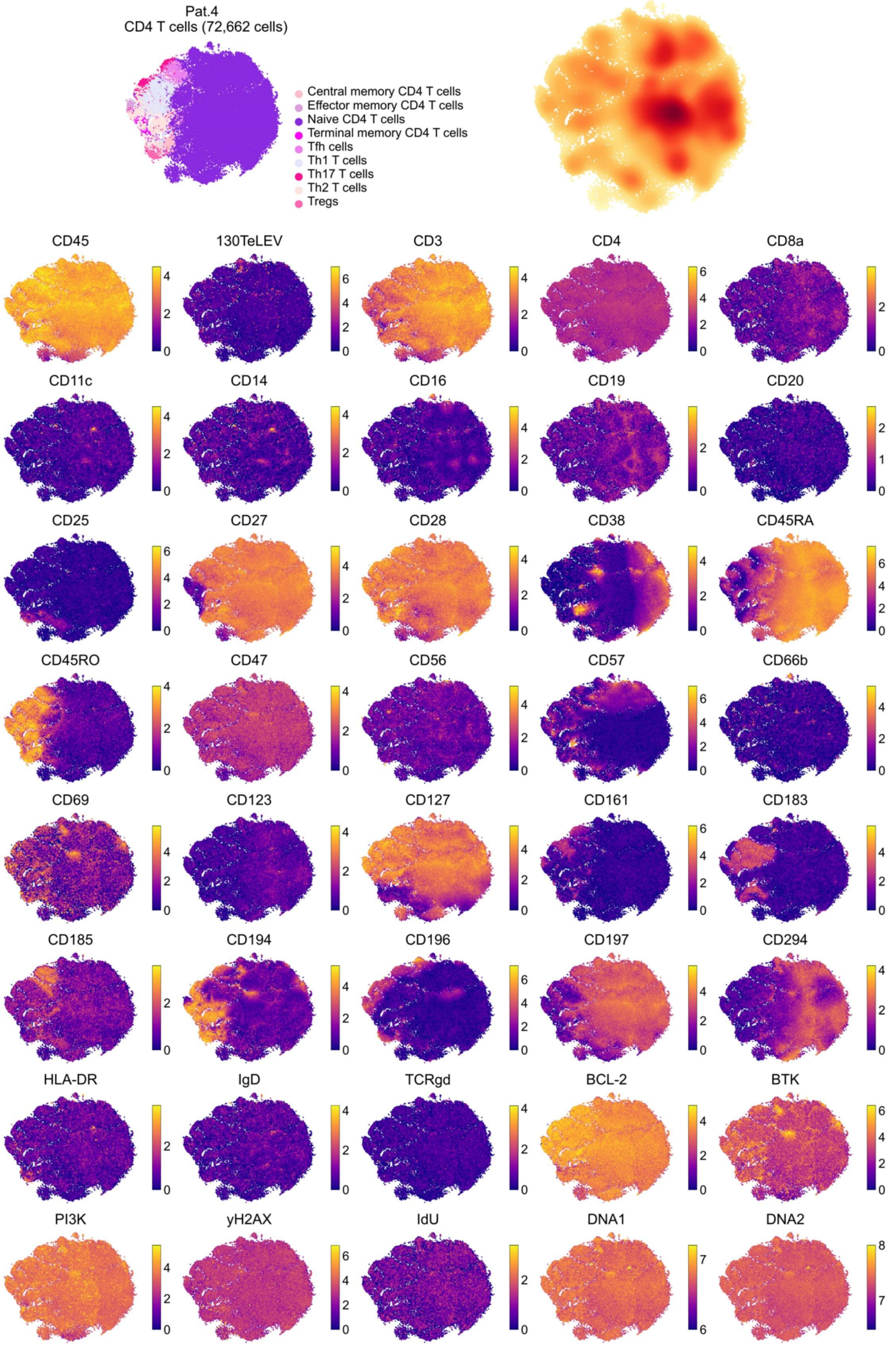

Pat.4
CD4 T cells (72,662 cells)
Central memory CD4 T cells
Effector memory CD4 T cells
Naive CD4 T cells
Terminal memory CD4 T cells
Tfh cells
Th1 T cells
Th17 T cells
Th2 T cells
Tregs
CD45
130TeLEV
CD3
CD4
CD8a
CD11c
CD14
CD16
CD19
CD20
CD25
CD27
CD28
CD38
CD45RA
CD45RO
CD47
CD56
CD57
CD66b
CD69
CD123
CD127
CD161
CD183
CD185
CD194
CD196
CD197
CD294
HLA-DR
IgD
TCRgd
BCL-2
BTK
PI3K
yH2AX
IdU
DNA1
DNA2

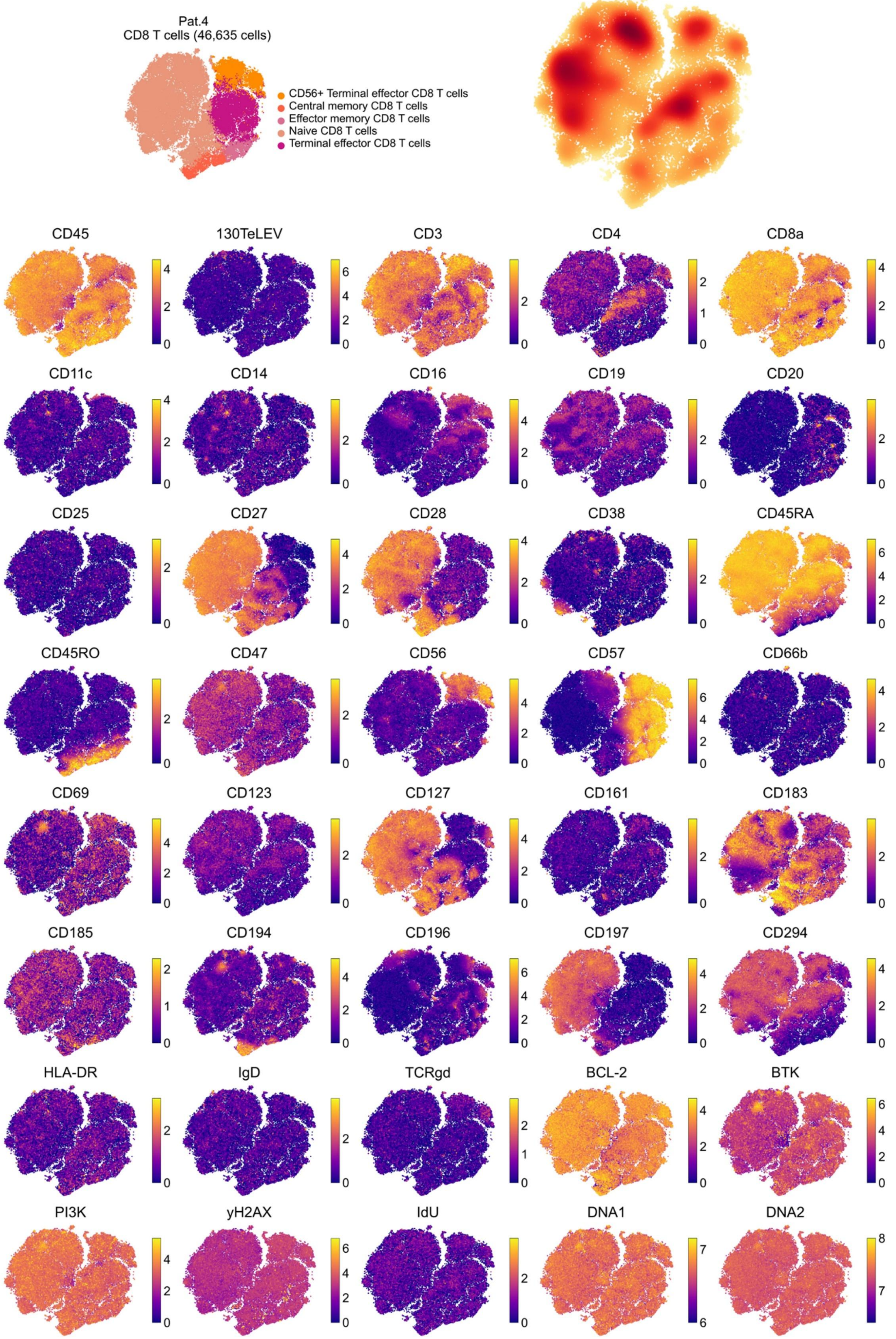

Pat.4
CD8 T cells (46,635 cells)
CD56+ Terminal effector CD8 T cells
Central memory CD8 T cells
Effector memory CD8 T cells
Naive CD8 T cells
Terminal effector CD8 T cells
CD45
130TeLEV
CD3
CD4
CD8a
CD11c
CD14
CD16
CD19
CD20
CD25
CD27
CD28
CD38
CD45RA
CD45RO
CD47
CD56
CD57
CD66b
CD69
CD123
CD127
CD161
CD183
CD185
CD194
CD196
CD197
CD294
HLA-DR
IgD
TCRgd
BCL-2
BTK
PI3K
yH2AX
IdU
DNA1
DNA2

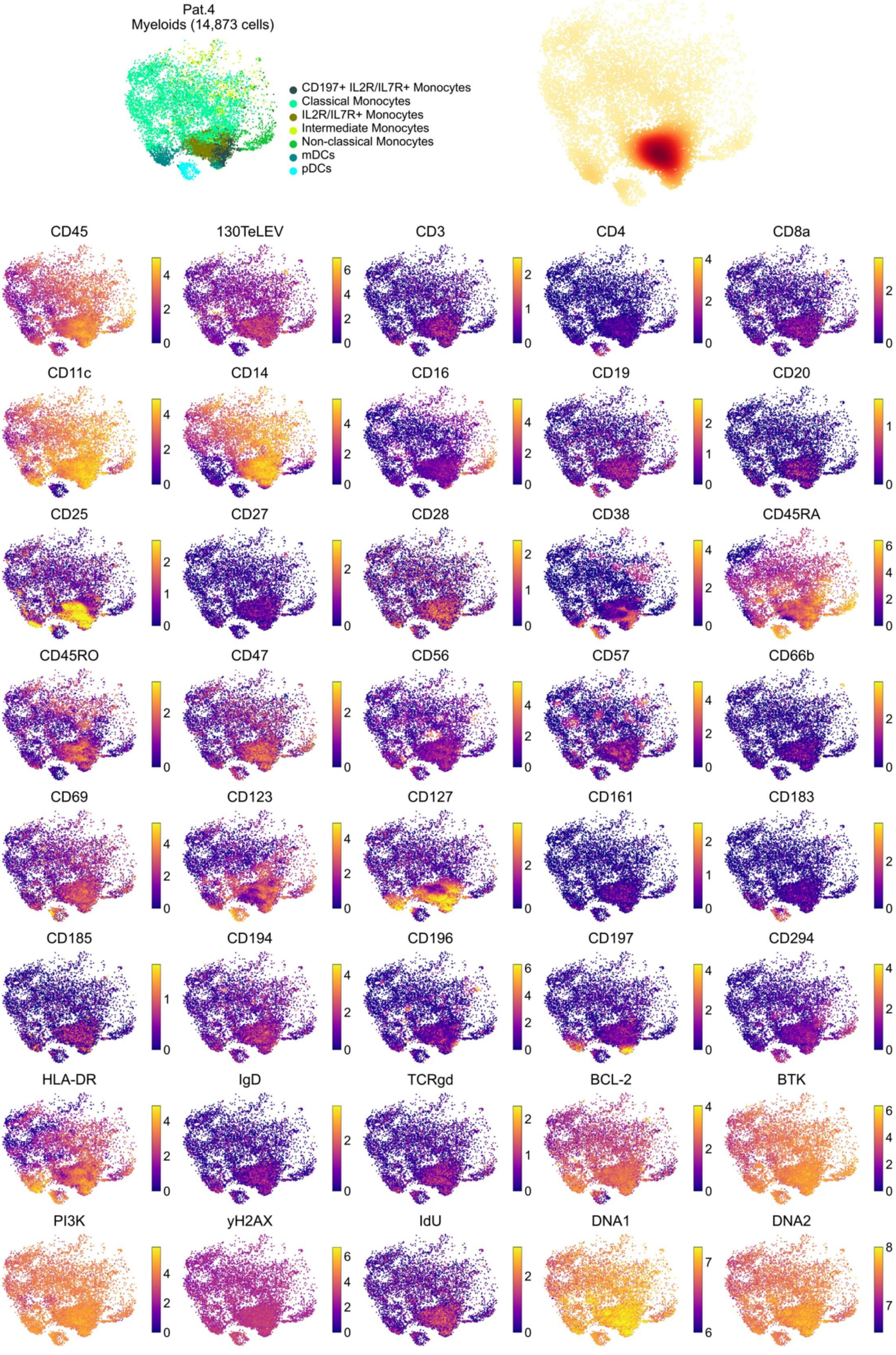
Pat.4
Myeloids (14,873 cells)
CD197+ IL2R/IL7R+ Monocytes
Classical Monocytes
IL2R/IL7R+ Monocytes
Intermediate Monocytes
Non-classical Monocytes
mDCs
pDCs
CD45
130TeLEV
CD3
CD4
CD8a
CD11c
CD14
CD16
CD19
CD20
CD25
CD27
CD28
CD38
CD45RA
CD45RO
CD47
CD56
CD57
CD66b
CD69
CD123
CD127
CD161
CD183
CD185
CD194
CD196
CD197
CD294
HLA-DR
IgD
TCRgd
BCL-2
BTK
PI3K
yH2AX
IdU
DNA1
DNA2

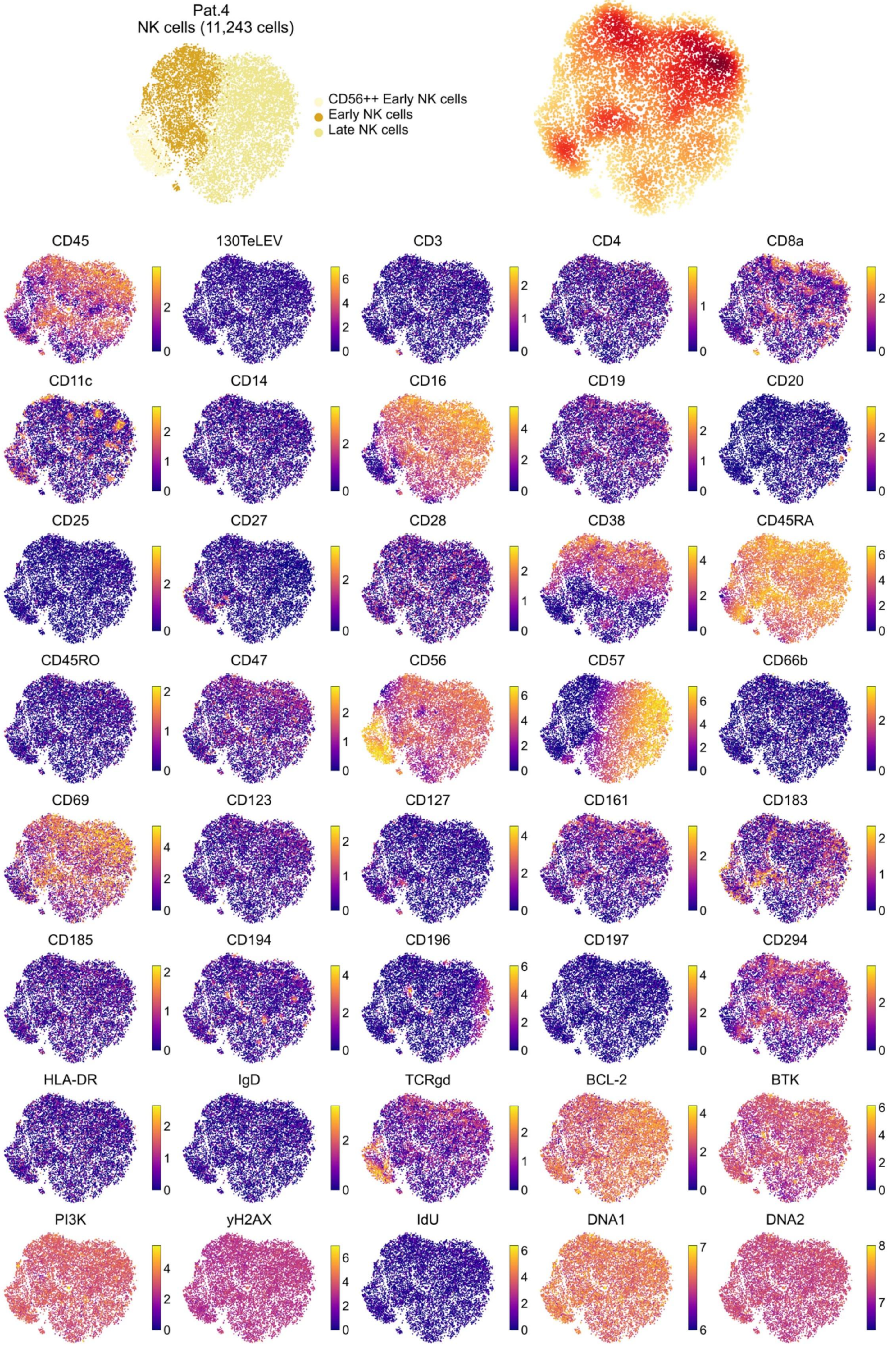

Pat.4
NK cells (11,243 cells)
CD56++ Early NK cells
Early NK cells
Late NK cells
CD45
130TeLEV
CD3
CD4
CD8a
CD11c
CD14
CD16
CD19
CD20
CD25
CD27
CD28
CD38
CD45RA
CD45RO
CD47
CD56
CD57
CD66b
CD69
CD123
CD127
CD161
CD183
CD185
CD194
CD196
CD197
CD294
HLA-DR
IgD
TCRgd
BCL-2
BTK
PI3K
yH2AX
IdU
DNA1
DNA2

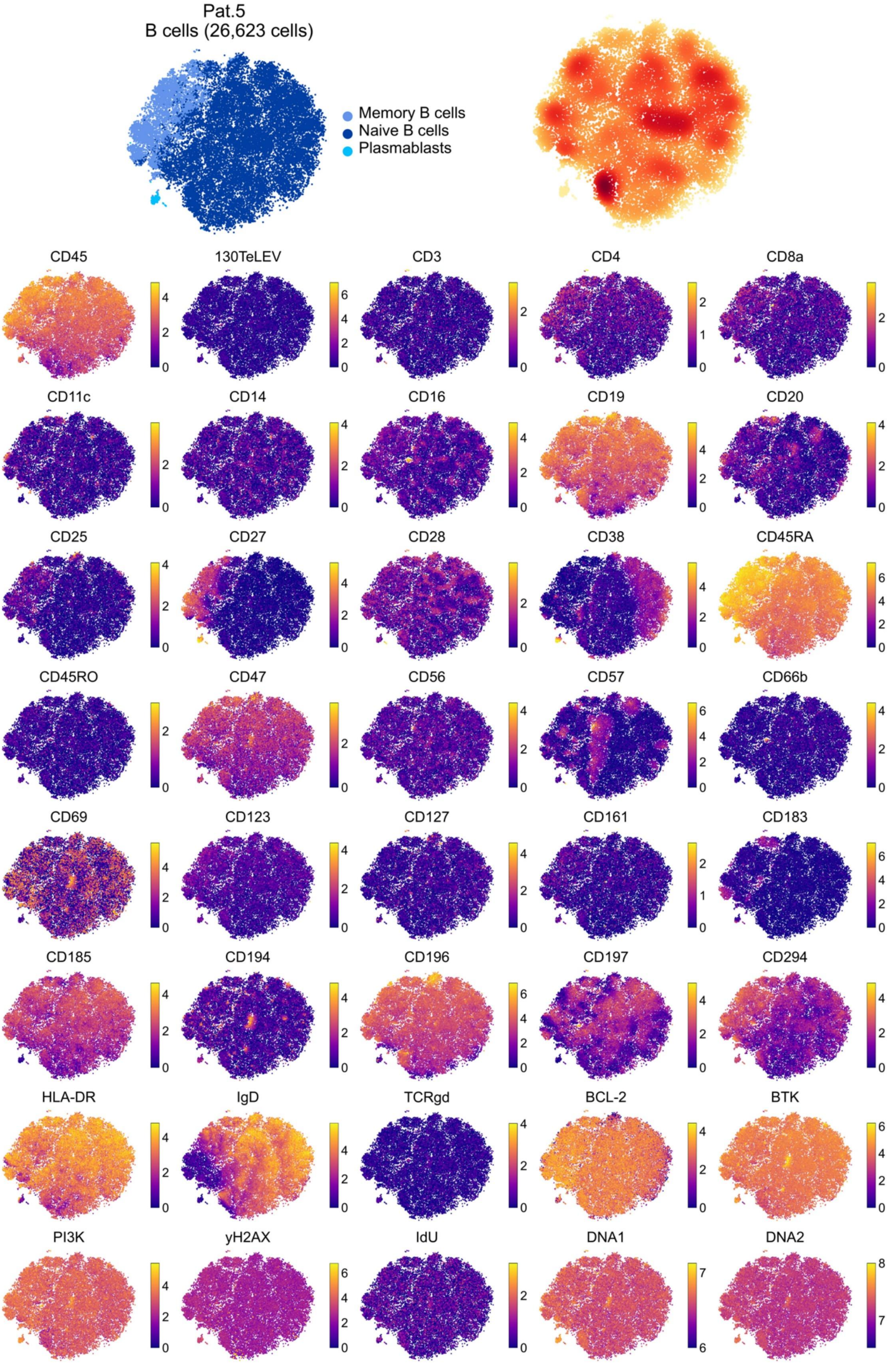
Pat.5
B cells (26,623 cells)
Memory B cells
Naive B cells
Plasmablasts
CD45
130TeLEV
CD3
CD4
CD8a
CD11c
CD14
CD16
CD19
CD20
CD25
CD27
CD28
CD38
CD45RA
CD45RO
CD47
CD56
CD57
CD66b
CD69
CD123
CD127
CD161
CD183
CD185
CD194
CD196
CD197
CD294
HLA-DR
IgD
TCRgd
BCL-2
BTK
PI3K
yH2AX
IdU
DNA1
DNA2

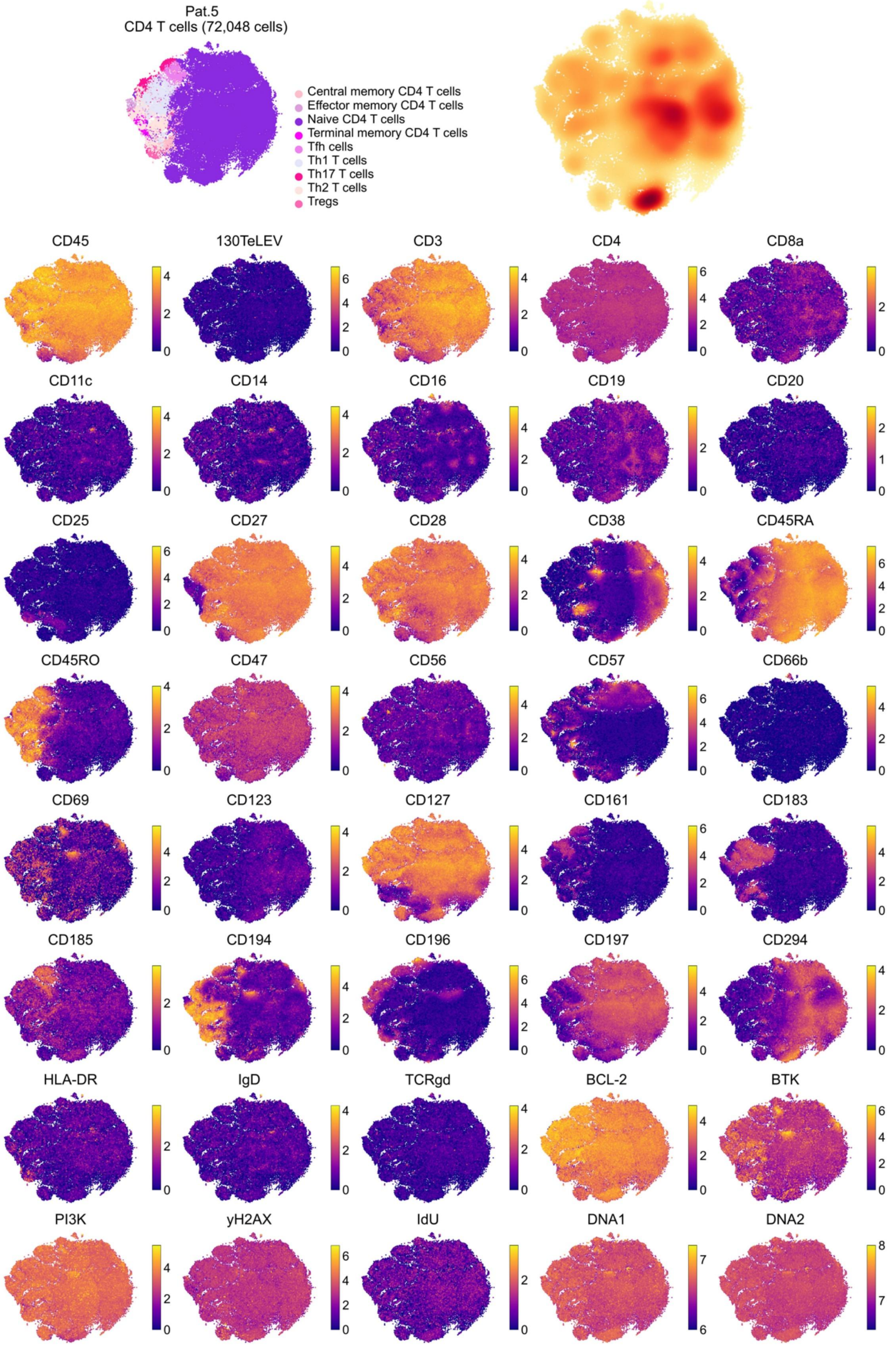
Pat.5
CD4 T cells (72,048 cells)
Central memory CD4 T cells
Effector memory CD4 T cells
Naive CD4 T cells
Terminal memory CD4 T cells
Tfh cells
Th1 T cells
Th17 T cells
Th2 T cells
Tregs
CD45
130TeLEV
CD3
CD4
CD8a
CD11c
CD14
CD16
CD19
CD20
CD25
CD27
CD28
CD38
CD45RA
CD45RO
CD47
CD56
CD57
CD66b
CD69
CD123
CD127
CD161
CD183
CD185
CD194
CD196
CD197
CD294
HLA-DR
IgD
TCRgd
BCL-2
BTK
PI3K
yH2AX
IdU
DNA1
DNA2

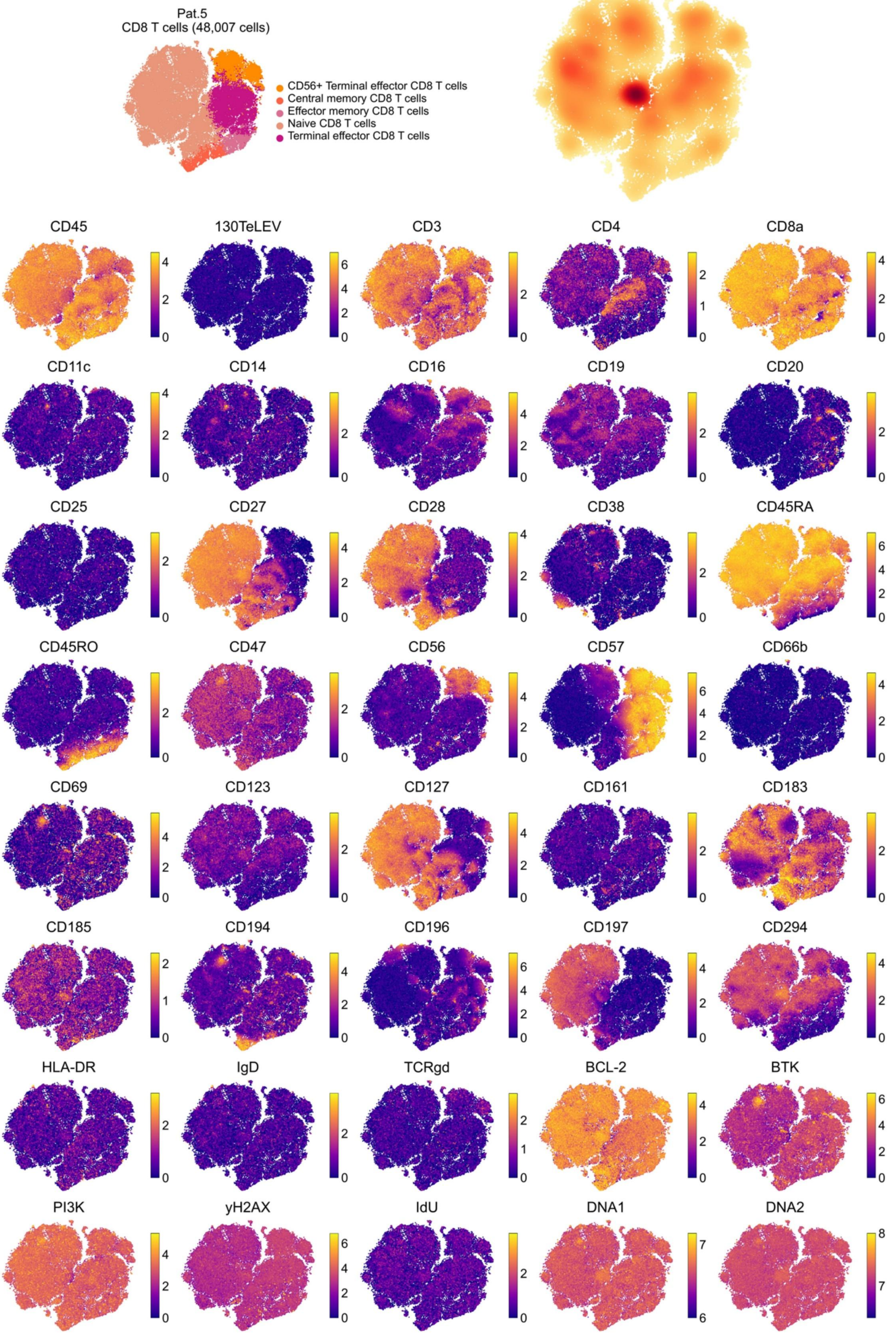
Pat.5
CD8 T cells (48,007 cells)
CD56+ Terminal effector CD8 T cells
Central memory CD8 T cells
Effector memory CD8 T cells
Naive CD8 T cells
Terminal effector CD8 T cells
CD45
130TeLEV
CD3
CD4
CD8a
CD11c
CD14
CD16
CD19
CD20
CD25
CD27
CD28
CD38
CD45RA
CD45RO
CD47
CD56
CD57
CD66b
CD69
CD123
CD127
CD161
CD183
CD185
CD194
CD196
CD197
CD294
HLA-DR
IgD
TCRgd
BCL-2
BTK
PI3K
yH2AX
IdU
DNA1
DNA2

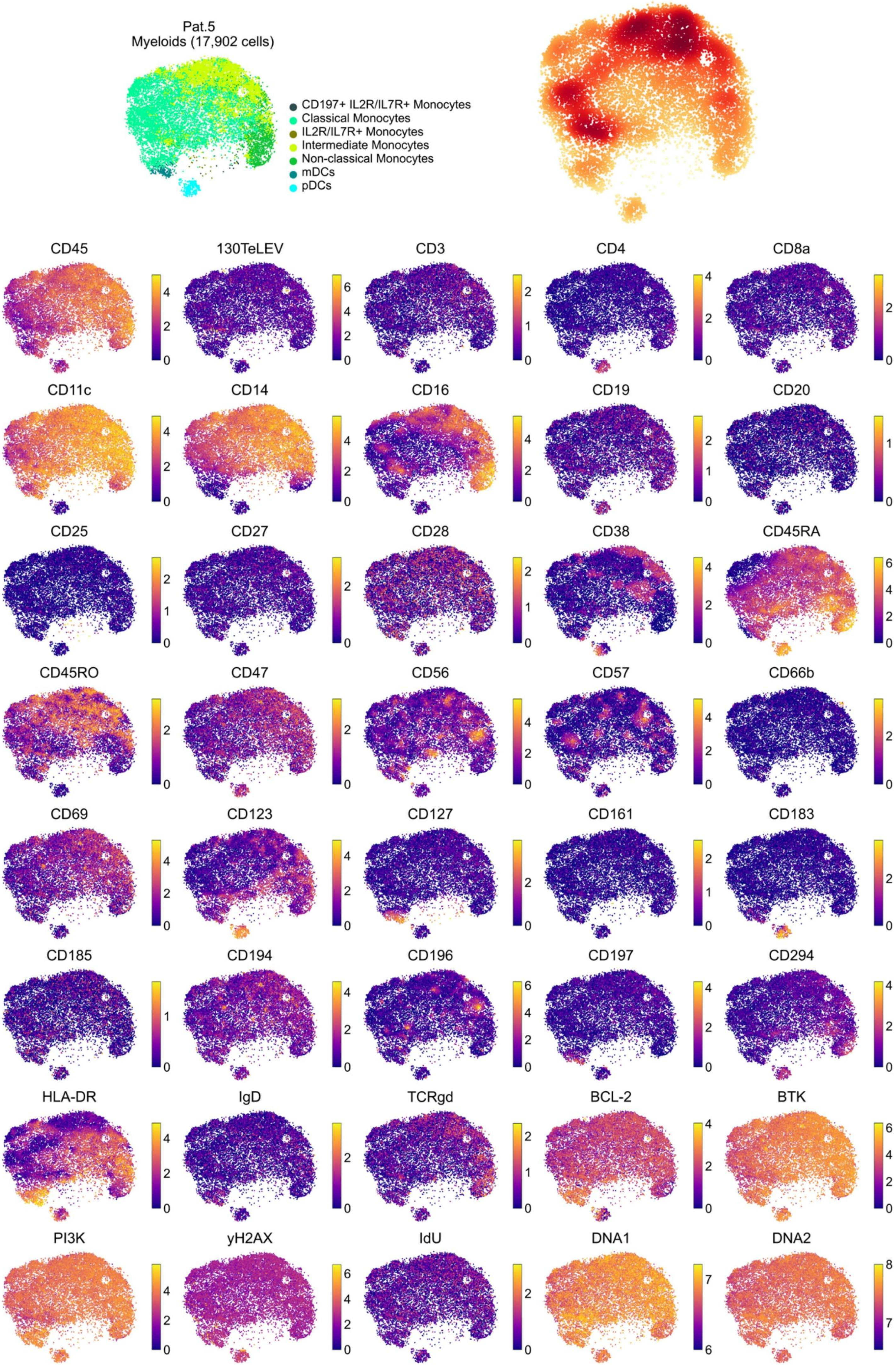
Pat.5
Myeloids (17,902 cells)
CD197+ IL2R/IL7R+ Monocytes
Classical Monocytes
IL2R/IL7R+ Monocytes
Intermediate Monocytes
Non-classical Monocytes
mDCs
pDCs
CD45
130TeLEV
CD3
CD4
CD8a
CD11c
CD14
CD16
CD19
CD20
CD25
CD27
CD28
CD38
CD45RA
CD45RO
CD47
CD56
CD57
CD66b
CD69
CD123
CD127
CD161
CD183
CD185
CD194
CD196
CD197
CD294
HLA-DR
IgD
TCRgd
BCL-2
BTK
PI3K
yH2AX
IdU
DNA1
DNA2

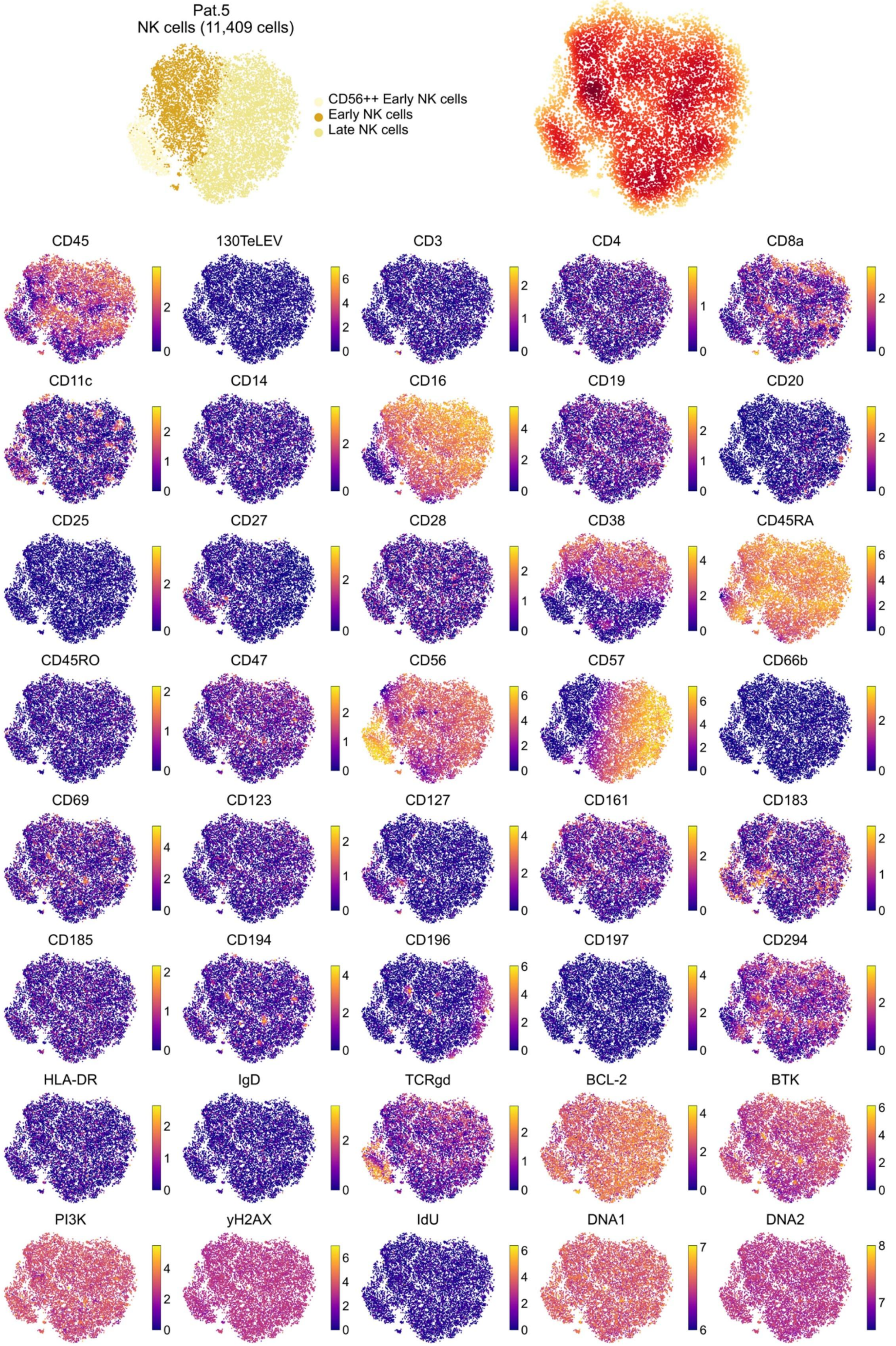
Pat.5
NK cells (11,409 cells)
CD56++ Early NK cells
Early NK cells
Late NK cells
CD45
130TeLEV
CD3
CD4
CD8a
CD11c
CD14
CD16
CD19
CD20
CD25
CD27
CD28
CD38
CD45RA
CD45RO
CD47
CD56
CD57
CD66b
CD69
CD123
CD127
CD161
CD183
CD185
CD194
CD196
CD197
CD294
HLA-DR
IgD
TCRgd
BCL-2
BTK
PI3K
yH2AX
IdU
DNA1
DNA2

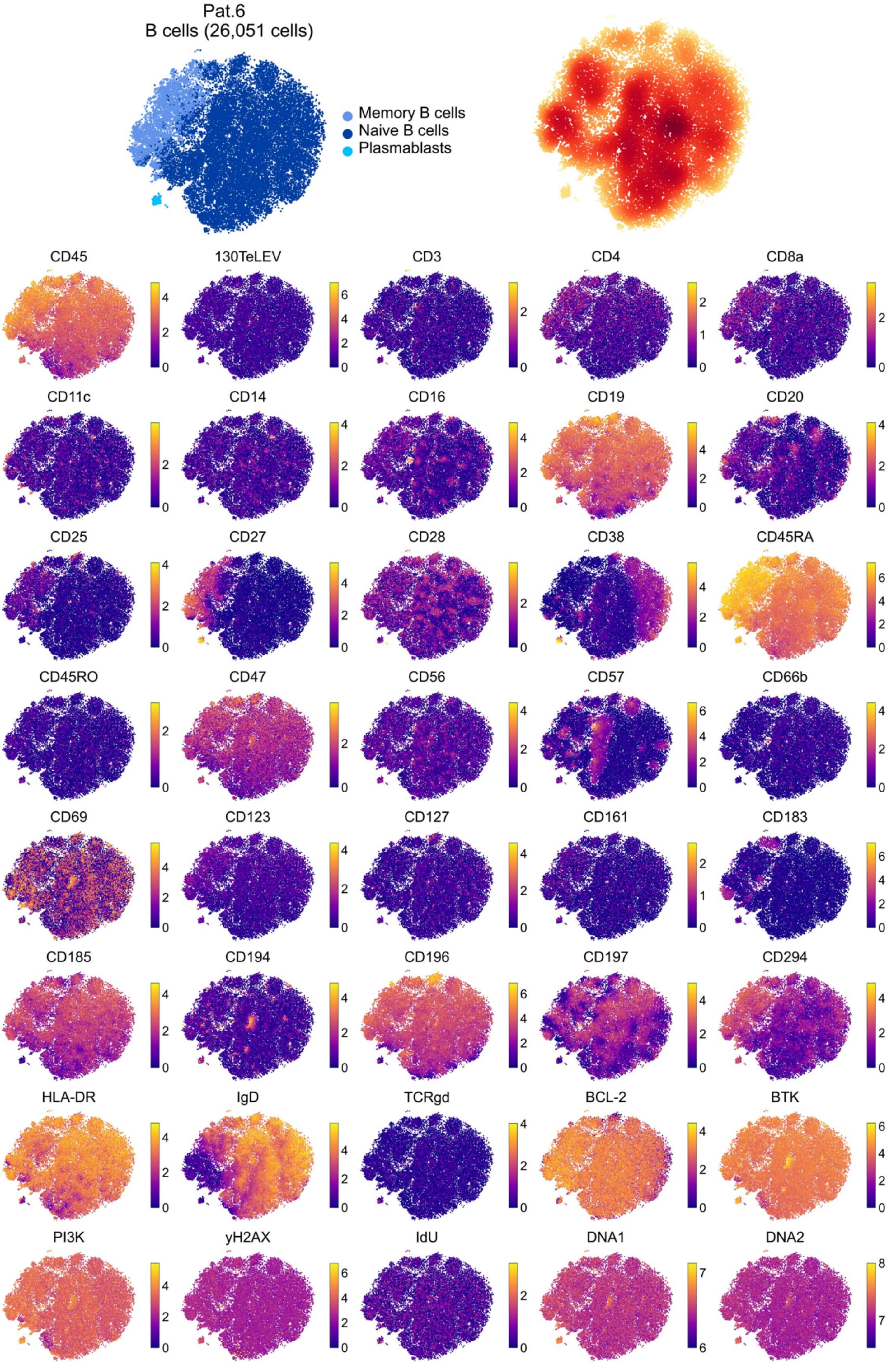
Pat.6
B cells (26,051 cells)
Memory B cells
Naive B cells
Plasmablasts
CD45
130TeLEV
CD3
CD4
CD8a
CD11c
CD14
CD16
CD19
CD20
CD25
CD27
CD28
CD38
CD45RA
CD45RO
CD47
CD56
CD57
CD66b
CD69
CD123
CD127
CD161
CD183
CD185
CD194
CD196
CD197
CD294
HLA-DR
IgD
TCRgd
BCL-2
BTK
PI3K
yH2AX
IdU
DNA1
DNA2

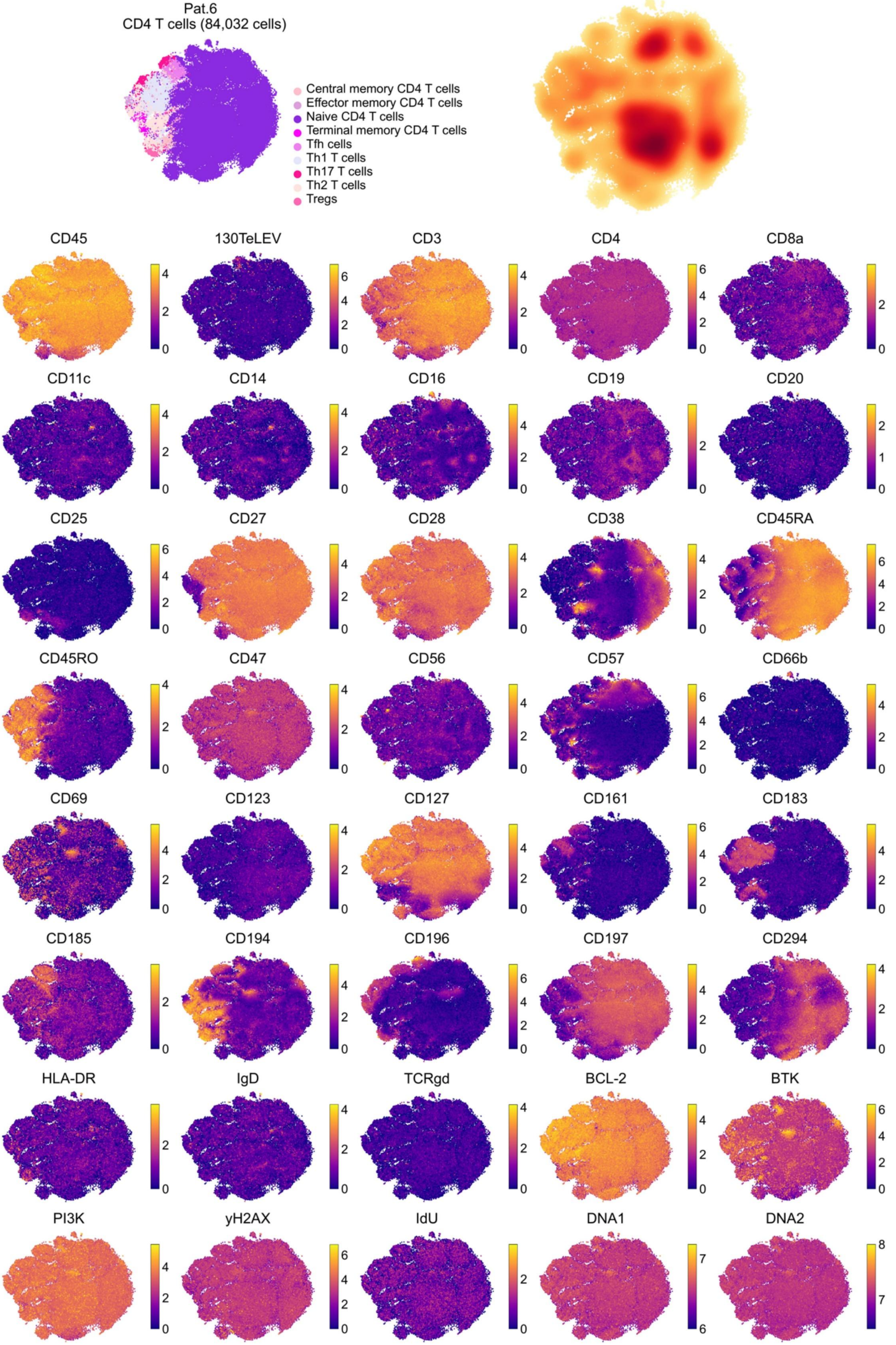
Pat.6
CD4 T cells (84,032 cells)
Central memory CD4 T cells
Effector memory CD4 T cells
Naive CD4 T cells
Terminal memory CD4 T cells
Tfh cells
Th1 T cells
Th17 T cells
Th2 T cells
Tregs
CD45
130TeLEV
CD3
CD4
CD8a
CD11c
CD14
CD16
CD19
CD20
CD25
CD27
CD28
CD38
CD45RA
CD45RO
CD47
CD56
CD57
CD66b
CD69
CD123
CD127
CD161
CD183
CD185
CD194
CD196
CD197
CD294
HLA-DR
IgD
TCRgd
BCL-2
BTK
PI3K
yH2AX
IdU
DNA1
DNA2

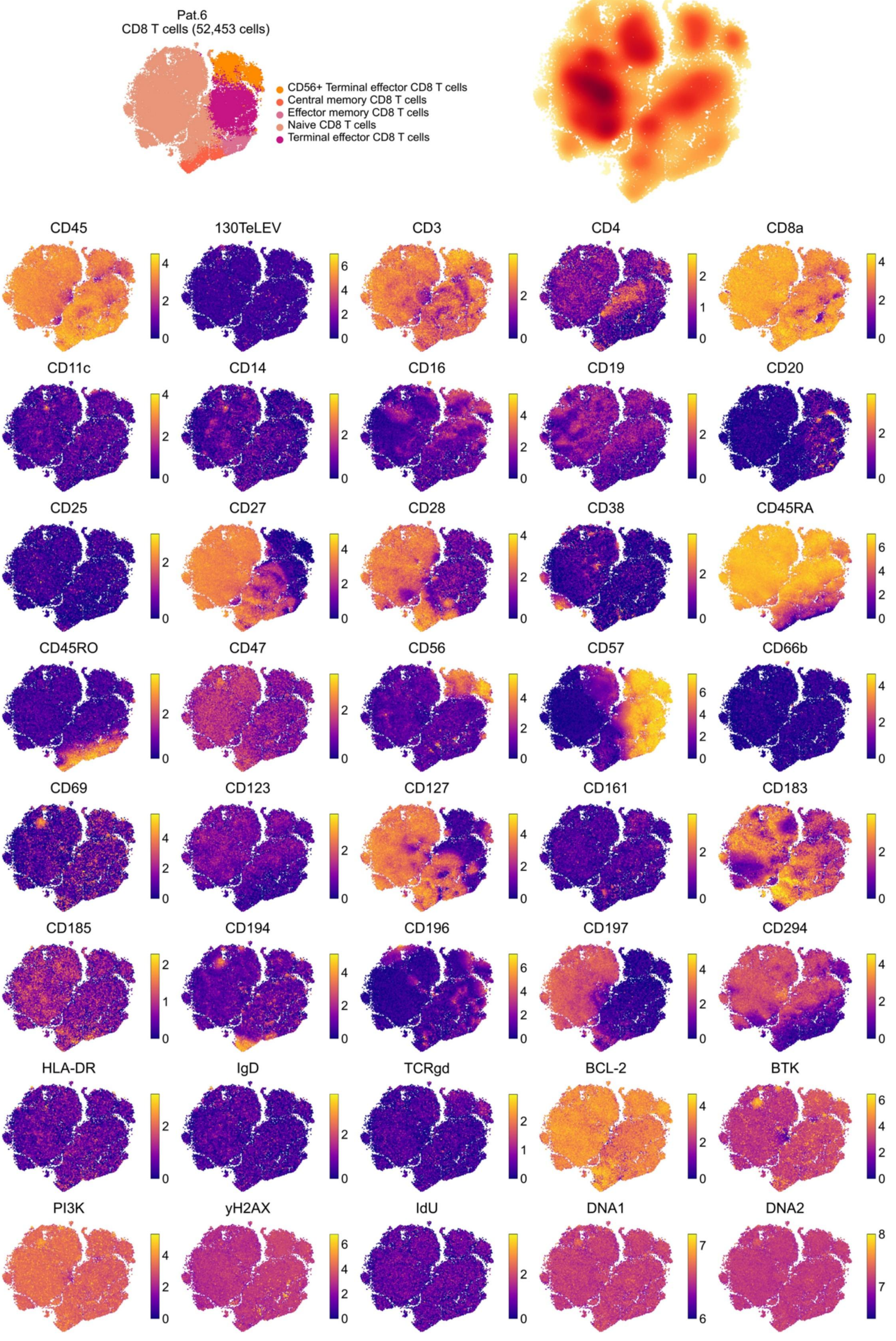

Pat.6
CD8 T cells (52,453 cells)
CD56+ Terminal effector CD8 T cells
Central memory CD8 T cells
Effector memory CD8 T cells
Naive CD8 T cells
Terminal effector CD8 T cells
CD45
130TeLEV
CD3
CD4
CD8a
CD11c
CD14
CD16
CD19
CD20
CD25
CD27
CD28
CD38
CD45RA
CD45RO
CD47
CD56
CD57
CD66b
CD69
CD123
CD127
CD161
CD183
CD185
CD194
CD196
CD197
CD294
HLA-DR
IgD
TCRgd
BCL-2
BTK
PI3K
yH2AX
IdU
DNA1
DNA2

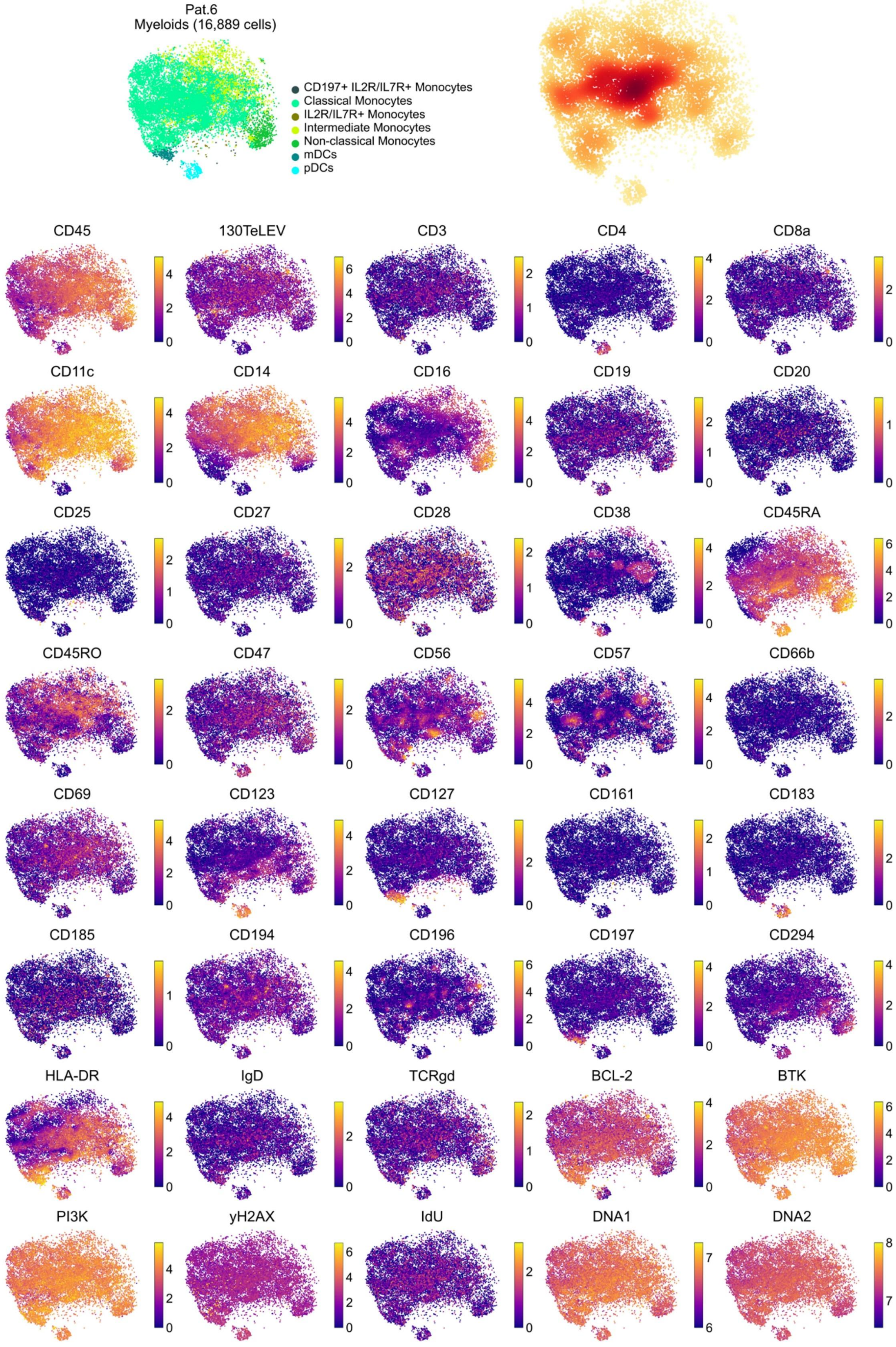

Pat.6
Myeloids (16,889 cells)
CD197+ IL2R/IL7R+ Monocytes
Classical Monocytes
IL2R/IL7R+ Monocytes
Intermediate Monocytes
Non-classical Monocytes
mDCs
pDCs
CD45
130TeLEV
CD3
CD4
CD8a
CD11c
CD14
CD16
CD19
CD20
CD25
CD27
CD28
CD38
CD45RA
CD45RO
CD47
CD56
CD57
CD66b
CD69
CD123
CD127
CD161
CD183
CD185
CD194
CD196
CD197
CD294
HLA-DR
IgD
TCRgd
BCL-2
BTK
PI3K
yH2AX
IdU
DNA1
DNA2

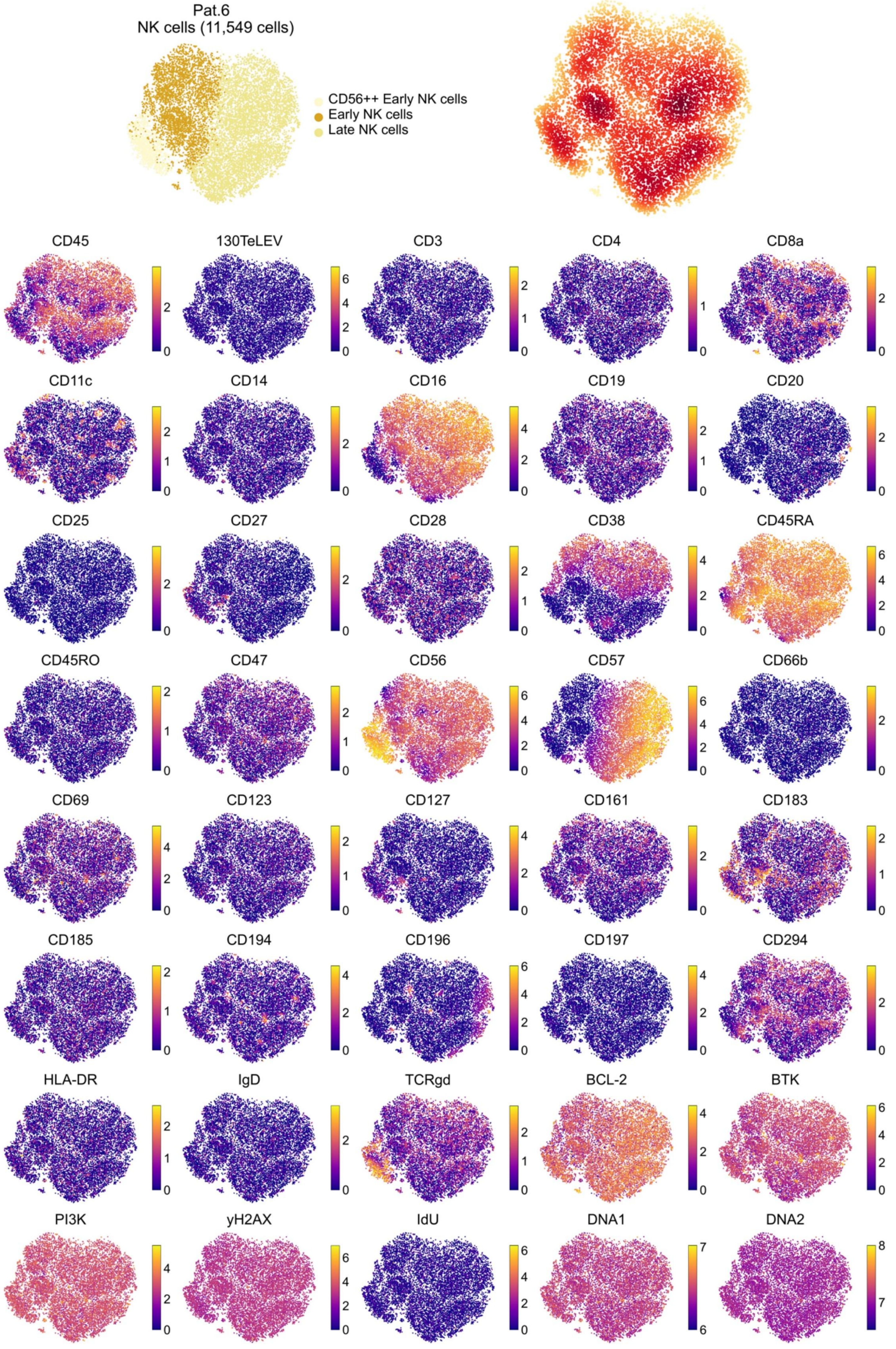

Pat.6
NK cells (11,549 cells)
CD56++ Early NK cells
Early NK cells
Late NK cells
CD45
130TeLEV
CD3
CD4
CD8a
CD11c
CD14
CD16
CD19
CD20
CD25
CD27
CD28
CD38
CD45RA
CD45RO
CD47
CD56
CD57
CD66b
CD69
CD123
CD127
CD161
CD183
CD185
CD194
CD196
CD197
CD294
HLA-DR
IgD
TCRgd
BCL-2
BTK
PI3K
yH2AX
IdU
DNA1
DNA2

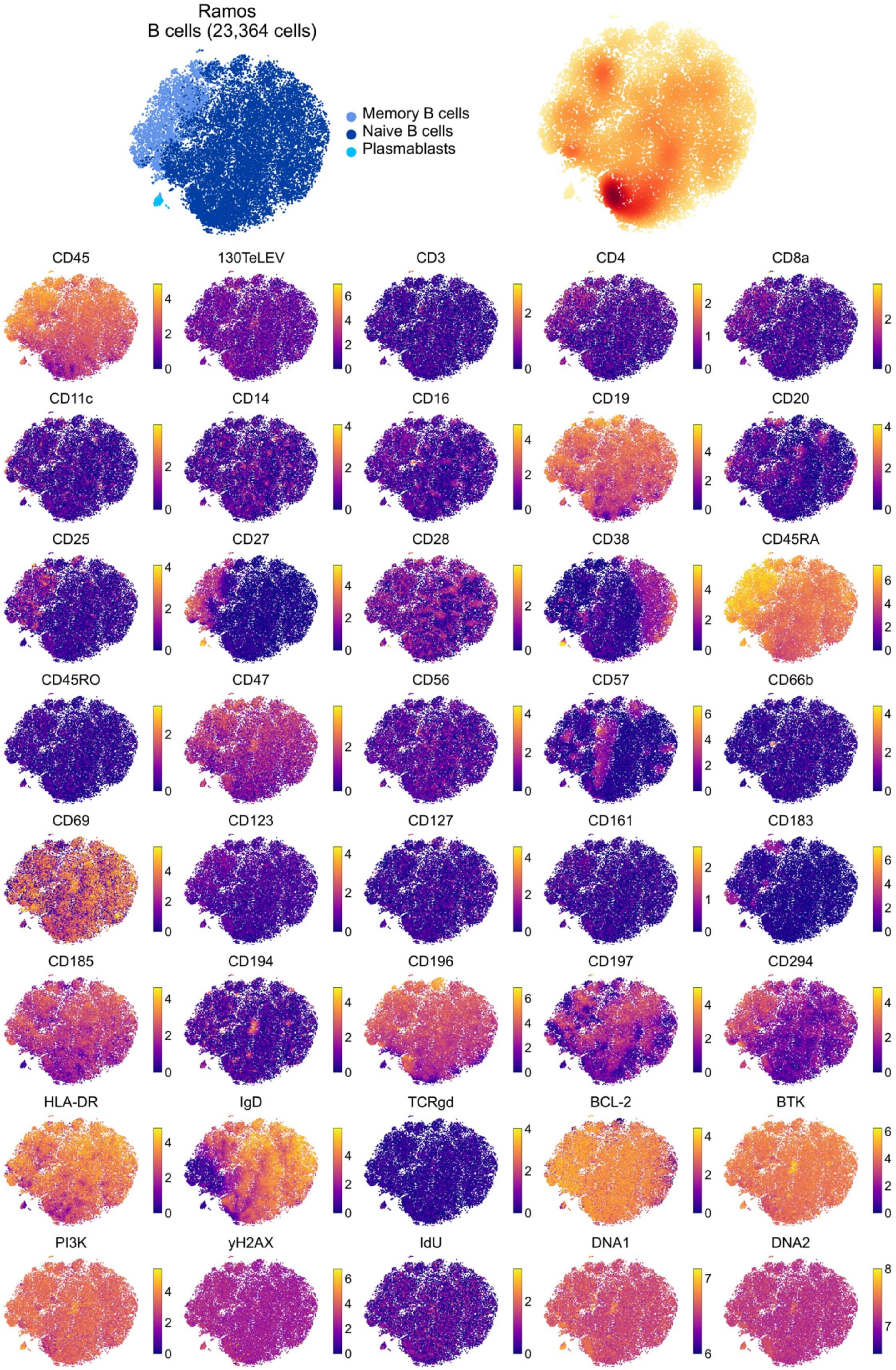
Ramos
B cells (23,364 cells)
Memory B cells
Naive B cells
Plasmablasts
CD45
130TeLEV
CD3
CD4
CD8a
CD11c
CD14
CD16
CD19
CD20
CD25
CD27
CD28
CD38
CD45RA
CD45RO
CD47
CD56
CD57
CD66b
CD69
CD123
CD127
CD161
CD183
CD185
CD194
CD196
CD197
CD294
HLA-DR
IgD
TCRgd
BCL-2
BTK
PI3K
yH2AX
IdU
DNA1
DNA2

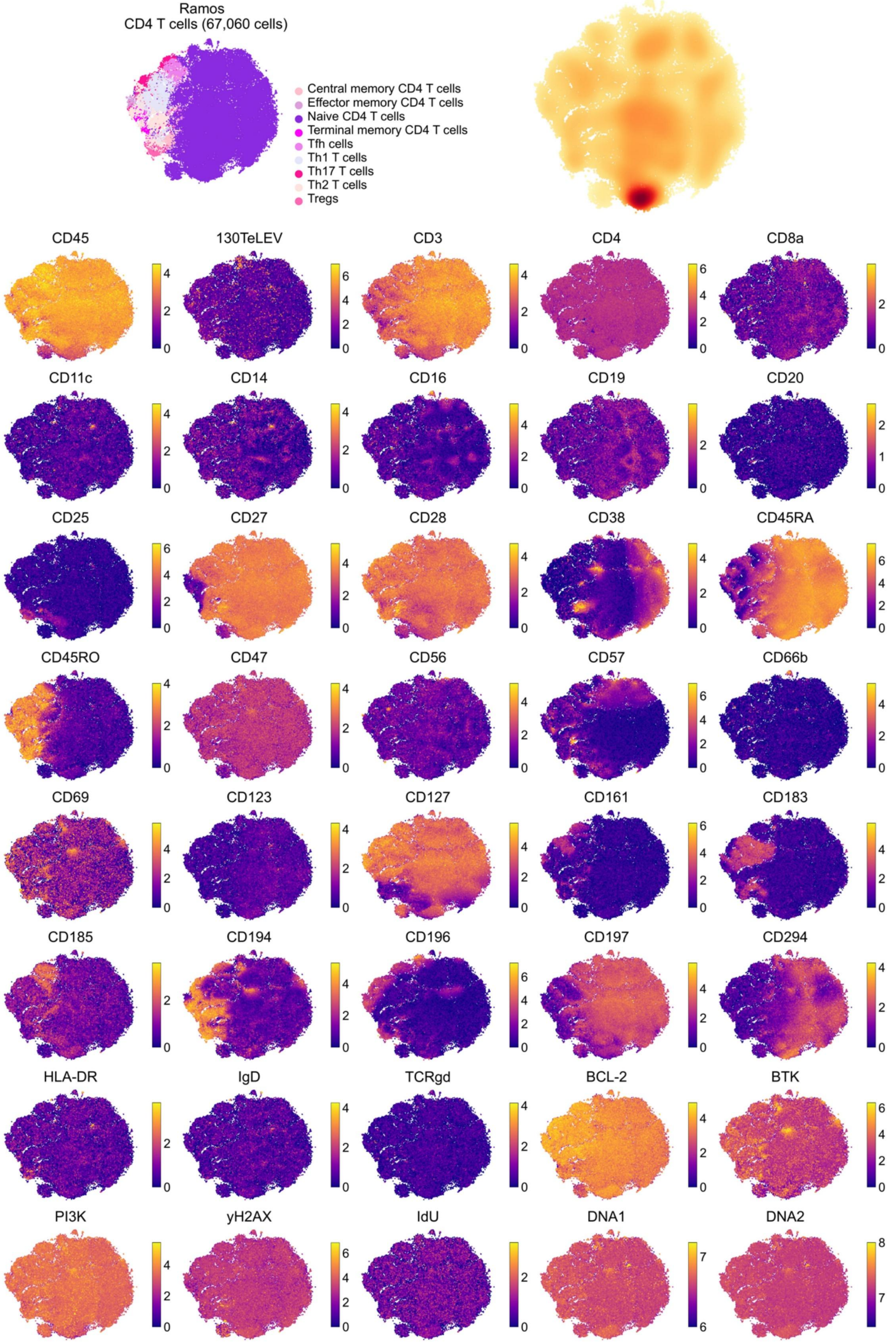
Ramos
CD4 T cells (67,060 cells)
Central memory CD4 T cells
Effector memory CD4 T cells
Naive CD4 T cells
Terminal memory CD4 T cells
Tfh cells
Th1 T cells
Th17 T cells
Th2 T cells
Tregs
CD45
130TeLEV
CD3
CD4
CD8a
CD11c
CD14
CD16
CD19
CD20
CD25
CD27
CD28
CD38
CD45RA
CD45RO
CD47
CD56
CD57
CD66b
CD69
CD123
CD127
CD161
CD183
CD185
CD194
CD196
CD197
CD294
HLA-DR
IgD
TCRgd
BCL-2
BTK
PI3K
yH2AX
IdU
DNA1
DNA2

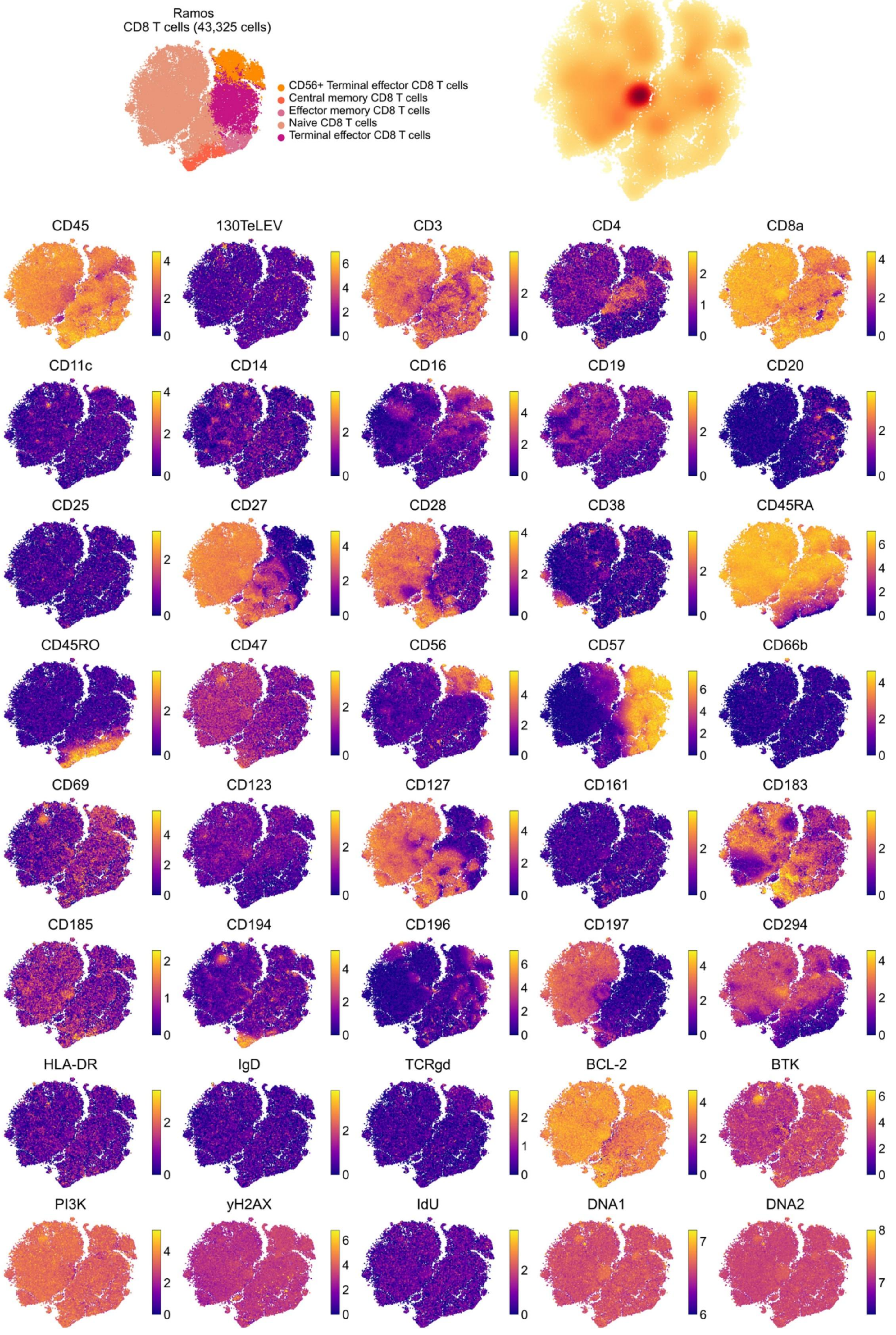

Ramos
CD8 T cells (43,325 cells)
CD56+ Terminal effector CD8 T cells
Central memory CD8 T cells
Effector memory CD8 T cells
Naive CD8 T cells
Terminal effector CD8 T cells
CD45
130TeLEV
CD3
CD4
CD8a
CD11c
CD14
CD16
CD19
CD20
CD25
CD27
CD28
CD38
CD45RA
CD45RO
CD47
CD56
CD57
CD66b
CD69
CD123
CD127
CD161
CD183
CD185
CD194
CD196
CD197
CD294
HLA-DR
IgD
TCRgd
BCL-2
BTK
PI3K
yH2AX
IdU
DNA1
DNA2

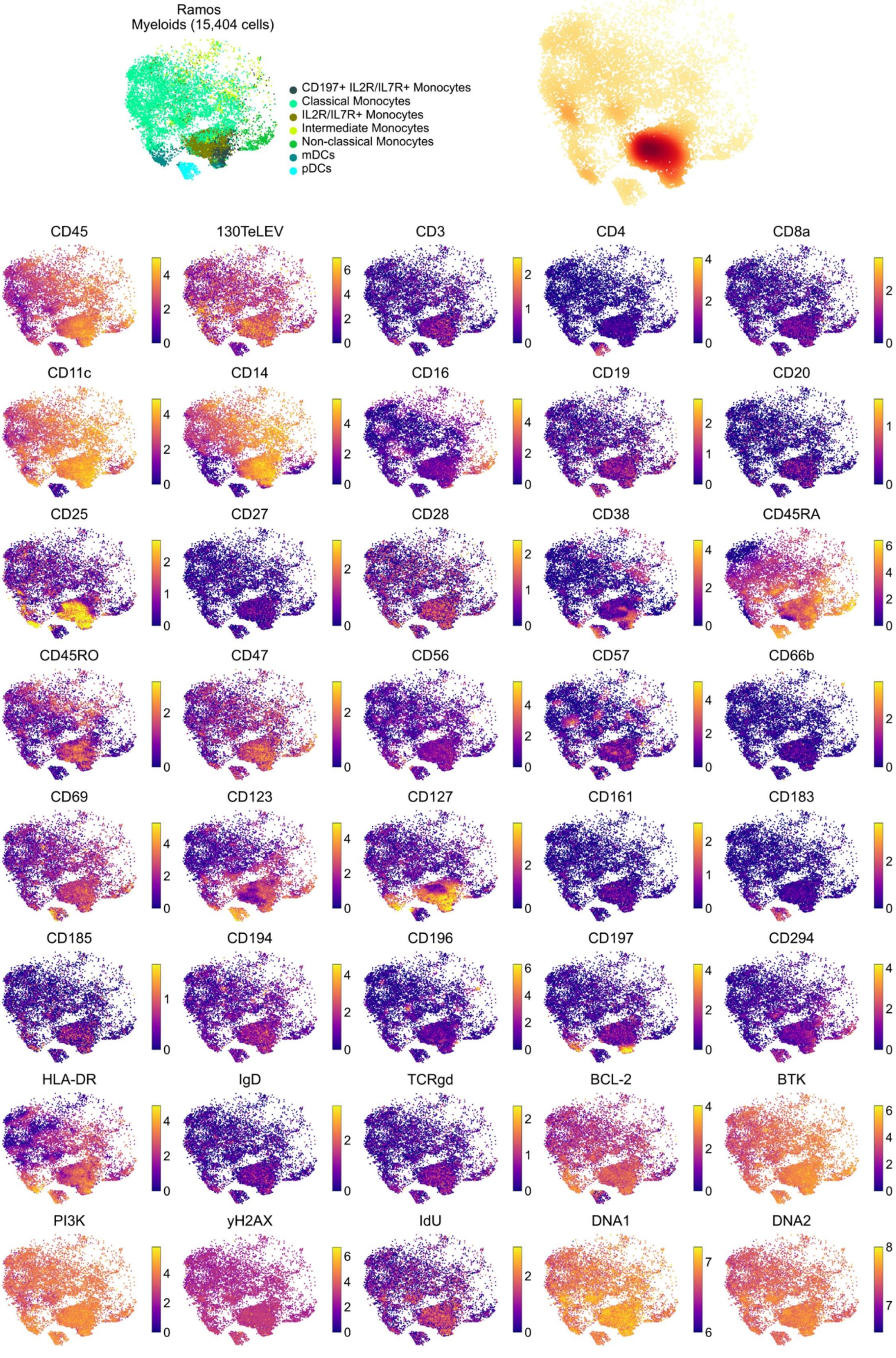
Ramos
Myeloids (15,404 cells)
CD197+ IL2R/IL7R+ Monocytes
Classical Monocytes
IL2R/IL7R+ Monocytes
Intermediate Monocytes
Non-classical Monocytes
mDCs
pDCs
CD45
130TeLEV
CD3
CD4
CD8a
CD11c
CD14
CD16
CD19
CD20
CD25
CD27
CD28
CD38
CD45RA
CD45RO
CD47
CD56
CD57
CD66b
CD69
CD123
CD127
CD161
CD183
CD185
CD194
CD196
CD197
CD294
HLA-DR
IgD
TCRgd
BCL-2
BTK
PI3K
yH2AX
IdU
DNA1
DNA2

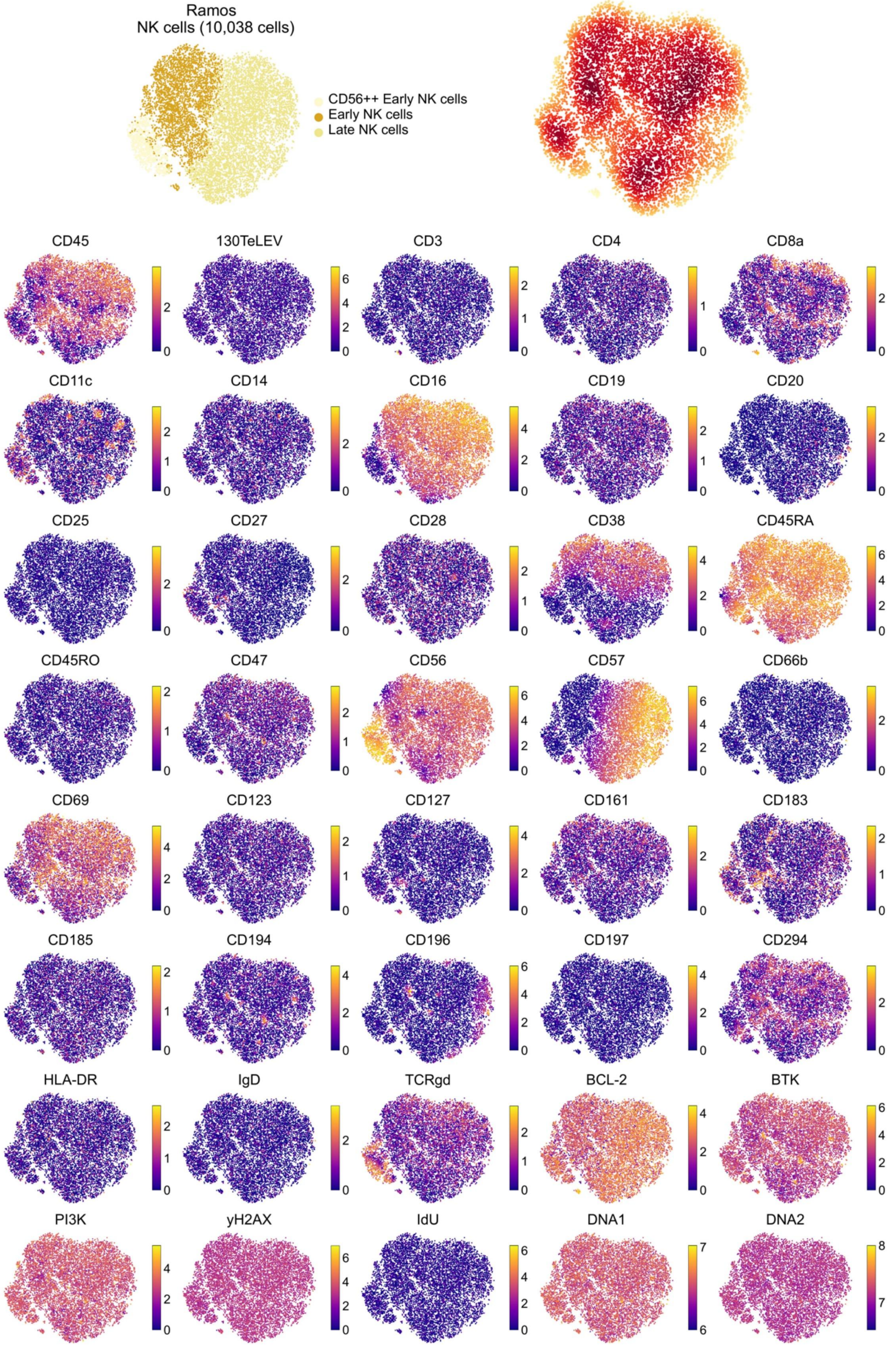

Ramos
NK cells (10,038 cells)
CD56++ Early NK cells
Early NK cells
Late NK cells
CD45
130TeLEV
CD3
CD4
CD8a
CD11c
CD14
CD16
CD19
CD20
CD25
CD27
CD28
CD38
CD45RA
CD45RO
CD47
CD56
CD57
CD66b
CD69
CD123
CD127
CD161
CD183
CD185
CD194
CD196
CD197
CD294
HLA-DR
IgD
TCRgd
BCL-2
BTK
PI3K
yH2AX
IdU
DNA1
DNA2

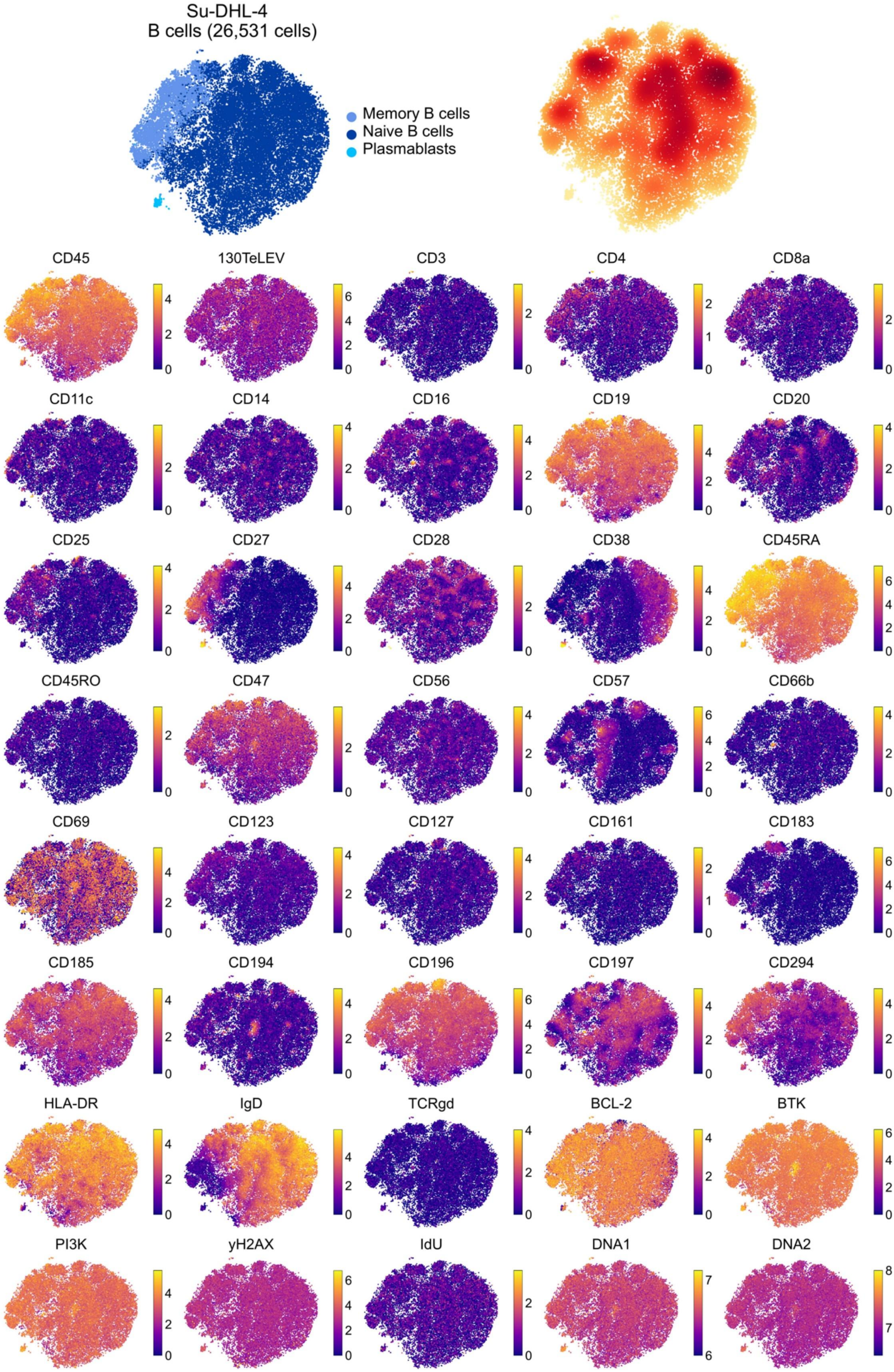
Su-DHL-4
B cells (26,531 cells)
Memory B cells
Naive B cells
Plasmablasts
CD45
130TeLEV
CD3
CD4
CD8a
CD11c
CD14
CD16
CD19
CD20
CD25
CD27
CD28
CD38
CD45RA
CD45RO
CD47
CD56
CD57
CD66b
CD69
CD123
CD127
CD161
CD183
CD185
CD194
CD196
CD197
CD294
HLA-DR
IgD
TCRgd
BCL-2
BTK
PI3K
yH2AX
IdU
DNA1
DNA2

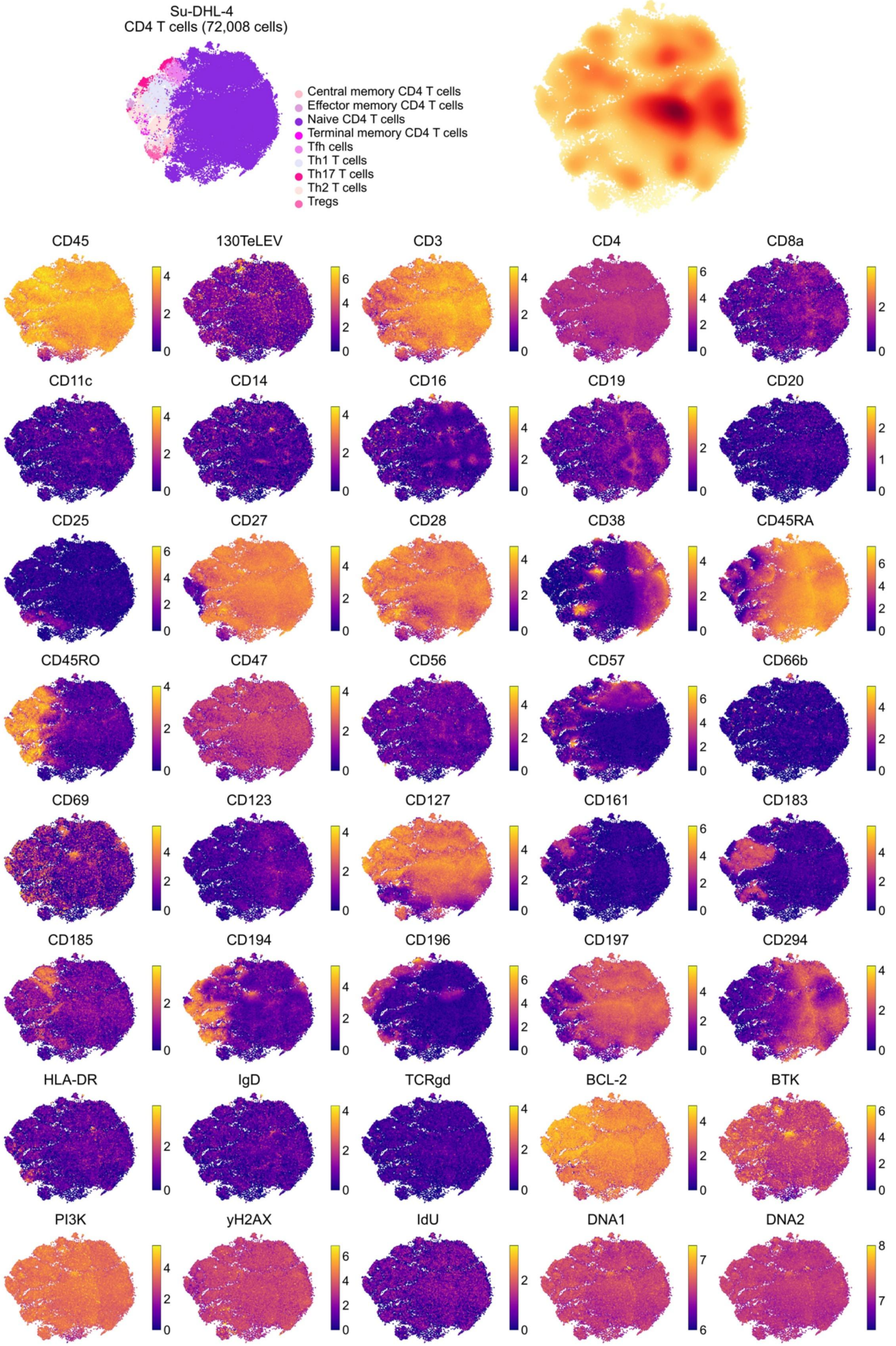

Su-DHL-4
CD4 T cells (72,008 cells)
Central memory CD4 T cells
Effector memory CD4 T cells
Naive CD4 T cells
Terminal memory CD4 T cells
Tfh cells
Th1 T cells
Th17 T cells
Th2 T cells
Tregs
CD45
130TeLEV
CD3
CD4
CD8a
CD11c
CD14
CD16
CD19
CD20
CD25
CD27
CD28
CD38
CD45RA
CD45RO
CD47
CD56
CD57
CD66b
CD69
CD123
CD127
CD161
CD183
CD185
CD194
CD196
CD197
CD294
HLA-DR
IgD
TCRgd
BCL-2
BTK
PI3K
yH2AX
IdU
DNA1
DNA2

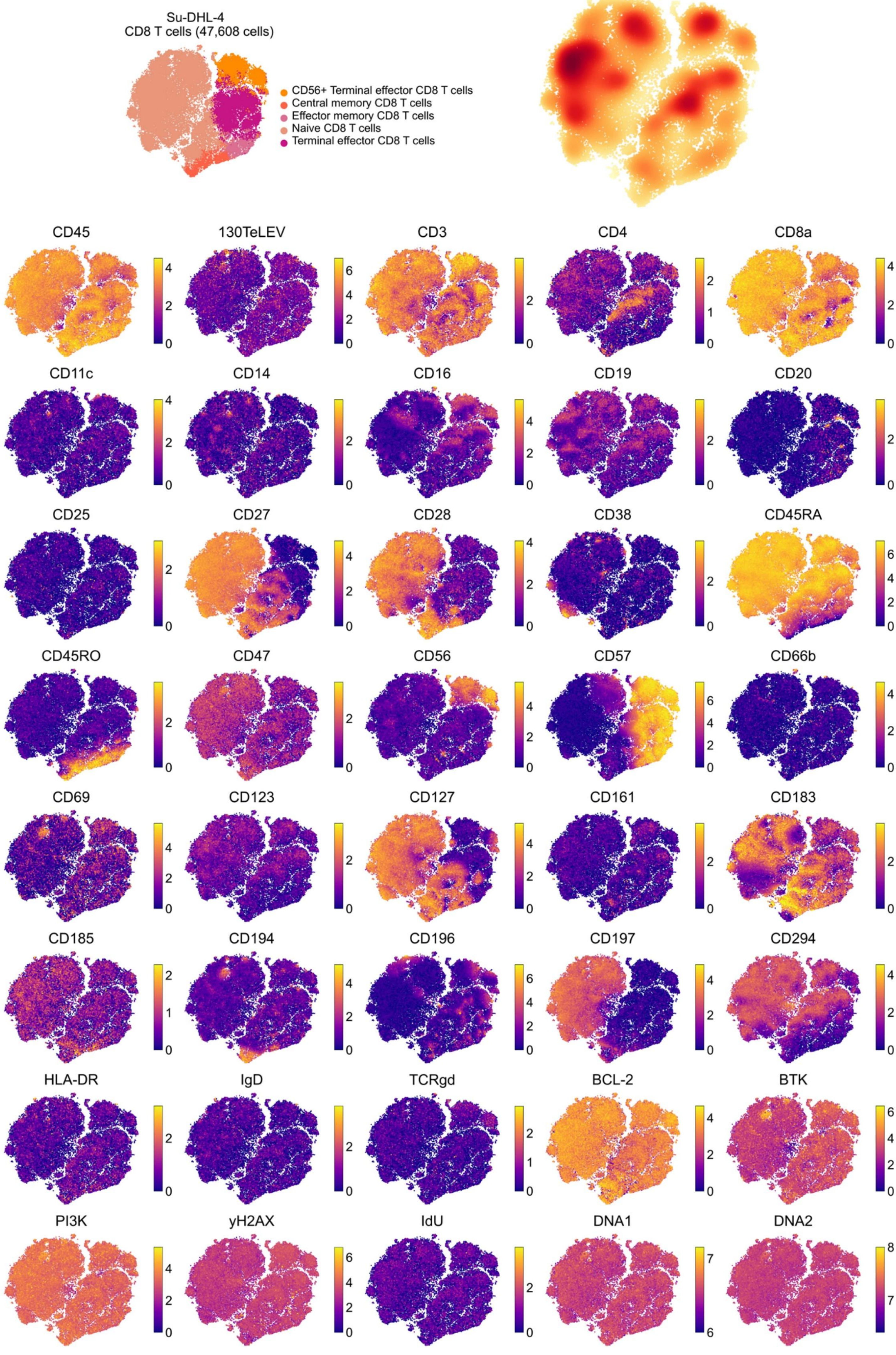

Su-DHL-4
CD8 T cells (47,608 cells)
CD56+ Terminal effector CD8 T cells
Central memory CD8 T cells
Effector memory CD8 T cells
Naive CD8 T cells
Terminal effector CD8 T cells
CD45
130TeLEV
CD3
CD4
CD8a
CD11c
CD14
CD16
CD19
CD20
CD25
CD27
CD28
CD38
CD45RA
CD45RO
CD47
CD56
CD57
CD66b
CD69
CD123
CD127
CD161
CD183
CD185
CD194
CD196
CD197
CD294
HLA-DR
IgD
TCRgd
BCL-2
BTK
PI3K
yH2AX
IdU
DNA1
DNA2

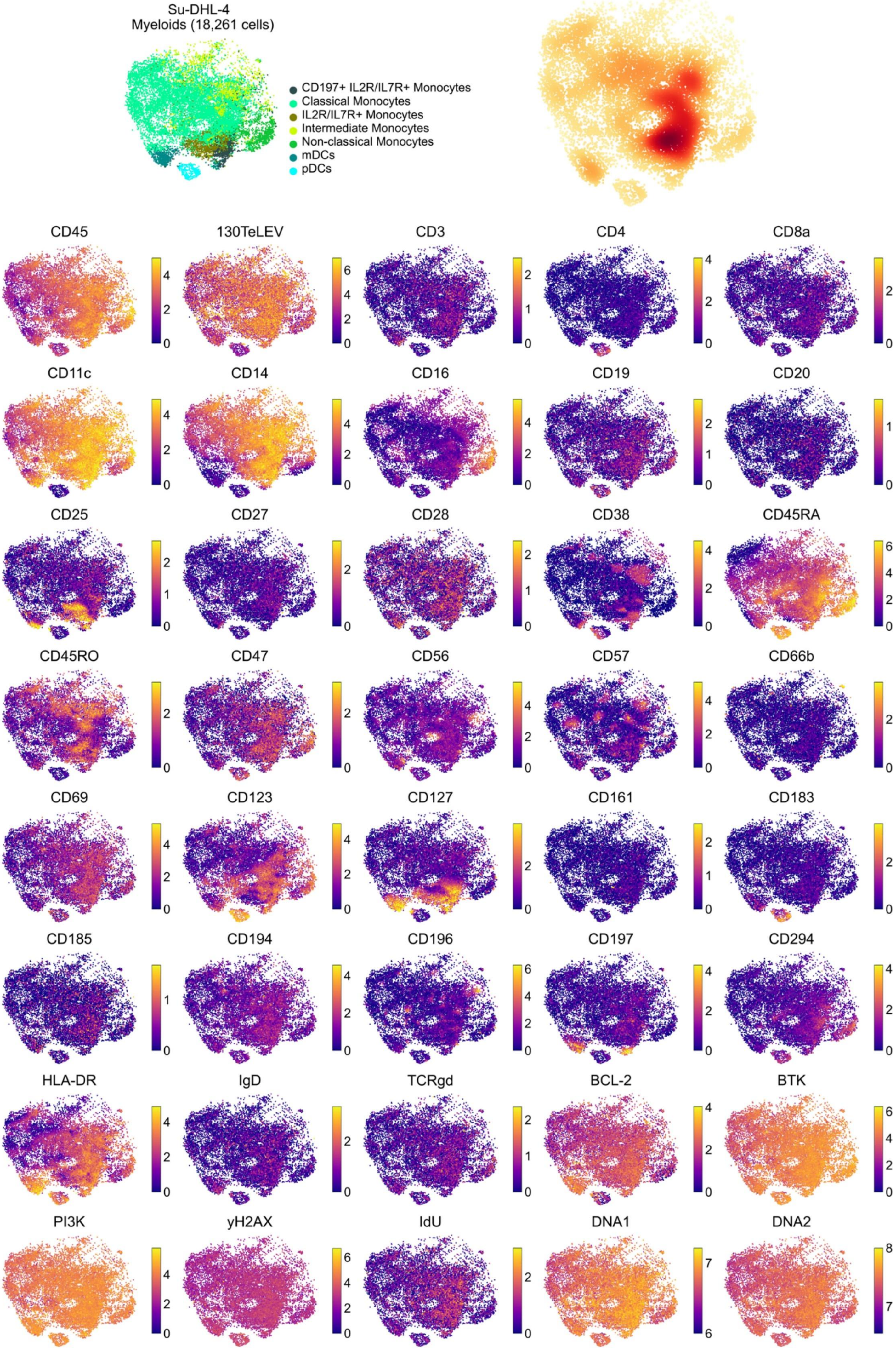
Su-DHL-4
Myeloids (18,261 cells)
CD197+ IL2R/IL7R+ Monocytes
Classical Monocytes
IL2R/IL7R+ Monocytes
Intermediate Monocytes
Non-classical Monocytes
mDCs
pDCs
CD45
130TeLEV
CD3
CD4
CD8a
CD11c
CD14
CD16
CD19
CD20
CD25
CD27
CD28
CD38
CD45RA
CD45RO
CD47
CD56
CD57
CD66b
CD69
CD123
CD127
CD161
CD183
CD185
CD194
CD196
CD197
CD294
HLA-DR
IgD
TCRgd
BCL-2
BTK
PI3K
yH2AX
IdU
DNA1
DNA2

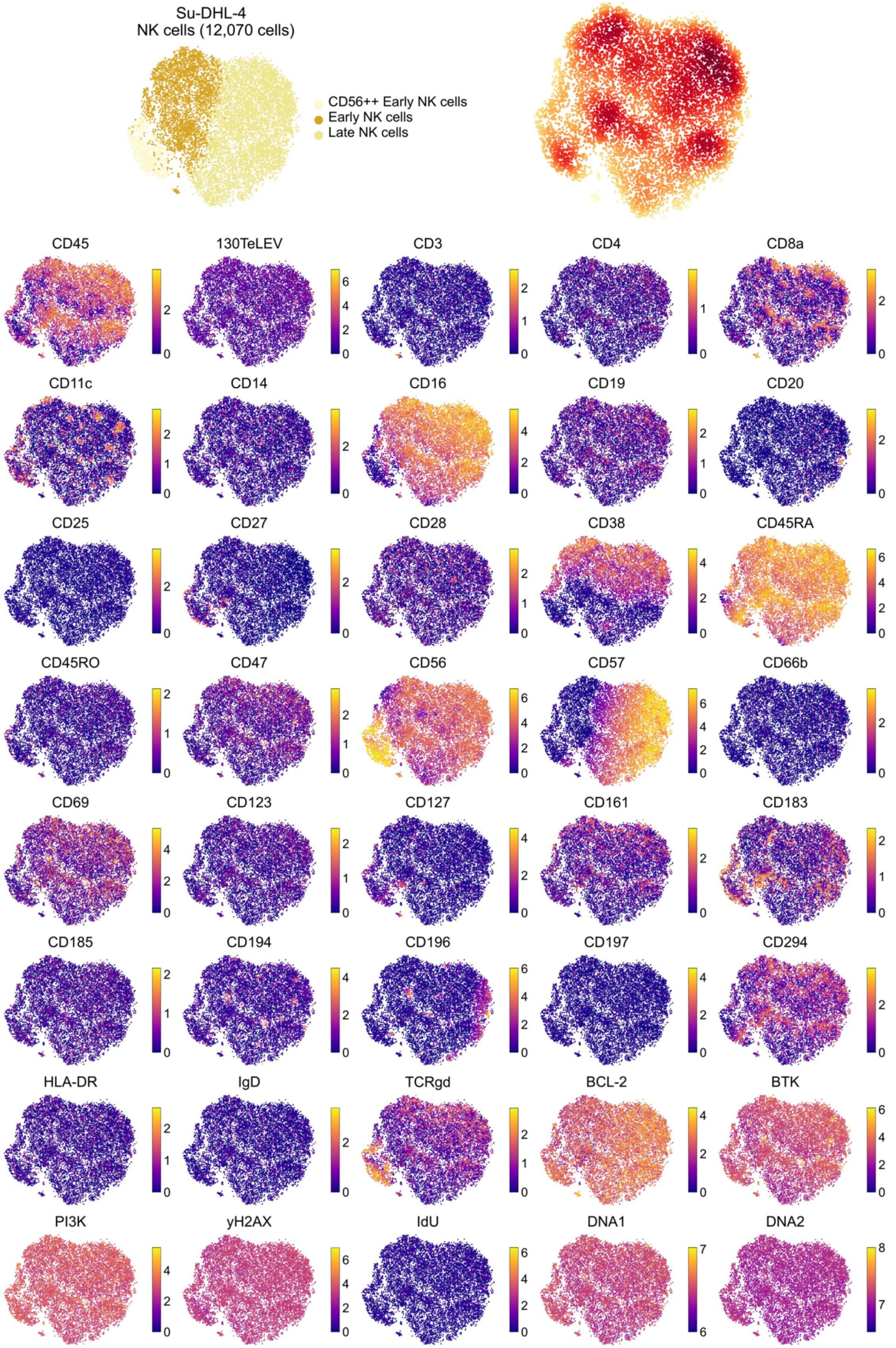
Su-DHL-4
NK cells (12,070 cells)
CD56++ Early NK cells
Early NK cells
Late NK cells
CD45
130TeLEV
CD3
CD4
CD8a
CD11c
CD14
CD16
CD19
CD20
CD25
CD27
CD28
CD38
CD45RA
CD45RO
CD47
CD56
CD57
CD66b
CD69
CD123
CD127
CD161
CD183
CD185
CD194
CD196
CD197
CD294
HLA-DR
IgD
TCRgd
BCL-2
BTK
PI3K
yH2AX
IdU
DNA1
DNA2

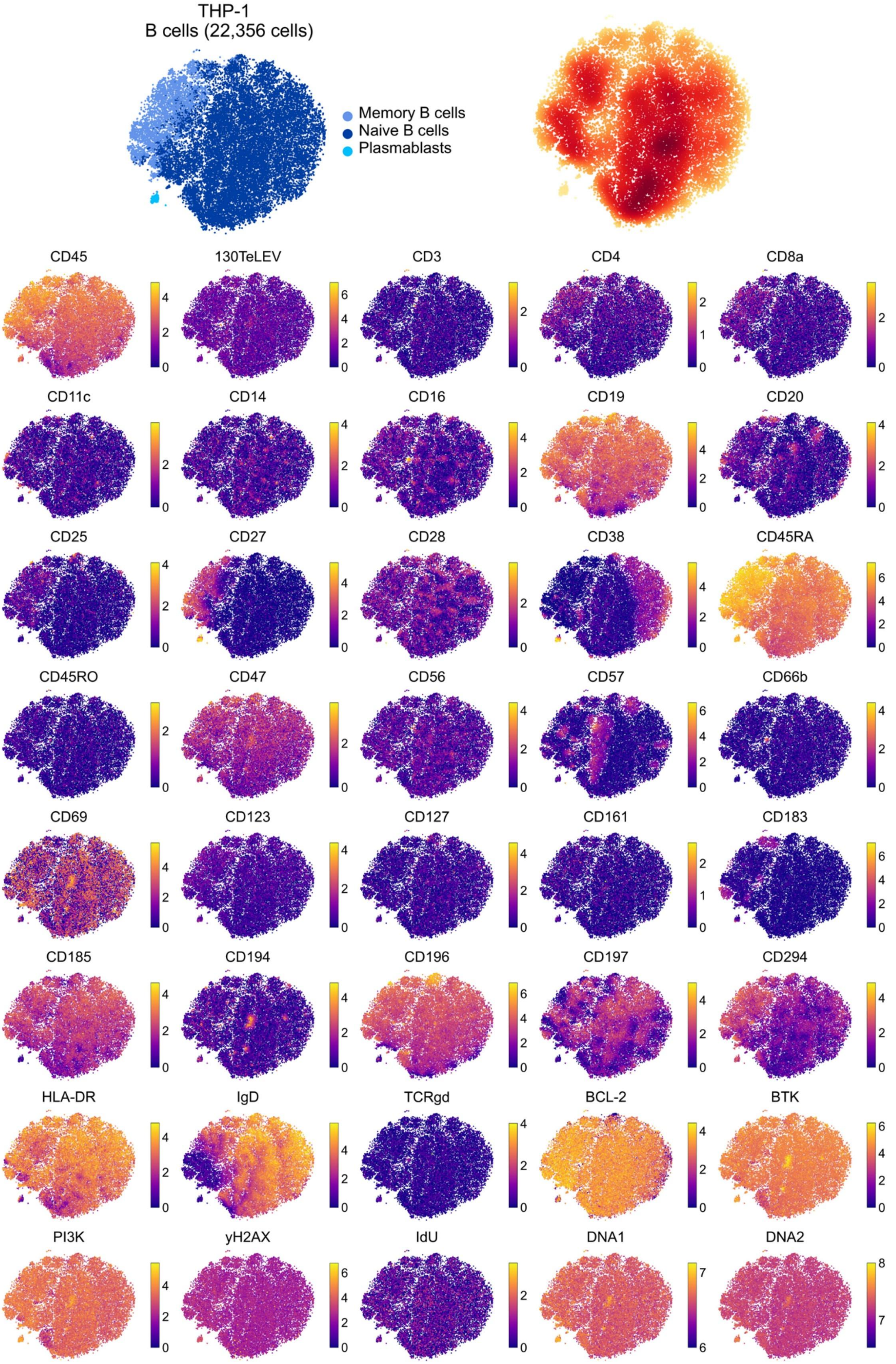
THP-1
B cells (22,356 cells)
Memory B cells
Naive B cells
Plasmablasts
CD45
130TeLEV
CD3
CD4
CD8a
CD11c
CD14
CD16
CD19
CD20
CD25
CD27
CD28
CD38
CD45RA
CD45RO
CD47
CD56
CD57
CD66b
CD69
CD123
CD127
CD161
CD183
CD185
CD194
CD196
CD197
CD294
HLA-DR
IgD
TCRgd
BCL-2
BTK
PI3K
yH2AX
IdU
DNA1
DNA2

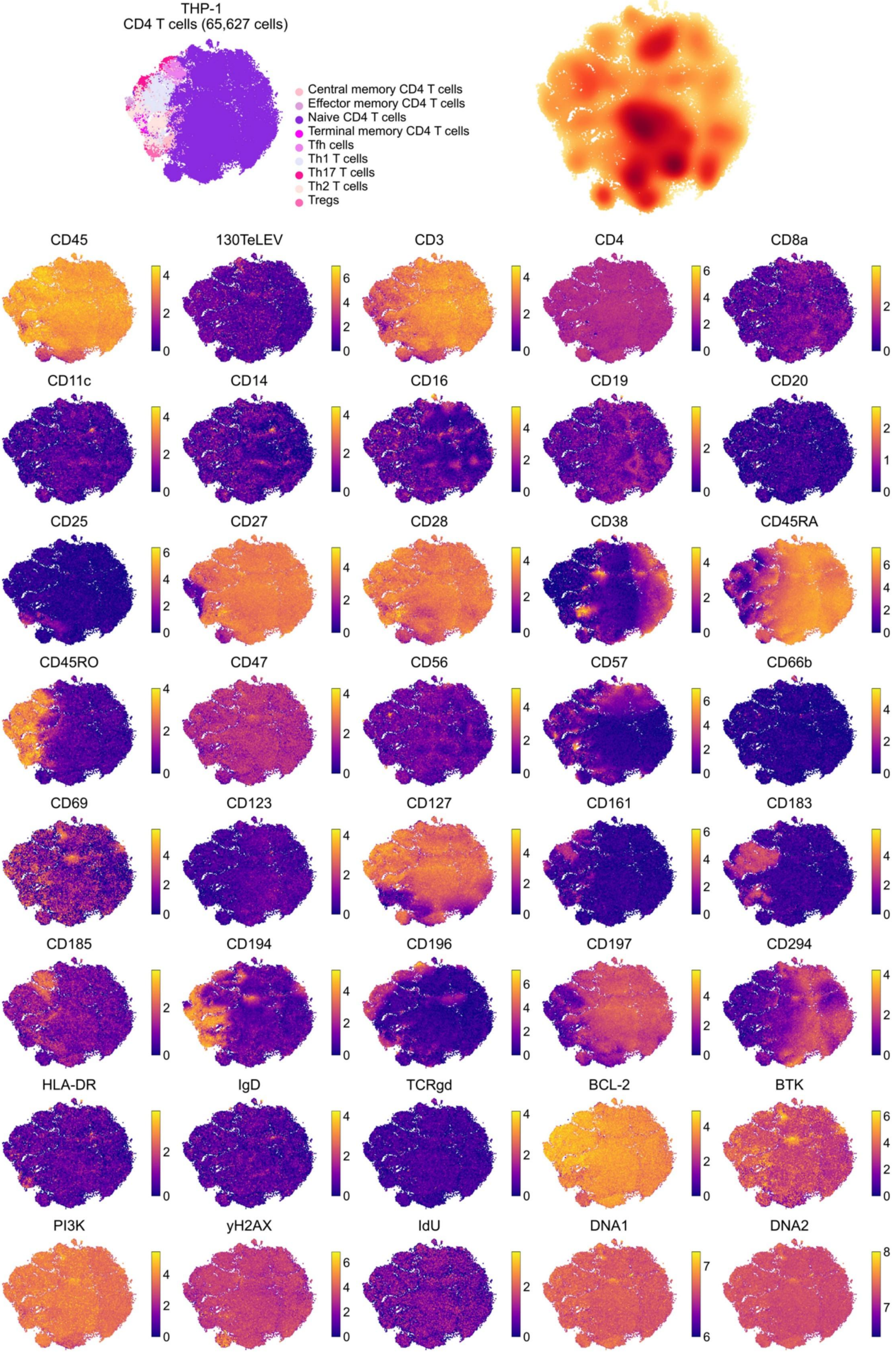

THP-1
CD4 T cells (65,627 cells)
Central memory CD4 T cells
Effector memory CD4 T cells
Naive CD4 T cells
Terminal memory CD4 T cells
Tfh cells
Th1 T cells
Th17 T cells
Th2 T cells
Tregs
CD45
130TeLEV
CD3
CD4
CD8a
CD11c
CD14
CD16
CD19
CD20
CD25
CD27
CD28
CD38
CD45RA
CD45RO
CD47
CD56
CD57
CD66b
CD69
CD123
CD127
CD161
CD183
CD185
CD194
CD196
CD197
CD294
HLA-DR
IgD
TCRgd
BCL-2
BTK
PI3K
yH2AX
IdU
DNA1
DNA2

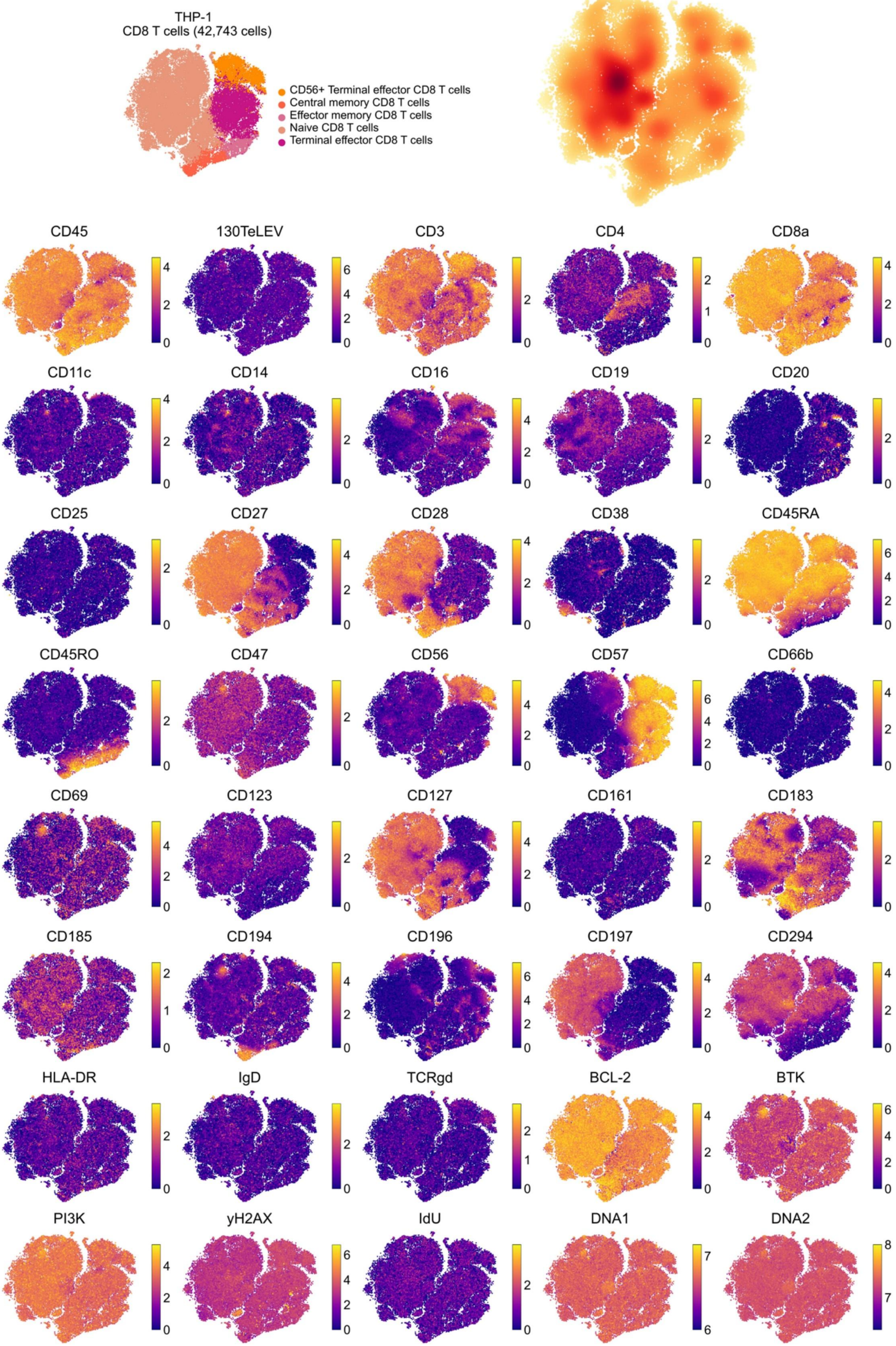
THP-1
CD8 T cells (42,743 cells)
CD56+ Terminal effector CD8 T cells
Central memory CD8 T cells
Effector memory CD8 T cells
Naive CD8 T cells
Terminal effector CD8 T cells
CD45
130TeLEV
CD3
CD4
CD8a
CD11c
CD14
CD16
CD19
CD20
CD25
CD27
CD28
CD38
CD45RA
CD45RO
CD47
CD56
CD57
CD66b
CD69
CD123
CD127
CD161
CD183
CD185
CD194
CD196
CD197
CD294
HLA-DR
IgD
TCRgd
BCL-2
BTK
PI3K
yH2AX
IdU
DNA1
DNA2

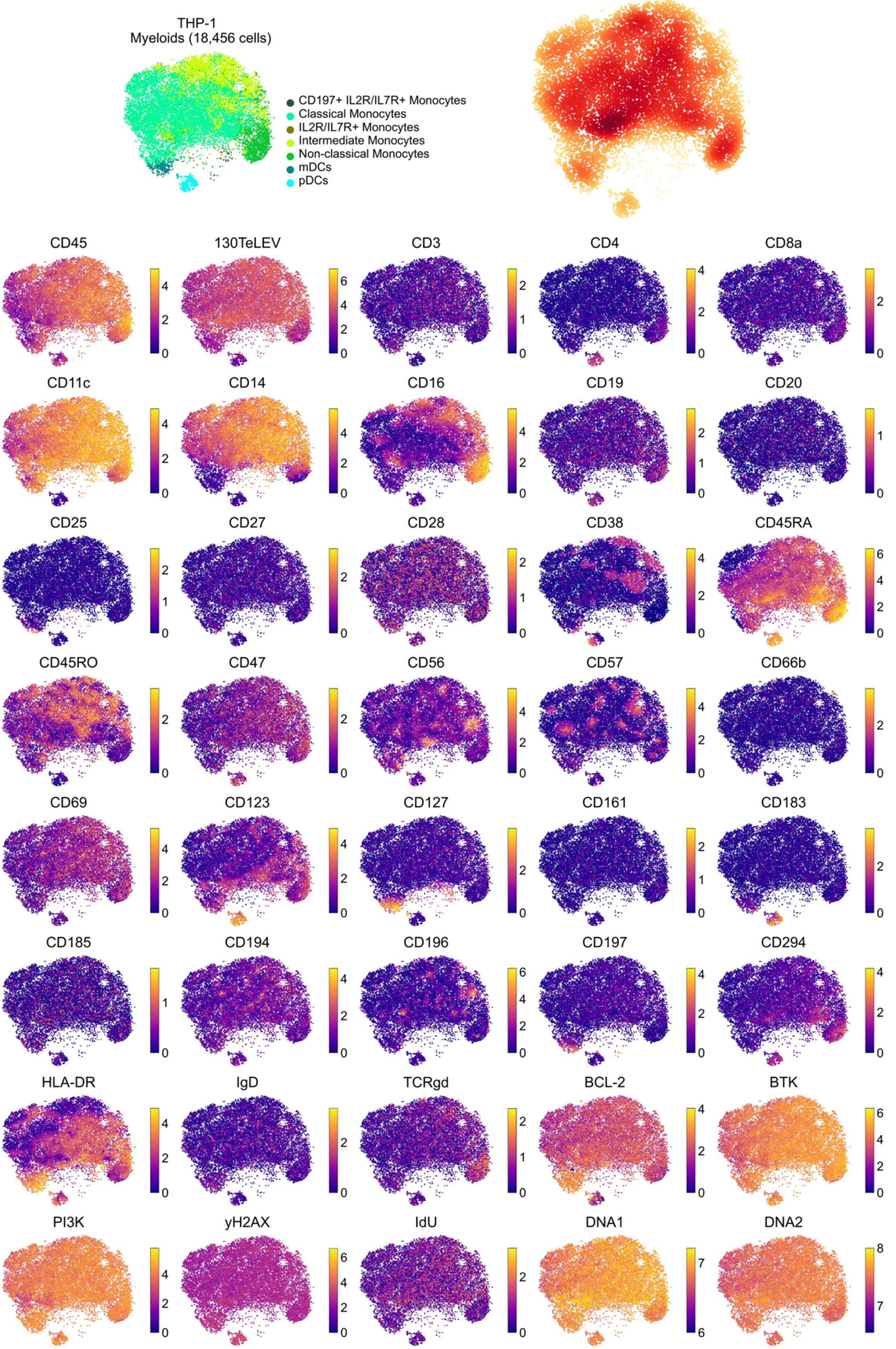
THP-1
Myeloids (18,456 cells)
CD197+ IL2R/IL7R+ Monocytes
Classical Monocytes
IL2R/IL7R+ Monocytes
Intermediate Monocytes
Non-classical Monocytes
mDCs
pDCs
CD45
130TeLEV
CD3
CD4
CD8a
CD11c
CD14
CD16
CD19
CD20
CD25
CD27
CD28
CD38
CD45RA
CD45RO
CD47
CD56
CD57
CD66b
CD69
CD123
CD127
CD161
CD183
CD185
CD194
CD196
CD197
CD294
HLA-DR
IgD
TCRgd
BCL-2
BTK
PI3K
yH2AX
IdU
DNA1
DNA2

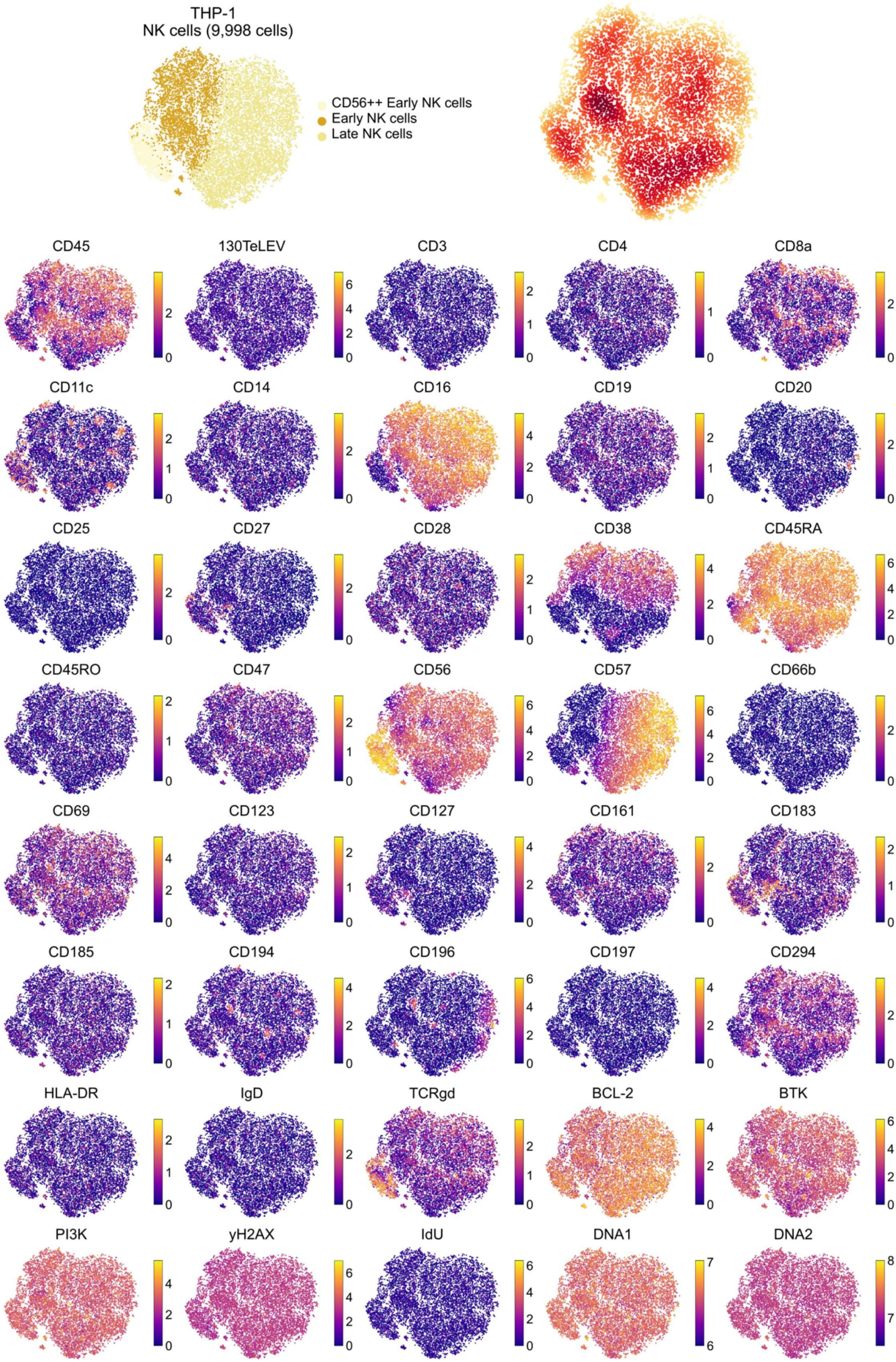
THP-1
NK cells (9,998 cells)
CD56++ Early NK cells
Early NK cells
Late NK cells
CD45
130TeLEV
CD3
CD4
CD8a
CD11c
CD14
CD16
CD19
CD20
CD25
CD27
CD28
CD38
CD45RA
CD45RO
CD47
CD56
CD57
CD66b
CD69
CD123
CD127
CD161
CD183
CD185
CD194
CD196
CD197
CD294
HLA-DR
IgD
TCRgd
BCL-2
BTK
PI3K
yH2AX
IdU
DNA1
DNA2

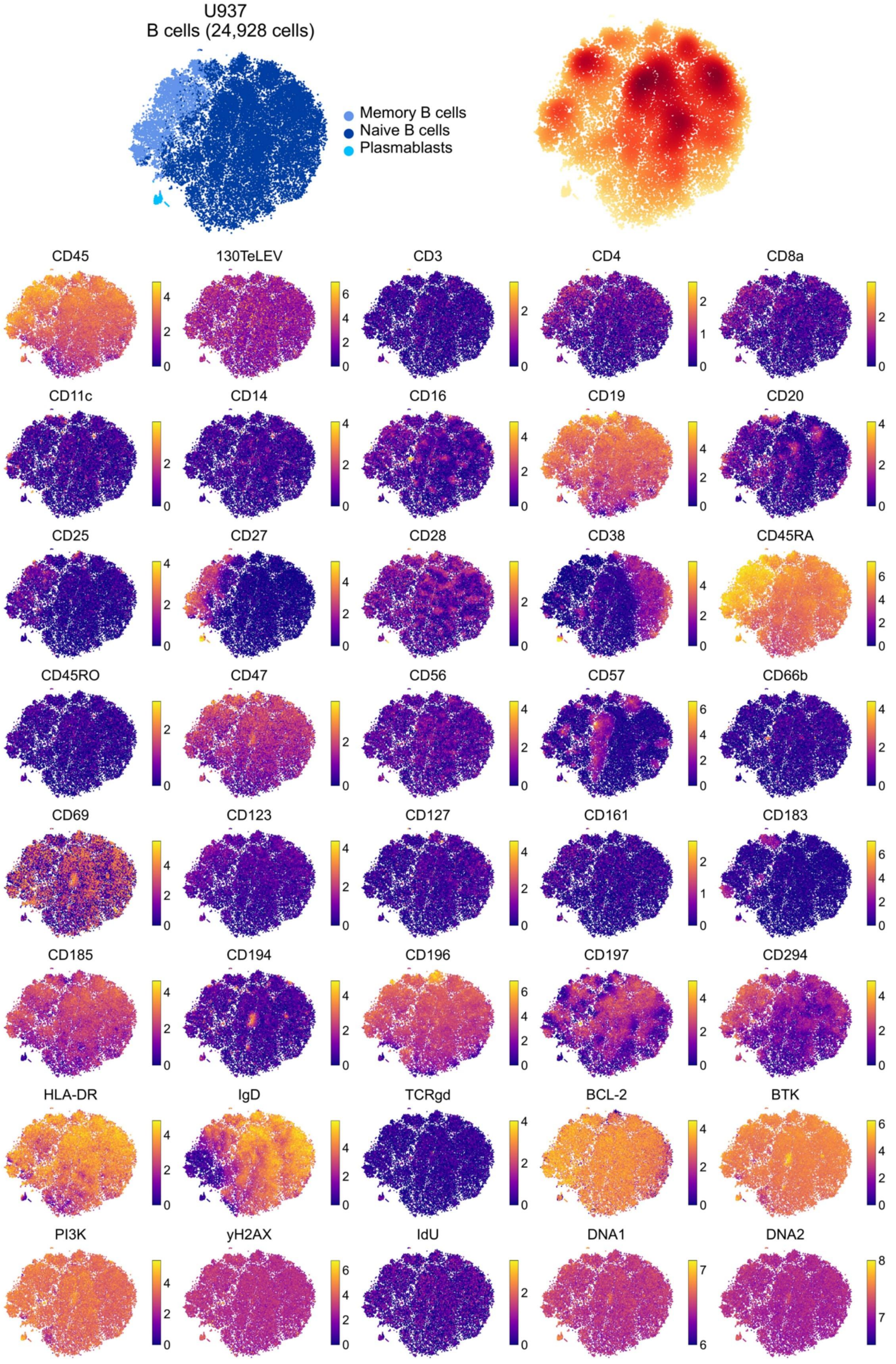
U937
B cells (24,928 cells)
Memory B cells
Naive B cells
Plasmablasts
CD45
130TeLEV
CD3
CD4
CD8a
CD11c
CD14
CD16
CD19
CD20
CD25
CD27
CD28
CD38
CD45RA
CD45RO
CD47
CD56
CD57
CD66b
CD69
CD123
CD127
CD161
CD183
CD185
CD194
CD196
CD197
CD294
HLA-DR
IgD
TCRgd
BCL-2
BTK
PI3K
yH2AX
IdU
DNA1
DNA2

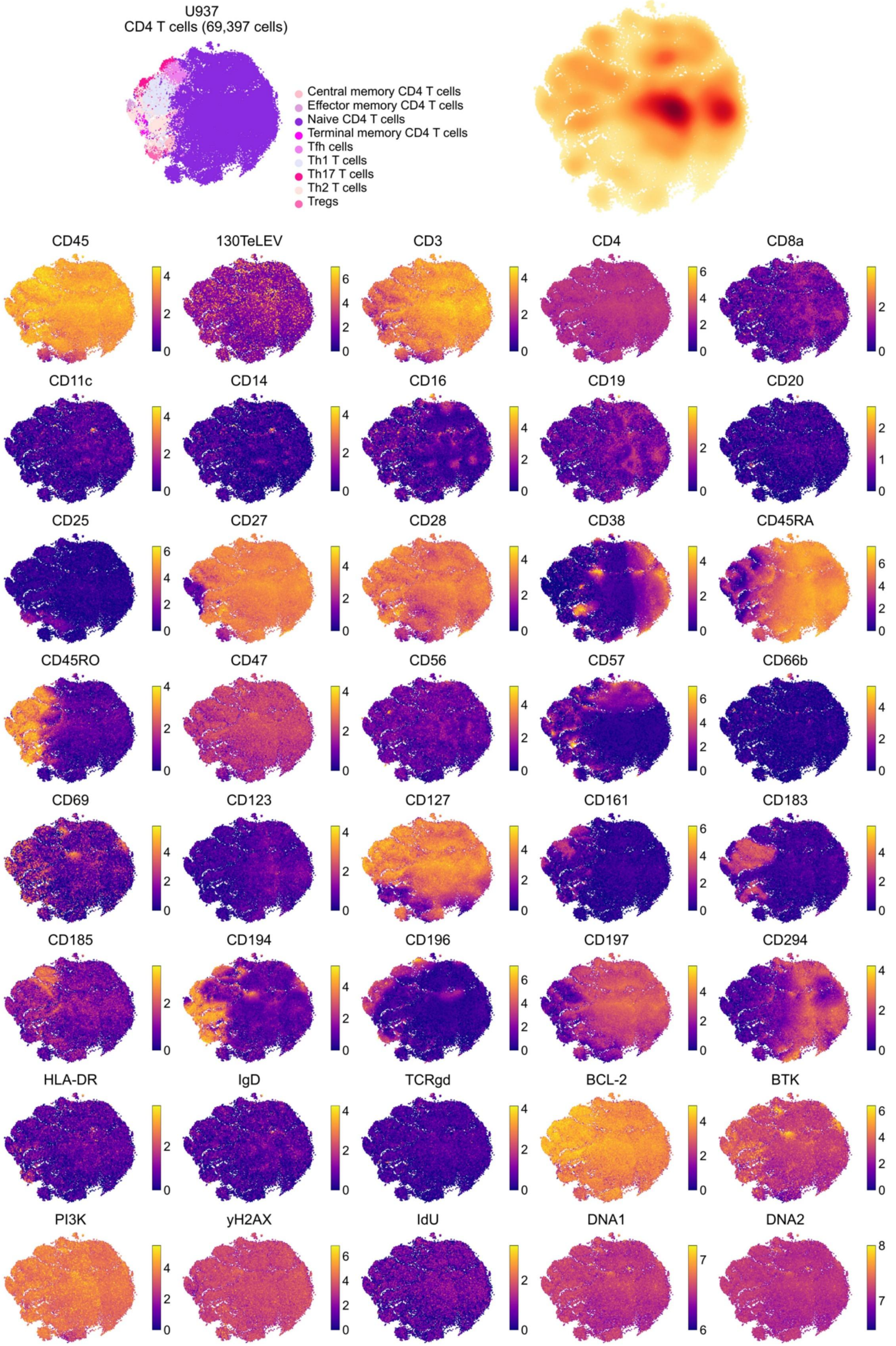
U937
CD4 T cells (69,397 cells)
Central memory CD4 T cells
Effector memory CD4 T cells
Naive CD4 T cells
Terminal memory CD4 T cells
Tfh cells
Th1 T cells
Th17 T cells
Th2 T cells
Tregs
CD45
130TeLEV
CD3
CD4
CD8a
CD11c
CD14
CD16
CD19
CD20
CD25
CD27
CD28
CD38
CD45RA
CD45RO
CD47
CD56
CD57
CD66b
CD69
CD123
CD127
CD161
CD183
CD185
CD194
CD196
CD197
CD294
HLA-DR
IgD
TCRgd
BCL-2
BTK
PI3K
yH2AX
IdU
DNA1
DNA2

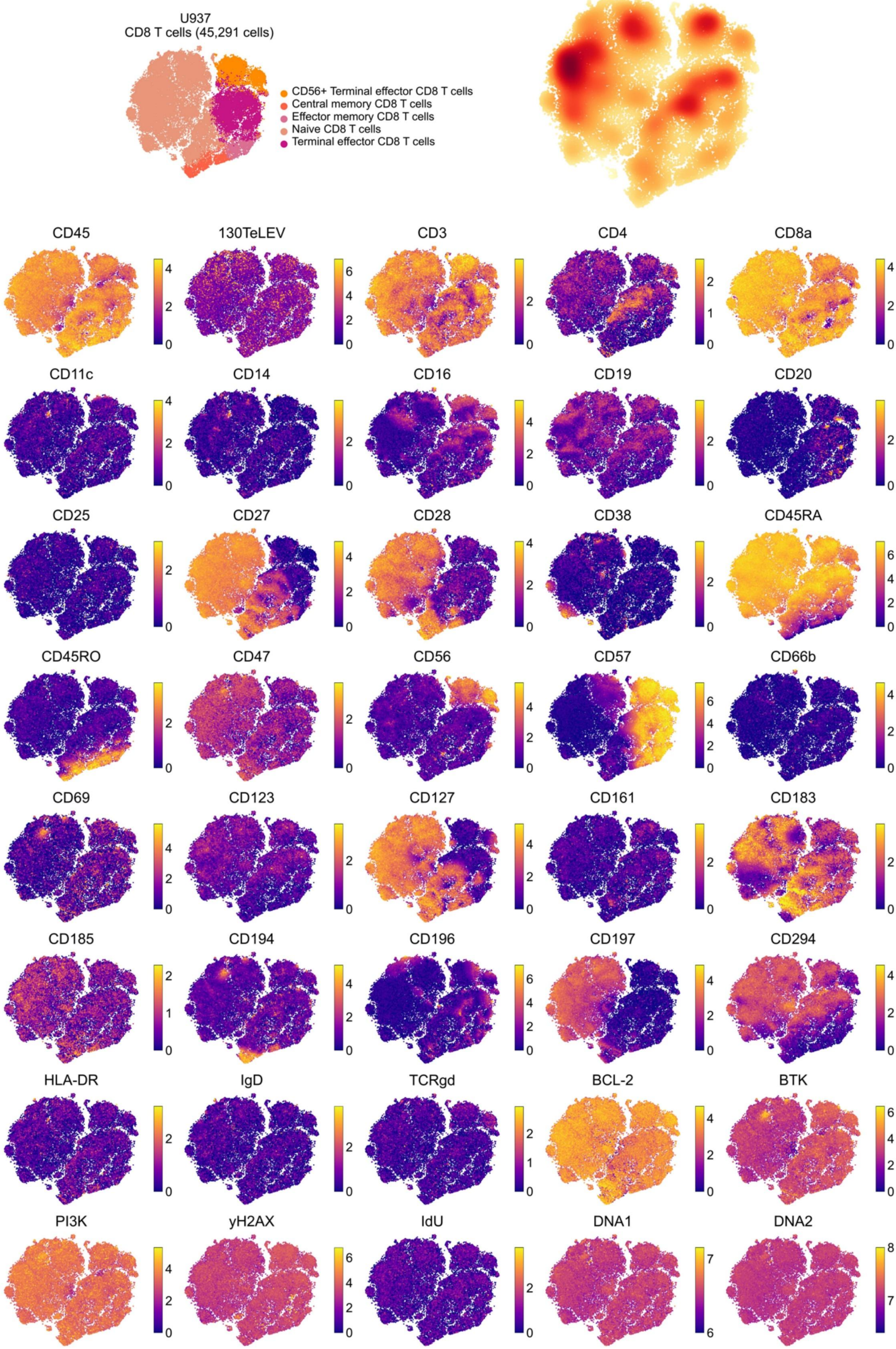

U937
CD8 T cells (45,291 cells)
CD56+ Terminal effector CD8 T cells
Central memory CD8 T cells
Effector memory CD8 T cells
Naive CD8 T cells
Terminal effector CD8 T cells
CD45
130TeLEV
CD3
CD4
CD8a
CD11c
CD14
CD16
CD19
CD20
CD25
CD27
CD28
CD38
CD45RA
CD45RO
CD47
CD56
CD57
CD66b
CD69
CD123
CD127
CD161
CD183
CD185
CD194
CD196
CD197
CD294
HLA-DR
IgD
TCRgd
BCL-2
BTK
PI3K
yH2AX
IdU
DNA1
DNA2

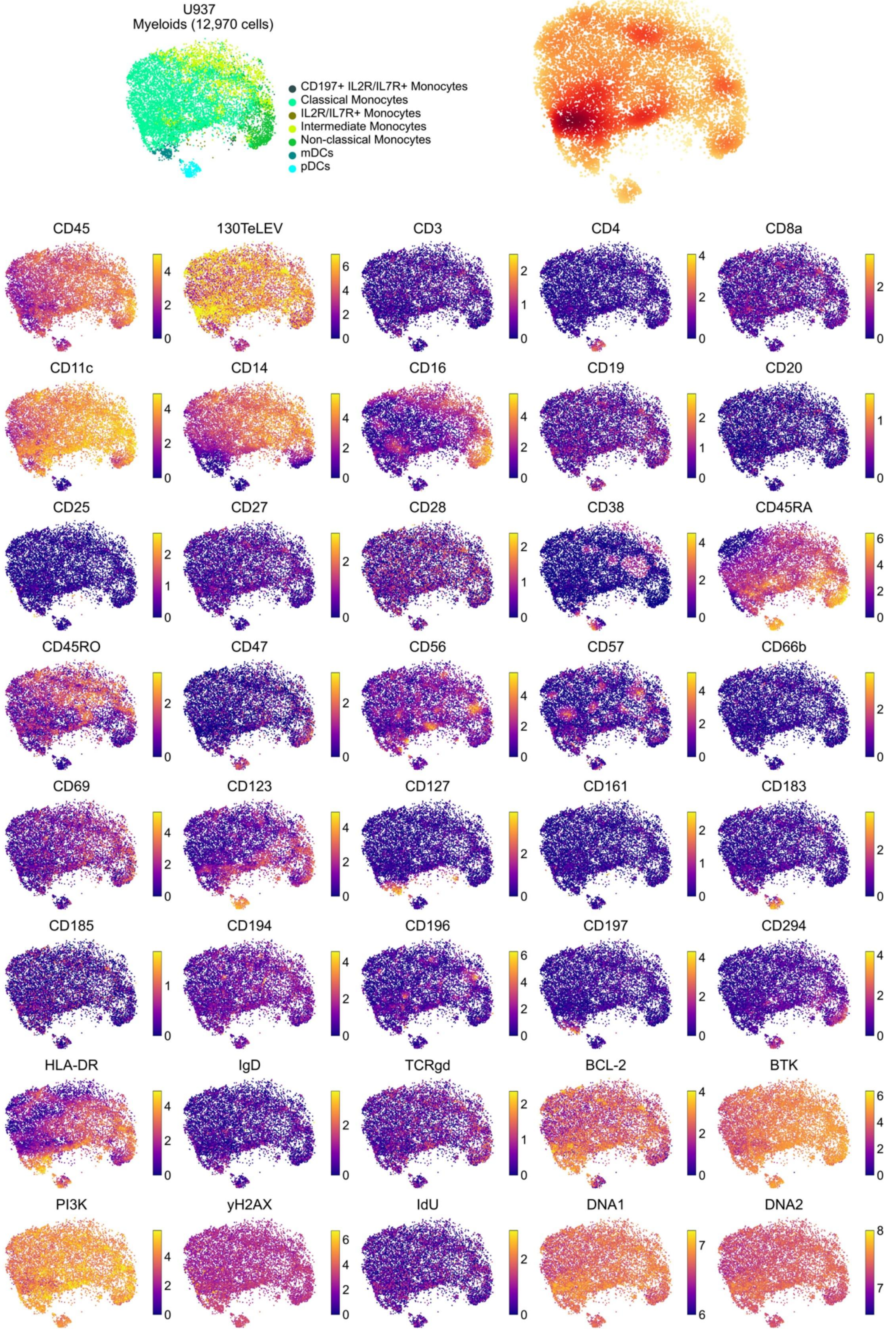
U937
Myeloids (12,970 cells)
CD197+ IL2R/IL7R+ Monocytes
Classical Monocytes
IL2R/IL7R+ Monocytes
Intermediate Monocytes
Non-classical Monocytes
mDCs
pDCs
CD45
130TeLEV
CD3
CD4
CD8a
CD11c
CD14
CD16
CD19
CD20
CD25
CD27
CD28
CD38
CD45RA
CD45RO
CD47
CD56
CD57
CD66b
CD69
CD123
CD127
CD161
CD183
CD185
CD194
CD196
CD197
CD294
HLA-DR
IgD
TCRgd
BCL-2
BTK
PI3K
yH2AX
IdU
DNA1
DNA2

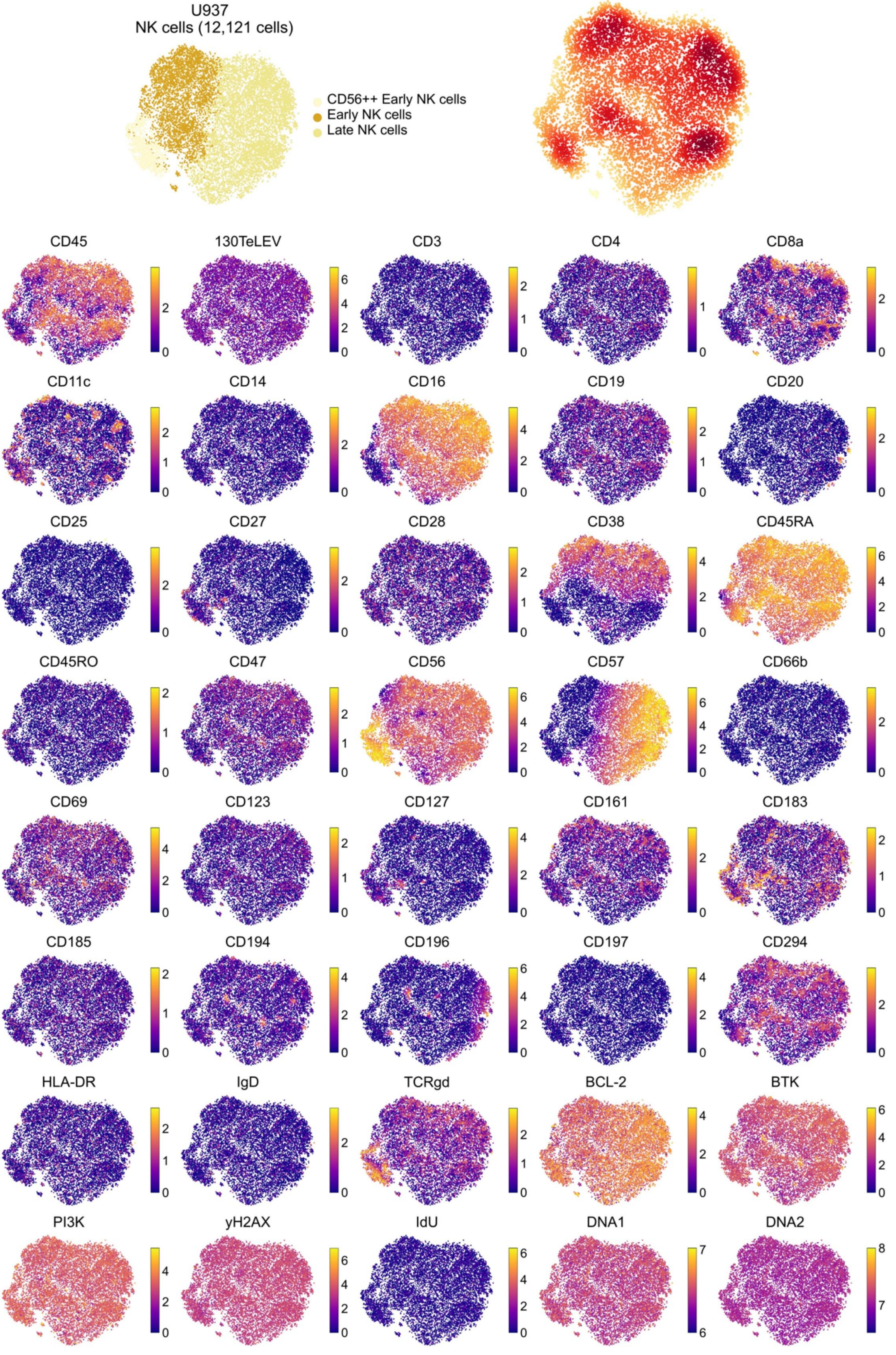

U937
NK cells (12,121 cells)
CD56++ Early NK cells
Early NK cells
Late NK cells
CD45
130TeLEV
CD3
CD4
CD8a
CD11c
CD14
CD16
CD19
CD20
CD25
CD27
CD28
CD38
CD45RA
CD45RO
CD47
CD56
CD57
CD66b
CD69
CD123
CD127
CD161
CD183
CD185
CD194
CD196
CD197
CD294
HLA-DR
IgD
TCRgd
BCL-2
BTK
PI3K
yH2AX
IdU
DNA1
DNA2

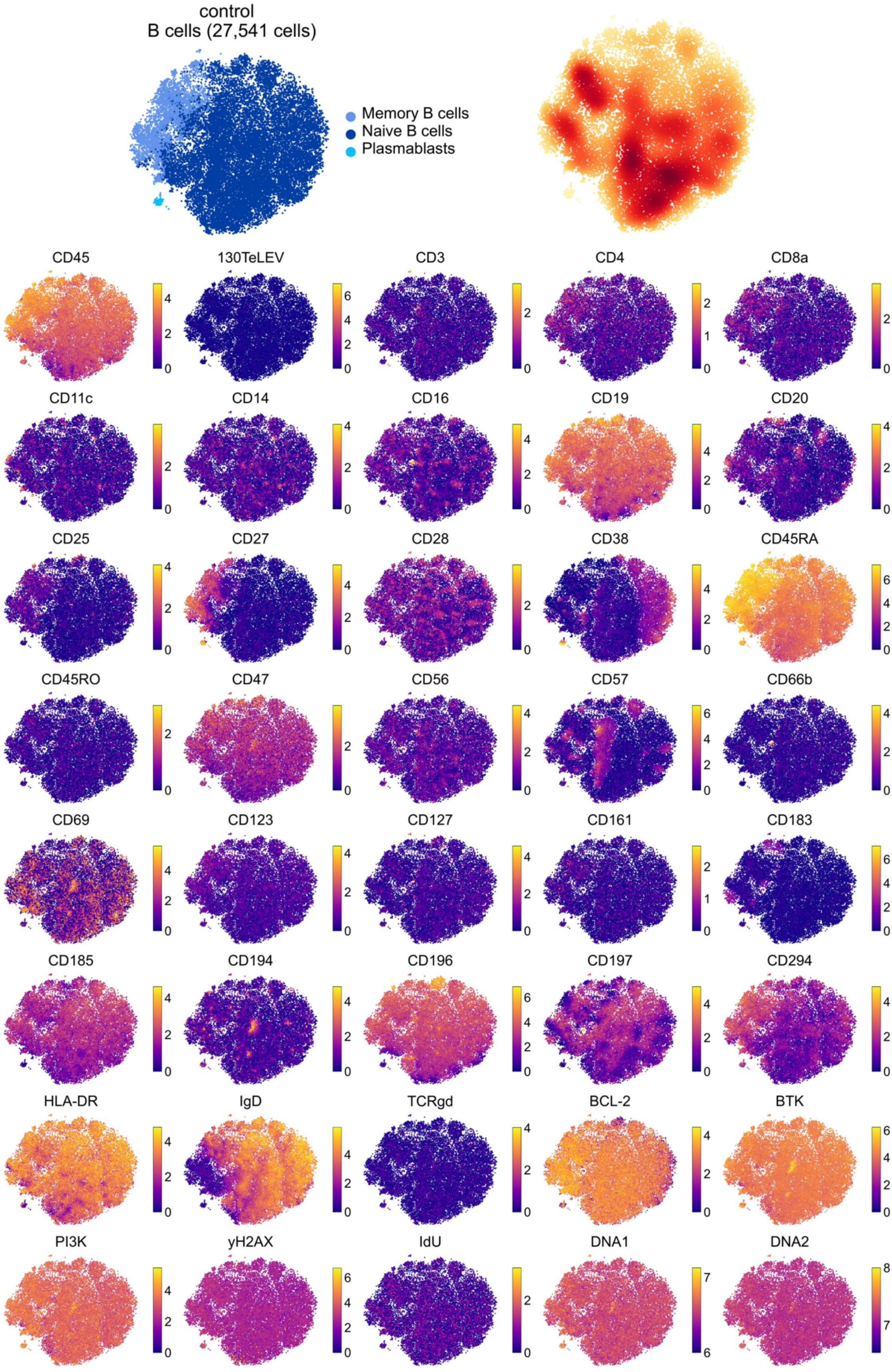
control
B cells (27,541 cells)
Memory B cells
Naive B cells
Plasmablasts
CD45
130TeLEV
CD3
CD4
CD8a
CD11c
CD14
CD16
CD19
CD20
CD25
CD27
CD28
CD38
CD45RA
CD45RO
CD47
CD56
CD57
CD66b
CD69
CD123
CD127
CD161
CD183
CD185
CD194
CD196
CD197
CD294
HLA-DR
IgD
TCRgd
BCL-2
BTK
PI3K
yH2AX
IdU
DNA1
DNA2

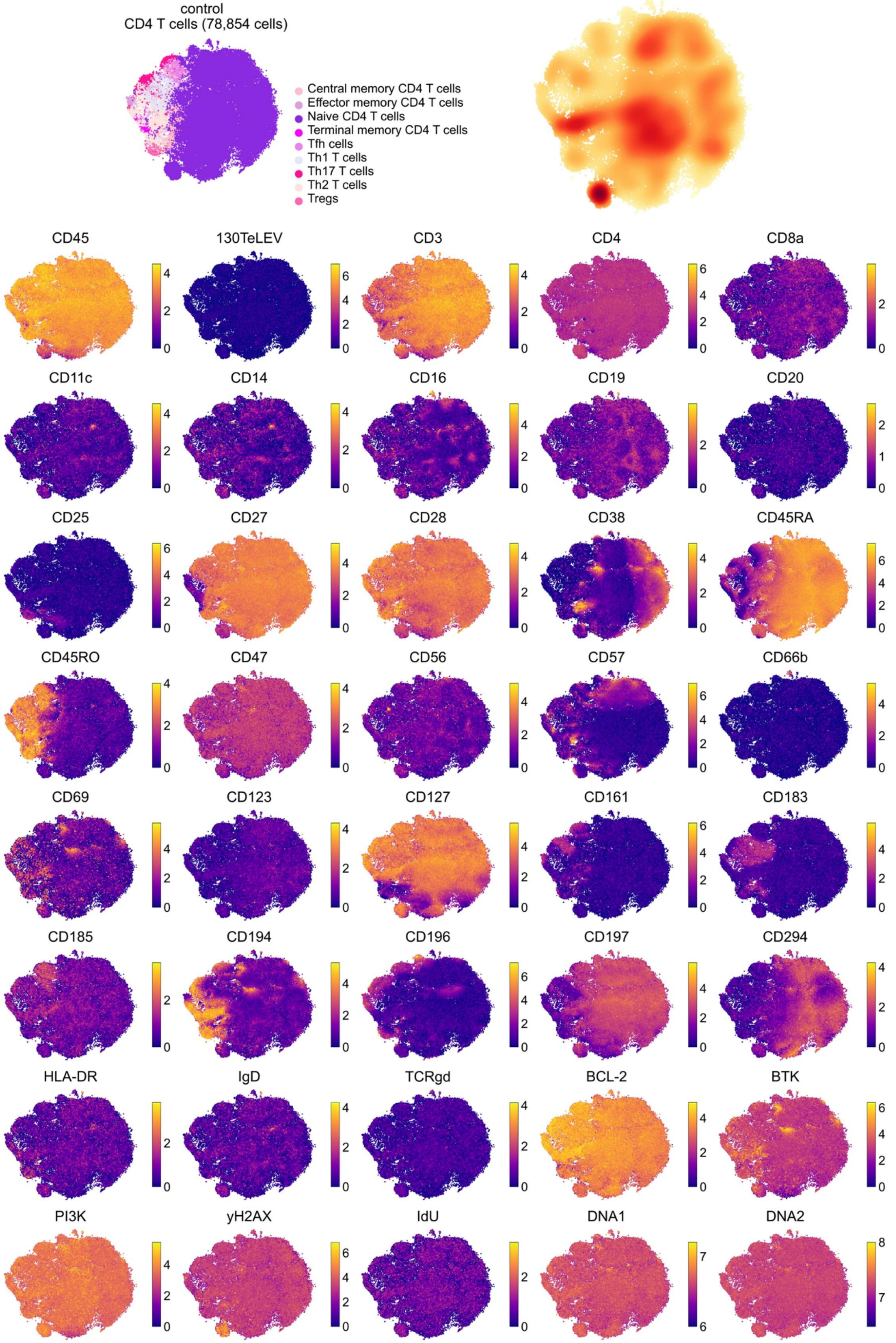
control
CD4 T cells (78,854 cells)
Central memory CD4 T cells
Effector memory CD4 T cells
Naive CD4 T cells
Terminal memory CD4 T cells
Tfh cells
Th1 T cells
Th17 T cells
Th2 T cells
Tregs
CD45
130TeLEV
CD3
CD4
CD8a
CD11c
CD14
CD16
CD19
CD20
CD25
CD27
CD28
CD38
CD45RA
CD45RO
CD47
CD56
CD57
CD66b
CD69
CD123
CD127
CD161
CD183
CD185
CD194
CD196
CD197
CD294
HLA-DR
IgD
TCRgd
BCL-2
BTK
PI3K
yH2AX
IdU
DNA1
DNA2

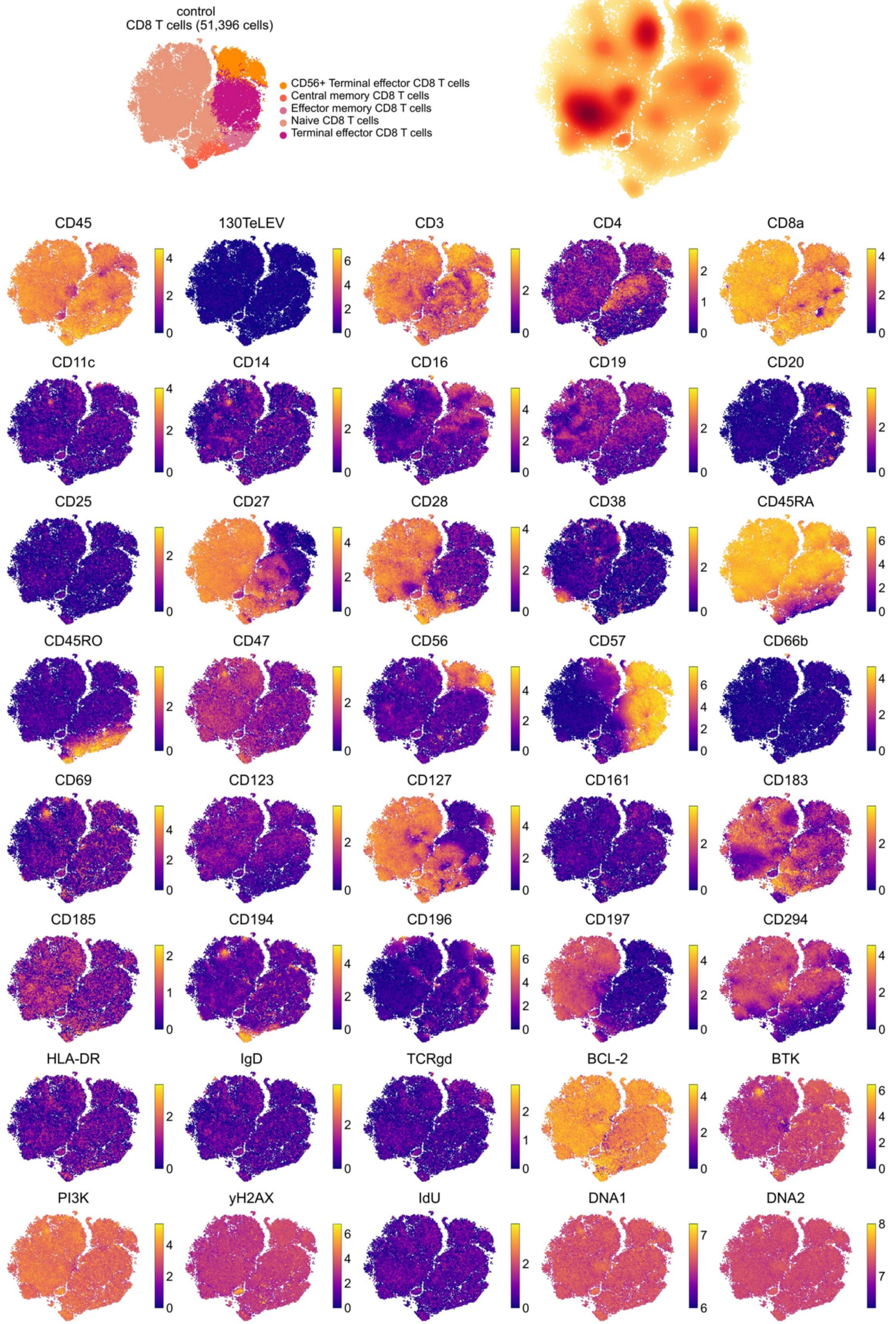
control
CD8 T cells (51,396 cells)
CD56+ Terminal effector CD8 T cells
Central memory CD8 T cells
Effector memory CD8 T cells
Naive CD8 T cells
Terminal effector CD8 T cells
CD45
130TeLEV
CD3
CD4
CD8a
CD11c
CD14
CD16
CD19
CD20
CD25
CD27
CD28
CD38
CD45RA
CD45RO
CD47
CD56
CD57
CD66b
CD69
CD123
CD127
CD161
CD183
CD185
CD194
CD196
CD197
CD294
HLA-DR
IgD
TCRgd
BCL-2
BTK
PI3K
yH2AX
IdU
DNA1
DNA2

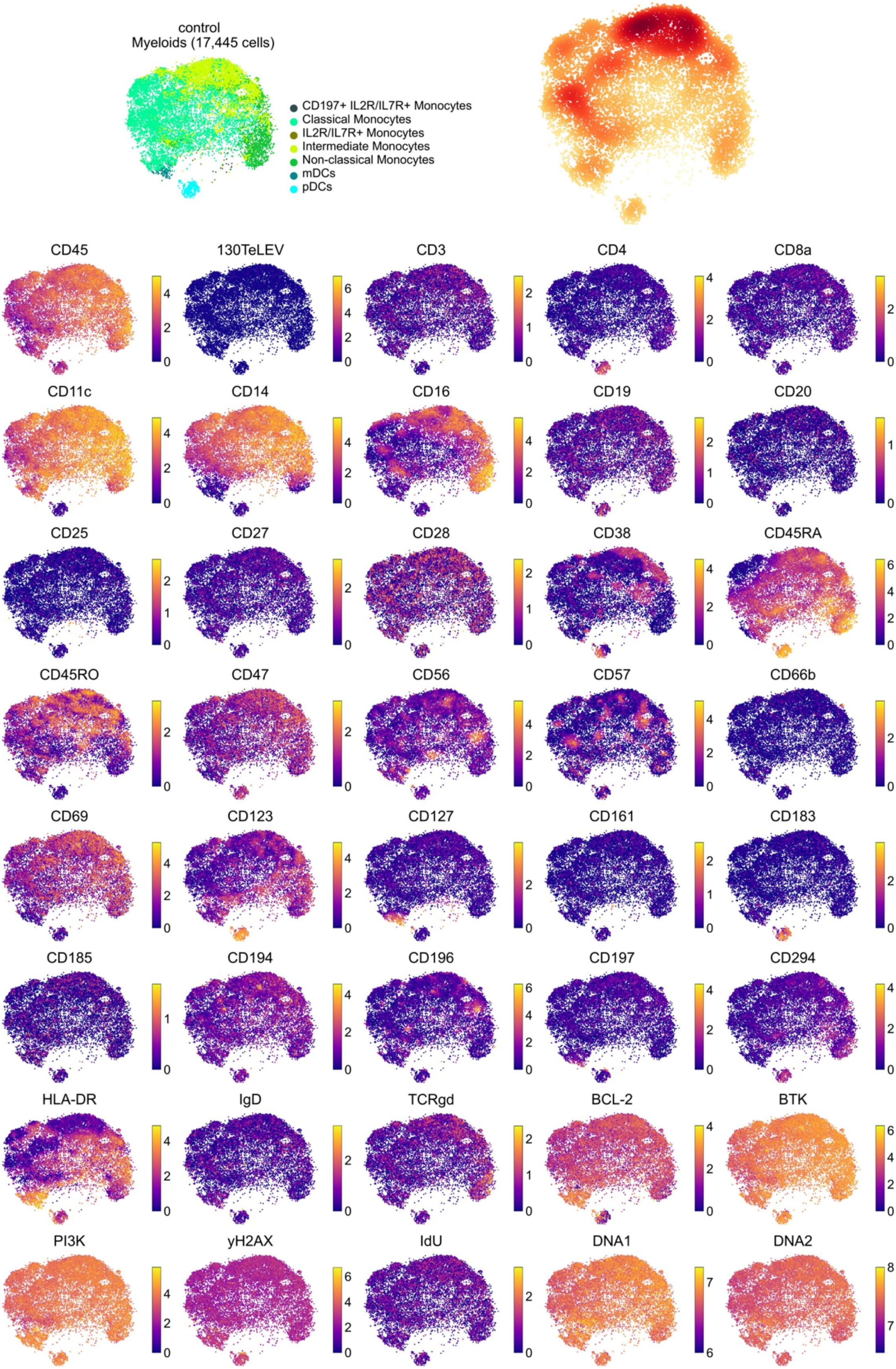

control
Myeloids (17,445 cells)
CD197+ IL2R/IL7R+ Monocytes
Classical Monocytes
IL2R/IL7R+ Monocytes
Intermediate Monocytes
Non-classical Monocytes
mDCs
pDCs
CD45
130TeLEV
CD3
CD4
CD8a
CD11c
CD14
CD16
CD19
CD20
CD25
CD27
CD28
CD38
CD45RA
CD45RO
CD47
CD56
CD57
CD66b
CD69
CD123
CD127
CD161
CD183
CD185
CD194
CD196
CD197
CD294
HLA-DR
IgD
TCRgd
BCL-2
BTK
PI3K
yH2AX
IdU
DNA1
DNA2

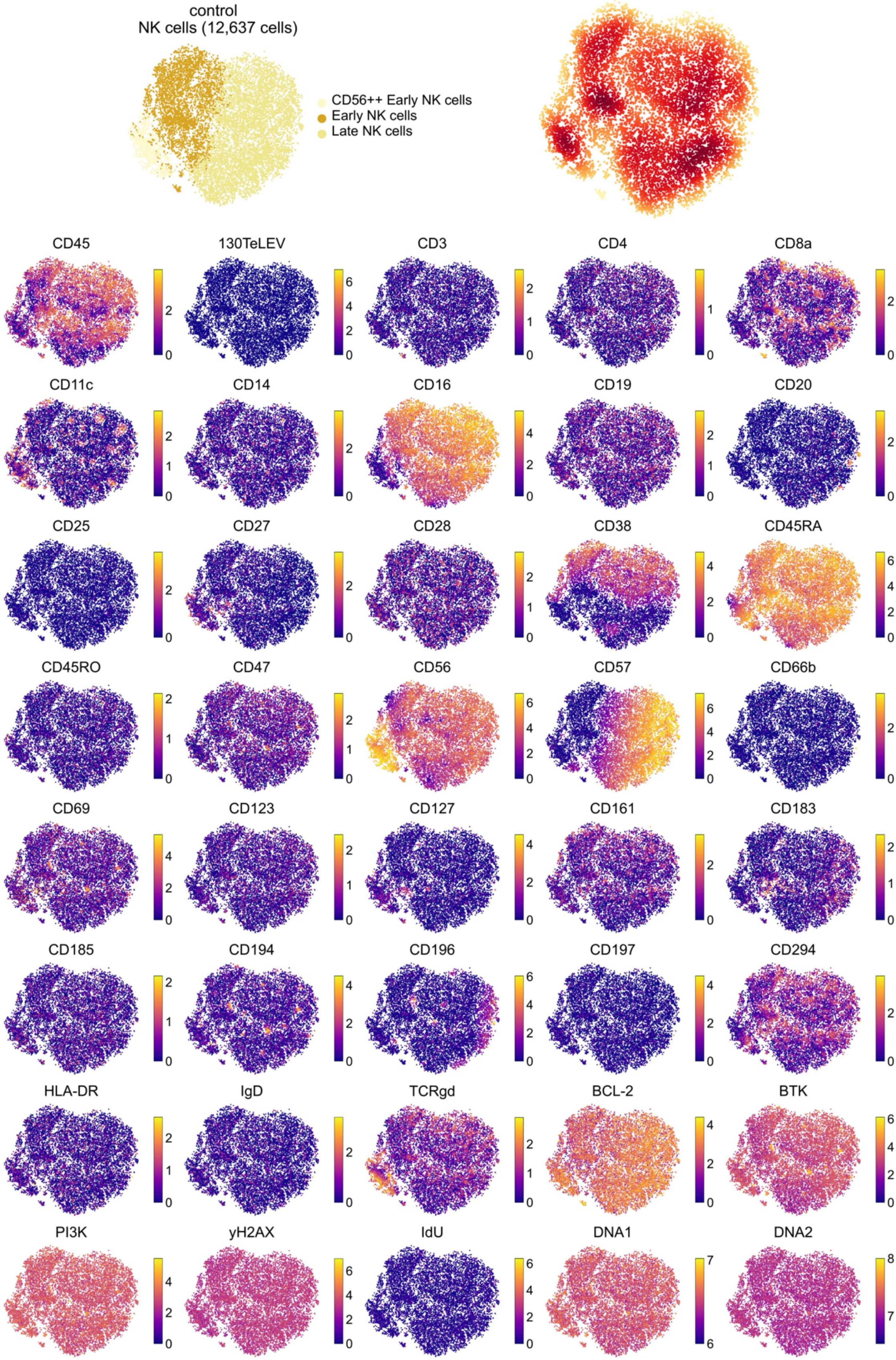
control
NK cells (12,637 cells)
CD56++ Early NK cells
Early NK cells
Late NK cells
CD45
130TeLEV
CD3
CD4
CD8a
CD11c
CD14
CD16
CD19
CD20
CD25
CD27
CD28
CD38
CD45RA
CD45RO
CD47
CD56
CD57
CD66b
CD69
CD123
CD127
CD161
CD183
CD185
CD194
CD196
CD197
CD294
HLA-DR
IgD
TCRgd
BCL-2
BTK
PI3K
yH2AX
IdU
DNA1
DNA2
